# Rectifying Stress into Structural Memory Drives Specialization in Cells and Adaptive Computational Systems

**DOI:** 10.64898/2026.01.29.702626

**Authors:** Remy Tuyeras, Alvaro Morcuende, Claudia Llinares, Asa Segerstolpe, Manolis Kellis, Teresa Femenia, Leandro Z. Agudelo

## Abstract

Functional specialization in continuous systems requires balancing adaptation to environmental stress with the preservation of encoded information. Yet, the physical constraints governing how living systems reconcile selective information retention with energetic structural reorganization remain unclear–a trade-off that may offer transferable design rules for adaptive computing. Here we present a unified framework that connects cellular specialization to adaptive machine learning through principles of nonequilibrium optimization. We find that asymmetric fluctuations in key parameters act as tunable drives that, when phase-aligned, are rectified into durable functional specialization. Biologically, we show that in adipose mesenchymal cells, transient nutrient deficits trigger cytosolic pH fluctuations. These oscillations, regulated by the ATP12A proton pump, modulate nuclear compartment dynamics to coordinate mitochondrial specialization and lock in epigenetic memory for phenotype persistence. Notably, modulating ATP12A amplifies these drives, enhancing functional specialization and metabolic homeostasis in disease models. Computationally, we formalize analogous principles using a compositional framework based on operadic algebra, where structural updates act as a topological ratchet converting drive fluctuations into persistent architectures. This reveals a learning regime where gains in predictive information are accompanied by auditable residual costs for non-predictive states and reorganization. As a proof of concept, we deploy a self-organizing machine learning system that discovers latent hierarchy and reconstructs data through adaptive compartment dynamics. We show that internal organization fluctuations increase with residuals, an information-processing ledger guiding adaptive computation. Across both domains, local organization gains are directionally linked to dissipation proxies (pH variance in vivo and residuals in silico), consistent with operation under shared constraints. Overall, this work identifies asymmetric-drive rectification as a fundamental mechanism of adaptive specialization, providing thermodynamically informed design rules for both bioengineering and adaptive AI.

**One-Sentence Summary:** Asymmetric-drive rectification is a fundamental mechanism by which cells and adaptive computational systems convert internal stress into structural memory, encoding functional specialization

## Introduction

Living systems face a fundamental challenge when optimizing function: how to specialize while conserving the ability to adapt to environmental changes. This challenge is prominent, for example, during metabolic stress, when cells need to reorganize their transcriptional machinery to improve fitness and preserve phenotypic flexibility. Therefore, understanding the physical principles behind this nonequilibrium specialization could profoundly impact medicine and machine learning (ML), yet a unified adaptive framework formalizing cross-domain principles remains elusive.

The pursuit of ML has long drawn inspiration from biology (*1–6*), primarily using the brain as a blueprint for computation and pattern recognition. However, the challenges of robust adaptation including managing uncertainty, continuous learning, and efficient self-organization, are not solved by these approaches alone. Biocomputing is an interdisciplinary field combining biology and computer science, where studying dynamic behavior within living systems can help identify operational rules of adaptation and information processing (*1–6*). For example, the cell and the organizational principles of the nucleus could provide the essential blueprint for adaptive architectures and the management of symbolic knowledge lifecycle (akin to DNA storage, regulation, and evolution). These insights can lead to innovative algorithms that enhance our understanding of complex behaviors and adaptive intelligence (*1–6*).

### Specialization under constraint: nuclear compartments as guiding scaffolds

A key step lies in formalizing the principles of cellular adaptive specialization. Cells are open nonequilibrium systems that exchange energy with their environment (*7–10*), while regulating specialization via the dynamic organization of nuclear compartments (*11–13*). This organization is facilitated by multi-scale features (*14*) including protein domains (DNA-binding domains; intrinsically disordered regions or IDRs), liquid-like phase separation (LLPS), and 3D genome architecture (*15, 16*). These transcriptional compartments coordinate gene regulation via several mechanisms, including transcription factor interactions (*15*), chromatin contacts (*16*), and epigenetic enzyme cooperation (*17–19*).

These nuclear compartments can also be fine-tuned by environmental stressors linked to low-energy states such as cold and fasting, which alter cellular volume (i.e., crowding), molecular interactions, and viscosity (*20–23*), thereby influencing compartment features and function (*24, 25*). Thus, multi-scale nuclear compartments integrate diverse components under variable environmental conditions to support cell specialization (*26*). Yet, how cells optimize compartment dynamics to achieve stable specialization while remaining adaptable is not clear.

### Epigenetic memory as a specialization readout

A critical challenge for this is reconciling how transient environmental perturbations induce persistent phenotypic states while maintaining flexibility. Phenotype conservation is influenced by different epi-transcriptional mechanisms such as epigenetic memory, which involves the maintenance of heritable modifications along the genome after developmental or environmental stress (*27–29*). Besides developmental and transgenerational memory (*27–29*), transcriptional memory (*30*) is associated with acute responses to stress (inflammation and metabolic stress (*31, 32*)), priming genes for reactivation (*31–35*) and phenotype preservation (*30*). This suggests an underlying optimization principle that balances acute adaptation and long-term functional specialization.

### Metabolic stress: a testbed for nonequilibrium optimization

Metabolic stress provides an ideal context to study nonequilibrium optimization (*36*). To preserve metabolic homeostasis, cells activate transcriptional programs (*37*) in response to metabolic stress such as exercise (*38*), cold (*39*) and fasting (*40*). These metabolic stressors are linked to energy-deficit and upregulate coactivators (*37*) to coordinate transcription factors (*41*) and transcriptional condensates (*42*), thereby modulating metabolic gene programs. These regulated gene programs lead to acquisition of new functions and preservation of cell-specific functional states.

Another challenge during energy deficit is controlling how metabolic activity impacts the cellular milieu, for example ionic imbalance or pH variations. In these cases, cell homeostasis is restored via proton/ion movement across the cell membrane and organelles (e.g. lysosomes) (*43*) via pumps such as V-ATPase and Tmem175 (*44, 45*). This adaptation is also associated with perinuclear lysosomal activity (*46*), which further modulates transcriptional responses for long-term cellular adaptation (*47*). Overall, rather than just enduring these perturbations, cells exploit them, suggesting that low-energy states fine-tune local organization for gain or preservation of function. Yet, the mechanisms of how metabolic fluctuations lead to persistent or adaptive specialization remain largely unknown.

### Cells as models of driven dissipative computation

Understanding nonequilibrium principles remains a fundamental challenge (*48–50*). Functional specialization is generally associated with compartmentalization (*48, 51*) and underlies many fundamental concepts in both biology (*52, 53*) and machine learning (*54, 55*). Cells acquire specific functions by dynamically modulating the formation and disassembly of transcriptional compartments (*16*). The ability to oscillate between specialized (e.g., low-entropy compartments) and non-specialized (high-entropy) states is critical, not only for preserving function but for adaptation and survival (*8, 10, 56*). Moreover, environmental dynamics influences how organizations respond to external drives (*7–10*), introducing timescale complexities that challenge computational formalisms.

An emerging perspective treats cells as systems that perform information processing via dissipative processes (*57–62*). Under sustained energy flux, molecular collectives drive selection, memory, and control through physical constrained computation (*63, 64*). In such systems, organization incurs in thermodynamic costs (erasure, writing, and adaptation) with entropy production (*65–67*). Fluctuations or external variations (gradients, crowding) are not only noise but usable drives. Periodic or asymmetric variations can be rectified/absorbed into directional adaptation computed into net organizational gains (*57–62, 68–73*).

A central outcome is that these systems exploit external energy from drives to compute organization at a lower energy cost (*74–76*). This dissipation-driven organization is at the basis of multiscale computation and energy-efficiency of living systems (*77, 78*). Thus, adopting this framework operationally and asking whether dissipative organization may be a recurring constraint for adaptive specialization across living and computational systems remains a priority (*79–84*)–bridging self-organization and adaptive learning.

### A cross-domain framework for adaptive specialization

Formalizing these processes requires identifying key nonequilibrium rules from empirical observations and adopting a mathematical language that can express hierarchical multi-scale structure, dynamic reorganization, and thermodynamic constraints. Current ML paradigms treat learning as an abstract optimization (static loss minimization under regularizers) decoupled from information-processing costs and experimentally grounded physical budgets. This supports the need for new computational paradigms linking adaptation in cellular and machine systems through shared nonequilibrium constraints.

Here, we combine theory and experimentation to show that asymmetric fluctuations can act as tunable drives being rectified into adaptive specialization. We proceed in four integrated stages: (i) biological and empirical identification of candidate principles for cellular specialization (focusing on dynamic nuclear organization; section 1); (ii) mathematical formalization of such principles (section 2); (iii) computational implementation in ML paradigms (section 3); and (iv) discovery and validation of modulatory design rules across substrates (sections 3-5).

Biologically, we show that cell systems use energetic-state oscillations to modulate nuclear compartments, enhancing functional specialization and homeostasis. We formalize underlying operational principles using operadic algebra and a nonequilibrium optimization inspired by Maxwell’s demon (*85*), which couples gains in organization to information-processing costs (*86, 87*). This translates into a self-organizing learning system that operates through adaptive compartment dynamics. We find that asymmetric fluctuations are control drives that adaptive systems rectify into structural and functional specialization. We validate this biologically, identifying similar drives in cells, revealing that drive-modulators influence transcriptional organizations, epigenetic-phenotype memory, and resilient states. Across domains, increased local organization is linked to dissipation proxies (dissipation not as total heat but as lower-bound process-specific proxies), suggesting that adaptive specialization proceeds under shared constraints and provides design guidelines for programming biological function and engineering thermodynamically informed adaptive AI.

## Results

### 1. Decoding nonequilibrium specialization: a multidomain approach to adaptive systems

To understand how environmental state-variations change specialization of function in cell systems, it is required to map the existing organizational hierarchy from genome architecture to phenotypic outcomes. We hypothesized that some specialized cell functions are the result from the coordinated activity of spatially clustered genes or genomic hubs, constituting the active structural core of transcriptional compartments. To test this, we developed a multimodal framework that systematically links and tests genomic hub identity to transcriptional compartment formation and functional specialization (**Fig. 1A**).

**Fig. 1.**
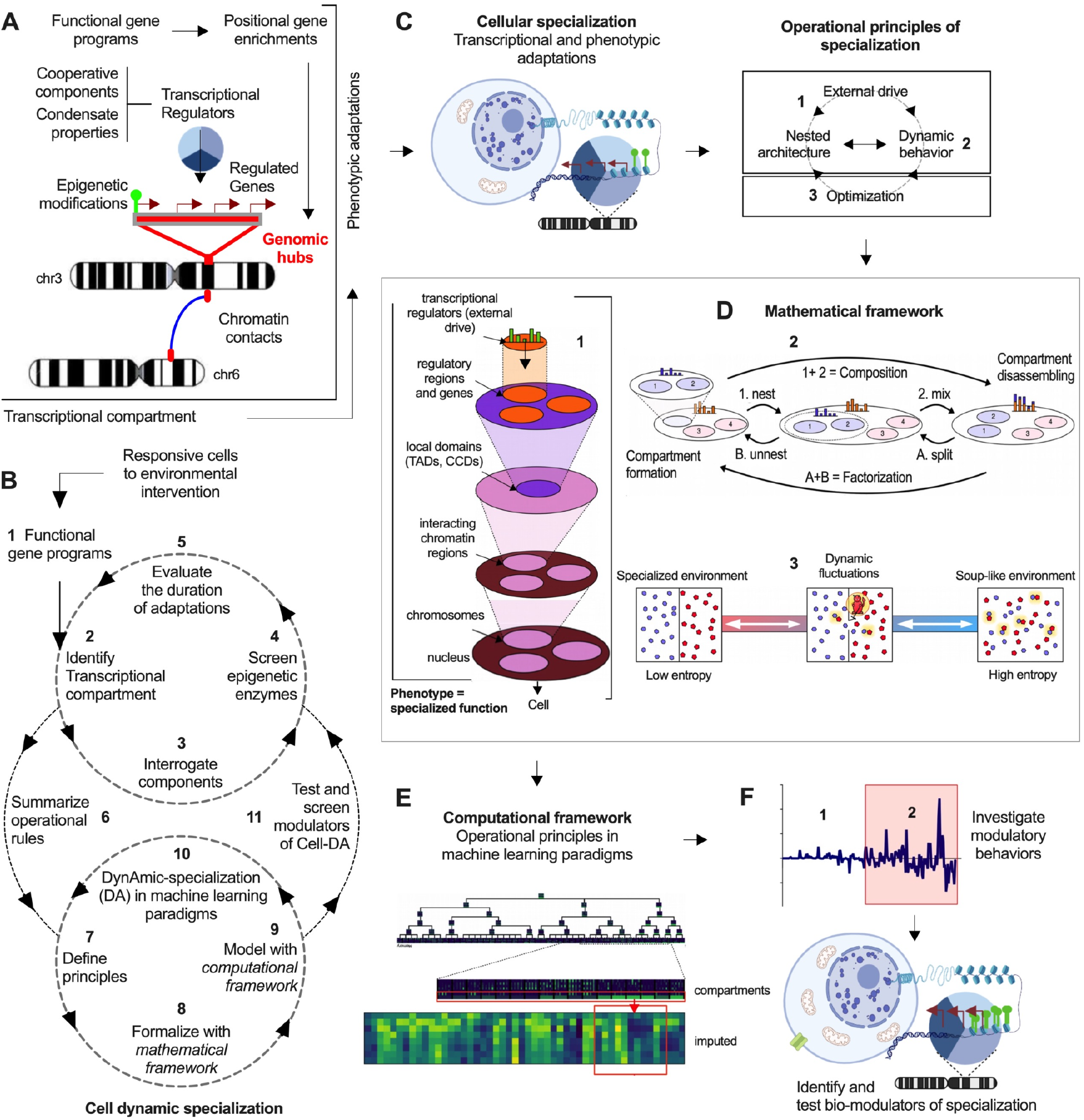
Cell dynamic-specialization (CellDa): a multi-domain approach for investigating principles of specialization. **(A)** Schematic illustration of transcriptional compartment components associated with specific phenotypic adaptations. **(B)** Schematic illustration of the multi-domain approach (CellDa) to study the operational principles for specialization in biological and machine learning systems. **(C)** Illustration representing “Duration of adaptations”. Right panel “Define operational principles” from experimental observations. **(D)** Formalize principles with a mathematical framework. (1) Illustration representing the principles “external drives and nested architecture.” (2) Illustration showing an example of modeling the principle “dynamic operations.” (3) Illustration of a modeling example of the “optimization” principle. **(E)** Formalize principles with a computational framework. Illustration showing implementation of operational principles and “dynamic specialization in ML paradigms.” **(F)** Illustration showing the identification of modulators of “dynamic specialization in machine learning paradigms,” followed by testing of biological modulators (“test and screen modulators of CellDA”).

Our approach integrates several layers of organization, which starts by identifying genomic hubs from gene signatures using positional gene enrichments (*88*). In line with previous studies (*88–91*), these identified genomic hubs can be used to characterize their associated transcriptional components including chromatin contacts, gene expression patterns, bound transcriptional regulators, protein-protein interactions, condensate properties, and epigenetic-linked modifications (**Fig. 1A**). By linking these molecular adaptations to phenotypic changes in diverse environmental conditions, we can quantify and decode the operational principles and logic by which cell systems fine-tune compartmentalization for specialization of function (**Fig. 1A**).

#### 1.1. The CellDa framework

Building on this foundation, we developed Cell Dynamic Specialization (CellDa, **Fig. 1B**), an iterative cross-domain framework that integrates experimental biology, mathematical formalization, and computational implementation to identify principles of specialization (**Fig. 1B**). CellDa operates through four integrated cycles:

[1] Biological discovery (Steps 1-5; **Fig. 1B**): This phase characterizes specialization of cell function in cells during environmental changes, mapping transcriptional and phenotypic adaptations to find operational features (**Fig. 1C**). [2] Mathematical formalization (Steps 6-8; **Fig. 1B**): This phase extracts and formalizes these operational rules of cellular specialization with a principled approach (**Fig. 1D**). [3] Computational implementation (Steps 9-10; **Fig. 1B**): We implement and evaluate the behavior of these principles in machine learning paradigms, revealing emergent behaviors and modulators of specialization (**Fig. 1E, F**). [4] Biological validation (Steps 11; **Fig. 1B**): Computational modeling guides the identification of biological analogs, in particular molecular modulators that can be targeted for enhanced cell programming and disease therapeutics (**Fig. 1F**).

The iteration for bidirectional insights shows that computational systems reveal dynamic organization principles that help identify biological modulators of analogue behavior. Conversely, cell system experiments validate and constraint theoretical models for enhanced computation. The end result is a unified framework that can be applicable to bioengineering and adaptive machine learning.

#### 1.2. SCAT-MSCs as a model for environmentally driven metabolic specialization

To investigate dynamic principles of metabolic specialization, we first defined specific criteria cell systems must meet: (i) robust transcriptional response to nutrient environmental stress, (ii) display an intrinsic ability for persistent functional adaptation, and (iii) directly impact organismal metabolic homeostasis and disease. We then evaluated a set of studies where transcriptomic profiles were conducted after metabolic stress aimed at restoring homeostasis.

This revealed single-cell studies showing that adipose tissue mesenchymal stem cells (AT-MSCs) are highly responsive to caloric restriction (*92*), exercise (*93*), and anti-aging interventions (*94*). In particular, white adipose tissue MSCs respond robustly to these interventions, with strong mitochondrial gene expression (*94*). Moreover, MSCs play a key role in disease prevention through cell plasticity, immune and vascular modulation, tissue regeneration, and homeostasis maintenance (*95*). These features make AT-MSCs a suitable model to study cell specialization as they actively specialize under stress conditions. In this study, we decided to focus on subcutaneous adipose tissue (SCAT) MSCs given their accessibility, metabolic response, and disease relevance. To first confirm their suitability as a model for studying metabolic specialization, we subjected lean mice to 24-hour fasting (a physiological stress that induces transient energy deficit) and isolated cellular populations via cell sorting approaches.

Compared to adipocytes and immune cells, SCAT-MSCs exhibited the strongest response to this intervention: increased expression of mitochondrial genes, elevated epigenetic remodelling with increased H3K4me3 activation marks, and reduced H3K27me suppressive marks at the same regulated gene promoters (**Fig. 2A, fig. S1A, B**). These responses were translated to enhanced mitochondrial activity (**fig. S1C**), supporting a functional response to fasting. These results indicate that SCAT-MSCs are a suitable model for investigating how transient metabolic stress induces persistent molecular and phenotypic adaptations and how this can be generalized to identify principles on how stress modulates specialization of function.

**Fig. 2.**
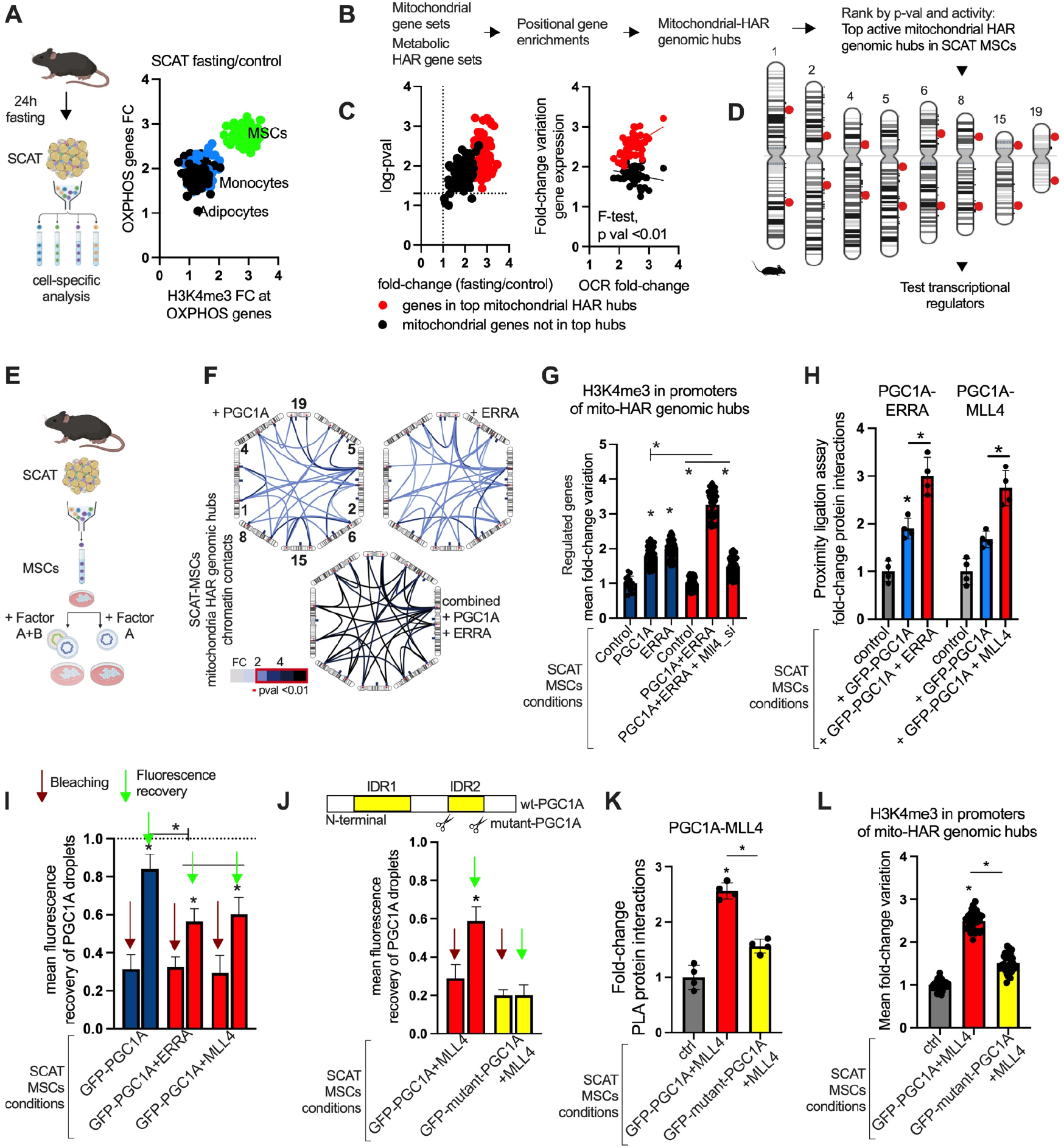
Functional genomics of transcriptional compartments linked to mitochondrial function in SCAT-MSCs. **(A)** Left panel illustrates fasting intervention and cell sorting. Right panels shows fold-to-fold change plot of OXPHOS gene expression and H3K4me3 relative levels in OXPHOS genes in sorted cells from subcutaneous adipose tissue after fasting (MSCs fold-change over control for all data points had a p-value<0.05). **(B)** Illustration of genomic hub identification. Mitochondrial gene sets and metabolic HAR genes were used to identify shared genomic hubs by positional gene enrichments (named mitochondrial HAR genomic hubs or mito-HAR hubs). Genomic hubs were filtered by p-val enrichment and activity. **(C)** Left panel, expression of mito-HAR hubs genes in SCAT-MSCs after fasting (24h) in wildtype lean mice as in **A**. Right panel, fold-to-fold change between mito-HAR hub gene expression and oxygen consumption rate (OCR) in plated cells from mice. **(D)** Illustration of top active mito-HAR genomic hubs. **(E)** Isolation of SCAT-MSCs, plating, and engineering with specific transcriptional regulators. Cells were transfected with expression vectors using the nucleofection for primary cells. **(F)** *in situ* ChIP-loop chromosome conformation assays for chromatin contacts between mitochondrial-HAR hubs in SCAT-MSCs transfected with transcriptional regulators individually or combined (24h). Contacts are represented as fold-change (FC) between overexpression and control groups. Contacts with FC > 2 have p-value <0.01. **(G)** Mean fold-change of H3K4me3 marks in promoters of regulated genes as in **E**. Comparing the effect of RNAi-knockdown for the epigenetic regulator Mll4 (mean FC H3K4me3 levels in promoters > 20 genes/group). **(H)** Proximity ligation assays (PLA) for PGC1A protein interactions in SCAT-MSCs transfected with GFP-PGC1A, ESRRA and MLL4 individually or combined. **(I)** Fluorescence recovery and photobleaching assays (FRAP) in transfected SCAT-MSCs cells individually or combined. Data shows PGC1A-droplet fluorescence recovery (10 seconds post bleaching). **(J)** Upper panel, PGC1A protein illustration with N-terminal region and disordered regions 1 and 2 (IDRs); scissors represent the deletion location to generate mutant PGC1A construct (without IDR2). Lower panel, mean droplet FRAP recovery of wildtype and mutant PGC1A constructs as in **E. (K)** PLA for protein interactions in SCAT-MSCs overexpressing either wildtype or mutant PGC1A with MLL4 combined. **(L)** Mean fold-change of H3K4me3 marks in promoters of mito-HAR hubs from SCAT-MSCs. For all panels, cell experiments were done with 3 independent replicates. Data show mean values and SEM. Unpaired two-tailed student’s t-test for comparing two groups, and ANOVA fisher’s least significant difference (LSD) for multiple groups comparisons. * indicates p-value <0.05.

#### 1.3. Transcriptional compartments associated with mitochondrial function specialization

After identifying SCAT-MSCs as targets to explore functional specialization, we asked whether there are specific genomic regions orchestrating a coordinated response. We hypothesized that genes critical for long-term metabolic adaptation would display two global features: (i) evolutionary constraint, which should reflect the fundamental relevance of energy homeostasis, and (ii) spatial clustering in genomic hubs, enabling the coordinate regulation via 3D genome architecture.

##### 1.3.1. Evolutionary context

Interestingly, mitochondrial function represents an evolutionary contradiction broadly speaking: core metabolic signatures are highly conserved across mammals given their essential role in energy production, yet metabolic capacity shows species-specific adaptation to diverse environments and energetic demands (*96*). This evolutionary constraint indicates that mitochondrial traits have been fine-tuned to optimize energy efficiency, metabolism, and organismal survival. Human-accelerated regions (HARs) are some of the fastest-evolving regions of the human genome, (*97*), many of which function as enhancers regulating gene expression as human-specific adaptations. Given the relevance of metabolic specialization for cross-species conservation, we reasoned that HARs and associated genes can be used as proxies to identify mitochondrial-associated genomic hubs coordinating metabolic adaptation. This will predict that such regions would (i) show spatial clustering along chromosomes, (ii) exhibit dynamic chromatin contacts during metabolic stress, and (iii) respond coordinately to environmental challenges.

##### 1.3.2. Mitochondrial-HAR genomic hubs

In our companion studies, we showed that specific genes associated with HARs spatially cluster in genomic regions that are syntenic in mice, are linked to metabolic function, and show coordinated activity in adipocytes during metabolic stress (*89–91*). These genomic hubs, defined as metabolic-HAR hubs, colocalize within interacting chromatin regions and, during nutrient deprivation, form chromatin contacts enabling the coregulation of genes influencing cell resilience (*89–91*) (**fig. S1D**). However, whether these evolutionarily tuned and metabolically-linked regions also organize mitochondrial associated genes remained unknown.

We therefore asked whether metabolic-associated HAR genes and nuclear-encoded mitochondrial genes (*98*) (from mitocarta) colocalize in specific genomic hubs. Such colocalization and contextual activation would indicate that (i) evolutionary selection has spatially organized metabolic regulators for gene coregulation during environmental stress and that (ii) there are multi-scale constraints on metabolic cellular response (*89, 99*) (**Fig. 1B** “step 1, 2,” **fig. S1E**). To test this, we used positional gene enrichment analysis (*88*), identifying chromosomal regions significantly enriched for this gene set, termed here ‘mitochondrial-HARs’ or mito-HAR genomic hubs (**Fig. 2B, fig. S1F, table S1**).

We next asked whether these genomic hubs coordinate function as transcriptional compartments during metabolic stress. In SCAT-MSCs isolated from fasting (24h) mice, we found highly active mito-HAR genomic hubs, based on: (i) chromatin architecture: increased chromatin contacts (evaluation of top active mito-HAR hub gene promoter regions interaction using a targeted 3C approach). (ii) Transcriptional output: enhanced genomic hub gene expression. (iii) Functional coupling: strong association with mitochondrial respiration in functional assays (**Fig. 2C, D, fig. S1G**).

Overall, these results suggest that evolutionarily tuned hubs organize mitochondrial genes into high-order nuclear compartments, whose coordinated action is associated with metabolic function specialization. This multi-scale genomic constraint (from evolutionary selection to 3D nuclear organization to phenotypic outcome) provides a structural framework for investigating the molecular regulators and operational principles for adaptive specialization.

##### 1.3.3. Transcriptional and epigenetic regulators of mitochondrial-HAR genomic hubs

Given that these genomic hubs provide the structural template for transcriptional compartments, we next aimed at investigating their linked transcriptional machinery. We hypothesized that these compartment hubs required (i) a transcription factor and (ii) a coactivator to nucleate protein complexes and recruit epigenetic modifiers. Our previous work identified the PGC1A coactivator as a regulator of metabolic-HAR hubs activity, influenced by co-factor interaction and metabolic states (*89–91*). Given this, we searched for (i) well known PGC1A-interacting transcription factors (using repositories for protein-protein interactions) that are (ii) linked to metabolic-HAR regions (based on our previous findings (*89*)), (iii) bind promoters of mitochondrial genes, and (iv) show elevated expression during fasting. This revealed one main candidate: the estrogen-related receptor alpha (ESRRA) transcription factor (**Fig. 1B** “step 3,” **fig. S1E**). ESRRA is an orphan nuclear receptor that binds ESRR response elements at mitochondrial gene promoters and has been shown to interact with PGC1A to activate oxidative metabolism programs (*100*).

To test their cooperative regulation at mito-HAR-hubs, we performed chromatin immunoprecipitation (ChIP) SCAT-MSCs from fasting mice. Both factors showed significant enrichment in mito-HAR hub promoter regions (**fig. S1H)**, suggesting co-regulation of the same genes, within specific genomic hubs. We next asked whether their functional cooperation would exacerbate the activity of target genes and associated transcriptional compartments. Gain-of-function experiments using plasmid vectors via nucleofection into extracted SCAT-MSCs (**Fig. 2E, fig. SI)** showed their cooperation, leading to increased chromatin contacts, enhanced promoter binding, and elevated expression of hub genes (**Fig. 2F, fig. S1J, K**). These results suggest that mitochondrial HAR hubs are co-regulated by heterotypic cofactors during environmental stress, with each factor enhancing the activity of the other.

While coregulators drive acute changes, we asked whether persistent epigenetic modifications might extend these adaptations. Transcriptional memory (more robust activity in primed genes after re-exposure to a stimuli) is often associated with histone modifications that persist after transcriptional activity. For instance, histone H3K4me methyltransferases (HMT) and histone H3K27 demethylases mediate chromatin opening at actively transcribed genes through interactions with the transcriptional machinery and condensate formation (*17–19*) We hypothesized that PGC1A/ESRRA recruit histone-modifying enzymes to write marks linked to persistent phenotypes.

To identify epigenetic enzymes, we performed RNAi screens (for 12 histone methyltransferases and demethylases) in SCAT-MSCs cells co-expressing PGC1A and ESRRA (**Fig. 1B** “step 4”). Cells were assayed for H3K4me3 (activation mark) and H3K27me3 (repression mark) at mito-HAR hub promoters. This revealed 2 key modifiers; the histone methyltransferase MLL4 (known as Kmt2d) and the histone demethylase UTX (known as Kdm6a) regulate epigenetic marks of activity and repression, respectively (**Fig. 2G, fig. S2A-C**). Both enzymes had the strongest effect on their respective marks among candidates.

Since MLL4 and UTX cooperatively regulate stem cell epigenetic plasticity through heterogeneous condensates (*17*), we primarily focused on MLL4 as it writes activation marks linked to transcriptional output. MLL4 knockdown via siRNAs substantially reduced epigenetic marks induced by PGC1A and ESRRA (**Fig. 2G**), indicating that MLL4 is required for translating transcriptional activity into stable chromatin activation. To determine whether MLL4 physically associates with the PGC1A/ESRRA complex we used proximity ligation assays (PLA) to detect protein interactions (**fig. S2D**). In cells, this revealed relative protein interactions between PGC1A-ESRRA, and PGC1A-MLL4, which exhibited a synergistic effect in co-transfections (**Fig. 2H, fig. S2D-F**).

##### 1.3.4. Model for cooperative compartment assembly

Together, these results establish a tripartite regulatory module composed of heterotypic cooperative factors: ESRRA provides DNA-binding specificity, PGC1A coactivator nucleates, stabilizes the complex, and coordinate cofactor recruitment, while MLL4 writes persistent H3K4me3 marks to maintain open chromatin states in evolutionary tuned regions related to mitochondrial function. In addition, this indicates cooperative amplification as each module was found to enhance the other. This configuration has functional implications as it allows signal integration, multiple input or modulatory drives can be processed through the same effector machinery. Moreover, it enables tunability of the system as altering the activity or properties in one of the components can shift the equilibrium between their cooperative state. Finally, it can create memory or persistent states, which is so important for continuous systems, enabling priming and reactivation.

#### 1.4. Transcriptional plasticity influences adaptation of phenotype

Given our observations, we next asked how these factors organize spatially to coordinate the activity of distant and genomic loci. Recent work has shown that transcriptional coactivators form liquid-like condensates (membraneless compartments) that purposely concentrate regulatory machinery at active, distant loci (*15, 16*). In addition, it has been shown that PGC1A contains IDRs at its C-terminus that drives liquid-liquid phase separation, droplet formation, and gene coregulation with interacting factors (*42*) (**fig. S2G**). We therefore hypothesized that PGC1A nucleate such mito-HAR hub transcriptional compartments, and that recruitment of ESRRA and MLL4 would alter condensate material properties in ways that affect function.

##### 1.4.1. PGC1A forms dynamic condensates that recruit cooperative regulators

We first confirmed PGC1A-droplet behavior using live-cell imaging assays (fluorescence recovery after photobleaching (FRAP)) of GFP-tagged PGC1A in SCAT-MSCs (**Fig. 2I. fig. S2G-H**). As expected, GFP-PGC1A formed discrete nuclear puncta alone (0.5-2 μm diameter; **fig. S2H**), and exhibited rapid FRAP recovery characteristic of liquid-like behavior (**Fig. 2I**). We next asked whether binding partners would affect PGC1A droplet dynamics and properties. For example, when co-expressed MLL4 with GFP-PGC1A, droplet morphology remained unaffected (**fig. S2H**). However, combined MSCs transfection of interacting regulators (GFP-PGC1A + ESRRA or + MLL4) reduced GFP-PGC1A droplet recovery more than individual transfection (**Fig. 2I**). This reduced droplet recovery indicates increased viscosity, a transition state from liquid-like to gel-like material properties (*24*). Importantly, the increase in droplet viscosity directly correlated with enhanced MLL4 protein interaction (**fig. S2I**), suggesting that local protein recruitment alters the physical state of the condensate, opening questions on the functional outcome of this tunable behavior.

##### 1.4.2. Intermediate droplet viscosity optimizes epigenetic activity

This raised one key question, namely, whether there is an optimal viscosity linked to epigenetic function. If droplets are too liquid, components might not be stably associated with chromatin, and if too solid, they might lose the dynamic features required for enzyme positioning and activity. We tested this by evaluating the association between H3K4me3 levels at mito-HAR promoters and droplet FRAP recovery across different conditions. Strikingly, maximal H3K4me3 marks in promoters of regulated mito-HAR hub genes were found strongly associated with intermediate viscosity (reduced recovery but still dynamic) of PGC1A fluorescence recovery in dual-transfected SCAT-MSCs (PGC1A+MLL4; **fig. S2J**). This inverted relationship (high epigenetic marks with reduced viscosity) suggests that partial gelation creates an optimal environment for MLL4 recruitment, with enough concentration for activity and dynamic enough for substrate turnover.

To test whether the IDR is required for this optimal viscosity and in line with previous studies (*42*), we generated a PGC1A mutant without the IDR2, while preserving the rest of the C-terminal domain (**fig. S2G**). This mutant revealed solid-like behavior with arrested fluorescence recovery in MSCs with combined transfection (**Fig. 2J**), impaired MLL4 protein interactions, and reduced epigenetic activity in mito-HAR hubs (**Fig. 2K, L**), suggesting that transition to a solid-like or gel-arrested droplet viscosity state impairs transcriptional plasticity.

Together, these results establish a mechanistic model, where PGC1A IDR enables the formation of dynamic condensates whose viscosity can be fine-tuned by cofactor interaction/recruitment. Partial gelation by interactions and increased epigenetic marks suggest that the creation of an optimal local environment is crucial for the epigenetic machinery while maintaining the molecular dynamics for chromatin modifications. This material-state transition offers a biophysical mechanism linking transcription compartment regulation to persistent epigenetic memory and functional specialization.

##### 1.4.3. Duration of molecular and phenotypic adaptations

Given our observations on cooperative regulation and functional adaptation, we next evaluated whether this plasticity is associated with epigenetic memory–chromatin modifications that persist post-stimuli (**Fig. 1B** “step 5”). To test this, we nucleofected SCAT-MSCs with expression vectors (peak expression at 24-48h and decay by 5-7 days) and tracked H3K4me3 marks, gene expression, and mitochondrial function over 15 days (**fig. S3A**).

Compared to individual (PGC1A alone) and dual (PGC1A+ESRRA) expression, combined transient expression of PGC1A, ESRRA, and MLL4, dramatically prolonged H3K4me3 marks, gene expression in mito-HAR hubs, and the duration of enhanced mitochondrial function (elevated through day 15; **Fig. 3A, B, fig. S3A-E**). As these adaptations exceeded the median lifespan of changes induced by PGC1A and ESRRA alone, this indicates epigenetic maintenance through MLL4 is associated with durable metabolic specialization.

**Fig. 3.**
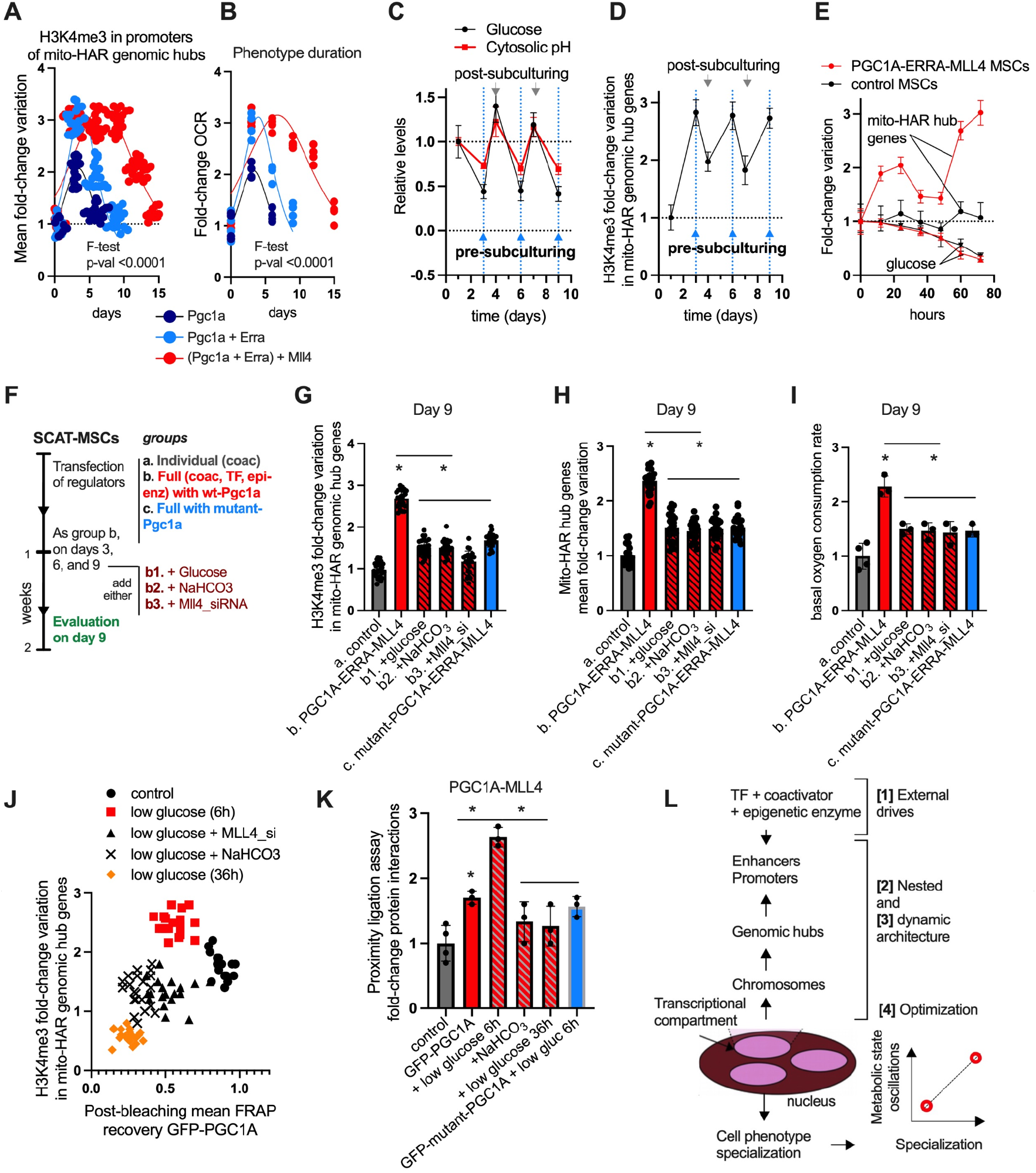
Environmental oscillations are required for mitochondrial function specialization in SCAT-MSCs. **(A)**Time-series for H3K4me3 mean fold-change (FC) in mitochondrial-HAR hub genes in transfected SCAT-MSCs with regulators individually or combined (mean FC of H3K4me3 levels in promoters > 20 genes per group). **(B)** Time-frame for FC of basal oxygen consumption rate (OCR) in the same conditions as in **A. (C)** Glucose, and cytosolic pH variation over time (days) in murine SCAT-MSCs transfected with PGC1A, ESRRA and MLL4 combined, and in control cells (dotted line). Arrows show pre- and post-subculturing day variation. **(D)** H3K4me3 FC over time (days) as in **C**. Data shows mean FC of H3K4me3 in promoters >10 genes per group. **(E)** FC gene expression (mitochondrial-HAR hub genes) and glucose levels over time (hours) in SCAT-MSCs control and transfected with regulators (P-value for group comparison <0.01). **(F)** Schematic illustration for perturbation experiments. SCAT-MSCs cells overexpressed PGC1A, individually (here representing control), and combined (as in **E**), or combined with mutant-PGC1A. During the days before sub-culturing (as shown in **C**), media was supplemented with glucose or NaHCO_3_ (1 mM). We used Mll4-siRNA in the same days. On day 9, SCAT-MSCs were collected and assayed for (**G**) H3K4me3 (FC in promoters>20 genes per group), (**H**) gene expression (FC, >20 genes per group), and (**I**) basal basal oxygen consumption rate (OCR). **(J)** H3K4me3 FC in mito-HAR hub promoters (y-axis) and mean fluorescence recovery after photobleaching (x-axis) from GFP-PGC1A droplets in SCAT-MSCs treated in different conditions. **(K)** PLA for PGC1A-MLL4 interactions in the same conditions as in **J. (L)** Summary illustration showing operating principles [1-4] for specialization of mitochondrial function in SCAT-MSCs. For all panels, cell experiments were done with 3 independent replicates. Data show mean values and SEM.Unpaired two-tailed student’s t-test for comparing two groups, and ANOVA fisher’s least significant difference (LSD) for multiple groups comparisons. Nonlinear regression (Lorentzian-Cauchy model) and extra sum-of-squares F test to compare conditions. * indicate p-value <0.05.

The persistence of epigenetic marks of gene activity beyond transcription factor expression suggested that these loci are ‘primed’ for robust reactivation upon subsequent stimulation, which is a hallmark of transcriptional memory. To test this, we challenged cells on day 15, when H3K4me3 were no longer increased, with CL-316,243, a beta-3 adrenergic receptor agonist that has been shown to activate mitochondrial gene programs (*101, 102*). In cells that expressed both PGC1A and ESRRA (day 1-3 of cell culture) and evaluated post beta-3 treatment (day 15 of cell culture), the agonist induced modest responses: OCR increased 1.8-fold only on day 17 and no changes were seen on day 20 (**fig. S3F**). However, in cells with PGC1A+ESRRA+MLL4 (15 days prior), the agonist induced a more robust and prolonged functional response (**fig. S3F**) with the corresponding transcriptional changes (**fig. S3G**), suggesting both prolonged adaptations and priming for reactivation.

This hypersensitivity and reactivation show features of transcriptional memory: prior exposure to the regulatory triad establishes a chromatin state that enables enhanced responsiveness to subsequent metabolic stress, even after the initial factors have been cleared. This priming mechanism could enable cells to mount stronger responses to recurrent metabolic stress or environmental variations. These results led us to ask whether enhanced mitochondrial function represents irreversible differentiation or reversible specialization. To distinguish this, we evaluated whether our interventions promote MSCs to retain stem cell properties or adipocyte cell commitment. Lipid accumulation assays in engineered MSCs showed no differences compared to control cells and much less compared to cells induced for adipocyte differentiation (**fig. S3H, I**). SCAT-MSCs transfected with regulators showed expression of stem-like markers and transcription factors, such as Oct4, Twist, Sox2 and Nanog, as well as preadipocyte factors such as Cebpb, Cebpa, Klf4 and Klf5 (**fig. S3J**), suggesting a stem-like preadipocyte state.

Functionally, colony forming assays (a measure or stem cell self-renewal) showed enhanced self-renewal in SCAT-MSCs from fasting mice and in engineered cells (**fig. S3K**). Overall, these results indicate that metabolic specialization and stem cell maintenance are compatible. The PGC1A/ESRRA/MLL4 regulatory axis establishes a specialized yet plastic state, where enhanced mitochondrial capacity can be coupled with preserved developmental potential. This reversible specialization may enable MSCs to meet metabolic demands while retaining the ability to respond to future developmental cues.

#### 1.5. Nutrient oscillation modulates transcriptional plasticity for cellular specialization

Cooperative regulation and epigenetic memory lead to functional specialization, yet they raise a deeper question on how cells sustain such specialization over time and over environmental changes? In standard culture, MSCs experience fluctuating nutrient availability since they consume glucose between medium changes. Instead of being a technical artifact, we hypothesized that such metabolic oscillations might serve as an environmental tuning signal that reinforces specialization and is analogous to how periodic training enhances learning in adaptive systems.

##### 1.5.1. Glucose oscillations drive cyclic transcriptional activation

As mentioned before, during low glucose and catabolic states, cellular pH becomes acidic, impacting macromolecular interactions, viscosity, and transcriptional processes (*22, 43*). To evaluate the associated processes, we confirmed glucose level variation in SCAT-MSCs before (low-glucose) and after subculturing (passaging days or high glucose, **Fig. 3C**), with cellular pH closely following glucose variation patterns (**Fig. 3C, fig. S4A**). Importantly, on subculturing days when glucose was low with cytosolic acidity, we observed increases in mito-HAR hubs–H3K4me3 levels, expression of transcriptional regulators, PGC1A-MLL4 interactions, and mito-HAR hubs chromatin contacts (**Fig. 3D, fig. S4B-D**), supporting phased-specific activity. These observations suggest that metabolic oscillations actively tune transcriptional compartments, with each low-energy phase reinforcing specialization of function. This positions nutrient fluctuation not as a stressor to be tolerated, but as a signal that drives optimization. These nutrient fluctuations can be extrapolated to organisms as they experience cyclic periods of fasting, which has been associated with circadian rhythms and overall health.

##### 1.5.2. Priming and reactivation of transcriptional states

To understand how metabolic oscillations influence engineered expression, we examined priming and reactivation of mito-HAR hubs during the first three days post-transfection. This revealed a biphasic pattern: (i) SCAT-MSCs with combined expression showed transient gene and H3K4me3 regulation with a two-fold increase after 24 hours, representing direct transcription factor-driven activation; (ii) the latter is followed by decreased expression or transient suppression; and a (iii) subsequent three-fold amplified increase after 60 hours (**Fig. 3E, fig. S4E**). Around the last phase (iii), glucose levels were reduced across both groups (**Fig. 3C**), suggesting that transient expression of cooperative regulators generates primed chromatin states, including their own expression, reactivated during subsequent nutrient deficits. The more robust expression of mito-HAR hub genes upon reactivation suggests a role for transcriptional memory in metabolic specialization. This amplification loop could enable progressive specialization over multiple metabolic cycles and enable cells to be specialized while preserving adaptation.

##### 1.5.3. Environmental perturbations impact sustained specialization

The association between glucose oscillation and transcriptional plasticity suggests causation, yet we still need to determine whether metabolic state oscillation is required for persistent specialization. To evaluate this, we designed a perturbation experiment to dissect the contributions of specialization components including transcriptional regulators, epigenetic enzymes, nutrient oscillation, and pH fluctuation (**Fig. 3F**).

We established a baseline using expression of regulators (ESRRA and MLL4) combined with either wildtype or mutant (IDR) PGC1A (**Fig. 3F**). During the 24 hours before the subculturing day (day 3, 6, and 9), we applied one of the three interventions: (i) Glucose supplementation (preventing deficit); (ii) pH buffering (1 mM NaHCO_3_, preventing acidification); (iii) MLL4 knockdown (siRNA, depleting the methyltransferase). On day 9, we assessed mito-HAR hub transcriptional activity and associated functional activity (**Fig. 3F**). This revealed a multi-factor effect, in which the activity linked to mito-HAR hubs (H3K4me3 levels, gene expression, chromatin contacts, and phenotype output) was impaired by each of the perturbation conditions: lack of low-glucose and cytosolic acidity oscillations (metabolic state variation), no MLL4 expression, and the absence of PGC1A disordered region (**Fig. 3G-I, fig. S4F-H**).

These results indicate that specialization of function requires the convergence of several multi-scale components: transcriptional regulators provide specificity, glucose oscillation provides temporal patterning, pH acidification provides physicochemical modulation, MLL4 provides epigenetic stability, and PGC1A IDRs provides adaptive compartmentalization. Removing any component leads to failing to achieve robust, persistent specialization.

##### 1.5.4. Epigenetic memory persists through cell fate transitions

Another test of epigenetic memory is whether this adaptation persists after cell fate transitions. To evaluate this, we induced adipocyte differentiation in SCAT-MSCs previously subjected to the oscillating regimen with PGC1A+ESRRA+MLL4 expression (days 0-12; **fig. S4I**). Remarkably, mito-HAR hub activity (gene expression and H3K4me3 levels) and mitochondrial activity persisted after adipocyte cell commitment (day 10 post-induction; **fig. S4I-M**). This suggests that metabolic memory acquired in stem cells influences the functional properties of their differentiated progeny. Mechanistically, this indicates that acidic cellular pH, downstream to glucose deficit, might be crucial for the observed adaptations. Specifically, trans-differentiation persistence shows that nutrient oscillations and cooperative transcriptional regulation establish stable epigenetic states that survive genome-wide reprogramming during cell fate transitions, a hallmark of robust cellular memory.

##### 1.5.5. pH-dependent modulation of droplet dynamics influence persistent specialization

Given our observations on how the disordered region of PGC1A is required for specialization and that intermediate droplet viscosity is optimal for MLL4 recruitment, we aimed at determining whether nutrient stress naturally modulates droplet material properties to achieve this optimal state. To test this, we tracked the effects of low glucose (over 36h) on droplet dynamics and associated functions. In low-glucose SCAT-MSCs, PGC1A droplets adopted a gel-like viscosity after 6h, which changed to arrested dynamics after 36 hours (36h; **Fig. 3J**). Notably, gel-like droplet recovery was associated with increased H3K4me3 levels in mito-HAR hubs and was dependent on cytosolic acidic pH (blunted by pH correction) and Mll4 expression (**Fig. 3J**). In agreement, low glucose increased PGC1A-MLL4 interaction, was influenced by acidic pH (reduced by pH buffering) and was reduced by 36h low-glucose or by IDR mutation (**Fig. 3K**). Together, these results reveal that acidic pH during low-energy states drives a time-dependent droplet gelation that creates a transient window of optimal viscosity for epigenetic enzyme recruitment and chromatin modification.

#### 1.6. Summary: Operational rules from biological observations

Taken together, these results show a multi-scale framework for metabolic specialization in SCAT-MSCs. For example, at the genomic level, evolutionarily tuned mito-HAR hubs provide spatial templates for coordinated gene regulation. At the protein level, transcriptional regulators form cooperative complexes and their assembly is regulated by nutrient variation. At the biophysical level, PGC1A IDR enables the formation of dynamic condensates with material properties (viscosity) that are modulated by pH to create a transient window of optimal cofactor-recruitment and epigenetic activity. At the temporal level, nutrient oscillations provide repeated low-energy states that recursively reinforce specialization through priming and reactivation. On the functional level, molecular adaptations translate into long-lasting mitochondrial function that persists cell fate transitions.

Notably, this system displays adaptive nonequilibrium processes, where hierarchical and nested organization spans multiple scales. It is a system that requires external inputs that tune internal states (nutrient oscillation), operates through dynamic assembly/disassembly of functional structures (transcriptional compartments), produces persistent states (epigenetic memory), and optimizes specialization through repeated cycles instead of sustained stimulation (**Fig. 3L**). To understand whether these features represent general principles of specialization applicable to other systems (including computational) rather than being system-specific rules, we sought to abstract the essential organizational logic and formalize it mathematically. Our goal was to identify a minimal set of operational rules that capture the essential system dynamics and to test whether the same principles can drive specialization in silico. In the following section, we formalize these features mathematically, extract generalizable principles, and implement them computationally to discover cross-domain modulators of specialization.

### 2. Mathematical modeling of operational principles for cellular specialization

We used a principled approach to abstract operational rules from our experimental observations (**Fig. 1B** “step 6,”), aiming to develop a framework for adaptive specialization. This is designed to be extensible (adding additional rules), and is used here to identify candidate organizing principles of specialization in adaptive systems. In particular, how transient recurrent organization (compartments) may contribute to specialization and preservation of function.

We focused on the following operational principles from our specialization readouts (**Fig. 3L**): [1] Multiscale nested architecture: dynamic compartments are scaffolded by the genome architecture, which exhibits hierarchical structure. [2] Cooperative operations: mechanisms enabling the formation and disassembly of cooperative organizations. [3] External drives: transcriptional regulators and metabolic states act as external drives that promote the formation, dissolution, and tuning of transcriptional organizations. [4] Optimization principle: metabolic state oscillation acts as an optimization drive. For example, glucose oscillations reactivate primed transcriptional states, modulating transcriptional compartment formation and epigenetic activity, optimizing persistent cellular specialization (**Fig. 3L**).

A dynamic and structural model can encode principles [1] through [3], but our observations support that these processes are constrained by energy flow and dissipation. Compartment formation, epigenetics, and reorganization do not take place for free, and likely require nonequilibrium dynamics and energy expenditure. Thus, our framework aims to include an operational budget for these operations. In the next sections, we (i) formalize these operational rules (**Fig. 1B** “steps 7, 8”), (ii) embed them in an optimization principle compatible with thermodynamic constraints (costs accounting), and (iii) implement them computationally to explore their qualitative behavior in ML paradigms.

#### 2.0. Theoretical foundations and design principles

We aim to develop a thermodynamically informed framework that helps interpret aspects of cellular specialization while providing design heuristics for adaptive systems. This is informed by three needs:

(i) Substrate-agnostic formulation: The framework is expressed in a way that does not depend on the details of any particular implementation, with the aim of enabling comparison across information-processing systems (cells or computational models). This approach uses mathematical abstractions to capture key organizational principles without depending on a specific system. (ii) Thermodynamic accounting: Unlike current ML approaches, which do not take into account physical constraints or processing costs, our framework aims to introduce a bookkeeping layer for tracking proxies linked to information processing costs. The intent is to check internal consistency with the second-law at the level of sign and accounting (*86, 87*), tracking how local organization is balanced by dissipation elsewhere. (iii) Prescriptive architectures via constructive operations: The framework cannot only describe static systems but offer constructive principles for designing optimal architectures, using intrinsic operations that discover sufficient structure for a given input.

These objectives led us to operadic algebra as it provides a flexible mathematical language from category theory that cleanly expresses hierarchical composition/factorization and multi-scale organization (*103–105*). Operads offer tools needed to model systems that build architecture from simpler structures, decompose them when required, and keep consistency across hierarchies. We further explore how combining this structural formalism with a nonequilibrium optimization principle could enable the formalization of how adaptive systems trade-off organization, adaptation, and information-processing costs.

#### 2.1. Modeling multiscale compartmentalized systems as nested structures

Biological systems use nested hierarchical architectures to represent related yet dynamic organizations and where structures at one scale usually serve as components of the next scale up (Biological observation/rule [1]). To model multi-scale organizations, we use nested blocks, where each block has a circular structure (cytosol or nucleoplasm) containing smaller circular structures for nesting (organelles or transcriptional compartment components; **Fig. 4A, B, fig. S5A**). We call these structures “*cells*” or “*artificial cells*,” and the whole system represents a multi-scale compartmentalized system with specialized functions that can be visualized as a tree (**Fig. 4B**). Tree topology naturally enforces nested relationships that enables recursive operations propagating through hierarchies needed for global function.

**Fig. 4.**
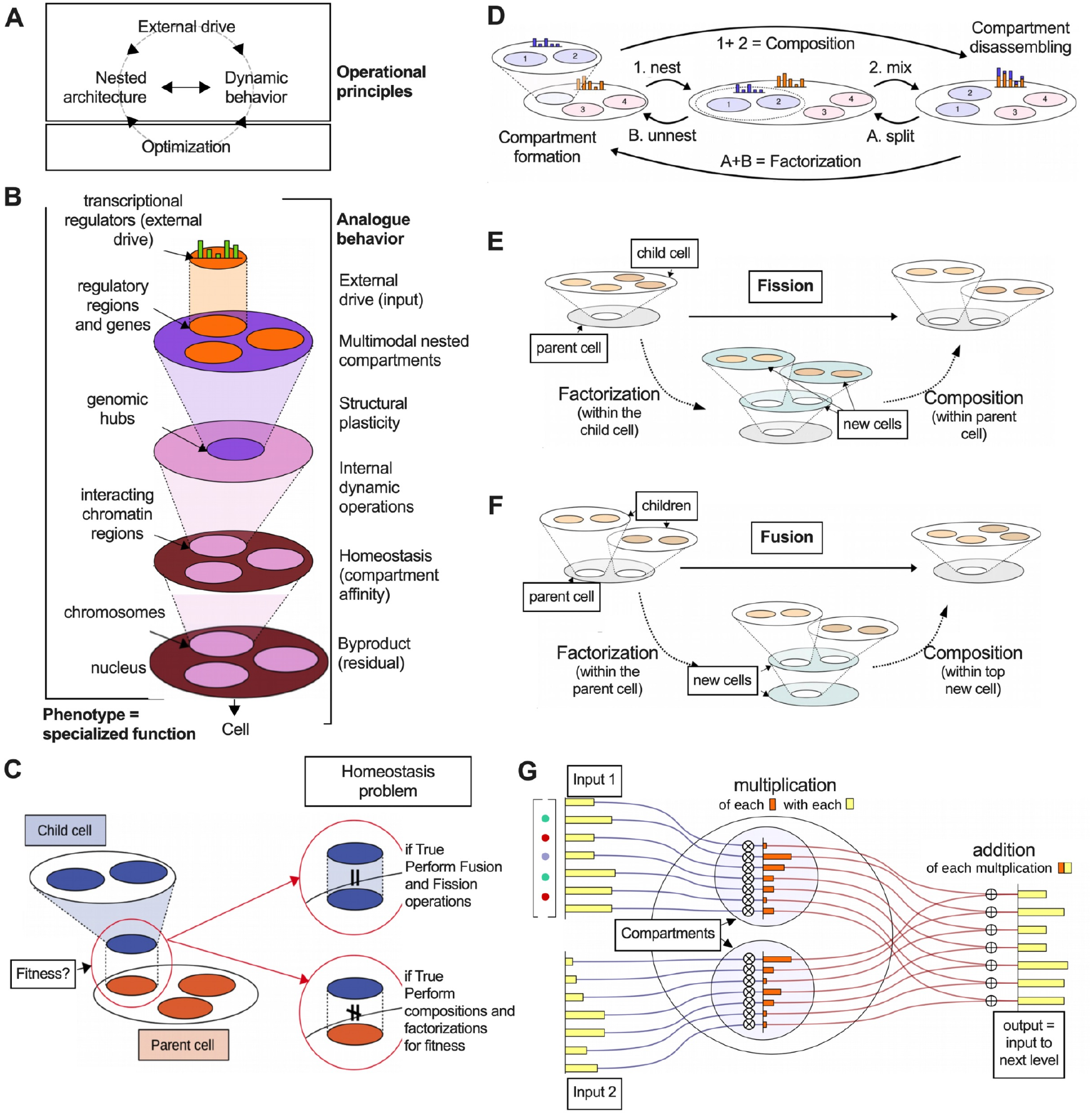
Modeling operational principles for specialization in multi-scale compartmentalized systems. **(A)** Operational principles. **(B)** Schematic illustration summarizing analogue behavior of phenotype specialization in SCAT-MSCs using a nested, multi-scale, and tree-like framework with dynamic properties. **(C)** Illustration of the concept of fitness. One cell (child) fits the compartment of another cell (parent) if the encoding vectors are equal. If they don’t fit, basic dynamic operations can be performed (see next panel) to reach homeostasis. If the cells of the tree-like structure reach homeostasis, we use 2-level operations fission and fusion of compartments for further tuning. **(D)** Basic dynamic operations in framework. Left-to-right, composition of compartments by nesting inner-cells followed by mixing of the contents within the outer-cell, which generates cytosolic residuals. Right-to-left, factorization (compartment formation) by isolating and splitting compartments and part of cytosolic contents, followed by unnesting and creating new cells (new-level structure). **(E)** Illustration showing fission of compartments in the child cell by factorization (into 2-level tree) and composition within the parent cell. **(F)** Illustration showing fusion of compartments in the children cells by factorization of the parent cell (into 2 new cells) followed by composition of the children cells with the middle cell. **(G)** Illustration showing how an abstract cell of dimension 7, equipped with 2 organelles, processes a 2 x 7-matrix of real numbers. Its output is then a vector of dimension 7, and it is the result of a component-wise multiplication of the columns of the matrix (i.e. called “input 1”and “input 2”) with the organelles of corresponding index, followed by a component-wise sum of these multiplication.

##### 2.1.1. Mathematical definition

The outer environment and the inner compartments of every artificial cell are characterized by vectors of real numbers with fixed dimension. Each vector component represents a degree of specialization, such that the complete vector captures the specialization profile of its associated environment. For a every non-negative integer *N*, we define an artificial *cell c of dimension N* as a quadruple *(n, res(c), cyt(c), org(c))* which consists of the following: [1] a *n* is a non-negative integer, representing the number of organelles; [2] *res(c)* is a non-negative real number called the residual; [3] *cyt(c)* is a *N*-dimensional vector of real values representing the cytosolic content; [4] *org(c)* is a *n*-tuple *(org(c)*_*1*_, *org(c)*_*2*_, …, *org(c)*_*n*_) of *N*-dimensional vectors of positive real values where the *i-th* component *org(c)*_*i*_ represents the *i-*th *organelles*.

##### 2.1.2. Hierarchical organization and specialization

We model multi-scale compartmentalized systems that can specialize in *N* different functions as trees of *cells* of dimension *N* (comprising roots, junctions, and leaves, each labeled by an artificial cell of dimension *N*). Each artificial *cell* junction has a number of organelles equal to the number of nested *child cells* (e.g., organelles, **fig. S5B**). To assess specialization within an artificial *cell*, we define the content *K(c)* (of a *cell c)* as the *N*-dimensional vector resulting from the sum of the cytosolic content *cyt(c)* and all organelle vectors nested in that cell (*org(c)*_*1*_, *org(c)*_*2*_, …, *org(c)*_*n*_). This content vector describes everything “inside” of the artificial *cell* system, independent of its compartmentalized structure.

##### 2.1.3. Fitness and homeostatic states

We model temporal states when an artificial nested *cell* (*d*) fits the (*i-*th) organelle of another artificial *cell (c)* if the content *K(d)* is equal to the organelle vector (*org(c)*_*i*_). This evaluation criteria creates a concept that we call local “fitness,” which specifies whether an artificial parent *cell d* is a match with the *i-*th organelle of the child *cell c*. From this we can define that a tree of *cells* can be in two states: [1] Homeostatic state: when the content of the *child cell* equals the content of the *organelle* of the *parent cell*. [2] Non-homeostatic state: when this equality evaluation does not hold (**Fig. 4C**).

In the next sections, we use this concept of fitness or suitability to determine whether artificial *cells* can maintain their current states or require updates through a top-down approach, which creates tunable operations or “contracts” between *nested cells*. This process allows us to define operations that preserve consistency within the tree structures and those that promote re-arrangements given different inputs and model computation fidelity.

#### 2.2. Modeling dynamic behavior with operadic operations

Biological systems use dynamic assembly and disassembly through structural operations that transform nested architectures yet maintain consistency across scales (Biological observation/rule [2]). Trees of artificial *cells* represent multi-scale systems, which can serve as hierarchical data storage units. To integrate dynamic operations for structural optimization, analogous to adaptive transcriptional compartments preserving multi-scale consistency (e.g., formation and disassembly of a compartment, fusion and fission of several compartments), we use concepts from operad theory (*103–105*). These are oriented towards tree-like structures, implementing compositions and factorizations, while maintaining global consistency.

##### 2.2.1. Composition operations

We composed two artificial *cells* by inserting the nested *cell* into an organelle of the parent *cell* and mixing their content (**Fig. 4D**, supplementary text, part 1, Def.1.6). The composition of *cell d* with *cell c*, at the *i-*th organelle of *c*, denoted cell *c* ◦_*i*_ *d* produces a cell whose: [1] The number of organelles is equal to the sum *n + m* (where n and m are the organelle counts of c and d, respectively). [2] Organelle collection of *org(c* ◦_*i*_ *d)* consists of all the organelles of *c*, except the *i-*th organelle *org(c)*_*i*_ is replaced with the organelles of *d*, (where *n* and *m* denote the number of organelles of *c* and *d*, respectively: *(org(c)*_*1*_, …, *org(c)* _*i*−*1*_, *org(d)*_*1*_, …, *org(d)*_*m*_, *org(c)*_*i+1*_, …, *org(c)* _*n*_ )) (2.2.1-A). [3] The cytosolic content *cyt(c* ◦_*i*_ *d)* equals to the sum of vectors *cyt(d)* and *cyt(c)*. [4] The residual *res(c* ◦_*i*_ *d)* is the sum of the *res(d)* and *res(c)*.

##### 2.2.2. Factorization operations

From compositions, we derived the reverse process, factorizations (**Fig. 4D**). Intuitively, a factorization represents the formation of a compartment within a *cell c*, which, once created, becomes the *i-th* organelle of a *cell c’*. Formally, a factorization of a *cell c* is a pair of *cells (c’, d)* as well as an index _*i*_ satisfying the equation *c = c’* ◦_*i*_ *d* (**Fig. 4D**).

##### 2.2.3. Parallel compositions

Beyond compositions at a specific organelle of an artificial cell, we consider simultaneous compositions of *cells* as follows: for a given *cell c* with *n* organelles and a given *n*-tuple of *cells d = (d*_*1*_, …, *d*_*n*_ ), we formalize the simultaneous composition of *c* with the collection of cells *d*_*1*_, …, *d*_*n*_ by *c* ◦ *d = (*…*(c*◦_*n*_*d*_*n*_)◦_*n*−*1*_ *d*_*n*−*1*_*…)*◦_*1*_*d*_*1*_ (3.2.3-A). Since the order of composition does not affect the results, our operation is operadic, which facilitates modeling exchanges between *cells* through dynamic loops of compositions and factorizations (**Fig. 4D**, loop of arrows).

##### 2.2.4. Content exchange mechanism

When a *cell d* represents the *i-*th *organelle* of *cell c*, then we can model content exchange between c and d by: [1] first composing *d* at the *i-th* organelle of *c*. [2] Subsequently factorizing the composite *c* ◦_*i*_ *d* as a pair *(c’, d’)* such that *d’* possesses the same *residual* and *organelles* as *d* and such that the relation *c* ◦_*i*_ *d = c’* ◦_*i*_ *d*’ holds. Passing from the pair *(c, d)* to the pair *(c’, d’)* simulates content exchange, which we use to evaluate fitness or homeostasis between two cells as mentioned before. In Supplementary text (section 4, part 1), we draw a parallel between these types of tuning mechanisms and the concept of homeostasis in cellular and artificial systems.

##### 2.2.5. Fission and fusion operations

Factorizations and compositions are further used to model fission (*division*) and fusion (*merging*) of artificial *cells* in trees containing at least two hierarchical levels (a *parent cell* and *child cells*). Division (fusion): we *divide* a *child cell d*, associated with the *parent c*, by [1] first factorizing the *cell d* into two *child cells d*_*1*_, *d*_*2*_ and a *parent cell c’* and then [2] composing the new *parent cell c’* (**Fig. 4E**). Merging (fission): we *merge* two *child cells d*_*1*_, *d*_*2*_, associated with different *organelles* of a *parent c*, by [1] first factorizing the *parent c* as a *parent cell c’* and a *child cell d* (*d* contains the two *organelles d*_*1*_ and *d*_*2*_), and [2] then composing the *child cells d*_*1*_ and *d*_*2*_ with the *parent cell c* (**Fig. 4F**). Overall, composition operations within tree-like structures provide a simplified yet powerful language for modeling dynamic compartmentalizing behavior analogous to molecular nuclear operations in cell systems (**fig. S5C**).

#### 2.3. Modeling interactions with external drives

Our formalization shows that compositions and factorizations allow exchanges between *artificial cells*. However, these interactions are internal and do not capture interactions with external drives (e.g., data inputs), so it requires mechanisms for external signals to influence structure (Biological observation/rule [3]-input-driven tuning). To model this, we used an analogy between external inputs (akin to nutrients, signals, data) and information directed to the *cell* (**Fig. 4A, B**).

##### 2.3.1. Information processing and forward propagation

An information intake produces an output that possesses the structure of the input and can be extended throughout the tree via forward-propagation. Specifically, we model this process by representing an *abstract cell* of dimension *N* equipped with *n organelles* as a collection of *n×N*-matrices. Each matrix represents a state of the environment providing external “information” drives to which the *cell* can respond. We regard these *n×N*-matrices as *n-*dimensional vectors *a = (a*_*1*_, *a*_*2*_, …, *a*_*n*_) whose *i-th* component *a*_*i*_ is a *N*-dimensional vector *(a*_*i,1*_, *a*_*i,2*_, …, *a*_*i,N*_ ). Each quantity *a*_*i,u*_ represents an amount of input drive to be processed (referred to as the *u-*th function of the *i-*th organelle of the *cell c; see* Supplementary text).

##### 2.3.2. Action of cells on inputs

We then define the intake of an *n×N*-matrix *a* by the *cell c* through a series of multiplications and additions between the coefficients of *a* and the parameters of *c* (**Fig. 4G**). Specifically, for each function *u* (∈ *{1*, …, *N}*) and each organelle index *i* (∈ *{1*, …, *n}*), we define the action of the *i-th* organelle of *c* on input *a* (relative to the *u-th* function), as the multiplication of the quantity *a*_*i,u*_ with the ratio *org(c)*_*i,u*_ */SK(c)*, This ratio quantifies iteratively the relative degree of specialization in the *i-th* organelle of *c* for the function *u* (**Fig. 4G, fig. S5D**):

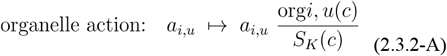

and where *SK(c)* is the sum of the components of *K(c)* (e.g. *K*_*1*_ *(c) + K*_*2*_ *(c) +* … + *K*_*N*_ *(c)*; **Fig. 4G, fig. S5D**) such that:

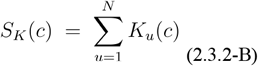

We then define the action of the *cell c* on the input *a* as the sum of actions done by each organelle:

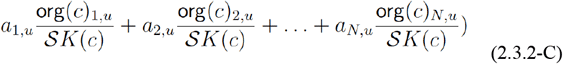

The cell-level action for function *u* sums organelle actions and is the vector of all actions (**fig. S5D**):

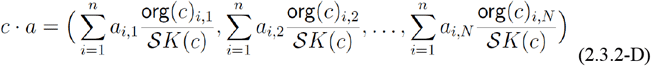

##### 2.3.3. Extension to trees of artificial cells

We extend *cell* actions to entire trees of *cells* by transferring the actions of *child cells* to their corresponding *parent cells* hierarchically (**fig. S5E**). The organelles of the *parent cell* act on the outputs from the *child cells* to generate the *parent cell’s* action (which becomes the new input to another *cell*, and so on). The tree structure ensures that the number of outputs (from *sibling cells*) matches the number of *organelles* in the *parent cell* (**fig. S5E**), ensuring structural and mapping constraints.

#### 2.4. Modeling optimization through nonequilibrium compartmentalization

Our mathematical framework uses actions on inputs (which are computed and propagated through a tree of *cells*) to fine-tune internal structural adaptations, yet it does not optimize specialization. Given our experimental observations on the recursive role of compartment formation and specialization (Biological observation/rule [4]-optimization through oscillation), we look for a theoretical framework that captures how recursive organization (the cyclical formation and dissolution of compartment) acts as a self-optimizing mechanism. In particular, where artificial cells could achieve specialized states by minimizing organization entropy (discrepancy between compartmentalized states), guiding structural evolution through nonequilibrium dynamics.

Drawing inspiration from Maxwell’s demon thought experiment (*85, 106*), where the action of a demon conceptually optimizes the system’s distribution and organization, we aimed at developing and implementing a nonequilibrium optimization principle that distinguishes between organization states (**Fig. 5A**). Specifically, we model [1] low entropy configurations as compartmentalized organizations and [2] high entropy configurations as non-compartmentalized, which, in our cellular experiments, linked function and specialization to the oscillation between these two configuration states (**Fig. 5A**).

**Fig. 5.**
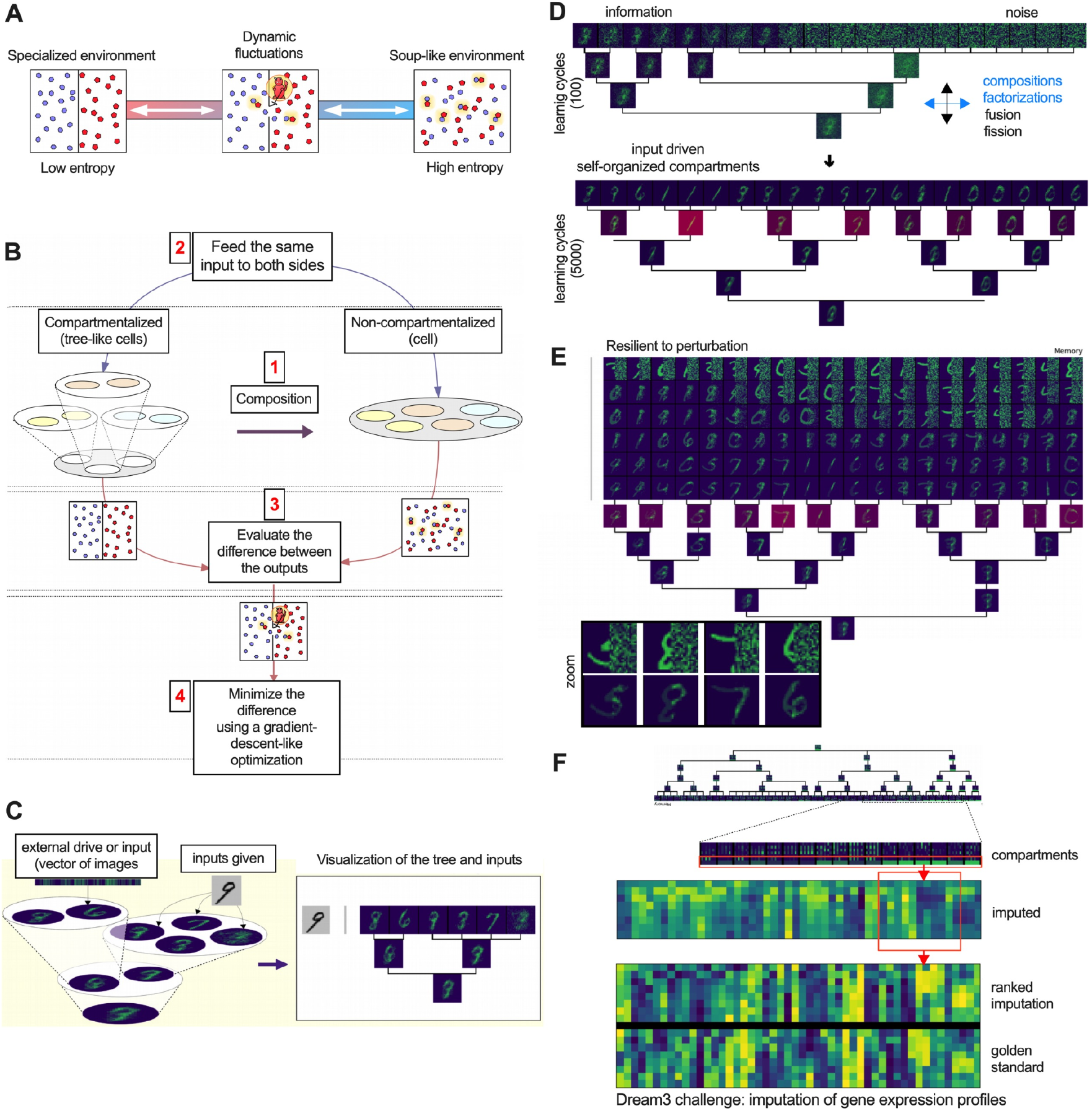
Optimization by a nonequilibrium-inspired loss function leads to self-supervised learning systems. **(A)** Illustration showing a conceptual parallel between Maxwell’s demon thought experiment and the dynamic properties of living systems. The panel shows three entropic states of a system, with the middle system representing a flexible configuration fluctuating between the other 2 states. **(B)** Schematic showing how each junction of a tree of cells is optimized during a 4 step procedure, based on compartmentalization logics and nonequilibrium tuning. First, each junction of the tree is turned into a cell by composing the child cells with the parent cell, as shown in step 1. The main differences between the junction and its composition is their respective levels of compartmentalization (the junction having one more level), which allows us to compare the effect of compartmentalization similar to Maxwell’s demon thought experiment. The two systems are compared by giving them the same inputs (step 2) and evaluating the difference between the outputs (step 3). A gradient-descent-like optimization deduces the changes to be made in the system in order to minimize this difference by using factorizations and compositions simulating formation and removal of compartments. **(C)** Tree representations and visualization of the memory of Intcyt. At every learning cycle, we give the tree of cells images such as the number 9 shown in black and white. **(D)** Development of Intcyt architecture on the MNIST dataset from a single cell to a hierarchical network of cells. State of the tree for the learning cycles 100 (upper panel) and 5000 (lower panel). During the development, Intcyt learns, abstracts and memorizes concepts from the inputs. The redder cells in the tree (at the bottom) are associated with relatively high values of the residual, which is equal to 1 + res(c)=SK(c). **(E)** Visualization of Intcyt outputs on MNIST images whose right sides are hidden with white noise. The panel shows the development of reconstructions and zoomed examples (lower subpanel) of the set of images by Intcyt for different compartmentalization parameters (we use ω = 0 and ω = 30, see Supplementary text, section 4b). **(F)** Extraction of learned information from the self-supervision of Intcyt on the DREAM3 dataset. Imputed gene expression, ranked expression and comparison with the golden standard provided with the DREAM3 challenge.

##### 2.4.1. Mathematical formulation

We formalized this principle mathematically using the canonical equation for operadic algebras (see supplementary text, section 4, part 1), and tested its application in computational systems. We conceptualized cell systems that combine both low- and high-entropy configurations to minimize the difference between the effects of their internal behavior in the presence or absence of compartmentalization (**Fig. 5A, B**).

We evaluate this difference by comparing the actions of trees of cells with different compartmentalized structures. Specifically, for a junction comprising a tree of *cells* made of a *parent cell c* and a collection of *child cells d = (d*_*1*_, …, *d*_*n*_), [1] we compared the junction action (on an input matrix *a = (a*_*1*_, …, *a*_*n*_); **Fig. 5B**, left side of the tree-cell; step [1]) with [2] that of the composed cell, *c* ◦ *d* on the same matrix *a* (**Fig. 5B**, right side of the tree-cell; step [1]) through the following operation: *U(c, d)(a)* : *= (c*◦*d)* · *a*−*c* · *(d*_*1*_ · *a*_*1*_, …, *c*_*n*_ · *a*_*n*_) (2+.4.1-A).

##### 2.4.2. Operadic algebra framework

The operation *U* is well-established in operad theory and is used to define “algebras of operads” through the equation *U (c, d)(a) = 0*, which must hold for every input *a*. In our context, exact solutions to such an equation would imply near-extreme specialization (*cells c* and *d*_*1*_, …, *d*_*n*_ with associated vectors containing zeros in all but one dimension).

We thus adopt a more nuanced approach by considering the equation *U (c, d)(a) = 0* as an ‘ideal situation’ that *cells c* and *d*_*1*_, …, *d*_*n*_ should converge towards. Consequently, we seek solutions *(c, d)* for which the scalar product *U (c, d)(a)* · *U (c, d)(a) = U (c, d)(a)*^*2*^ is minimized (as close as possible to *0*). The notion of converging toward a ‘*minimizing situation for a given environment*’ aligns with Maxwell’s demon principle, where finding the parameters that minimize the multivariate function *(c, d)* → *U (c, d)(a)*^*2*^ yields optimally organized configurations (**Fig. 5B**).

##### 2.4.3. Optimization strategies

In Supplementary text (part 1, subsection 1.6), we describe how to minimize the multivariate function *(c, d)* → *U (c, d)(a)*^*2*^ through two types of complementary optimizations:

[1] Numerical (gradient-descent-like) optimization: Our gradient-descent-like optimization differs from conventional gradient descent because we compute the differential of *(c, d)* → *U (c, d)(a)*^*2*^ on a manifold that is not in the Euclidean plane. Our differential is computed as an infinitesimal variation of the function *(c, d)* → *U (c, d)(a)*^*2*^ relative to an infinitesimal cytosolic exchange between the *cell* variables *c* and *d*. We then use the resulting differential to modify the parameters of the *cells (c, d)* to obtain a solution *(c’, d’)* with a smaller value *U (c’, d’)(a)*^*2*^. After complete re-parametrization, the tree may enter a non-homeostatic state such that any fitness disagreement is corrected iteratively through *cytosolic* exchanges between *parent cells* and *child cells*.

[2] Structural optimization (using fission and fusion events): In our structural optimization, we show how the value of the function *(c, d)* → *U (c, d)(a)*^*2*^ can be reduced by merging (fusion) or dividing (fission) the *child cells d*_*1*_, *d*_*2*_,…, *d*_*n*_. In the supplementary text, we give strategies to identify *cells* to be merged or divided as well as methods to compute the division of a *cell d*_*i*_.

Overall, our framework operates as follows: Interaction: a tree of *cells* interacts with external drives (inputs) through initial operations (actions) for forward propagation. Recognition: it recognizes input similarities. Self-organization: the system self-organizes and specializes by alternating between (1) numerical and (2) structural optimizations. The computational implementation is presented in the following sections. Importantly, unlike other graph-based formalism, our operadic framework carries action and state with explicit semantics using composition/factorization, which enables tree-consistent structural plasticity under external drives providing a mathematically closed space for structural learning.

##### 2.4.4. Diagnostics and interpretation inspired by the information bottleneck (IB)

Even though our optimization objective is not a formal implementation of the IB principle, it can be viewed through its lens as a diagnostic tool, given that it can offer an approximation for the information theoretic notion of optimality under compression and prediction trade-offs. In IB, the goal is to maximize predictive information while minimizing stored information, and trading compression against accuracy. In our framework, minimizing *U (c, d)(a)*^*2*^ under a thermodynamic budget (Δ *S*_total_ ≥ 0) can be viewed as an analogue trade-off that can be used as a learning ledger as it conserves predictive information while discarding irrelevant information.

Heuristically, (i) the *U*^*2*^ term promotes predictively-useful representations as a proxy, while (ii) the residual increments offer an audit track of irreversible updates, discouraging retention of non-predictive content or unneeded information-processing costs (because discarding state during learning requires irreversible operations, which can be viewed as entropy-like costs in the Landauer sense; see Supp. Methods M3). In addition, the operadic operations enable a form of structural compression via hierarchical abstraction, for instance when parent cells represent coarse-grained summaries of child cell information.

The process of learning carries a measurable cost: every time the system performs a logically irreversible operation (e.g. the zeroing out cytosolic content into a residual store, merge and discard, clearing of transient buffers) we treat this as information being erased from the active workspace. We track these events as a ledger and report them as a proxy in a Landauer-normalized dissipation (LND) manner, as bits-equivalent per irreversible operation from model signals (see Supp. Methods M3-M4 for a definition of the three event types and for the LND calibration; only for diagnostic purposes and no formal equivalence).

In classic IB, the goal is to find a representation Z which predicts relevant outputs Y with high fidelity and with the least amount of retained information about the input X. In our setting, the specialized action by the tree, namely the action by the tuned supercell in response to allostasis, plays a role analogous to a candidate representation Z (for the purpose of an IB-inspired diagnostic). We use standard variational bounds to estimate predictive information I(Z;Y) and the retained information I(Z;X). The difference I(Z;X)-I(Z;Y) is used as a proxy for non-predictive memory (Supp. Methods M6; a diagnostic without optimizing mutual information directly).

Empirically, decreasing *U*^*2*^ is coupled with an increase in estimated predictive information I(Z;Y) and a decrease of non-predictive memory proxy through our learning optimization (see next section). At the same time, operadic factorization reduces the number of active compartments and thus performs a structural type of compression: increased predictive power using less retained memory resources. The trade-off weight is not a forced ad-hoc hyperparameter but is provided as a budget by the empirically determined LND proxy, which can be viewed as bits per operation from the residual signals and is anchored in a physically motivated proxy instead of an arbitrary penalty (Supp. Methods M4, LND). We can then view our optimization as an IB-inspired interpretation where the effective trade-off weight emerges from information-processing constraints embedded in the learning objective. This perspective supports the use of IB-like diagnostics alongside dissipation ledgers as a step towards efficient learning by minimizing waste from irrelevant information (*107, 108*) (Supp. Methods (M1-7); a quantitative approach on dissipation and specialization, including entropy split, LND calibration, and IB-like diagnostics, appear later in section 3.7.2 and **Supp Method Figure. 1A-F**).

##### 2.4.5. Diagnostics and interpretation inspired by thermodynamic consistency and information-processing

After defining an optimization rule that promotes organization, we next ask whether this optimization can be made directionally consistent (at least as an explicit bookkeeping of learning) with basic nonequilibrium constraints. To do so, we introduced a relative cost-accounting (in the spirit of modern approaches to Maxwell’s paradox and the generalized second law statements (*57, 58, 86, 87*)), where, for example, compartment organization can minimize local disorder while information-processing steps (representation resets during updates) are assigned entropy-like proxies. In this framing, the accounted sum is non-decreasing while tracking irreversible updates, without stating that the proxy captures every flow related to physical entropy in a particular substrate.

We decompose the entropy-like accounting of a tree T as: *S*_total_ (*T*) = *S*_comp_ (*T*) + *S*_res_ (*T*) (2.4.5-A), where *S*_comp_ measures the disorder of the compartmentalized structure, which is decreasing with the increasing specialization, and *S*_res_ tracks cumulative entropy-production proxies from logically irreversible operations in the learning process, increasing when the system processes and organizes information (*57, 58, 86, 87*) (Supp. Methods M1; *S*_res_ is only intended as a conservative ledger-proxy tied to quantified irreversible events).

At each optimization processing step (composition *c* ◦_*1*_ *d*, evaluation of *U*, gradient-like update, homeostasis restoration) we usually observed the following: (i) *S*_comp_ tends to decrease by routing similar inputs to more specialized compartments (content distributions sharpen and local order increases), such that organization in the internal representation is approximating Δ *S*_comp_.< 0. (ii) *S*_res_ tends to increase because of irreversible operations at each cycle, implementing decision-making and memory updates (*86, 87, 107*). Thus, each decision has processing cost-like proxies and are conceptually relevant steps for Landauer-type lower bounds (see Supp. Methods M3). We then monitor a generalized second-law balance as a consistency check, Δ*S*_total_ = Δ *S*_comp_ + ≥ *S*_res_ ≥ 0 (2.4.5-B), such that when specialization takes place (Supp. Methods M2), the rise in the residual ledger keeps the accounted changes non-negative:

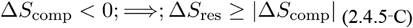

Connection to Landauer limit (as a lower-bound proxy): Landauer’s principle states that, in an ideal setting, the minimum entropy cost for erasing Δ *H*_erase_ bits is *k*_B_*T* ln 2 · Δ*H*_erase_ as heat dissipated (a minimum entropy increase *k*_B_*T* ln 2 · Δ*H*_erase_ (*86*)). In our discrete operadic updates, the system implements such erasure events (conservative count of information loss) with residual increases serving as the corresponding ledger increment. We report this in LND units (bit/irreversible operations) calibrated from event logs and stress-like signals (Supp. Methods M1-4). Importantly, we do not interpret LND-proxy or claim it as absolute Joules; the goal is to allow internal consistency check and cross-reference comparisons (further theoretical and empirical support is presented in Fig. 5-6, Table 1, section 3.7.2., and **Supp. Method Figure. 1A-F**).

**Table 1.** Diagnostics: Sign-level closure. Arrows indicate direction relative to baseline (cells: 5.5 mM glucose; Intcyt: early learning). Dissipation proxies are pH-fluctuation amplitude in cells or residual/stress indices from logically irreversible updates in Intcyt. Specialization readouts include H3K4me3 at mito-HAR promoters, gene activity within mito-HAR hubs, OCR, FRAP recovery, duration of specialization (cells) and consolidation of compartment, while computationally we minimize the U^2^ operation for specialization between early baseline learning and self-organizing learning (SOL) phases.. Closure marks matched sign-level changes (↑/↑ or ↓/↓). For intermittent glucose deficit (IGD) and ATP12A^−^/^−^ + IGD, low glucose (LG; 12 h), dissipation arrows indicate mechanism plus observed metrics. *Calibration note*. LND is treated as a lower-bound, proxy-based metric; Supp. Method section details LND-struct (counting irreversibilities) and LND-signal (stress-derived estimates), while cellular LND uses cytosolic pH amplitude as a proxy. See Table S2 in supplementary materials for ablation experiments.

| System | Condition / Perturbation | Dissipation proxy | Specialization readout | Functional specialization | Closure |
| --- | --- | --- | --- | --- | --- |
| Cells | Control (glucose) | — | — | — | — |
| Cells | Low glucose (LG) | ↑ | ↑ | ↑ | ✓ |
| Cells | vATPase-RNAi (LG) | ↓ | ↓ | ↓ | ✓ |
| Cells | Tmem175-RNAi (LG) | ↓ | ↓ | ↓ | ✓ |
| Cells | ATP12A-RNAi (LG) | ↑ | ↑ | ↑ | ✓ |
| Cells | IGD (pre-subculture) | ↑ | ↑ | ↑ | ✓ |
| Cells | ATP12A-/- + IGD | ↑↑ | ↑↑ | ↑↑ | ✓ |
| Intcyt | Early (baseline) | — | — | — | — |
| Intcyt | Learning | ↑ | ↑ | ↑ | ✓ |
| Intcyt | ablation | ↓ | ↓ | ↓ | ✓ |

##### 2.4.5a. Learning vs dissipation trade-off: TUR-inspired heuristics

Thermodynamic uncertainty relation (TUR) establishes a theoretical connection between driven currents and their entropy production. Our operadic learning dynamics are measured through proxies, so we use a TUR-inspired heuristic diagnostic to motivate a qualitative trade-off (not a formal TUR bound for the model). Our diagnostic refinement asks whether there is any correlation between the precision of learning and the dissipative cost measured.

From our entropy decomposition approach (eq. 2.4.5-A,B,C), we derive an approximation consistent with the TUR-style bounds (*109, 110*) (fundamental trade-off between learning speed and energy cost), which is supported by empirical information (cross-reference: in the next sections we show that periods of specialization match larger residual increments). Let the scaling (specialization rate)^2^ · (entropy production) ≥ C (2.4.5-D), where specialization rate ≈ | Δ*S*_comp_|/Δ*t*, and the proxy is residual accumulation (in our computational model) or pH-fluctuation amplitude (in the cell systems). In each learning window we define a learning current J_t_ as a signed measure of improvement, which is associated with -Δ*U(c, d)(a)*^*2*^ or a reduction in *S*_*comp*_. For each window we estimate the mean,*μJ* variance 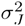 and the dissipation ∑_*w*_ expressed in the number of bits using LND. TUR-style ratio is then reported in the form of:

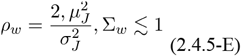

For stationary driven dynamics satisfying TUR-style diagnostic assumptions, this ratio is expected to be <1, values of > 1 hint on underestimation of erasures or presence of non-stationarity (e.g., regime shifts). In our setting, we presented ratio values as a diagnostic of consistency, and not as an indicator of bound saturation. When the computation enters an oscillatory window, learning-per-bit diagnostic is well behaved, and when we disrupt the drive, *ρ*_*w*_ increases, which is consistent with a mixture of proxy mismatch and uncontrolled window transients (**Fig. 6A–D, F–I**). We therefore treat this TUR-style behavior as a supporting heuristic check supported by our data. Of importance, a formal bound saturation or a formal theorem for operadic dynamics is left to future work (see Supp. Methods M4, M7, and section 3.7.2).

**Fig. 6.**
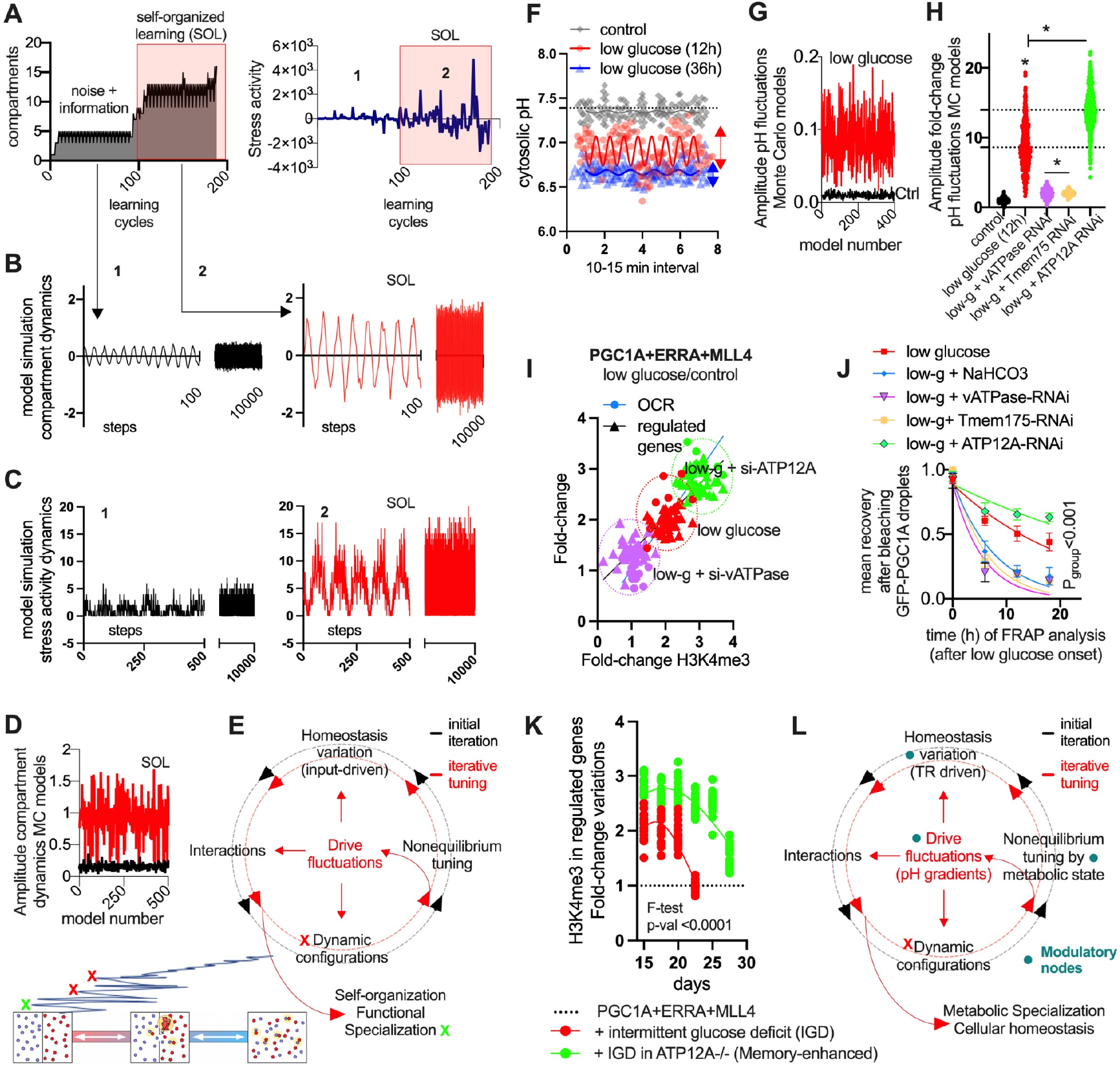
Nonequilibrium fluctuations enhance specialization and homeostasis. **(A)** Intcyt parameters throughout learning cycles. Left, number of compartments over time. Right, stress-activity trace (see supplementary text). Red box self-organized learning (SOL) phase. Model simulation (nonlinear sine-wave models) of compartment **(B)** and stress activity **(C)** dynamics throughout learning cycles. Left, low-amplitude steady-state transitions. Right, show self-organization with increased amplitude of fluctuations. **(D)** Comparison of fluctuation amplitudes in early states of learning (black) versus SOL phases (red) from 500 Monte Carlo models using parameters identified in **B. (E)** Schematic of dynamic specialization steps. Input-driven variation triggers interacting behavior of the system and dynamic partial configurations (red X, analogous to local minima in optimization). These are tuned by a nonequilibrium-inspired loss function, which leads to conformational transitions, and iterative fluctuations, promoting self-organization and specialization (green X). **(F)** Time-series (every 10 to 15 min) of cytosolic pH variation in SCAT-MSCs after 12 hours or 36 hours in low glucose (control 5.5 mM, low glucose is 2 mM). Red and blue arrows show high and low variance oscillatory behavior. Nonlinear sine-wave models were fit and used for Monte Carlo (MC) modeling. **(G)** Amplitude variation of MC models from dynamic variation of cytosolic pH (as in **F**) for control or low glucose conditions. **(H)** Mean amplitude variation of pH-fluctuations models as in **G**, from SCAT-MSCs with or without RNAi for proton pumps. **(I)** Fold-to-fold change plot for expression of mitochondrial-HAR hub genes (Y-axis) and basal oxygen consumption rate (OCR, y-axis). In X-axis H3K4me3 variation levels after low glucose (12h). SCAT-MSCs overexpressed PGC1A, ESRRA, and MLL4 (in combination) with or without RNAi for proton pumps. **(J)** FRAP analysis of PGC1A droplets after the onset of glucose deficiency in SCAT-MSCs (X-axis). At each time-point, mean droplet recovery (10 seconds post-bleaching) in different conditions: control (5.5mM glucose), low glucose (2mM), low glucose with NaHCO3 (1mM), and with RNAi for proton pumps. **(K)**, Time-course H3K4me3 fold-change in mitochondrial-HAR hub promoters in SCAT-MSCs after combined overexpression of regulators, with intermittent glucose deficit (IGD) alone or in ATP12A-/-MSCs (ME; Memory-enhanced MSCs; mean fold change in promoters of at least 20 promoters per group). **(L)** Illustration for metabolic function specialization. Dynamic configurations from interactions between transcriptional regulators (TR) and transcriptional compartments components. Dynamic behavior of transcriptional droplets trend towards arrested materials as they age (*24*) (red “X”: arrested dynamic is analogous with local minima). Metabolic state variation (nutrient shortage) through proton accumulation (byproduct of catabolism) and pH gradients (through lysosome activity) generate asymmetric fluctuating drives that modulate transcriptional compartments and enhance cell specialization. For all panels, cell experiments were done with 3 independent replicates. Data show mean values and SEM. Unpaired two-tailed student’s t-test for comparing two groups, and ANOVA fisher’s least significant difference (LSD) for multiple groups comparisons. Two-way ANOVA was used to estimate significance between groups constrained by time-based measurements, and Tukey test for multiple comparisons. Nonlinear regression (Lorentzian-Cauchy model) and extra sum-of-squares F test to compare models between conditions. * indicates p-value <0.05.

### 3. Adaptive compartmentalization leads to self-organizing machine learning systems

#### 3.1. Computational nonequilibrium optimization

Current ML systems treat learning as an optimization decoupled from physical constraints (*57, 87, 107, 108*). This approach inevitably leads to systems that, even though powerful (such as deep learning) are thermodynamically inefficient, with fixed architectures, and are prone to catastrophically forgetting learned information. This suggests the need for new computational frameworks that can implement learning with physical thermodynamic constraints, where, for example, architecture emerges from data itself and it is not just being shaped by preconceived designs. In contrast, biological systems dynamically adjust their ‘architecture’ (compartment organization) in response to inputs while remaining thermodynamically constrained. We therefore asked if it is possible to take mathematical formalizations of cellular adaptive principles to build computational systems with adaptive architectures and thermodynamic consistency.

We developed a computational framework (intcyt) that uses nonequilibrium tuning of compartmentalization as a self-organizing machine learning system (**Fig. 1B**, “steps 9, 10,” supplementary text, part 2). Conventional machine learning optimizes abstract losses, while Intcyt, optimizes a physically constrained objective where architecture emerges through fission/fusion operations under an explicit entropy budget. It shows different capabilities such as thermodynamically compatible at the level of our accounting, adaptive architecture, and adaptive learning. Notably, these properties are not engineered separately but instead emerge directly from our mathematical framework, showing that physical principles can guide the design of novel machine learning architectures.

Specifically, Intcyt uses a tree of *artificial cells* with organelle vectors that learn and self-organize data inputs from datasets (e.g., MNIST, fashion-MNIST, gene expression data). Our goal was to show that cell-inspired out-or-equilibrium principles can be formalized, computed, and lay the foundation for physical adaptive ML, while providing a scalable optimization toolkit for further development aimed at the integration of our approach with more advanced AI architectures.

#### 3.2. Learning mechanism and adaptive evolution

For every input and junction *(c,(d*_*1*_,…,*d*_*n*_)) in the tree, we use [1] a gradient-descent-like algorithm together with [2] factorization and composition operations to update organelle memory. The information learned by the tree is stored in the organelles of each *cell* and, in a homeostatic state, the information in the *organelle* of each *parent cell* equals the sum of the information contained in their *child cells* (**Fig. 5C**). Intcyt begins with a single *cell* (receiving data inputs) and adaptively evolves into a tree of *cells* via fission and fusion operations, consisting of iterative factorizations and compositions (**Fig. 5D**). Importantly, if the *root cell* of a tree has no *parent cell* for division, a fission operation consists only of a factorization step since the complementary composition step would require a *parent cell*.

This asymmetric use of compositions and factorizations at the root cell leads to the generation of new compartments, making it critical for further optimization. Elsewhere in the tree, *cells* merge and divide symmetrically, enabling exchange of organelles. Notably, every fusion and fission of cells minimizes the algebra operator *U* (see optimization strategies from previous section), which optimizes learning locally and globally by merging *cells* with similar information and dividing *cells* with distinct information into different compartments. Altogether, these dynamic mechanisms lead to structural reinforcement of learning and information self-organization (see supplementary mathematical section).

#### 3.3. Information organization and abstraction

Our computational framework organizes its architecture by integrating external inputs in a hierarchical and adaptable database using composition operations. The system then separates information from noise, merges organelles with similar information, and reinforces learned information across higher tree levels (**Fig. 5D, fig. S6A, B**). As some organelles become more representative of the cell’s content, the *cell* containing them develops a preference towards these functional specializations (**fig. S6C**). Notably, our framework creates higher-level abstractions in *parent cells* from lower-level learned and self-organized concepts (e.g. the *child cells*) by enforcing homeostatic evaluations. This process takes place during gradient-like descent updates, and it is further optimized by managing cytosolic contents at every learning cycle (residual information accumulation).

#### 3.4. Managing residual cytosolic information

Throughout gradient-like descent and operadic optimization, we fine-tuned this process by the iterative removal of residual cytosolic content. To prevent memory collapse and Intcyt from converging onto empty vectors during these removals, we eliminate these contents in a way that preserves their numerical content (no numerical loss). Specifically, for each *cell c*, the vector *cyt(c)* is made of either negative or non-negative values, reflecting previous exchanges. If the sum of *cyt(c)* components (*Scyt(c)*) equals zero, then turning the vector *cyt(c)* into a zero vector does not create any numerical loss in the system. Conversely, if *Scyt(c)* is non-zero, we prevent numerical loss and memory collapse by adding *Scyt(c)* to the residual *res(c)* as follows: *Scyt(c)* = *e*_*i1*_ + … + *e*_*in*_ − *e*_*j1*_ − … − *e*_*jm*_.

#### 3.5. Residuals as cost indicators (dimensionless)

Residual arises from logically irreversible updates (see *2.4.4. section*) that are viewed as dimensionless lower bound proxy for learning stress (work performed by the system, monitoring progress). This is intended as a relative cost signal to compare conditions and events in the model, while also being considered a metric of computational effort to create structure. While local compartments decrease disorder through specialization, residuals accumulate elsewhere in the system, consistent with processing constraints. This interpretation is compatible with Landauer-style arguments (*86, 87*), that irreversible operations carry a thermodynamic cost of information processing, offering a principled approach to track or discard events during learning.

As shown in our empirical observations, residuals indicate specialized compartments (redder organelles; work was performed; **Fig. 5D, fig. S6D**, supplementary text, section 4, part 1, subsection 1.5). The conversion of cytosolic content *cyt(c)* can be viewed as a byproduct reaction of the form e _i 1_ + … + e _i n_ → e _j 1_ + … + e _j m_, releasing dissipated work equal to *res(c)*. In Intcyt, cytosol zeroing with transfer to res(c) and merge-and-discard during fusion are, again, logically irreversible updates. So, the transformation provides a calibrated lower bound on entropy production, which is associated with the computational decisions of structurally learning inputs into specialized compartments (**Fig. 5B-D and fig. S6D**).

Interestingly, we can draw parallels where the total entropy of the system (compartments + residuals) remains consistent with the second law: The Residual accumulation pattern, monotonic increase during self-organization learning, supports Δ*S*_comp_ < 0; ⇒; Δ*S*_res_ ≥ | Δ *S*_comp_| (3.4-C). Then, the conversion of cytosolic content evokes parallels related to the Landauer-irreversible erasure, where, as compartments organize Δ*S*_comp_ < 0 (organization/specialization), residual increases Δ*S*_res_ > 0 (processing cost), maintaining: Δ*S*_total_ = Δ*S*_comp_ + Δ*S*_res_.> 0

The internal consistency check in our framework is designed such that increased specialization (decreased local entropy) is accompanied by increased residuals, providing an intuitive way to monitor the cost of organization (**Fig. 5D, fig. S6D**). Calibration to Landauer units (*k*_*B*_ ln 2per erased bit as relative bounds to phase/condition) is described in supplementary methods (section M); here we report normalized residuals to compare learning phases without claiming absolute energy (see Discussion). Yet, this consistency check distinguishes our framework from current paradigms, which aim at optimizing loss functions without accounting for processing constraints. Intcyt, on the other hand, shows that inherent processing constraints can enable adaptive learning architectures.

#### 3.6. Integration with machine learning paradigms

Intcyt’s transformation of information and restructuring works together with the gradient-like optimization to minimize the algebra operator *U* values, which enhances the learning specialization in organelles with similar inputs (within inputs dimensions) and decreases the specialization in others (dimensions with less similarity).

##### 3.6.1. Nonequilibrium tuning

Empirically, reactions associated with residuals ensure that losses or gains in compartment specialization have a counter-effect on others, which creates both an out-of-equilibrium feedback effect that fosters competition between compartments and prevents the system from settling into suboptimal configurations. This competition results in the retention of specialized compartments for the inputs, leading to their gradual memorization and abstraction when repeated iteratively. Gradually, this selective mechanism promotes adaptive and evolutionary-like structural rearrangements, with memorized and abstracted information across hierarchical nested scales.

##### 3.6.2. Interpretability

Importantly, cooperation and communication between organelles improves the learning and the structural representation specific to each dataset, which in turn helps prevent overfitting (**fig. S6C, D**). Gradient descent increments were coupled with learning policies to improve specialization (supplementary text, section 4d). Integration of policies were possible thanks to the easy interpretation of memorized information by organelles, whose vectorized content fully represent data inputs displayed in the figures (images or numerical, **fig. S6C, D**). In addition, since every organelle vector represents data inputs, no preprocessing is needed for querying, and assessing information is straightforward.

##### 3.6.3. Performance validation

Besides fast learning, structural representations, and abstraction from datasets, the robustness of Intcyt was supported through data reconstruction tasks. In only a few learning cycles, the system reconstructed data with missing information from MNIST, fashion-MNIST (visual pattern completion, **Table S2**), and the DREAM3 challenge (*111*) (gene expression imputation; **Fig. 5E, F, fig. S7A, B**). Baseline results (K-NN, linear SVM, random forest, 2-conv CNN) are representative published benchmark values on MNIST and Fashion-MNIST with standard splits and raw-pixel inputs. Intcyt values are reported as mean ± SD over independent runs with different random seeds/data orders (see **Table S2**). For DREAM3 data, Intcyt fully recovered partially-hidden gene expressions outperforming other algorithms in the challenge (**Fig. 5F, fig. S7C**). Intcyt integrates hierarchical clustering, data reconstruction, and adaptive architecture discovery using principles from our observations on how nonequilibrium tuning of compartments leads to specialization. Notably, our optimization principle results in an adaptive, self-supervised machine learning system. In the next section, we evaluate modulatory behaviors in Intcyt to identify fundamental relationships between fluctuation dynamics and specialization.

#### 3.7. Nonequilibrium fluctuations modulate specialization

To investigate modulatory behavior linked to specialization in Intcyt, we evaluated parameters throughout the learning process (**Fig. 1B** “step 10”). We found that compartment formation and disassembling, stress activity parameters, and residual accumulation increased in tandem during self-organizing behavior (**Fig. 6A, fig. S8A**). We also observed that specialization is associated with asymmetric fluctuations in compartment transitions, prompting us to investigate these patterns using nonlinear models and Monte Carlo simulations (**Fig. 6B-D**).

##### 3.7.1. Fluctuation dynamics

We found that the amplitude of fluctuations grows as self-organizing behavior and accumulation of residuals increase. To estimate this, we analyzed compartmentalization and stress activity dynamics during learning cycles. During early learning cycles we observed: compartments: 1–5 (mean 4.0\pm1.1); compartment oscillation amplitude; 1.6; Stress-activity s.d.: 279 (model units); entropy-production proxy (|stress|/cycle): 150 units/cycle. During self-organization cycles we observed: compartments: 7–16 (mean 11.3\pm2.2); compartment oscillation amplitude: 3.4; stress-activity s.d.: 1,067; Entropy-production proxy: 750 units/cycle. Fold-change variations between SOL vs early was: compartment oscillation amplitude: 2.1×; stress-activity fluctuation amplitude (s.d.): 3.82×; entropy-production rate (|stress|/cycle): 5.0× (**Fig. 6A-D**). These increases show that SOL is associated with amplified nonequilibrium fluctuations (Monte-Carlo simulation shows a 3.4× amplitude increase; **Fig. 6D**).

##### 3.7.2. Processing cost interpretation

The amplitude of fluctuations increases with residual accumulation in the system. This suggests that the system’s work locally (residuals) can guide the asymmetric use of compositions and factorizations as a drive for specialization (**Fig. 6E**). This relationship supports TUR-style heuristic summaries and diagnostics (equation 2.4.5-D) where larger organizational gains (specialization) requires greater processing costs (expenditure).

Mechanistically, Intcyt can be viewed as using nonequilibrium tuning to reduce time spent in partially-fit configurations (analogous to local minima in conventional optimization; **Fig. 6E**). This tuning includes: [1] taking in external drives or inputs into compartments; [2] promoting dynamic operations to recognize input similarities; [3] irreversible updates that generate residual; [4] structural adaptation via compartmentalization; [5] an operadic gradient-descent loss function for global optimization by reducing the manifold *U*^*2*^ (supplementary mathematical framework, part 1, subsection 1.5).

Calibration, accounting, and dissipation definition: We quantify dissipation proxies similar to cumulative informational residual gained during structural updates (akin to Kullback-Leibler divergence (*108, 112*)). This serves as a theoretically grounded proxy for the Landauer cost (irreversible work required to discard non-predictive states; see section 2.4.4). We provide the quantitative accounting of processing-cost proxies and calibration in supplementary methods Fig. 1A-D, where we show phase segmentation by compartmentalization threshold including cost parameter decomposition, stress-to-bits slope, and LND (bits per operation; see Supp. Methods M4-7 methods, including rectification/efficiency analysis and next section).

##### 3.7.3. Asymmetry and rectified fluctuation promotes self-organization

The behavior of the system indicates that our optimization leads to specialization through asymmetric fluctuations, suggesting a transition regime during adaptive optimization. Conceptually, this pattern is reminiscent of broken-symmetry transitions in other driven systems linked to the emergence of adaptive behaviors (*10, 113, 114*). This process exhibits an initial state of uniform symmetry that spontaneously breaks into a locally organized domain of specialized compartments with low-symmetry when a parameter crosses a threshold. The control-like parameter in Intcyt is residual accumulation, a record that co-varies with the transition of organizational work (here used as a diagnostic ledger; not physical control).

We also observed that the amplitude of fluctuations increases with the accumulation of residuals, and not during random exploration. This temporal ordering differentiates purposeful adaptation from noise-drive drift or random drive. The accumulation of byproducts after work can be viewed as analogous to how low-energy states in cellular models promote energy expenditure and byproducts. Similar to how epigenetic marks record cellular activity, these byproducts of computation (residuals) serve as memory-like traces that bias structural adaptation towards configurations that recognize similar inputs. In other words, they modulate self-organization leading to structural specialization (green X in **Fig. 6E**). This suggests that nonequilibrium dynamics may offer a handle signature for efficient adaptation, in line with previous observations (*10, 113, 114*).

Under this interpretation, the coupling between work and exploratory amplitude of fluctuation creates a reinforcing feedback loop characteristic of self-organizing adaptation (*7, 10, 113, 114*). Early specialization produces residuals, residuals amplify fluctuations, and this magnitude difference enables larger structural rearrangements and specialization. The resulting outcome is associated with the acceleration towards optimally specialized architectures. Despite these features known to occur in living systems as they exploit broken-symmetry configurations and dissipative states to find optimal adaptations with low energy requirements (*10, 113, 114*), we treat these as model-specific insights on adaptation.

Interestingly, adaptive systems rarely organize by pushing a continuous uphill battle against entropy. In fact, it has been proposed that they harvest internal variations, oscillations, gradients, and mechanics inherent to dynamic systems (*7, 62*). In agreement with previous studies (*7*), we propose that asymmetric fluctuations may function as a controllable drive that can be rectified within the system for specialization, converting alternating and low-grade variation into net organizational gains (**Fig. 6E;** see section 6.3).

In this perspective, intermittent windows of optimal adaptation created by a fluctuating variable, periodically lower effective barriers for key operations. What we observed here is that when those operations are phase-aligned to these optimal windows, the system performs more productive reconfigurations per unit of activity than under tonic forcing. In our computational instances, internal fluctuation amplitude gates structural reconfigurations (fusion/fission, consolidation), and when amplitude peaks, the system tests higher impact rewirings (*7, 71*). When it troughs, it then stabilizes useful changes and configurations. Repeatedly, these rectified updates steer the architecture toward partitions that best capture input structure, without steady forcing (**Fig. 6E;** see section 6.3). Quantification: the conversion of drive to organization is summarized by the DOE (rectified organization efficiency), estimated as the slope of organization change versus dissipation proxy during SOL. We show TUR-style ratios as diagnostics of internal consistency and non-stationarity (not claim of bound saturation), which remains within expected ranges and indicates a compatible learning-to-dissipation trade-off instead of under-measured information loss (Supplementary methods M5 and M7 and **Supp. Method Fig. 1E, F**). This is supported by empirical ablation of fluctuations, structural updates, and residual accumulation, which impaired learning in Intcyt (**Table S2**), consistent with specialization driven by structural adaptation instead of numerical refinement alone.

This conceptual framing can enable testable predictions: (i) intermediate amplitude minimum where very low and very high amplitudes can be suboptimal as they are either insufficient to cross barriers or wasteful to be impactful. Then an intermediate window maximizes organization gained per unit cost (*68, 115*); (ii) phase alignments with fluctuations and structural operation may modulate net gains; (iii) oscillatory drives may provide greater organization per unit cost than sustained forcing as they concentrate work when the system is most responsive (*76, 110, 116–118*). These quantitative relationships offer a computational benchmark, where amplified fluctuations can drive efficient specialization. These testable hypotheses are supported by proxy-based analyses, but the intuition can be generalizable; fluctuation offers a shove, rectification is the net step, while specialization is the staircase. This also raises a question as to whether living systems use similar modulators to tune specialization through fluctuating drives.

### 4. Identification of analog bio-modulators of phenotype specialization

#### 4.1. In vitro bio-modulators enhancing phenotype specialization

In this section we aimed at investigating whether analog drives of nonequilibrium dynamics affect transcriptional compartments and phenotype specialization (**Fig. 1B** “step 11”). Our earlier experiments revealed that glucose level oscillation is crucial for specialization and that this is directly linked to cytosolic pH variations (**Fig. 3C**). In line with previous work (*89*), both substrates have been linked to several gradients occurring in low-energy states (*43*), influencing both transcriptional and metabolic adaptations (**Fig. 3**). Notably, increased activity of lysosomal and membrane proton pumps has been linked to the generation of proton gradients (*43*), which are crucial for intracellular crowding, protein macromolecular assembly, and cell survival (*22, 119, 120*). Similar to Intcyt behavior, whether pH fluctuations (not merely static acidification) are a drive of cellular specialization with proton pumps as potential modulators remain unknown.

##### 4.1.1. Glucose deficit induces amplified pH oscillations

We hypothesized that proton pump activity during glucose deficit generates pH fluctuations (temporal oscillations around a mean acidic state) analogous to Intcyt compartment fluctuations. This led us to examine cytosolic pH variations and their overall impact on transcription, epigenetic plasticity, and phenotype adaptation. We tracked cytosolic pH variations in MSCs every 10-15 mins in control and glucose deficit states (12 and 36h), followed by modeling temporal variations using nonlinear sine-wave and Monte Carlo (**fig. S8B**). This revealed increased cytosolic pH fluctuations in 12-hour glucose deficit compared to control conditions and 36-hour glucose deficit. Modeling of fluctuation dynamics showed increased amplitude of pH variation in 12h glucose deficit (**Fig. 6F-G**). This transient adaptation suggests active regulation rather than passive buffering, potentially implicating the involvement of proton pumps.

##### 4.1.2. Proton pumps differentially control pH dynamics

We screened the expression of 15 proton pump genes during glucose deficit states. This showed significant upregulation in the vATPase, Tmem175, and Atp12a pumps (**fig. S8C**). These pumps have distinct locations and functions: the vATPase controls H+ influx to lysosomes and is commonly associated with lysosomal activity and pH balance (*43, 44, 121*); the TMEM175 pump mediates lysosomal H+ efflux (*45*) and is linked to lysosome function and cellular pH balance (*43, 44*); the ATP12A, on the other hand, is a cell membrane proton (efflux) and potassium (influx) channel (*122*).

To test their functional roles, we performed siRNA knockdowns in MSCs co-expressing PGC1A+ESRRA+MLL4 under glucose deficit. We found that the expected epigenetic H3K4me3 marks at mito-HAR hubs are lost by vATPase (Atp6v1b1 subunit) and Tmem175 channel knockdowns, but are enhanced by Atp12a channel knockdown (**fig. S8D-F**). Thus, lysosomal pumps (vATPase, TMEM175) are needed for epigenetic adaptations while the membrane pump ATP12A inhibits this process, suggesting opposing roles in pH regulation, with the ATP12A exhibiting a maladaptive role for molecular outcomes (**fig. S8F**).

##### 4.1.3. Pumps determine pH fluctuation amplitude

To assess their role on pH fluctuations, we repeated pH time-series measurements in perturbed cells and quantified fluctuation amplitude via Monte Carlo modeling (**Fig. 6H**). This showed that, in low glucose (12h), the amplitude of pH fluctuations is lost with vATPase-RNAi and Tmem175-RNAi, while it is substantially amplified with Atp12a-RNAi (**Fig. 6H**). This data revealed a clear relationship in which pH fluctuation amplitude predicts specialization capacity.

Proton pumps that reduce pH fluctuations (vATPase/TMEM175 knockdown) abolish H3K4me3 elevation, while pumps that amplify pH fluctuations (ATP12A knockdown) enhance it. This parallels Intcyt adaptation, where computational perturbations that suppress fluctuations impair learning while those amplifying it accelerate learning. Yet, both systems show similar organizational logic, with asymmetric oscillations between states enabling efficient optimization dynamics rather than static configurations.

##### 4.1.4. ATP12A causes potassium-dependent pH dysregulation

The opposing effects between the pumps suggested they act through different molecular mechanisms. To understand ATP12A inhibitory role, we evaluated lysosomal response in cells under glucose deficit. In MSCs with low glucose, vATPase and Tmem175 knockdowns led to reduced lysosomal activity and excessive cytosolic acidity, while Atp12a knockdown had the opposite effect (**fig. S8G, H**). This suggests that lysosomal vATPase and TMEM175 pumps are linked to proton flow and pH balance, while ATP12A impairs lysosome function.

Interestingly, ATP12A overactivity has been linked to potassium excess in the cytosol, which influences electrochemical gradients and proton movement, impairing both lysosome function and pH homeostasis (*45, 123*). We therefore tested whether potassium excess and ATP12A influence lysosome function during nutrient glucose deficit. Exogenous potassium treatment in glucose-deficient MSCs reduced lysosomal activity and increased cytosolic acidity, similar to vATPase and Tmem175 knockdowns (**fig. S8G, H**). Critically, Atp12a knockdown rescued these effects (**fig. S8G, H**), supporting that Atp12a channels mediate potassium dependent dysregulation.

In line with this, extracellular potassium levels reduced progressively during glucose deficit (larger reductions in 36h>12h>6h) (**fig. S8I**), indicating cell uptake. This was associated with increased time-dependent expression of ATP12A (24h>12h; **fig. S8J**), correlating with potassium accumulation. ATP12A overexpression worsened potassium toxicity (reduced lysosomal activity and cytosolic pH), while TMEM175 overexpression rescued these effects (**fig. S8K, L**). These results suggest that ATP12A-mediated cytosolic translocation and accumulation of potassium during persistent low glucose states affects lysosome function and pH homeostasis (**fig. S8M**), indicating that ATP12A acts as a maladaptive brake on pH oscillations.

##### 4.1.5. pH fluctuations tune transcriptional compartment dynamics

Having established that specific pumps control the amplitude of pH fluctuation, we tested whether this impacts transcriptional mechanisms such as droplet viscosity, MLL4 recruitment, and H3K4me3 epigenome writing. In MSCs expressing PGC1A+ESRRA+MLL4 and in low glucose conditions, Atp12a-RNAi enhanced epigenetic H3K4me3 marks and gene expression in mito-HAR genomic hubs, as well as mitochondrial function (**Fig. 6I**). vATPase perturbation blunted adaptations during low glucose (**Fig. 6I**).

We previously showed that intermediate droplet viscosity was associated with MLL4 recruitment and epigenetic activity during nutrient deficit (**Fig. 2, 3**). We hypothesized that proton flow influences droplet plasticity. FRAP analysis at different time points after the onset of glucose deficit revealed that PGC1A droplet viscosity increased with time (**Fig. 6J**). At 6h droplets remain liquid, at 12h they reach intermediate viscosity, and at 36h they are gel-arrested droplet viscosity (**Fig. 6J**), suggesting a temporal window of optimal dynamics at 12h, precisely when pH fluctuations peak (**Fig. 6F**).

In agreement, dynamic droplet recovery was lost by pH correction (**Fig. 6J**), vATPase-RNAi, and Tmem175-RNAi (**Fig. 6J**), supporting the need for lysosomal activity and pH dynamics to preserve droplet dynamics and plasticity. Notably, Atp12a-RNAi conditions maintained to a greater extent intermediate droplet recovery after 12 hours in low glucose conditions (**Fig. 6J**). In agreement, we found that the increased amplitude of pH fluctuations in Atp12a perturbation was directly associated with intermediate droplet viscosity, increased PGC1A-MLL4 protein interactions, and enhanced epigenetic activity (**fig. S9A-C**). As expected, these links were lost by vATPase perturbation and in cells expressing the IDR2-mutant PGC1A (**fig. S9A-C**).

We next tested whether ATP12A modulation enhances acute transcriptional memory formation in our models. Notably, low glucose in combination with Atp12a knockdown in transfected MSCs increased substantially the reactivation of primed transcriptional states and transcriptional memory, in contrast to vATPase perturbation and nutrient stress alone (**fig. S9D**). These results suggest that increasing pH fluctuations in nutrient deficit by ATP12A pump perturbation enhances transcriptional plasticity of regulators, including material properties, epigenetic recruitment, chromatin modification, and memory encoding. This further indicates that the amplitude of drives during the initial priming phase determines the memory strength of the reactivation phase. This parallels the learning dynamics in our computational model where larger, controlled fluctuation improves model robustness. Biologically, the question remains whether these effects translate to sustained specialization and organismal-level adaptation.

##### 4.1.6. Intermittent metabolic stress extends epigenetic memory

We previously showed that metabolic specialization is tight to metabolic state oscillations in our culture system (**Fig 3**). Namely, glucose level and pH variation during culture passages created repeated stimuli that progressively enhance transcriptional plasticity of regulators controlling the sustained specialization of mitochondrial function. Rather than treating this as an artifact, we pushed deeper amplitude of variations by reducing even more glucose levels during subculturing days.

Given that biological organisms experience rhythmic metabolic states (diurnal fasting/feeding, exercise-induced metabolic stress followed by recovery, and variations in food availability) we hypothesized that if oscillatory optimization is a controllable general principle, changing key modulators of adaptation to oscillatory optimization (repeated cycles) should produce greater sustained specialization. We formalized this as intermittent glucose deficit (IGD), controlled cycles of glucose deficit and recovery during MSCs culture, and asked whether deeper metabolic oscillation in the presence or absence of ATP12A impact sustained metabolic specialization driven by our triad of regulators (**fig. S9E, F**).

To assess this, we used MSCs (expressing the triad of regulators) as control and in combination with low glucose during the days before subculturing (IGD; **fig. S9E, F**). We also used Atp12a-/-MSCs (expressing the triad of regulators) with or without IGD (**fig. S9E**). Notably, IGD in transfected MSCs increased the duration of H3K4me3 marks in mito-HAR genomic hubs, along with mitochondrial activity (20 days, **Fig. 6K, fig. S9F, G**). These effects were further amplified (25 days) in Atp12a-/-MSCs with IGD, which we referred to here as memory-enhanced MSCs (ME-MSCs, **Fig. 6K**). We then evaluated at day 20 long-term molecular adaptations within mito-HAR hubs. This revealed a similar pattern induced by IGD and further enhanced by Atp12-/-, such as elevated H3K4me3, gene expression, and chromatin contacts (**fig. S9H-J**). This was accompanied by substantial functional responses including increased mitochondrial respiration, mitochondrial DNA, and membrane potential (**fig. S10A-D**).

To confirm that pH variations via lysosome function mediate epigenetic memory via MLL5, we repeated the IGD protocol with specific perturbations during the deficit phases of each cycle (**fig. S9E**). This revealed that sustained adaptations were blunted by pH correction, vATPase and Mll4 perturbation (**fig. S9H-J, fig. S10A-D**), indicating that pH fluctuations via lysosome activity and MLL4 writing are required for epigenetic memory and specialization of function. Similar to prior observations, these cells remained in a stem-like preadipocyte state (on day 20, **fig. S10E**). In addition, we tested transcriptional memory reactivation with beta-3-agonists at day 30, which showed enhanced phenotypic recall with persistent adaptations (**fig. S10F**). Critically, these cells, when differentiated into adipocytes, transferred their enhanced mitochondrial function to adipocyte progeny (**fig. S10G-I**). This confirmed that ME-MSCs gained persistent metabolic enhancements while preserving developmental plasticity (**fig. S10G-I**).

##### 4.1.7. Multiscale mechanistic model

Taken together, these results show a more complete mechanistic model at multiple scales (**Fig. 6L, fig. S10J**). Briefly, nutrient oscillation leads to lysosomal pump activation and pH fluctuations (amplified by deleting a molecular break; the ATP12A). These amplified pH fluctuations regulate PGC1A droplet gelation properties, enabling an optimal window for MLL4 recruitment and H3K4me3 mark deposition, which ultimately enhance transcriptional memory. The oscillatory optimization then reinforces every state for progressive and specialization via recursive usage of the same biophysical principle; asymmetric fluctuations enable systems to escape local minima and converge towards optimal structural configurations (**Fig. 6L**). Importantly, we consider pH fluctuations as asymmetric drives given their kinetic differences between the stress-induced and quick acidification phase and the slower ATP12A-modulated variations. This temporal discrepancy is key and, unlike noise, it creates a ratchet effect on specialization readouts, preventing the reversal to its prior state during relaxation phases.

The parallel to Intcyt is more complete now. Just as ML with repeated input cycles produces residual accumulation guiding structural specialization, biological training with repeated metabolic cycles generates epigenetic accumulation that guides transcriptional specialization. Both systems optimize through oscillation, with dissipation (residuals or pH fluctuations) coupled to organizational gains (**Fig. 6D and 6L**). Our experiments proved that in cellular systems during favorable phases, transcriptional compartments assemble, recruit regulators, and install memory marks. During unfavorable phases, the system changes state without erasing progress. However, throughout repeated cycles the system converges toward a specialized state with each oscillation providing a retained increase in specialization, which accumulates into stable phenotype despite modest cost. This further suggests that the cross-domain convergence of asymmetric fluctuations and dissipation serves as an analogous common constraint. Before examining this, we test whether these mechanisms operate in living organisms where cells experience systemic signals, then evaluate cross-domain consistency checks (Section 5).

#### 4.2. Modulating specialization in SCAT-MSCs impacts organismal homeostasis

Our ex vivo work indicated that ATP12A is a modulator of pH fluctuations and specialization. Despite this, key questions remained, for example, whether the role of ATP12A is conserved in vivo, and whether interventions enhancing MSC specialization improve organismal metabolic health in disease models. We tested this via genetic validation, intermittent fasting interventions, and adoptive ME-MSC transfer.

##### 4.2.1. ATP12A expression is dysregulated in disease models

We first evaluated Atp12a expression across metabolic tissues and disease states, revealing that it was present in several organs, including small intestine, kidney, lung, liver, and white adipose tissue (**fig. S11A**). This also showed that Atp12a is preferentially upregulated in SCAT after a high-fat diet (HFD during 8 weeks) and aging (18 months; **fig. S11A**). Atp12a expression was further increased after fasting in SCAT from HFD and aged mice (**fig. S11B**). In the same models, we sorted cell populations from SCAT, which revealed a similar pattern of expression, exclusively in MSCs (**fig. S11C**). These results suggest that maladaptive elevation of ATP12A expression in disease states could limit required specialization responses.

##### 4.2.2. Genetic validation in acute fasting

To test whether ATP12A modulates MSC specialization in vivo and fasting responses, we used whole-body Atp12a knockout mice. In addition, we used a previously described model (Cre-mediated recombination driven by the Prx1 promoter (*124*)) to generate mice with deletion of Pgc1a and Esrra in adipose mesenchymal progenitors (**fig. S11D**). Compared to control fasted mice, SCAT-MSCs from Atp12a-/-mice showed enhanced mito-HARs hubs chromatin contacts after 24h of fasting (**Fig. 7A**). This response was absent in AT-MSCs-Pgc1a-/- and AT-MSCs-Esrra-/-mice, as well as in HFD mice (8 weeks), and aged mice (18 months; **Fig. 7A**), supporting the requirement of regulators and the inhibited response in disease states. In the same models, we evaluated transcriptional compartment activity and mitochondrial function, which revealed that these were only amplified in Atp12a-/-mice after fasting (**Fig. 7B, C**), while being absent in the other mice models (**Fig. 7B, C, fig. S11E, F**). This physiological response and absence in models with specific deletion support our ex vivo observations on key transcriptional compartments regulating mitochondrial specialization.

**Fig. 7.**
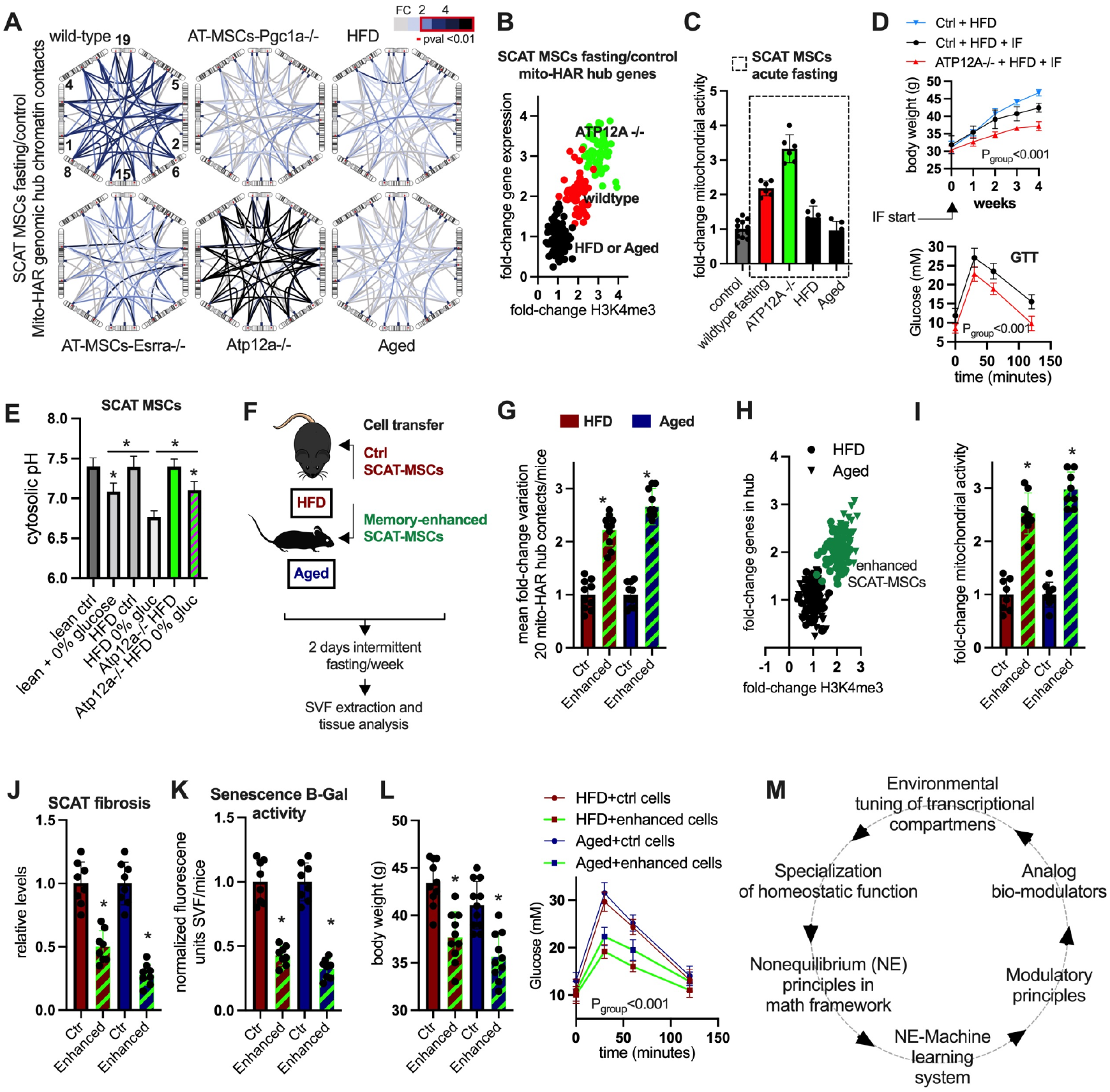
Nonequilibrium tuning enhances adoptive cell transfer and organismal homeostasis. **(A)** *in situ* 3C chromosome conformation assays for chromatin contacts between mitochondrial-HAR hubs in sorted SCAT-MSCs from lean, HFD (8 weeks) and aged (18 months) wildtype mice, Atp12a-/-mutant mice, along with AT-MSCs perturbations such as Pgc1a-/- and Esrra-/-mutant mice after fasting (24h). Contacts are represented as fold-change (FC) over control groups (not fasted littermates). **(B)** Fold-to-fold change plot of mitochondrial-HAR hub gene expression and H3K4me3 levels in regulated genes after fasting in sorted cells from subcutaneous adipose tissue as in **A** (FC >20 genes per group). **(C)** Mitochondrial activity (OCR) in sorted cells from mice as in **B. (D)** Upper panel, body weight evolution in wildtype and Atp12a-/-mutant mice fed a HFD for 12 weeks, data shown for the final 4 weeks during which the mice were treated with 2-day bouts of intermittent fasting (IF) every week. Lower panel, glucose tolerance test at week 4 after IF treatment. **(E)** Cytosolic pH response to glucose deprivation challenge (12h) in sorted and plated SCAT MSCs from lean, HFD wildtype and Atp12a-/-mutant mice. **(F)** Nonequilibrium (NE) tuning strategy for evaluation of adoptive cell transfer to SCAT of HFD and aged wildtype mice. Memory-enhanced (ME) SCAT-MSCs are Atp12a-/-, overexpressed Pgc1a, Esrra and Mll4, and underwent intermittent glucose shortage (12h) previous each cell culture passage (collected on day 6). Once transferred to SCAT, host mice received 2 days intermittent fasting per week. **(G)** Mean FC > 20 contacts from targeted in situ chromosome conformation assays between mitochondrial-HAR hubs in SCAT stromal-vascular-fraction from HFD and aged wildtype mice receiving control or ME-SCAT-MSCs as described in **F. (H)** Fold-to-fold change plot of mitochondrial-HAR hub gene expression and H3K4me3 in promoter genes as in **G. (I)** Mitochondrial activity (OCR) from SVF-cells from the same conditions as in **G. (J)** SCAT fibrosis, hydroxyproline relative levels, **(K)** senescence B-Gal activity in the same conditions. **(L)** The left panel shows body weight. The right panel shows glucose tolerance test in the same conditions as in **G. M**, Summary illustration. For all panels, animal experiments were done with n=5-8 mice per group, both female and male mice were included. Cell experiments were done with 3 independent replicates. Data show mean values and SEM. Unpaired two-tailed student’s t-test for comparing two groups, and ANOVA fisher’s least significant difference (LSD) for multiple groups comparisons. Two-way ANOVA was used to estimate significance between groups constrained by time-based measurements, and Tukey test for multiple comparisons. * indicate p-value <0.05.

Extracted MSCs from HFD, aged, and (Pgc1a and Esrra) KO mice showed an impaired response to a glucose deprivation challenge (*ex vivo*) with excess cytosolic acidity and reduced lysosome activity (**fig. S11G, H**). In contrast, MSCs from Atp12-/-mice showed an enhanced response to the same challenge, even withstanding deleterious effects induced by potassium excess (**fig. S11G, H**). These results show that fasting adaptations in SCAT-MSCs are impaired in diseased models and in models with deletion of regulators of mitochondrial function. It also shows that Atp12a perturbation *in vivo* enhances transcriptional adaptations, pH balance, and mitochondrial function in MSCs after fasting.

##### 4.2.3. Chronic intermittent fasting interventions

Our ex vivo approach showed that repeated metabolic cycles (IGD) progressively extend epigenetic memory in MSCs. We hypothesized that in vivo intermittent fasting (IF; periodic restriction) would similarly create cycles in MSCs, reinforcing their specialization. Moreover, if ATP12A limits fluctuation amplitude per cycle, combining IF with ATP12A deletion should produce synergistic benefits.

To evaluate this, we used a 2 week IF approach in lean control and Atp12-/-models (2 non-consecutive days fasting per week, **fig. S12A**). IF in control mice increased mito-HAR hub chromatin contacts, gene expression, H3K4me3 marks, and mitochondrial activity in SCAT-MSCs, which were all substantially more pronounced in Atp12-/-mice (**fig. S12B-F**). Compared to IF mice, extracted MSCs from Atp12-/-mice + IF showed better lysosome activity and better cytosolic pH balance after glucose deficiency challenge (**fig. S12G, H**). Physiologically, we observed that, after the intervention, body weight gain was modestly reduced in IF mice, but much more Atp12-/-mice + IF (**fig. S12I**). In parallel to our in vitro experiments, IF potentiates molecular adaptations and specialization in MSCs, which are enhanced by Atp12 deletion.

We next tested whether these interventions work in metabolic disease models. We subjected both control and Atp12a-/-mice to 8-week HFD followed by the same IF protocol (**fig. S12J**). Interestingly, Atp12a-/-mice on a HFD had similar body weight to control HFD mice before the IF protocol (**Fig. 7D**). However, compared to control HFD mice with IF, IF in Atp12a-/-HFD mice reduced body weight gain, improved glucose tolerance test (GTT, **Fig. 7D**), and increased mitochondrial function in the stromal vascular fraction (SVF) of SCAT (**fig. S12K**). An *ex vivo* nutrient deprivation challenge showed that, compared to cells from lean mice, SCAT-MSCs from HFD mice are unable to maintain cytosolic pH (**Fig. 7E**). This effect was not observed in cells from Atp12a-/-mice in HFD (**Fig. 7E**).

These results supported that genetic deletion of the ATP12A maladaptive break allows IF to efficiently enhance molecular and metabolic adaptations associated with mitochondrial function specialization and overall health parameters. Given that pharmacological cell-specific targeting remains challenging, we sought to implement an alternative strategy, whether our ex vivo approach enhances therapeutic outcomes from engineered cells.

##### 4.2.4. Enhancing adoptive therapeutic cell transfer

To evaluate ectopic interventions with engineered MSCs, we used a single SCAT adoptive cell transfer of control and ME-MSCs to HFD and aged recipient mice (**Fig. 7F**). Post adoptive transfer, the recipient mice were subjected to two weeks IF (two days of IF per week), to enhance metabolic cycling and reinforce ME-MSC function in vivo. At the end of this protocol, extracted SVF from recipient SCAT was analyzed, revealing molecular adaptations induced by ME-MSCs. Compared to control-MSC recipients, ME-MSCs recipients showed enhanced mito-HAR hubs chromatin contacts, elevated gene expression, and increased H3K4me3 levels (**Fig. 7G-I, fig. S13A, B**). Mitochondrial function and lysosome activity in the SVF of these models were similarly enhanced (**Fig. 7G-I, fig. S13A, B**). This indicates that the SVF from ME-MSCs recipient acquired molecular adaptations similar to the adopted cells (mitochondrial function gain) despite being in diseased environments.

Systemic effects were substantial as well. ME-MSCs recipients showed reduced adipose tissue expression of the Tnfa inflammatory marker (**fig. S13C**), reduced fibrosis, and reduced senescence levels (**Fig. 7J, K**). Metabolic parameters improved proportionally. Compared to control recipients, ME-MSCs recipients lost more body weight and improved their glucose tolerance test (GTT) significantly (**Fig. 7L**). Mechanistically, we found that ME-MSCs transfer activated thermogenic gene programs in recipient adipocytes (**fig. S13D, E**). Similar thermogenic gene programs were observed in mice with deletion of Atp12a and treated with IF in both lean and obese phenotypes (**fig. S13F, G**). This suggests that ME-MSCs transfer promotes the phenotypic programming of adipocytes towards thermogenic oxidative metabolism, so long associated with improved whole body metabolism and overall health.

Taken together, our in vivo validation experiments across genetic, nutritional, and cell therapy paradigms indicate that the principles identified through our integrated biological and computational framework translate to physiological contexts and hold therapeutic promise. The remaining question is whether our observations, where organization gains correlate with dissipation proxies in both Intcyt and MSCs, exhibit cross-domain thermodynamic consistency. This requires systematic comparison of our interventions.

### 5. Lower-bound diagnostic proxies on thermodynamic-consistency across domains

We examined specialization mechanisms in two dynamic systems. A biological (cellular) system that specializes by tuning transcriptional compartments dynamics through drives such cytosolic pH fluctuations, and a machine system, designed around corresponding operational principles, which learns through adaptive compartmentalization while tracking residual accumulation. Despite substantial mechanistic differences, we observed qualitative similar patterns; asymmetric fluctuations are associated with specialization readouts while fluctuation amplitude increases during optimization windows.

This convergence is consistent with the possibility that both contextual changes across systems are shaped by related nonequilibrium constraints. In particular, if specialization is dependent on recurrent increases in local organization (subsystems with lower organization entropy), then generalized second-law reasoning suggests that local gains are accompanied by dissipation elsewhere (higher entropy (*86, 87*)). We therefore asked a simple but directional question, whenever our systems become more organized, are our dissipation proxies increasing more? If our framework captures shared rules of adaptive specialization, both systems (machine and cellular) should show ‘sign-level’ agreement across perturbations, where operational accounting of proxies could be read off the data. We tested this by defining proxies for each domain and examining sign-level consistency.

#### 5.1. Dissipation proxies

A challenge in comparing thermodynamic properties across domains is to identify proxies for dissipation accounting in each system while remaining consistent across substrates. Our goal here was to document relative changes and evaluate sign-level consistency across perturbations (not absolute entropy; the mapping is proxy-based and not a claim of mechanistic identity).

##### 5.1.1. Built-in proxy for in silico adaptive learning

In Intcyt, information is discarded when the system performs irreversible operations (e.g., zeroing cytosolic content into a residual ledger). We treat the resulting residuals as a lower-bound proxy for information erasure (Landauer events). Two practical features characterize this: (i) the bound proxy is intrinsic to the algorithm, as every structural update that cannot be reversed leaves an auditable trace in the form of residuals; (ii) we also report this trace in LND units (see sections 2.4.4-2.4.5), allowing conservative comparisons (see Supp. Methods M1-7 and **supp. method Figure. 1**). For example, during self-organizing learning, all three variables in Intcyt increase together, residual accumulation, fluctuation amplitude, and specialization readouts (**Fig. 6A-D**), consistent with the notion that stronger organization is accompanied by greater irreversible-updates in the model.

##### 5.1.2. A cellular proxy for adaptive specialization

For living cells, direct measuring of entropy is more challenging as tracking every chemical event is nearly impossible. Instead, we used machine learning predictions and physiological observations to identify, test, and modulate a dissipative proxy during specialization. We use cytosolic pH fluctuation amplitude as an empirically tractable proxy for cellular energy-dependent activity based on several reasons: (i) energetic basis: pH fluctuations require active work by proton pumps. Conditions that amplified them are associated with higher metabolic rate and energy expenditure; (ii) dissipation-dependent responses: modulating pH fluctuations unidirectionally affects specialization readouts and phenotype robustness.

While amplitude of fluctuations is an indirect proxy, it responds predictably to several perturbations. For our purposes, probing sign-level consistency across perturbation offers a tractable parameter. For example, low glucose increases pH fluctuation amplitude (**Fig. 6F–G**); vATPase/TMEM175 perturbation diminishes it, and ATP12A perturbation amplifies it (**Fig. 6H**). Across all of these manipulations, the dissipation proxy co-varies with specialization readouts—mito-HAR hub gene and epigenetic activity and OCR (**Fig. 6I**), PGC1A-droplet FRAP recovery (**Fig. 6J**), and the duration of specialization (**Fig. 6K**). In sum, manipulations that open the dissipation window improve organization readouts and perturbations that reduce it impair specialization readouts.

#### 5.2. Proxy-based sign-level closure

To test directional thermodynamic consistency, we compiled perturbations across both systems and assessed whether changes in the dissipation proxy matched changes in specialization readouts (**Table 1**). This showed that in every condition tested across perturbation in both substrates, the direction of change in organization/specialization features matches the direction of change in the dissipation proxy. No condition shows organizational gains with reduced dissipation or specialization disparities. This sign-level agreement can be viewed as being directionally consistent with second-law expectations under proxy-based accounting, where local organization requires dissipation proxies, such that our framework is (i) a learning system that exposes its irreversibilities as a quantitative trace, and (ii) where the cell system behaves similarly, with the behavior being experimentally controlled via proton-pump activity.

#### 5.3. Fluctuation-driven efficiency: diagnostic on rectifying oscillations

Beyond sign-level closure, it is important to describe the main features of how a system is driven. A remaining question to establish is how dissipation (energetic drive) does work on organization, which opens two possibilities: (i) tonic forcing, in which continuous energy input pushes the system toward more organized states; (ii) oscillatory rectification (see section 3.7.3.), where periodic fluctuations open brief windows in which barriers to irreversible steps are lowered, such that the system accumulates organizational gains and specialization readouts.

Across domains our data supports that asymmetric fluctuations can act as a low-grade control drive that the system rectifies into organization. Rectification here refers to converting bidirectional fluctuations into net unidirectional progress in organization-related metrics. For example, in cells, externally paced oscillations are gated by proton-pump networks and converted into useful work including condensate remodeling, epigenetic marking, and persistent phenotypes. During these windows, ATP-dependent processes (H3K4me3 deposition, looping, etc) occur with reduced effective barriers, while the marks are able to persist well beyond the window and the gains are retained without continued input.

Computationally, windows of asymmetric updates play the analogous role. For instance, fission/fusion can convert these windows into compartment structures that better match the data manifold. Minimization of *U*^*2*^ then selects and keeps representative structures while discarding redundant configurations and tracking associated irreversible operations. Once this process is stabilized, the architectures that were selected persists across cycles unless the input distribution shifts, which computationally mirrors epigenetic mark retention in cells.

A reason oscillations may be more efficient than tonic forcing is that concentrating organizational work into short favorable intervals reduces total energy expenditure by: (i) barrier lowering, irreversible steps are carried out when activation barriers are low; (ii) passive retention of cyclic gains, either in readouts or structure, without the need for continuous drive; (iii) temporal gating of key steps when conditions are productive, avoiding suboptimal phases. A consequence of productive drives and phases is more organization per unit dissipation than sustained forcing of the same amplitude drive (*10, 113, 114*).

We summarize this with a relative drive-to-organization efficiency (DOE) diagnostic metric, DOE = Δ Organization/LND, where Δ Organization is a relative change in a metric associated with specialization (compartment consolidation; H3K4me3/OCR/memory readouts) and Δ LND is the bits-equivalent dissipation lower bound proxy. This parameter is treated here as a relative metric for substrate directional comparisons and not as an absolute thermodynamic efficiency. Conditions that amplify fluctuations (SOL; low glucose; ATP12A modulation) tend to exhibit higher DOE (more organization per unit dissipation) than conditions with suppressed fluctuations (steady state fluctuation in intcyt, vATPase/TMEM175 perturbations in cells; see **Table 2**, Supp. Methods M5 and **Supp. method Fig. 1E**).

**Table 2.** Relative drive-to-organization efficiency: All entries are fold-changes relative to baseline (cells: 5.5 mM glucose; Intcyt: early-learning phase). ΔOrganization (relative) represents specialization gains: for cells, fold-change in H3K4me3 at mito-HAR promoters (directional agreement with OCR); for Intcyt, organization proxy such as residual ledger and based on compartment dynamics (amplitude × operations; U^2^-linked consolidation) between early baseline learning and self-organizing learning (SOL) phases. LND (relative) is a Landauer-normalized dissipation proxy (lower-bound; no absolute energy) that is further normalized to each baseline: for cells, it is proportional to cytosolic-pH fluctuation amplitude over the window of analysis; for Intcyt, from the residual ledger increased by irreversible updates. DOE (relative) quantifies organization per unit dissipation relative to baseline, where values > 1 indicate higher efficiency. Ratio directions represent the stated baselines. Estimates are detailed in Supp. Methods (M).

| System | Phase/Condition | $\Delta \text{Organization}$ | LND | DOE |
| --- | --- | --- | --- | --- |
| Cells | Control (glucose) | 1x | 1x | 1x |
| Cells | Low glucose (LG) | 2.5x | 2.1x | 1.19x |
| Cells | vATPase-RNAi (LG) | 0.5x | 0.6x | 0.83x |
| Cells | ATP12A-RNAi (LG) | 4x | 2.4x | 1.67x |
| Intcyt | Early (baseline) | 1x | 1x | 1x |
| Intcyt | Learning | 9.5x | 5.1x | 1.86x |

#### 5.4. Implications of diagnostic consistency

This cross-domain agreement has several implications: (i) it suggests that information-theoretic and nonequilibrium perspectives may be applied to cellular biology, as an informed guide and a qualitative predictive tool; (ii) it reveals that the amplitude of cytosolic pH fluctuation may serve as a tractable proxy influencing specialization, enabling real-time monitoring, similar to residuals in our computational model monitoring structural adaptation. Additionally, features in asymmetric fluctuations can be used as a metric of organizational gains and specialization outcomes; (iii) it supports the notion that the dissipation accounting in our computational framework is not arbitrary as its proxy behavior aligns well with experimental perturbations across domains. More broadly, these design rules indicate that increasing key dissipation proxies (ATP12A modulation or residual policies) may open windows for optimal specialization readouts while suppressing dissipation closes them. It also shows that optimal approaches to design rules should aim for the creation, timing, and alignment of fluctuation windows to overlap the active recording of irreversible steps into persistent specialization when the systems are most responsive. This rectification rule may scale from cellular organization to adaptive architectures and suggests tunable handles for both cell programming and thermodynamically informed ML architectures.

## Discussion

Our combined biological, mathematical, and computational results support a shared qualitative picture: structural plasticity in nonequilibrium systems is not arbitrary, but rather influenced by controllable fluctuations that can be rectified into durable organization and specialization. We show this picture in two different substrates, cellular and adaptive ML systems, which behave following the same organizational logic.

Biologically, we characterize multimodal transcriptional information related to specific cell functions and establish a principled approach for identifying rules linking molecular adaptations to functional specialization. Our framework defines key operational rules, such as [1] nested compartments, [2] dynamic operations, [3] interaction with external drives or inputs, and a [4] nonequilibrium optimization based on how oscillation of metabolic states influence cell specialization via fine-tuning of transcriptional properties and epigenetic memory (**Fig. 7M** and **Fig. 8A**).

**Fig. 8.**
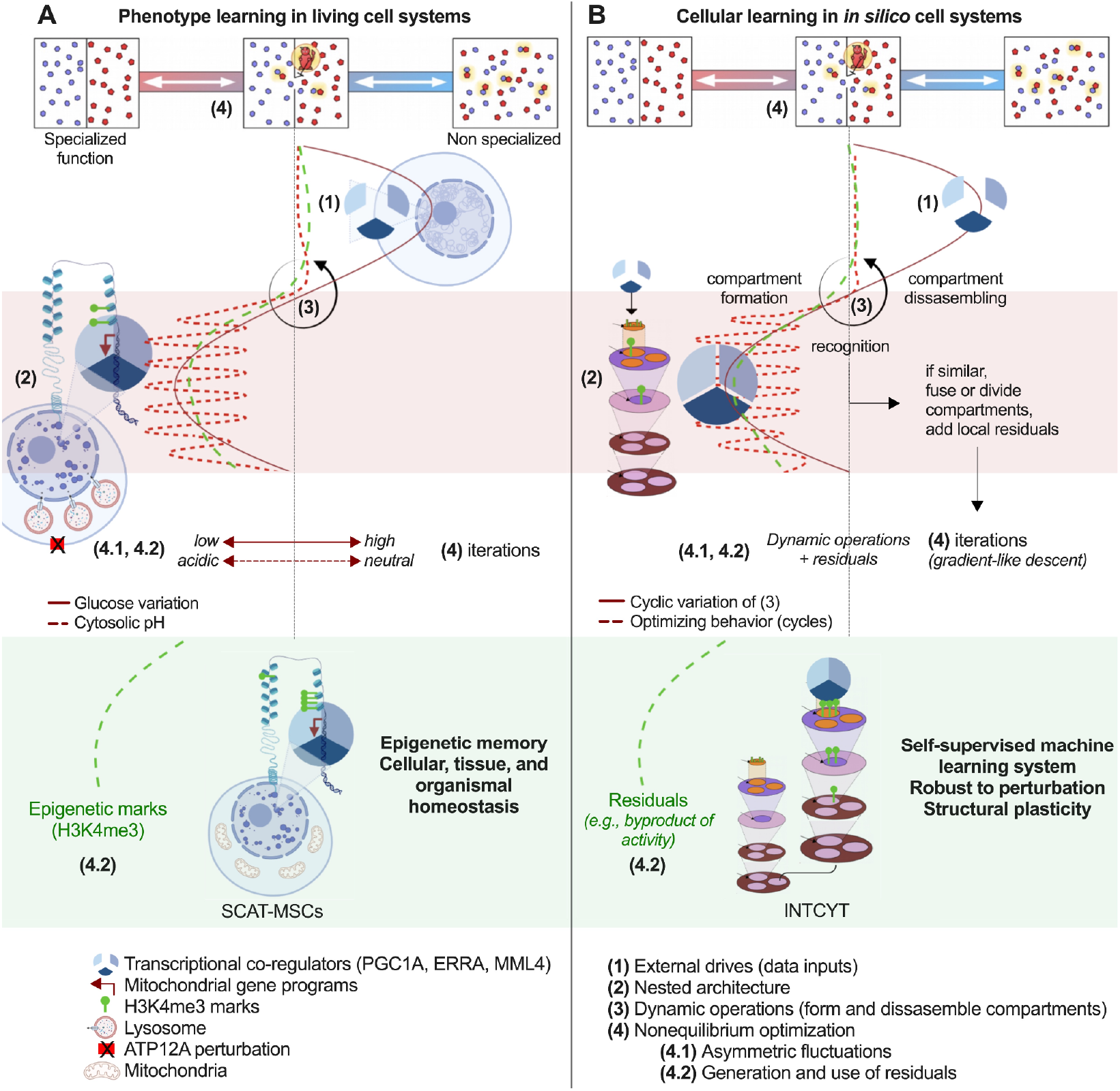
Nonequilibrium tuning enhances functional specialization in cellular and machine learning systems. Summary illustration displaying a multi-domain approach to investigate adaptive specialization in nonequilibrium systems. **(A)** In the left panel, a description of specialization in cell systems. The lower left panel shows labeled components. **(B)** In the right panel, specialization in Intcyt. The lower right panel shows the principles derived from the biological system and modeled by our computational framework in machine learning paradigms.

Theoretically, we investigate these operational rules by developing a mathematical framework using a tree-like multiscale structure and a nonequilibrium optimization approach. We use operadic models–a powerful language from algebraic topology and category theory–as they effectively express composition, factorization, multiscale representation, and adaptability, which all are valuable for studying adaptive systems that need consistency across scales (*103–105*). We develop an optimization principle inspired by Maxwell’s demon paradox as it integrates nonequilibrium concepts. In this system, oscillations between low-entropy and high-entropy configurations (forming and disassembling compartments) are guided by actions, allowing the identification of ideal configurations for a given dynamic system. This mathematical framework is inherently constructive, which provides a foundation for structural learning.

Computationally, Intcyt implements these adaptive compartmentalization principles as a self-organizing ML system. To form and disassemble compartments, it uses factorization and composition while iteratively evaluating input similarities (external drives), which, once recognized, generates new compartments (**Fig. 8B**). Structural transitions are coupled with a gradient-like descent optimization, allowing the system to progressively evaluate optimal configurations for given inputs. Computing operations generate residual byproducts (dissipation proxy ledger) when recognizing similar inputs and forming new compartments, which continuously guide structural specialization.

Our framework shows that asymmetric fluctuations drive specialization through thermodynamically informed, energy-efficient adaptation. Three key advances: (i) mechanistic identification of molecular effectors whose manipulation tunes specialization and memory of function, (ii) formalization of their logic via operadic algebra and a nonequilibrium optimization, providing a self-organizing ML system that recapitulates key biological dynamics, and (iii) thermodynamic sign-level consistency in our interventions (**Table 1**), with oscillatory organization achieving higher efficiency than steady-state conditions. This shows that the efficiency improves under fluctuation amplification in both cells and computation (**Table 2**), indicating more organization per unit dissipation.

### Framework for adaptive specialization via rectified fluctuations

This work connects and operationalizes phenomena from cellular specialization and adaptive computation under a candidate organizing principle: adaptive systems may optimize through controlled oscillation between organized and exploratory states, where amplitude and asymmetry of fluctuations can influence the extent of specialization. Instead of relying on continuous forcing, they exploit tunable asymmetric drives. Across modulations, amplifying fluctuations is linked to specialization, while suppressing them is associated with reduced organization.

In our tested conditions, oscillatory organization can be more efficient than continuous forcing. An interpretation is that it may concentrate organizational work into optimal windows (intermediate droplet viscosity, high amplitude of compartmentalization) where barriers are low, leading to retained informational gains between cycles. This, in turn, avoids wasteful updates when conditions are not optimal across both domains, supporting the view that oscillatory mechanisms are predominant in biological systems, where evolutionary optimization has discovered that temporal structure in energy expenditure enables cheaper organization than constant expenditure.

### Relationship to existing frameworks

These findings extend to several frameworks while providing mechanistic insights. For example, the Waddington epigenetic landscape describes differentiation as balls rolling into valleys (*125*). We provide instantiation where valleys represent transcriptional compartments with low U^2^ and where hills can represent energy barriers requiring pH fluctuation-driven crossings. Oscillations act as repeated metabolic cycles accumulating persistent epigenetic memory marks, and we show that the landscape evolves through use, with each cycle deepening valley optimization.

Another framework is metabolic control of cell fate (*126, 127*), which emphasizes that metabolic switches drive lineage decisions. We show key mechanistic insights that establish asymmetric oscillations between states as optimal for specialization of function, compared to static continuous states. Static and prolonged deficit produces limited specialization with dampened fluctuations, while intermittent deficit amplifies fluctuations and produces robust long-lasting memory of functional states.

Phase separation models (*14–16*) have revealed that condensates concentrate molecular machinery to perform specific cellular functions. We show that material properties, such as intermediate viscosity at optimal pH, optimize droplet function. Too-liquid condensates fail to retain MLL4 while too-solid condensates arrest dynamics and function. FRAP studies identified an optimal dynamic zone for epigenetic interaction and enzymatic activity, linking condensate material properties to energetic state.

The IB principle states that useful representations compress irrelevant information while preserving predictive information (*128, 129*). Here we provide an accounting perspective on IB-style trade-offs within learning; improvements in predictive information are paralleled by erasure-like operations needed for predictive updates. Minimizing U^2^ acts as a task-fit proxy for predictive structural information, where compartment actions match inputs, while residual accumulation is associated with discarded non-predictive state. This reframes bottleneck behavior from a purely abstract optimization to a process with auditable update costs. Even though we do not claim formal equivalence to the IB optimum, our optimization approximates some features that emerge from constraints not being imposed externally. In addition, both stochastic thermodynamics (*57, 58, 86, 87*) and Maxwell demon frameworks (*85, 106*) offer formal models of the costs of information processing but there have been few applications to adaptive systems. We implement experimentally lower-bounds for sign-level closure across interventions, where we show that specialization is coupled to key dissipation proxies in both cellular and computational adaptation.

## Implications

### Biomedical engineering

Cytosolic pH fluctuations during low-energy states regulate transcriptional plasticity, epigenetic memory, and phenotype specialization, and these can be tuned by molecular modulators (ATP12A pump). We show that modulators of drives enhance the response to adoptive cell transfer in murine disease models. Thus, ATP12A is a tractable target for multiple modalities including modulation for stem cell expansion, temporal systemic dosing paired with intermittent fasting, and memory retention of cell function applicable to CAR T cell therapies or other adoptive transfer interventions.

Our framework suggests that metabolic diseases reflect not only substrate imbalance but also dysregulation of key molecular oscillatory dynamics that enable efficient adaptation. Investigating and restoring oscillatory capacity might be as important as correcting metabolic substrates. Future work could include other operational rules in additional cell types, which could assess the impact of environmental conditions, genetic mutations, and epigenetic modifications on phenotype diversity. Understanding additional principles will provide ways to design, modulate, and identify programmable targets for therapeutic approaches.

### Artificial intelligence

Our framework provides foundational steps toward paradigms of adaptive learning such as thermodynamically grounded and biophysics-inspired AI (an adaptive framework where structure self-organizes via fission/fusion under an explicit budget constraints (LND), optimizing a Maxwell-inspired objective (U^2^) and delivering continual learning and interpretability with improved DOE). Current AI architectures treat learning as a cost-free optimization, which leads to inefficiencies and structural rigidity. We show that by embedding learning with adaptive physical constraints in a multiscale nested framework (that keeps consistency across scales), can lead to the emergence of computational properties more naturally, even in low budgeting computations.

Our algorithm shows how thermodynamic accounting and structural operations provide computationally desirable properties: (i) self-organization under constraint without hyperparameter search, where architecture adapts automatically via fission/fusion guided by U^2^ minimization under residual accounting; (ii) continual learning via selective retention rather than computational overwrite, where new inputs create new compartments naturally mitigating catastrophic forgetting; (iii) energy-aware training with oscillatory schedules that raise DOE, suggests a route to higher efficiency in hybrid neurosymbolic systems or in analog neuromorphic hardware; (iv) interpretability by construction: hierarchical tree structure provides inherent interpretability, each compartment represents learned features, U^2^ contributions explain why these exist and why they change.

Future work should aim at integrating Intcyt with deep learning architectures, which could help in addressing some of the problems observed during backpropagation. This integration could also optimize computational resources for hybrid systems, and establish the foundation for adaptive architectures guided by molecular principles. For example, deep learning can handle pattern recognition while operadic structure provides adaptive higher-level and continuous structural learning integration. This also couples well with hardware-algorithm co-design in neuromorphic architectures where the material analog could implement the same logic physically, allowing more efficient computation and energy-aware systems.

### Asymmetric oscillations as a general organizing principle

Our observation that oscillatory driving can outperform steady forcing in DOE (**Table 2**) offers one possible thermodynamic motivation for the ubiquity of biological rhythms. Instead of maintaining organized states continuously (expensive), systems may exploit temporal drives during windows where downstream processes are primed, accumulating persistent changes that do not require continuous energy to maintain. For example, circadian, cell-cycle, developmental programs, and neural oscillations, may function not only as clocks but as rectifiers, timing costly, irreversible steps to maximize organization per unit dissipation. This aligns well with temporal interventions (e.g., time-restricted feeding, cyclic stimulation, interval training) and with time-scale matching, showing that they outperform tonic regimens when the goal is persistent adaptation. This suggests a general design rule for adaptive control in living and artificial systems.

If oscillatory organization can approach favorable efficiency regimes in certain contexts, evolution may in some cases converge toward rhythmic strategies across scales. This was observed in our analysis and modulation of nested compartments enriched in evolutionary accelerating DNA regions (HARs), linked to highly conserved function across metazoa such as mitochondrial activity. The oscillatory tunability per species would allow traits to be conserved across species yet exhibit species-specific adaptation. Thus, this structural adaptive organization provides evolutionary flexibility within physical constraints. This design rule may go beyond biology with impactful routes for computational integration with current AI. Oscillatory forcing, which has already been observed across thermodynamic systems such as stochastic resonance and brownian motors (*115, 130*), shows that breaking symmetry to extract work from undirected drives is an efficient process. It would be interesting to see this integration where adaptive ML systems exploit similar physics while providing quantitative relationships for fine-tuning.

### Limitations and considerations

There are limitations that are worth consideration: (i) Proxy measurements: We infer dissipation based on measure of pH amplitude fluctuations and computational residual increments as proxies. These proxies directionally couple to specialization readouts and respond to perturbations accordingly, which would suggest causal association, this still requires absolute quantification with calorimetry. Our LND and DOE calculations (Table 2) are thus conservative lower bounds, consistent with physical constraints. The connection to the information bottleneck is intuitive and conceptual and a formal equivalence will be the grounds for future work. These theoretical developments are ongoing work and our current formulation provides heuristic bounds supported by data but does not claim mathematical completeness. (ii) Mechanism: We find a molecular link between pH and droplet viscosity but this requires further investigation. Potential options include pH-dependent IDR charge residues mutations (in line with previous work (*89*)), protonation-dependent conformations, or pH-sensitive crosslinkers. In addition, data showing that ΔIDR2 mutants lose pH-associated recruitment of MLL4 supports intrinsic IDR properties for specific interactions, but reconstitution experiments are still needed. (iii) Scope: In this work, we decided to focus on MSCs and mitochondrial specialization for specific reasons, but whether these principles extend to other cell types (neurons, immune cells) and programs (synaptic plasticity, immune memory) requires further evaluation. In addition, the identified operational features (nested compartments, dynamic assembly, oscillatory optimization) appear generalizable because oscillatory rhythms are widespread in biology, but analogs need to be found in each context. (iv) Computation: We show proof-of-concept with a strong integration of formalisms, but this has not been scaled to modern benchmarks, which would need hybrid architectures for practical implementations. Scaling to current benchmarks might also need the development of further learning policies and how these are implemented together with deep learning. One approach might be to start with adaptive learning of learned features from neural network architectures. (v) Theory: Rigorous formulation still needs to be performed, for instance formal proof that operadic optimization satisfies thermodynamic uncertainty relations and explicit mapping from residuals to entropy production costs in kT units. These theoretical concepts need to be further developed and tested as our current formulation provides heuristic bounds supported by data and does not claim mathematical completeness. (vi) Alternative interpretations: We discuss in the supplementary text alternative explanations and empirical evidence against them (Supplementary text (S1): section S6.5; see also section S6.3)

## Conclusion

We show that specialization in adaptive systems, both cellular and computational, is driven by asymmetric fluctuations through energy-efficient mechanisms. In cells, nutrient variation induces pH fluctuations tuning transcriptional compartments and epigenetic memory, modulable by ATP12A, intermittent fasting, and cell engineering. Formalization of principles via operadic optimization shows that compartment fluctuations coupled to a processing ledger drive adaptive architecture evolution. Across domains, organization gains correlate with dissipation proxies and achieve higher efficiency than steady-state alternatives.

Specialization is neither idiosyncratic to biology nor arbitrary in computation but instead it reflects fundamental constraints of information processing. Two key elements are (i) oscillatory rectification, which times irreversible steps to windows of lowered barriers, and (ii) an explicit cost ledger that makes efficiency optimizable. This framework provides practical and conceptual advances, including (1) mechanistic targets for metabolic disease, (2) design fundamentals toward thermodynamically-informed and energy-aware adaptive AI, and (3) empirical observations that biological and computational information processing share tunable constraints (see appendix **Table 3**).

The cross-domain sign-level consistency suggests that we have captured operational principles beyond implementation details in each system. This opens exploratory directions such as engineering synthetic fluctuation-guided systems, testing whether immune memory, neural plasticity, or specialization in ecosystems use similar rectifiers; and tightening bounds on learning efficiency with organization-rate limits and dissipation budgets. Specialization thus emerges not from stochasticity, but from asymmetric fluctuations rectified into persistent architectures by systems far from equilibrium, a principle with a shared language with which we can further explore adaptation across living, computational, and other physical systems.

## Acknowledgements

We thank all the members from the various research labs and collaborators who have contributed to this project. We thank Amy Grayson and coauthors for the help reviewing and editing the manuscript. **Funding:** LZA is supported by Daice Labs inc. with previous funding from Novo Nordisk and MIT CSAIL. MK is supported by MIT with previous funding from Novo Nordisk. TF is supported by Ramon y Cajal grants by the Spanish state research agency, and the “Severo Ochoa” programme for Centres of Excellence in R&D (SEV-2017-0723).

## Author contributions

Project design & Conceptualization: LZA. Cross-domain framework design and methodology: LZA. Mathematical methodology: RT with feedback from LZA, MK. Software Intcyt methodology: RT, LZA with feedback from MK. Experimental design and validation: LZA, AS, TF, AM, CL. Supervision of the work: TF, MK, LZA. Funding acquisition: TF, MK, LZA. Writing original draft: LZA. Writing & editing: TF, RT, MK, LZA. Authors reviewed the manuscript.

## Supplementary Materials

Figs. S1-S13

Tables S1-S2

Biological and statistical materials and methods.

Supplementary text (S1): Mathematical and computational framework

Theoretical, Mathematical, and Computational Framework

Appendix: Mathematical and computational framework.

## Data and materials

Data is available in the supplementary materials. The code and the documentation for Intcyt is hosted at the following github address: https://github.com/daicelabs/intcyt-v2

---

**A. Appendix Table 3.** Summary table showing advances across domains.

| Domain | Key Advances | Core evidence | Where |
| --- | --- | --- | --- |
| Biology | **Oscillatory drives** | Time-resolved pH peaks and bidirectional modulation of specialization outcomes | Fig. 6F–J; §4.1–4.2 |
|  | **Optimal window (viscosity)** | FRAP inverted-U; MLL4 recruitment and mark-writing | Fig. 6J; §1.4.2 |
|  | **ATP12A as tunable brake** | Modulation of ATP12A influences functional specialization | Fig. 6H–I; §4.2–4.4 |
|  | **Compartment scaffolds** | Spatial templates coordinating stress-responsive programs | §1.3.1–1.3.2 |
| Mathematics | **Operadic formalism** | Constructive, multi-scale composition/factorization | §2–2.4 |
|  | **Maxwell-inspired objective** | Nonequilibrium loss compares organized vs mixed states | §2.4 |
|  | **Information processing** | Residuals audit irreversible steps as a dissipation proxy | §2.4.4; §3.5 |
| Computation | **Self-organizing ML** | Fission/fusion yields adaptive architecture; residuals ↔ consolidation | §3.1–3.2; §3.5 |
|  | **Continual learning** | New inputs trigger new compartments rather than overwriting old ones | §3.2 |
| Cross-Domain | **Sign-level closure** | organization gains ↔ dissipation proxies | Table 1 |
|  | **Oscillatory efficiency** | DOE improves under fluctuation amplification | Table 2 |
|  | **Rectification principle** | Fluctuations into durable organization via temporal gating | §3.7.3; §5.3 |

