## Supplementary materials for "Rectifying Stress into Structural Memory Drives Specialization in Cells and Adaptive Computational Systems"

### Supplementary Figures

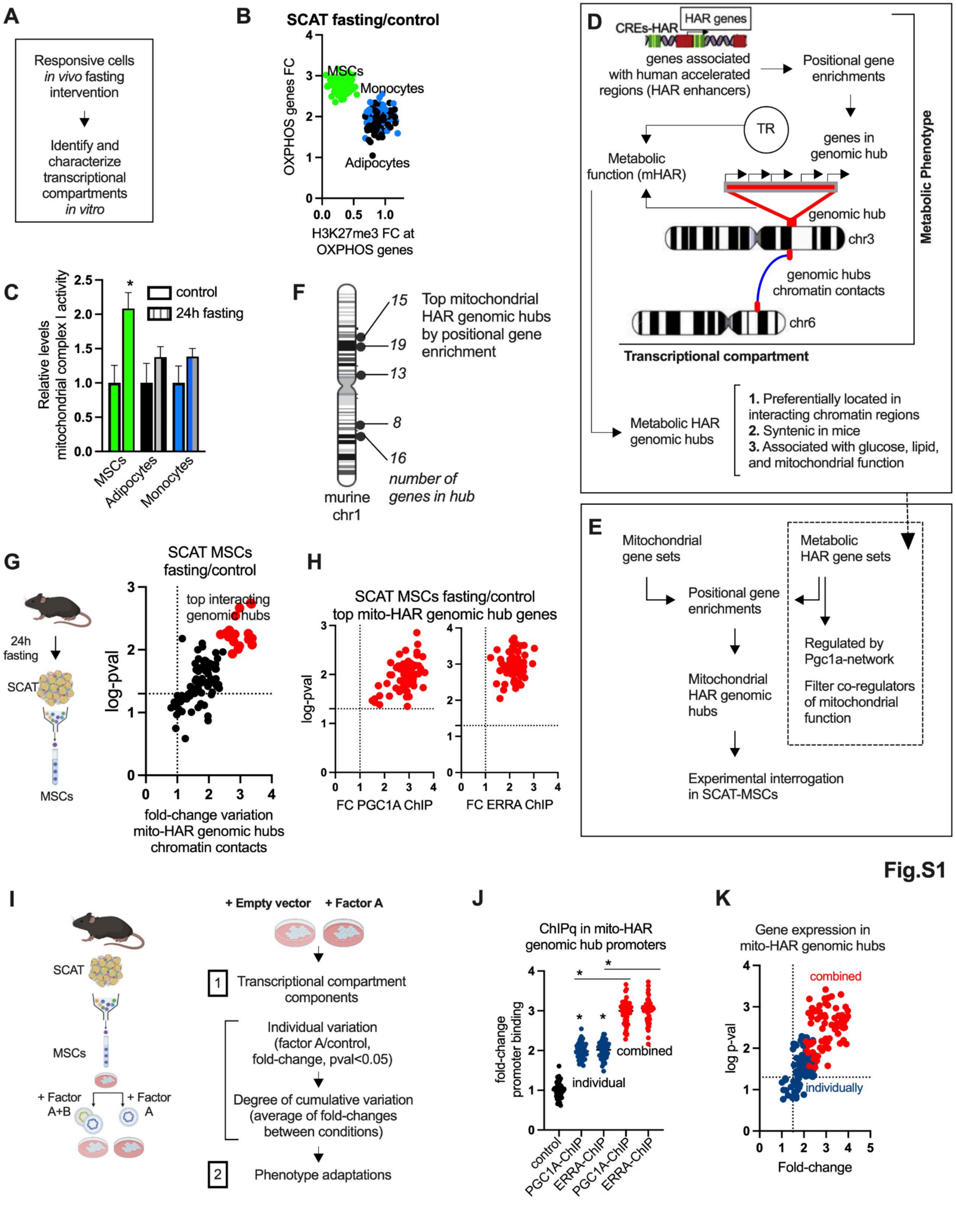

**Fig. S1. Transcriptional compartments associated with mitochondrial function.** *(related to Fig. 1).*
**(A)** Schematic illustration for identifying and experimentally validating transcriptional compartment adaptations in responsive cells.
**(B)** Left panel illustrates fasting intervention and cell sorting. Right panel, fold-to-fold change plot of OXPHOS gene expression and H3K27me3 levels after fasting (24h) in sorted cells from subcutaneous adipose tissue wildtype lean mice. Data showing mean fold-change variation of H3K27me levels and gene expression in at least 30 mitochondrial function genes per group. In MSCs all data points have p-value<0.05 over control.
**(C)** In sorted cells as in B, functional assay for mitochondrial complex I activity.
**(D)** Schematic illustration for functional genomic analysis of human accelerated regions. HAR-associated genes are located in specific genomic regions (metabolic HAR hubs) associated with metabolic function and controlled by transcriptional regulators (TR) such as PGC1A and PGC1A-associated transcription factors. These genomic hubs are located in interacting chromatin regions, are syntenic in mice, and are related to mitochondrial function.
**(E)** Schematic illustration for the identification of genomic hubs. Mitochondrial gene sets and metabolic HAR genes used to identify shared genomic hubs by positional gene enrichments (named mitochondrial HAR genomic hubs or mito-HAR hubs). Metabolic HAR genes are regulated by the PGC1A, and interacting metabolic transcription factors (e.g., PPARA, PPARG, ERRA). Regulators of mitochondrial genes such as ERRA and PGC1A were selected for experimental validation.
**(F)** Representative illustration showing top mitochondrial HAR genomic hubs in murine chromosome 1.
**(G)** Left panel illustrates fasting intervention and sorting of MSCs. Right panel, targeted *in situ* 3C chromosome conformation assays showing fold-change variation of chromatin contacts between top mito-HAR genomic hubs in SCAT-MSCs after fasting (24h) in wildtype lean mice as in **B**. Data shows mean fold change variation of contacts between genomic hubs (fasting/control, at least 4 contacts per genomic hub were assessed, and were used to calculate the mean fold change variation). In red, top interacting genomic hubs.
**(H)** PGC1A and ERRA ChIP-qPCR in promoters from mitochondrial-HAR genomic hubs from murine SCAT-MSCs (fasting over control).
**(I)** Isolation of SCAT-MSCs, plating, and engineering with specific transcriptional regulators. Cells were transfected with expression vectors using the nucleofector instrument and protocols for primary cells.
Right panel shows strategy for SCAT-MSCs engineering with specific transcriptional regulators, evaluation of transcriptional compartment components (gene expression, ChIP-qPCR, ChIP-loop, histone modifications), and phenotypic adaptations.
**(J)** PGC1A and ERRA ChIP-qPCR in genes from mitochondrial-HAR genomic hubs from murine SCAT-MSCs as in **I**. Data showing mean fold-change variation of binding in at least 20 promoter genes from mito-HAR genomic hubs.
**(K)** Gene expression for mitochondrial-HAR genomic hub genes in the same conditions as in **I**.
For all panels, animal experiments were done with n=5-8 mice per group, both female and male mice were included. Cell experiments were done with 3 independent replicates. Data show mean values and SEM. Unpaired, two-tailed student's t-test was used when two groups were compared, and ANOVA followed by Fisher's least significant difference (LSD) test for *post hoc* comparisons for multiple groups. * indicates p-value <0.05.

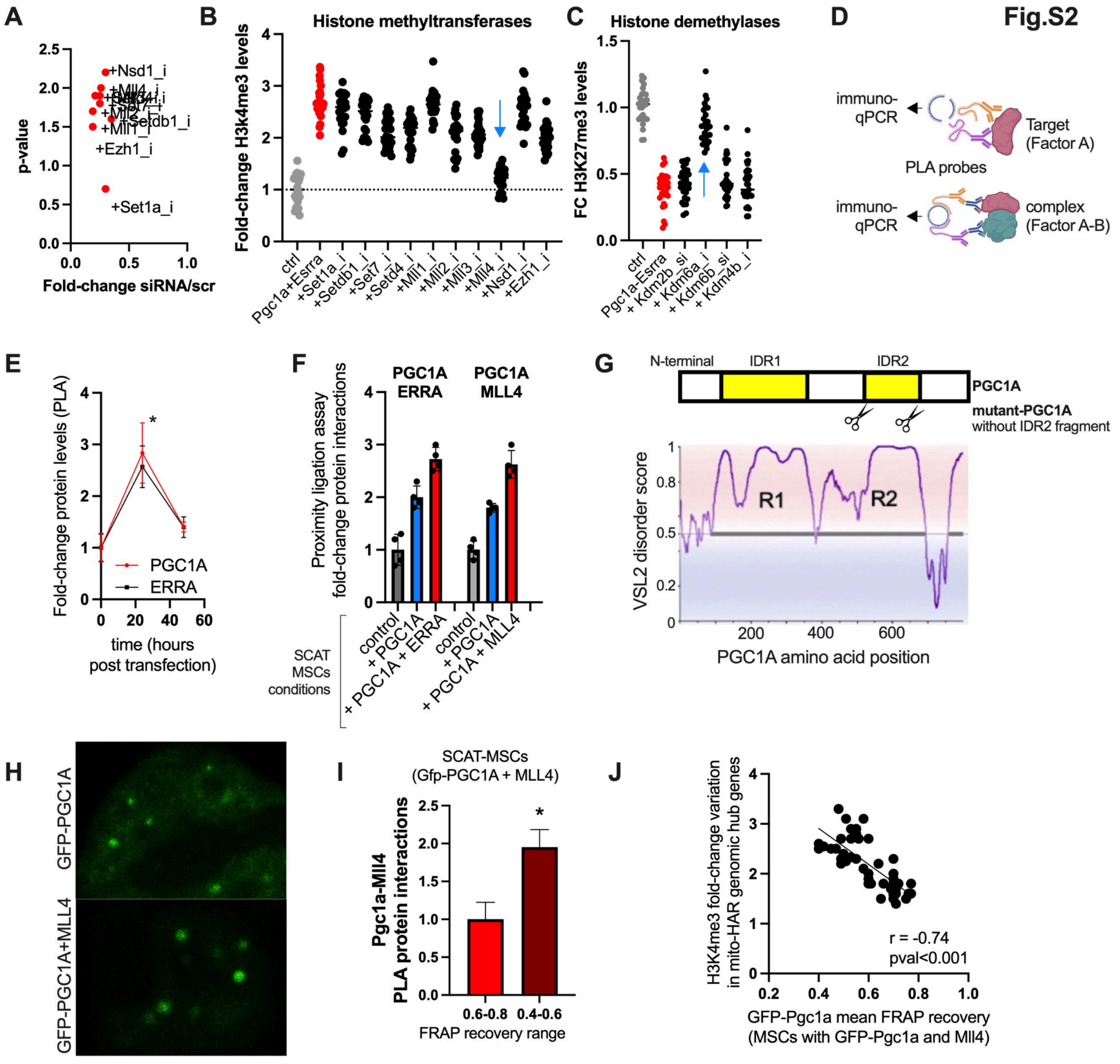

**Fig. S2. Transcriptional compartment associated with SCAT-MSCs mitochondrial function.** *(related to Fig. 2).*

**(A)** Gene expression knockdown efficiency for RNAi perturbations in transfected SCAT-MSCs.

**(B)** RNAi screen to identify active histone methyltransferases and **(C)** histone demethylases in SCAT-MSCs after 24 h of combined transfection with PGC1A and ERRA (siRNA/scramble). Data shows mean fold change variation of H3K4me3 and H3K27me levels in promoters of at least 20 regulated mito-HAR genomic hub genes per group.

**(D)** Illustration showing proximity ligation assay strategies for detection of individual (top panel) and in complex (lower panel) proteins.

**(E)** Proximity ligation assays (PLA) for PGC1A and ERRA individual proteins in transfected SCAT-MSCs. Probes conjugated with antibodies recognizing (GFP or His) and protein epitopes.

**(F)** Proximity ligation assays (PLA) for PGC1A protein interactions in SCAT-MSCs transfected with PGC1A (no GFP-tag), ERRA, and MLL4 individually or combined.
**(G)** Lower panel shows VSL2 disorder score for the transcriptional coactivator PGC1A amino acid sequence. Upper panel shows wildtype PGC1A protein illustration with IDR regions. The IDR2 region is flanked with scissors representing the deletion location of IDR2 for the generation of mutant PGC1A construct.
**(H)** Confocal microscopy images for GFP-PGC1A droplets in living SCAT-MSCs after 24 h of transfection with GFP-PGC1A and MLL4 individually or combined.
**(I)** Proximity ligation assays (PLA) for PGC1A-MLL4 protein interactions in SCAT-MSCs as in **C**. Relative levels of interactions were partitioned by FRAP (fluorescence recovery and photobleaching assays) recovery of GFP-PGC1A droplets in SCAT-MSCs.
**(J)** Fold-change H3K4me3 levels in mito-HAR genomic hubs promoters (Y-axis) and mean fluorescence recovery after photobleaching (X-axis) of GFP-PGC1A droplets in SCAT-MSCs treated as in **C** (r and p-value from pearson's correlation).
For all panels, cell experiments were done with 3 independent replicates. Data show mean values and SEM. Unpaired, two-tailed student's t-test was used when two groups were compared, and ANOVA followed by fisher's least significant difference (LSD) test for post hoc comparisons for multiple groups. Nonlinear regression (Lorentzian-Cauchy model) and extra sum-of-squares F test to compare models between conditions. * indicates p-value $<0.05$.

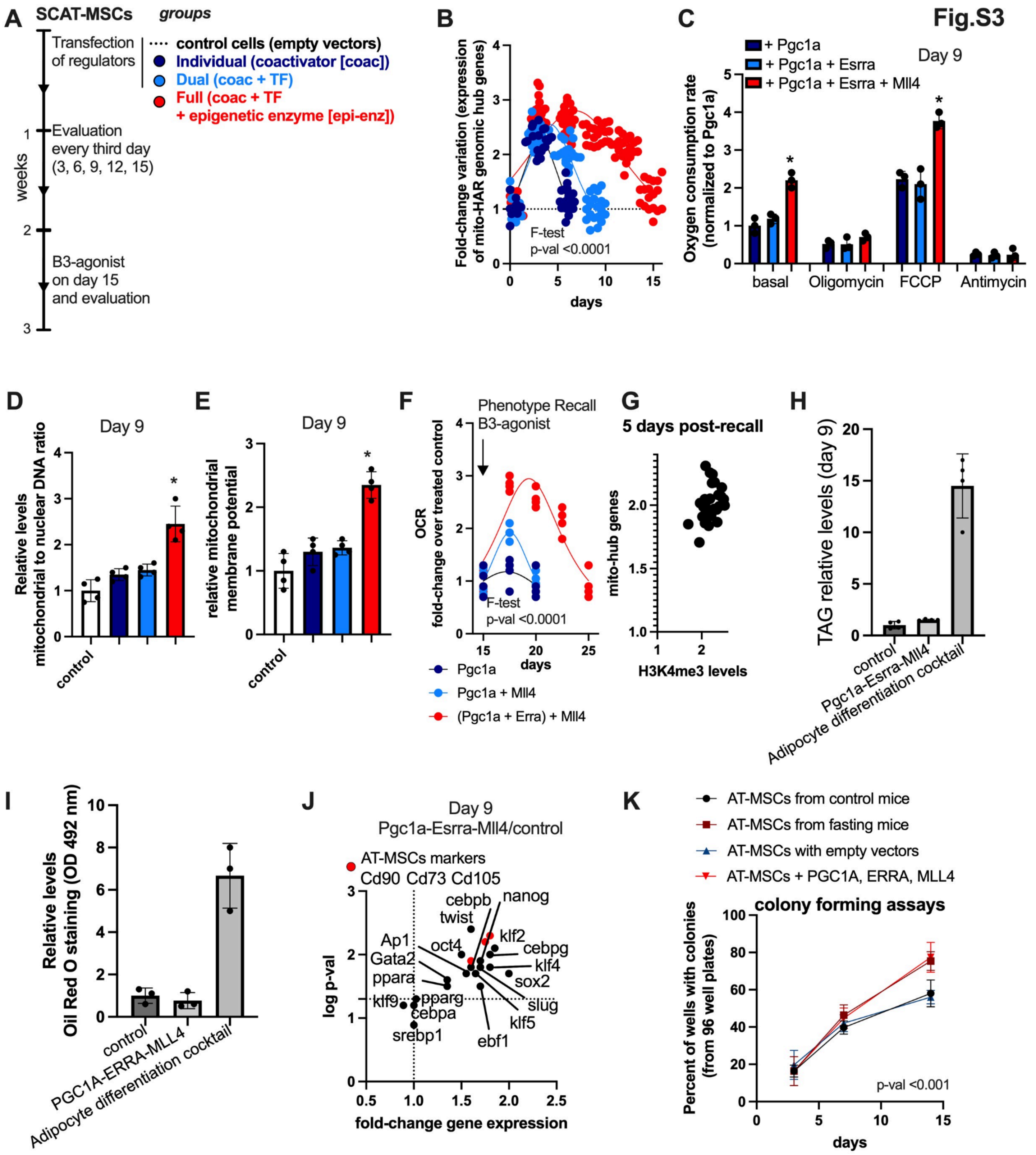
Fig.S3
A
SCAT-MSCs
groups
Transfection of regulators
control cells (empty vectors)
Individual (coactivator [coac])
Dual (coac + TF)
Full (coac + TF + epigenetic enzyme [epi-enz])
Evaluation every third day (3, 6, 9, 12, 15)
weeks
B3-agonist on day 15 and evaluation
B
Fold-change variation (expression of mito-HAR genomic hub genes)
F-test p-val <0.0001
days
C
+ Pgc1a
+ Pgc1a + Esrra
+ Pgc1a + Esrra + Mll4
Day 9
Oxygen consumption rate (normalized to Pgc1a)
basal
Oligomycin
FCCP
Antimycin
D
Day 9
Relative levels mitochondrial to nuclear DNA ratio
control
E
Day 9
relative mitochondrial membrane potential
control
F
Phenotype Recall B3-agonist
OCR fold-change over treated control
F-test p-val <0.0001
days
Pgc1a
Pgc1a + Mll4
(Pgc1a + Erra) + Mll4
G
5 days post-recall
mito-hub genes
H3K4me3 levels
H
TAG relative levels (day 9)
control
Pgc1a-Esrra-Mll4
Adipocyte differentiation cocktail
I
Relative levels Oil Red O staining (OD 492 nm)
control
PGC1A-ERRA-MLL4
Adipocyte differentiation cocktail
J
Day 9
Pgc1a-Esrra-Mll4/control
AT-MSCs markers Cd90 Cd73 Cd105
log p-val
fold-change gene expression
cebpb
nanog
twist
klf2
Ap1
oct4
cebpg
Gata2
klf4
ppara
sox2
klf9
pparg
slug
cebpa
klf5
srebp1
ebf1
K
AT-MSCs from control mice
AT-MSCs from fasting mice
AT-MSCs with empty vectors
AT-MSCs + PGC1A, ERRA, MLL4
colony forming assays
Percent of wells with colonies (from 96 well plates)
p-val <0.001
days

**Fig. S3. Mitochondrial specialization in SCAT-MSCs through heterotypic regulators.** *(related to Fig. 2).*

**(A)** Schematic illustration for experiments assessing the amplitude and duration of molecular and phenotypic adaptations in engineered SCAT-MSCs. Cells transiently overexpressed PGC1A (individual), PGC1A+ERRA (dual), PGC1A+ERRA+MLL4 combined (full), and control cells with empty vectors. Evaluation was performed every third day. **(B)** Time-frame fold-change variation in expression of mitochondrial-HAR genomic hub genes in SCAT-MSCs as described in **A**.

**(C)** At day 9 of cell culture of SCAT-MSCs described in **A**, seahorse measurement of oxygen consumption rate (OCR) at basal and in the presence of 1μM oligomycin (uncoupler), 1μM FCCP (maximal respiration inducer), or 1μM antimycin (respiration inhibitor).

**(D)** Relative mitochondrial/nuclear DNA ratio in SCAT-MSCs as in **A**, and compared to control (not transfected with transcriptional regulators).

**(E)** Relative mitochondrial membrane potential in SCAT-MSCs treated as in **A**, and compared to control (not transfected with transcriptional regulators).

**(F)** Time-frame fold-change variation of oxygen consumption rate (OCR) in the same conditions as in **A** and after treatment with beta-3 adrenergic receptor agonists CL-316,243 (10 nM, Tocris) on day 15 of cell culture.

**(G)** Mito-HAR genomic hubs gene expression and H3K4me3 fold change variation after 5 days of beta-3 agonist treatment in cells nucleofected with PGC1A+ERRA+MLL4.

**(H)** Triglycerides relative levels in SCAT-MSCs (control and combined transfection), and compared to SCAT-MSCs treated with adipocyte differentiation cocktail (7 days post-induction).

**(I)** Oil red O staining relative levels in SCAT-MSCs treated as in **H**.

**(J)** Transcription factor expression profile at day 9 in SCAT-MSCs after transfection with transcriptional regulators over control cells.

**(K)** Colony forming assays in SCAT-MSCs from control and fasting (24h) mice, and SCAT-MSCs with expression vectors. Cells were plated in 96 well plates and data is shown as the percent of wells with colonies for time points and conditions.

For all panels, cell experiments were done with 3 independent replicates. Data show mean values and SEM. Unpaired, two-tailed student's t-test was used when two groups were compared, and ANOVA followed by fisher's least significant difference (LSD) test for post hoc comparisons for multiple groups. Two-way ANOVA was used to estimate significance between groups constrained by time-based measurements, and Tukey test for multiple comparisons. Nonlinear regression (Lorentzian-Cauchy model) and extra sum-of-squares F test to compare models between conditions for time-frame experiments. * indicate p-value <0.05.

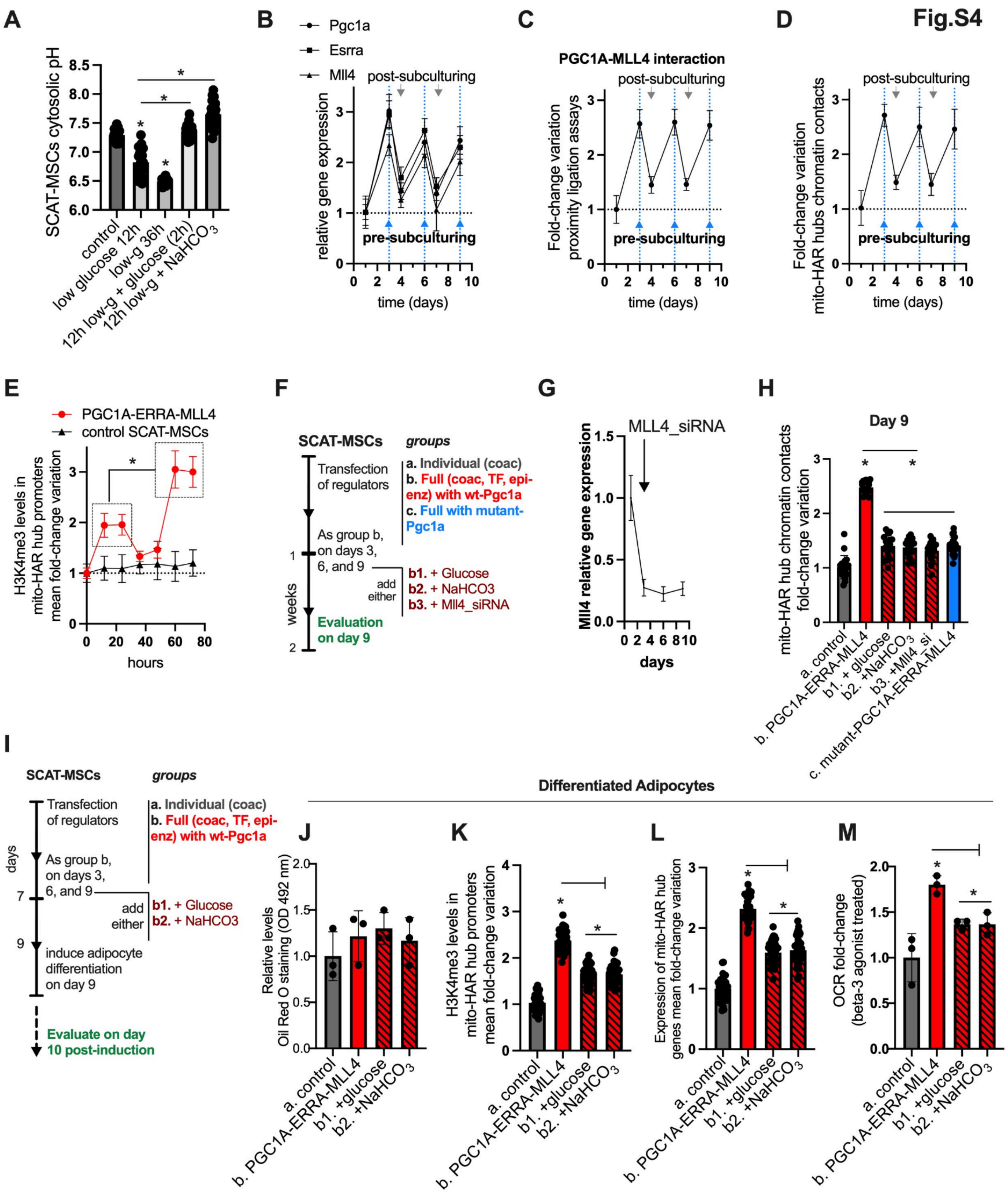
Fig.S4
A
SCAT-MSCs cytosolic pH
control
low glucose 12h
low-g 36h
12h low-g + glucose (2h)
12h low-g + NaHCO3
B
Pgc1a
Esrra
Mll4
post-subculturing
pre-subculturing
relative gene expression
time (days)
C
PGC1A-MLL4 interaction
Fold-change variation proximity ligation assays
D
Fold-change variation mito-HAR hubs chromatin contacts
E
PGC1A-ERRA-MLL4
control SCAT-MSCs
H3K4me3 levels in mito-HAR hub promoters mean fold-change variation
hours
F
SCAT-MSCs
groups
Transfection of regulators
a. Individual (coac)
b. Full (coac, TF, epi-enz) with wt-Pgc1a
c. Full with mutant-Pgc1a
As group b, on days 3, 6, and 9
add either
b1. + Glucose
b2. + NaHCO3
b3. + Mll4_siRNA
weeks
Evaluation on day 9
G
MLL4_siRNA
Mll4 relative gene expression
days
H
Day 9
mito-HAR hub chromatin contacts fold-change variation
a. control
b. PGC1A-ERRA-MLL4
b1. + glucose
b2. +NaHCO3
b3. +Mll4_si
c. mutant-PGC1A-ERRA-MLL4
I
SCAT-MSCs
groups
Transfection of regulators
a. Individual (coac)
b. Full (coac, TF, epi-enz) with wt-Pgc1a
As group b, on days 3, 6, and 9
add either
b1. + Glucose
b2. + NaHCO3
days
induce adipocyte differentiation on day 9
Evaluate on day 10 post-induction
Differentiated Adipocytes
J
Relative levels Oil Red O staining (OD 492 nm)
K
H3K4me3 levels in mito-HAR hub promoters mean fold-change variation
L
Expression of mito-HAR hub genes mean fold-change variation
M
OCR fold-change (beta-3 agonist treated)
a. control
b. PGC1A-ERRA-MLL4
b1. +glucose
b2. +NaHCO3

**Fig. S4. Mitochondrial specialization through environmental state oscillations.** *(related to Fig. 3).*

**(A)** Cytosolic pH variation in SCAT-MSCs in control (glucose 5.5 mM) and low-glucose media (2 mM) for 12, 24, and 36 hours. 12 hours low glucose cells were supplemented with glucose (5.5 mM) or $NaHCO_3$ (1 mM). **(B)** Fold-change variation of expression of transcriptional regulators over time (days) in murine SCAT-MSCs transfected with PGC1A, ERRA and MLL4 combined and control cells (dotted line not transfected cells). Arrows show pre- and post-subculturing day variation.

**(C)** Proximity ligation assays showing fold-change variation of protein interaction between transcriptional regulators over time (days) in murine SCAT-MSCs treated as in **A**.

**(D)** *In situ* 3C chromosome conformation assays showing fold-change variation of chromatin contacts between mito-HAR genomic hubs over time (days) in murine SCAT-MSCs treated as in **A**. Data shows mean fold change variation of at least 10 contacts per group.

**(E)** Fold-change variation of H3K4me3 in mitochondrial-HAR genomic hub promoters (10 promoters evaluated) over time (hours) in SCAT-MSCs control and transfected with regulators (P-value for group comparison <0.01).

**(F)** Schematic illustration for perturbation experiments during cell culture. SCAT-MSCs cells overexpressed PGC1A, individually (here representing control), and combined, or combined with mutant-PGC1A construct. During the days before sub-culturing, media was supplemented with glucose or $NaHCO_3$ (1 mM). In addition, we used Mll4-siRNA in the same days.

**(G)** Gene expression knockdown efficiency for Mll4 RNAi perturbations added during 24h before subculturing days.

(**H**) On day 9, SCAT-MSCs were collected and assayed for chromatin contacts between mito-HAR genomic hubs (mean fold change variation of at least 15 contacts per group)

**(I)** Schematic illustration for perturbation experiments during cell culture followed by adipocyte differentiation. SCAT-MSCs cells overexpressed PGC1A, individually (here representing control), and combined. During the days before sub-culturing, SCAT-MSCs with combined expression were supplemented with glucose or $NaHCO_3$ (1 mM). On day 9, cells were induced to adipocyte differentiation and were evaluated 10 days post-induction.

On day 10, adipocytes from modified SCAT-MSCs, as described in **I**, were collected and assayed for (**J**) oil red o staining levels, (**K**) H3K4me3 (fold change variation in promoters of at least 40 genes per group), (**L**) gene expression (fold change variation of at least 40 genes per group), and (**M**) oxygen consumption rate (OCR) after beta-3 adrenergic receptor agonists CL-316,243 (10 nM, Tocris) treatment (12h).

For all panels, cell experiments were done with 3 independent replicates. Data show mean values and SEM. Unpaired, two-tailed student's t-test was used when two groups were compared, and ANOVA followed by Fisher's least significant difference (LSD) test for post hoc comparisons for multiple groups. Nonlinear regression (Lorentzian-Cauchy model) and extra sum-of-squares F test to compare models between conditions for time-frame experiments. * indicates p-value <0.05.

A

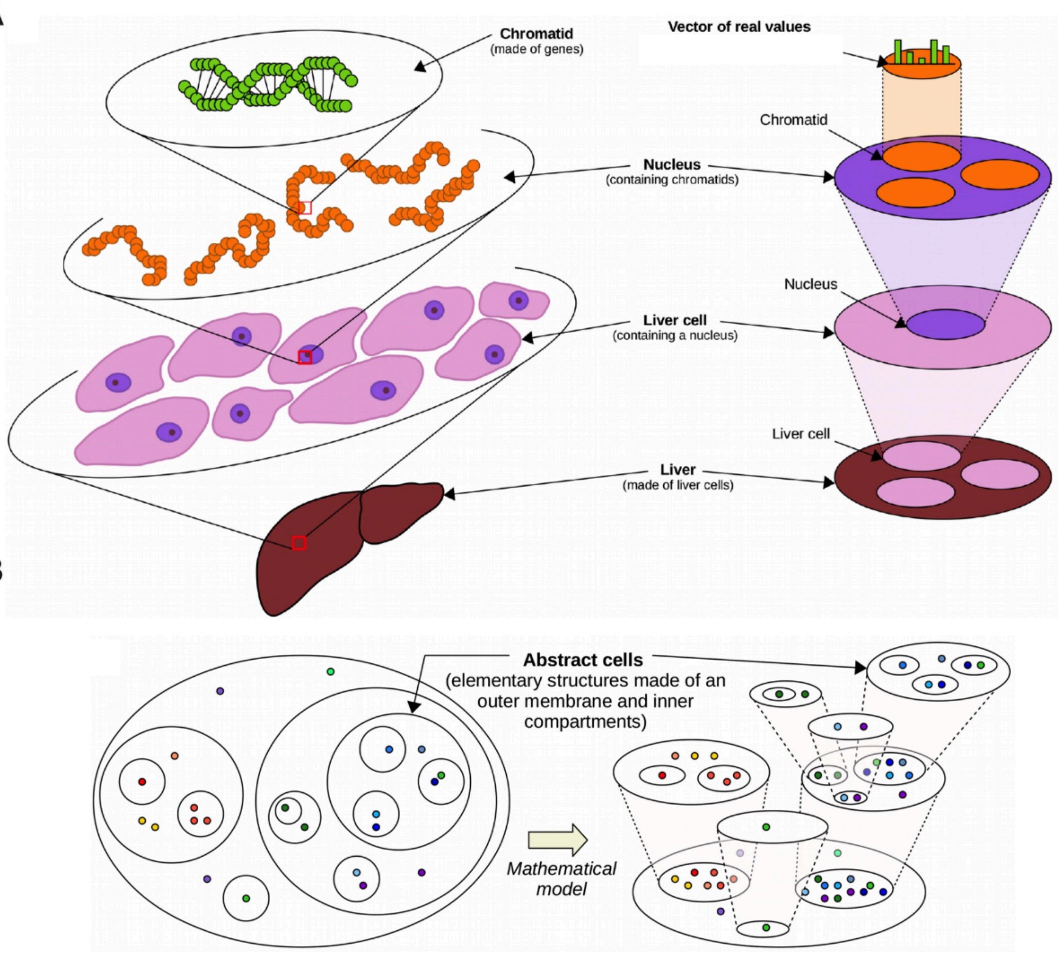

C

| Operadic Language | Dynamic behaviors |
|---|---|
| Composition | Compartment disassembling |
| Factorization ( = reverse composition) | Compartment formation |
| Composition **+** factorization | Exchanges between compartments / homeostasis |
| Inner factorization **+** outer composition | Fusion of compartments |
| Outer factorization **+** inner composition | Fission of compartments |

D E

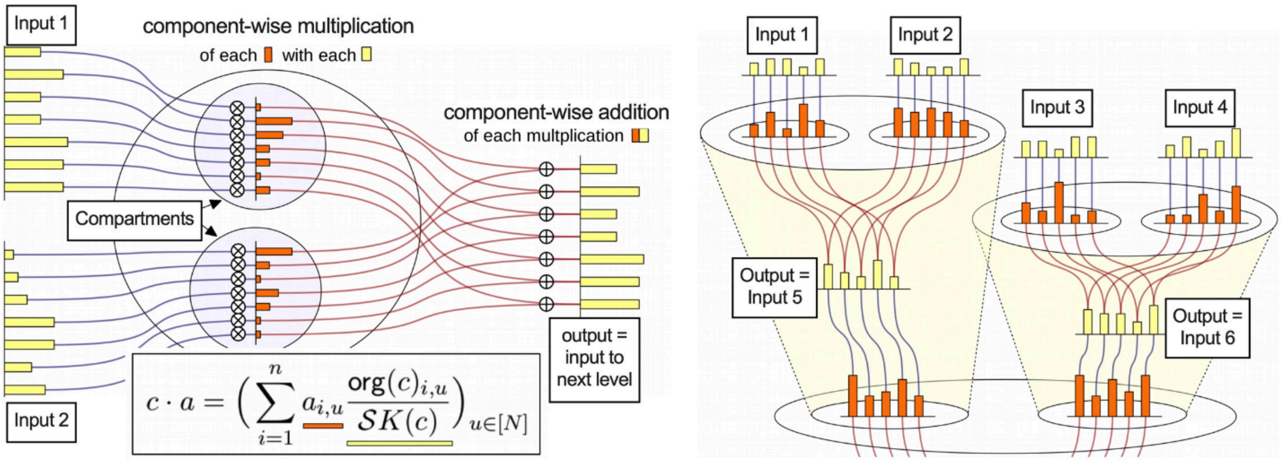

**Fig. S5. Modeling dynamic multi-scale compartmentalized systems** *(related to Fig. 4)***.**

**(A)** Schematic of a multi-scale organization (tissue level representation). On the left, successive scale enlargements zoom in the nested composition of the liver. On the right, these successive close-ups can be represented as a nesting of cell-like structures. These cell-like structures are made of a milieu and a collection of compartments meant to receive further nesting. The milieu and each compartment is equipped with a vector of real values representing the functional specialization within the system.
**(B)** Schematic representation of how we model a multi-scale compartmentalized ecosystem as a tree of (abstract) cells. We can imagine visualizing the multi-scale compartmentalized ecosystem by looking at the tree of cells from above, as suggested by the descending transparent cones. In the tree shown in the picture, each cell compartment describes the content of its hovering child cell.
**(C)** Table summarizing the dynamic behavior modeled by the operadic framework.
**(D)** Schematic showing how an abstract cell of dimension 7, equipped with 2 organelles, processes a 2x7-matrix of real numbers. The output, which is a vector of dimension 7, is the result of a component-wise multiplication of the columns of the matrix (i.e. called “input 1” and “input 2”) with the organelles of corresponding index, followed by a component-wise sum of these multiplication.
**(E)** Schematic illustrating how a tree of cells of dimension 7, equipped with 4 leaf cells, processes a 4x7-matrix by propagating the actions of its cells from the leaves to the root. As can be seen, outputs returned by child cells are turned into inputs for the parent cells.

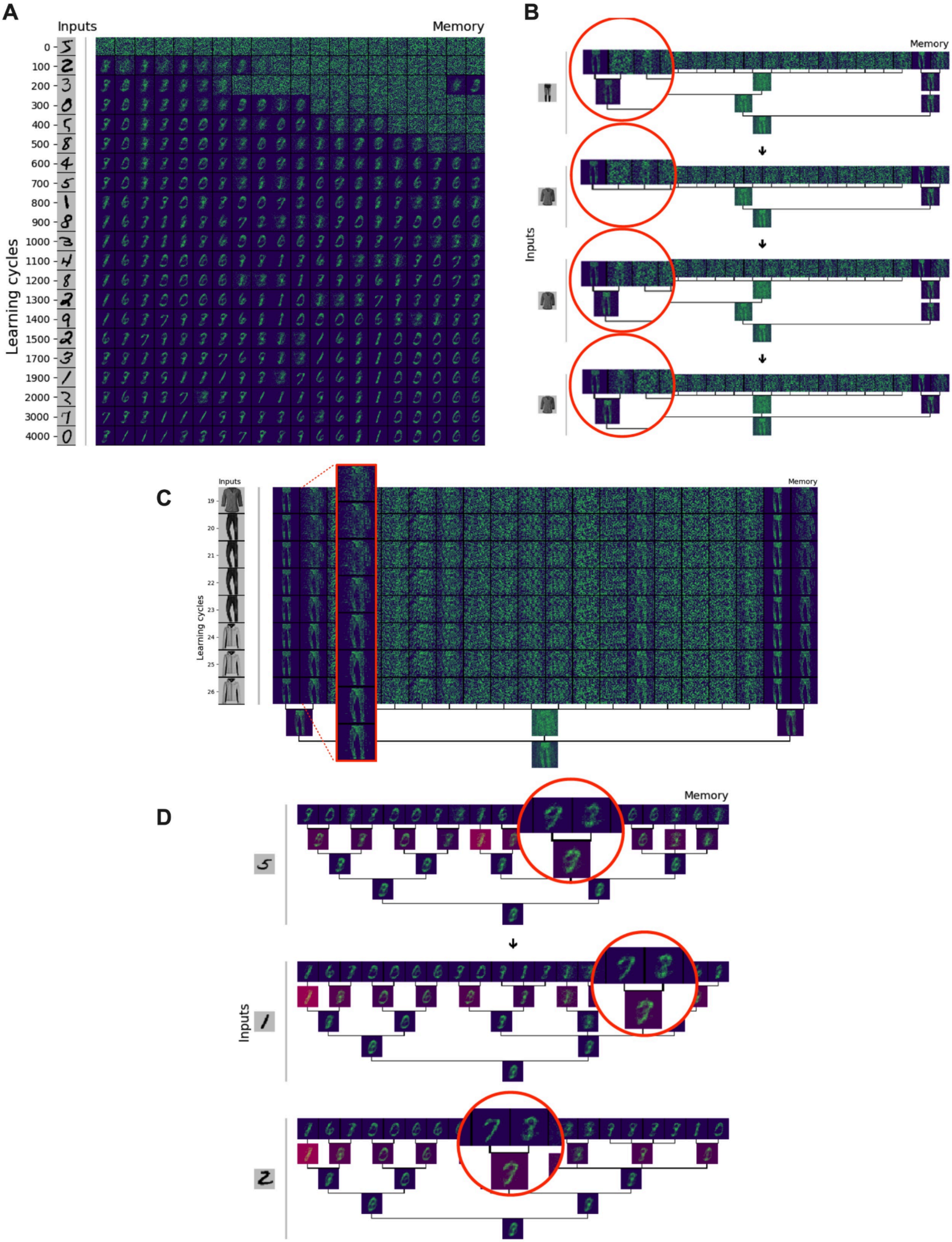
A
Inputs
Memory
Learning cycles
0
100
200
300
400
500
600
700
800
900
1000
1100
1200
1300
1400
1500
1700
1900
2000
3000
4000
B
Memory
Inputs
C
Inputs
Memory
Learning cycles
19
20
21
22
23
24
25
26
D
Memory
Inputs

**Fig. S6. Optimization of dynamic behavior by a nonequilibrium optimization.** *(related to Fig. 5).*

**(A)** Visualization showing the evolution of the organelles of the leaf cells as well as the state of the tree for the last cycle. The inputs given to the tree are taken from the MNIST dataset. As can be seen, Intcyt first separates noise from information, and eventually clusters the different memorized pieces of information into hierarchical compartments.
**(B)** Visualization showing an example of "recruitment" in the case where the inputs originate from the dataset fashion-MNIST. The leftmost organelle integrates a nearby cell and recruits the organelle of that cell with similar information in a new compartment.
**(C)** Visualization showing an example of a communication between cells right after the cycles shown in panel **B**. Specifically, the organelle containing the image influences its neighbor organelle to change its content and specialize towards a similar image.
**(D)** Visualization showing that communication between organelles can lead to smoothening the abstractions made by Intcyt. Specifically, the memorized image of similar inputs. The cycles shown in the figure correspond to the cycles 750, 1150 and 1250 of the simulation shown in panel **A**. By using such communication mechanisms, Intcyt avoids overfitting its inputs.

**A**

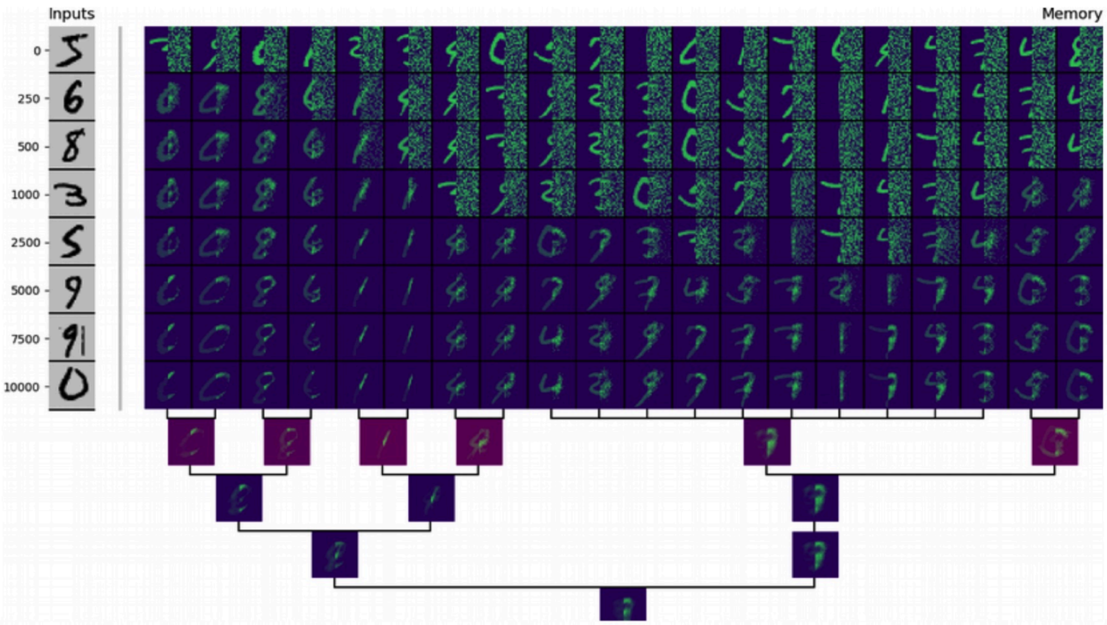

**B**

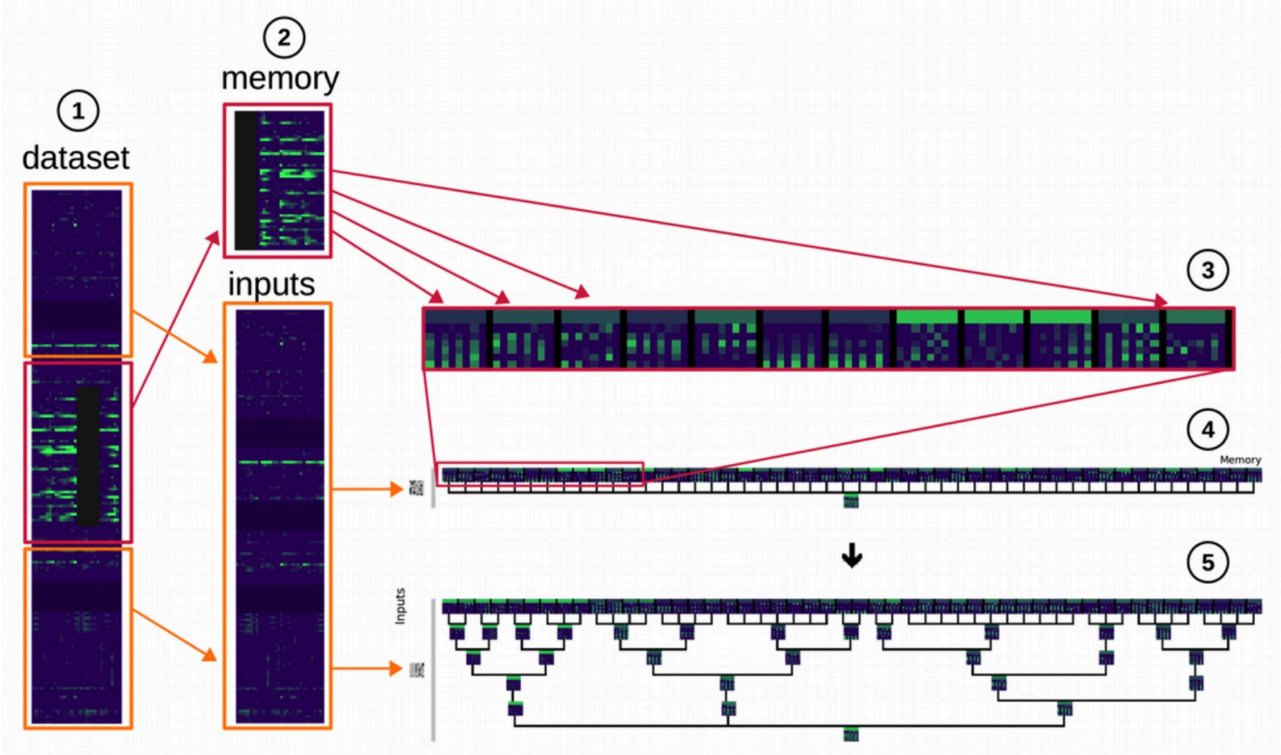

**C**

| Team | Score | Gene accuracy | Gene p-value | Time accuracy | Time p-value |
|---|---|---|---|---|---|
| **IntCyt** | **3.57** | **0.576** | **2.4E-06** | **0.489** | **2.9E-02** |
| GH | **3.25** | 0.563 | 6.5E-06 | 0.512 | 4.8E-02 |
| KNN | 3.18 | 0.558 | 1.1E-05 | **0.533** | 3.9E-02 |
| Team 263 | 1.85 | 0.421 | 7.5E-03 | 0.112 | 2.7E-01 |
| Team 297 | 1.68 | 0.333 | 5.6E-03 | 0.313 | 7.9E-02 |

**Fig. S7. Optimization of dynamic behavior by a nonequilibrium optimization.** *(related to Fig. 5)***.**

**(A)** Visualization of Intcyt outputs on MNIST images whose right sides are hidden with white noise. The panel shows the development of reconstructions and zoomed examples (lower subpanel) of the set of images by Intcyt for different compartmentalization parameters (we use $\omega = 0$ and $\omega = 30$, see Supplementary text, section 4b).
**(B)** Schematic showing the use of Intcyt on the DREAM3 dataset in order to impute partially-hidden gene expressions using the same type of technique as that used in panel **A**.
**(C)** Table showing the results of Intcyt on the DREAM3 dataset (bolded in red) and comparing them to the best results obtained by other algorithmic approaches for the challenge (bolded in black).

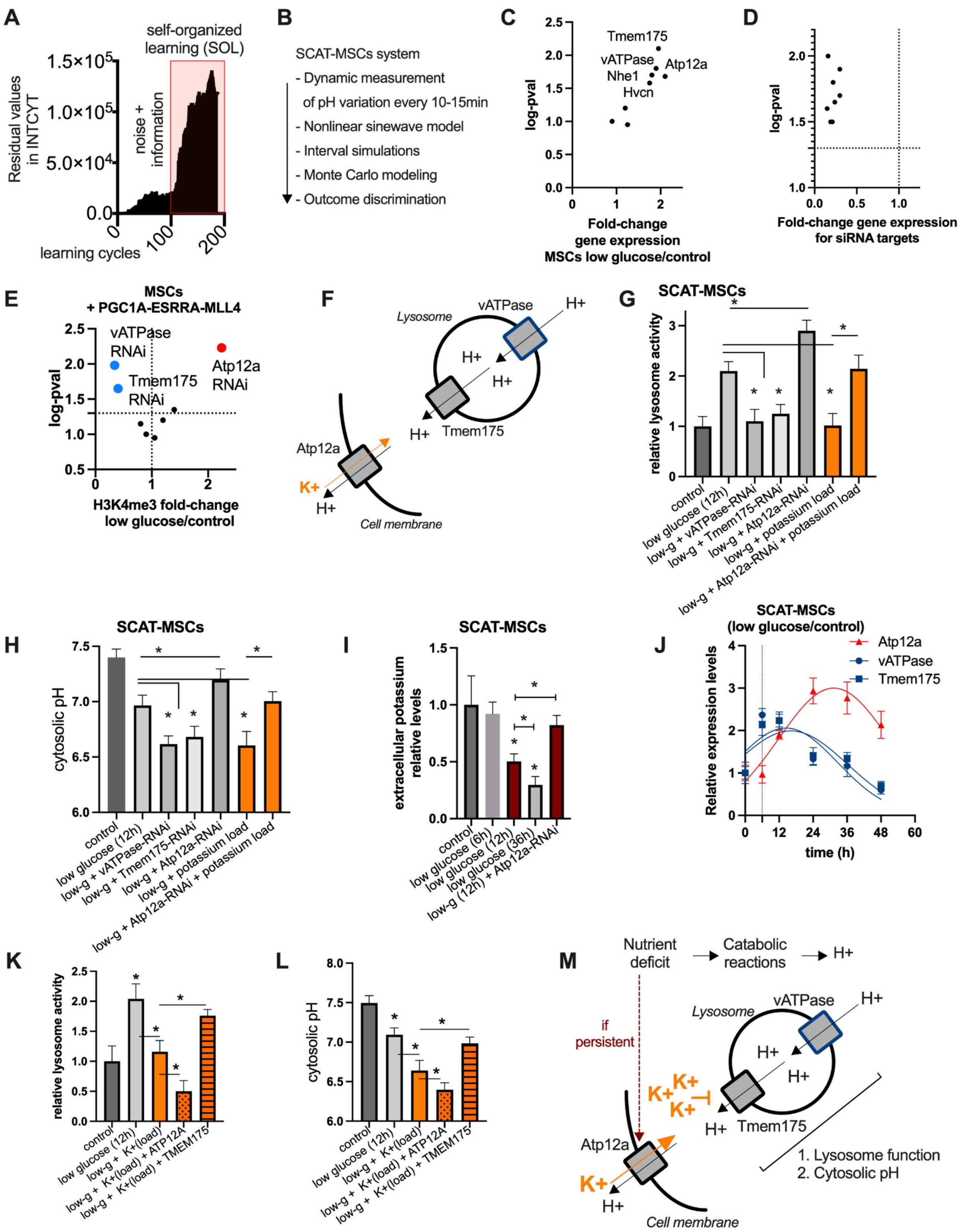
A
self-organized learning (SOL)
Residual values in INTCYT
noise + information
learning cycles
B
SCAT-MSCs system
- Dynamic measurement of pH variation every 10-15min
- Nonlinear sinewave model
- Interval simulations
- Monte Carlo modeling
- Outcome discrimination
C
Tmem175
vATPase
Atp12a
Nhe1
Hvcn
log-pval
Fold-change gene expression MSCs low glucose/control
D
log-pval
Fold-change gene expression for siRNA targets
E
MSCs + PGC1A-ESRRA-MLL4
vATPase RNAi
Tmem175 RNAi
Atp12a RNAi
log-pval
H3K4me3 fold-change low glucose/control
F
Lysosome
vATPase
H+
Tmem175
Atp12a
K+
Cell membrane
G
SCAT-MSCs
relative lysosome activity
control
low glucose (12h)
low-g + vATPase-RNAi
low-g + Tmem175-RNAi
low-g + Atp12a-RNAi
low-g + potassium load
low-g + Atp12a-RNAi + potassium load
H
SCAT-MSCs
cytosolic pH
I
SCAT-MSCs
extracellular potassium relative levels
control
low glucose (6h)
low glucose (12h)
low glucose (36h)
low-g (12h) + Atp12a-RNAi
J
SCAT-MSCs (low glucose/control)
Atp12a
vATPase
Tmem175
Relative expression levels
time (h)
K
relative lysosome activity
control
low glucose (12h)
low-g + K+(load)
low-g + K+(load) + ATP12A
low-g + K+(load) + TMEM175
L
cytosolic pH
M
Nutrient deficit
Catabolic reactions
H+
if persistent
Lysosome
vATPase
Tmem175
Atp12a
K+
Cell membrane
1. Lysosome function
2. Cytosolic pH

**Fig. S8. Nonequilibrium pH fluctuations mediated by lysosomal proton pumps.** *(related to Fig. 6)***.**

**(A)** Intcyt parameters throughout learning cycles. Panel shows residual values (see supplementary section 4c, subsection 1.5). Red box highlights self-organization of data by Intcyt (self-organized learning, or "SOL").
**(B)** Summary steps to measure and model dynamic pH variation in SCAT-MSCs. Time-frame measurement (every 10 to 15 min) of cytosolic pH variation in MSCs after glucose deficit. This is followed by the generation of nonlinear sine-wave models. This is then used for interval simulations (at least 10k steps) and by Monte Carlo approach to generate at least 500 models for discrimination of model features such as amplitude.
**(C)** Proton pumps gene expression screen (see supplementary materials) in SCAT-MSCs in low glucose for 12h.
**(D)** Gene expression knockdown efficiency for silenced proton pumps in SCAT-MSCs.
**(E)** H3K4me3 fold-change variation and log-p value of glucose deficient MSCs transfected with PGC1A, ERRA and MLL4 combined and with siRNAs for several proton pumps (H3K4me3 levels in mito-HAR genomic hub promoters of at least 15 regulated genes per group).
**(F)** Summary illustration showing known roles of identified proton pump.
**(G)** Relative lysosome activity in SCAT-MSCs comparing the effect of low glucose, proton pump knockdowns and potassium overload (25mM potassium).
**(H)** Cytosolic pH variation in SCAT-MSCs treated as in **G.**
**(I)** Extracellular potassium levels in SCAT-MSCs control and in low glucose alone (different time points) or with Atp12a proton pump knockdown.
**(J)** Relative expression levels of proton pumps in SCAT-MSCs in different time points of low glucose.
**(K)** Relative lysosome activity in SCAT-MSCs comparing the effect of potassium overload (25mM) in low glucose (2mM) and with transfection of either ATP12A or TMEM175.
**(L)** Cytosolic pH variation in SCAT-MSCs treated as in **K**.
**(M)** Summary illustration of proton pump roles during glucose deficit in SCAT-MSCs. vATPase and Tmem175 activity maintain lysosome function and cytosolic pH, whereas Atp12a reduces it by mediating the accumulation of cytosolic potassium and subsequent inhibition of Tmem175.
For all panels, cell experiments were done with 3 independent replicates. Data show mean values and SEM. Unpaired, two-tailed student's t-test was used when two groups were compared, and ANOVA followed by fisher's least significant difference (LSD) test for post hoc comparisons for multiple groups. * indicate p-value <0.05.

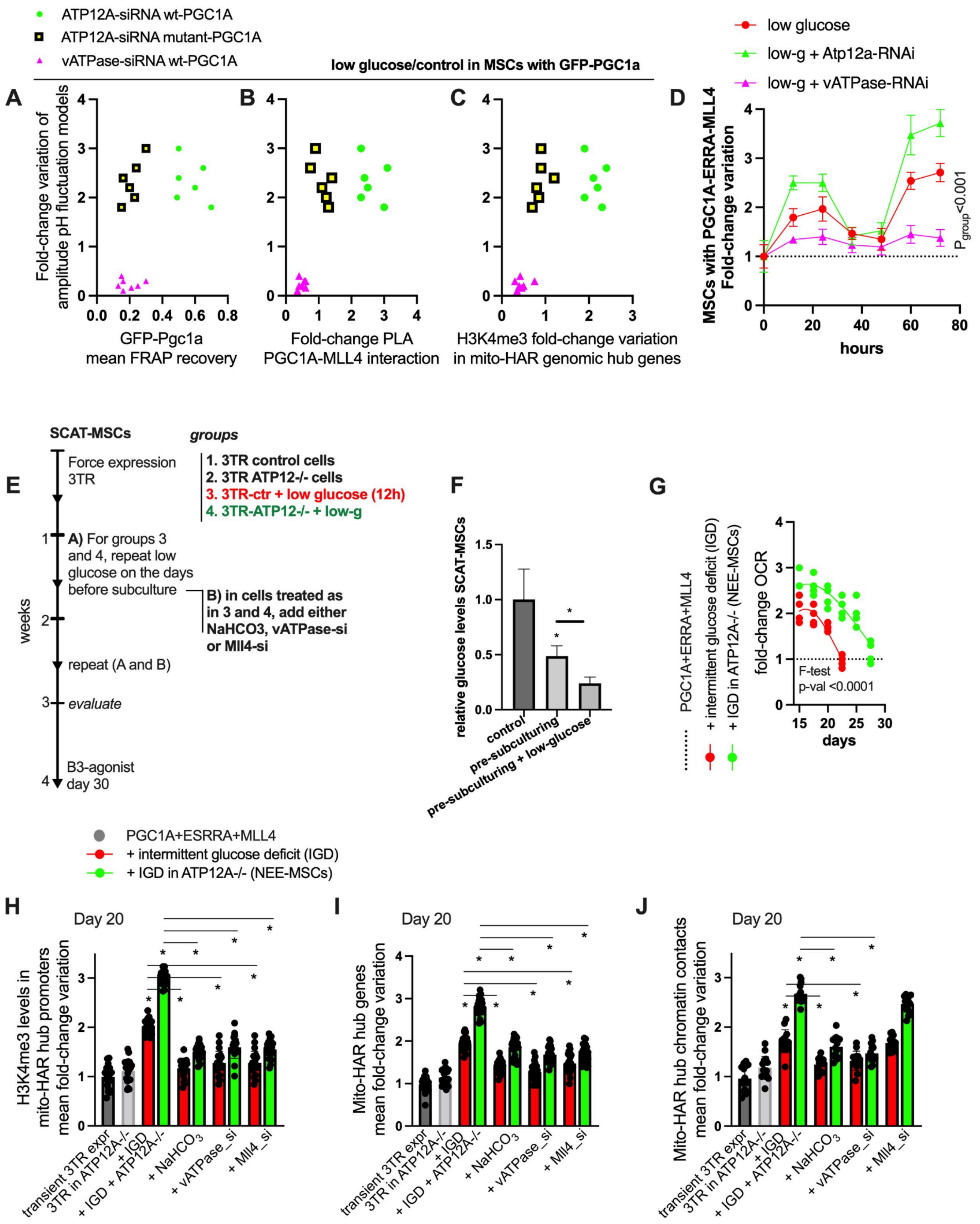
ATP12A-siRNA wt-PGC1A
ATP12A-siRNA mutant-PGC1A
vATPase-siRNA wt-PGC1A
low glucose/control in MSCs with GFP-PGC1a
A
B
C
D
Fold-change variation of amplitude pH fluctuation models
GFP-Pgc1a mean FRAP recovery
Fold-change PLA PGC1A-MLL4 interaction
H3K4me3 fold-change variation in mito-HAR genomic hub genes
low glucose
low-g + Atp12a-RNAi
low-g + vATPase-RNAi
MSCs with PGC1A-ERRA-MLL4 Fold-change variation
hours
$P_{group}$<0.001
SCAT-MSCs
groups
E
Force expression 3TR
1. 3TR control cells
2. 3TR ATP12-/- cells
3. 3TR-ctr + low glucose (12h)
4. 3TR-ATP12-/- + low-g
A) For groups 3 and 4, repeat low glucose on the days before subculture
B) in cells treated as in 3 and 4, add either NaHCO3, vATPase-si or Mll4-si
repeat (A and B)
evaluate
B3-agonist day 30
weeks
F
relative glucose levels SCAT-MSCs
control
pre-subculturing
pre-subculturing + low-glucose
G
PGC1A+ERRA+MLL4
+ intermittent glucose deficit (IGD)
+ IGD in ATP12A-/- (NEE-MSCs)
fold-change OCR
F-test
p-val <0.0001
days
PGC1A+ESRRA+MLL4
+ intermittent glucose deficit (IGD)
+ IGD in ATP12A-/- (NEE-MSCs)
H
I
J
Day 20
H3K4me3 levels in mito-HAR hub promoters mean fold-change variation
Mito-HAR hub genes mean fold-change variation
Mito-HAR hub chromatin contacts mean fold-change variation
transient 3TR expr
3TR in ATP12A-/-
+ IGD
+ IGD + ATP12A-/-
+ $NaHCO_3$
+ vATPase_si
+ Mll4_si

**Fig. S9. Nonequilibrium pH fluctuations modulate transcriptional adaptations in SCAT-MSCs.** *(related to Fig. 6)***.**

**(A)** SCAT-MSCs transfected with wild-type PGC1A or mutant-PGC1A in combination with either Atp12a or vATPase knockdowns. pH fluctuations were measured after 12h in low glucose media followed by Monte Carlo modeling and discrimination of fold-change variation (over unchallenged control) in amplitude parameters (Y-axis). This was plotted against mean droplet recovery by FRAP assays in cells treated similarly (data shows biological replicates).
**(B)** In cells as in **A,** amplitude of pH fluctuations vs mean fold-change variation of PGC1A-MLL4 protein interactions by proximity ligation assays (PLA; X-axis).
**(C)** In cells as in **A,** amplitude of pH fluctuations vs mean fold-change variation of H3K4me levels in mito-HAR genomic hub promoters (X-axis).
**(D)** Fold-change variation of gene expression (mitochondrial-HAR genomic hub genes) over time (hours) in SCAT-MSCs transfected with regulators comparing control (transfected and unchallenged) and low glucose (12h), in combination with either Atp12 or vATPase knockdowns (P-value for group comparison <0.01).
**(E)** Schematic illustration of perturbation experiments during cell culture. MSCs cells transfected with PGC1A, ERRA and MLL4 in combination (here representing control; 3TR, three transcriptional regulators), with normal (5mM glucose) and low glucose media (2mM) during overexpression (12h), and during the days before subculturing (every third day). This group was called intermittent glucose deficit (IGD). ATP12A-/- MSCs were also used with or without IGD. During the days before subculturing, we corrected pH by adding $NaHCO_3$ (1mM) and used Mll4-siRNA and vATPase-siRNA perturbation.
**(F)** Relative glucose levels in SCAT-MSCs in the days before and after subculturing and in low-glucose media.
**(G)** Time-frame fold-change variation of oxygen consumption rate (OCR) in SCAT-MSCs after combined transfection of transcriptional regulators, with intermittent glucose deprivation (IGD) alone or together with Atp12a perturbation (NEE; nonequilibrium-enhanced MSCs).
On day 20, MSCs as described in **E**, were collected and assayed for (**H**) H3K4me3 (mean fold change variation of H3K4me3 levels in promoters of at least 20 genes per group), (**I**) gene expression of mito-HAR hub genes (FC-variation of at least 20 genes per group), (**J**) chromatin contacts between mito-HAR genomic hubs (mean fold change variation of at least 15 contacts per group),
For all panels, Data show mean values and SEM. Unpaired, two-tailed student's t-test was used when two groups were compared, and ANOVA followed by Fisher's least significant difference (LSD) test for multiple groups. Two-way ANOVA was used to estimate significance between groups constrained by time-based measurements, and Tukey test for multiple comparisons. Nonlinear regression (Lorentzian-Cauchy model) and extra sum-of-squares F test to compare models between conditions for time-frame experiments. * indicates p-value <0.05.

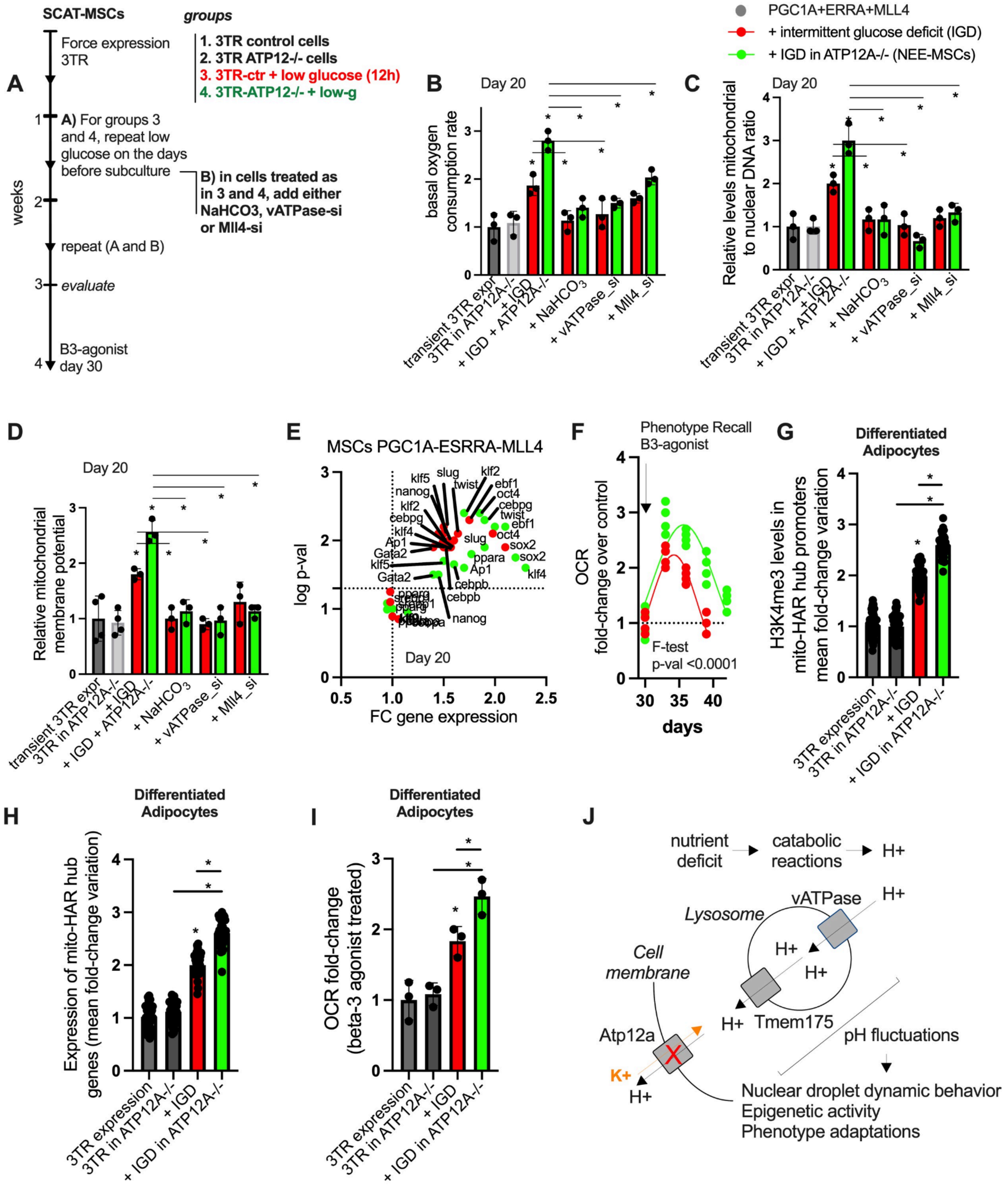
A
SCAT-MSCs
groups
Force expression 3TR
1. 3TR control cells
2. 3TR ATP12-/- cells
3. 3TR-ctr + low glucose (12h)
4. 3TR-ATP12-/- + low-g
A) For groups 3 and 4, repeat low glucose on the days before subculture
B) in cells treated as in 3 and 4, add either NaHCO3, vATPase-si or Mll4-si
repeat (A and B)
evaluate
B3-agonist day 30
weeks
PGC1A+ERRA+MLL4
+ intermittent glucose deficit (IGD)
+ IGD in ATP12A-/- (NEE-MSCs)
B
Day 20
basal oxygen consumption rate
C
Relative levels mitochondrial to nuclear DNA ratio
transient 3TR expr
3TR in ATP12A-/-
+ IGD
+ IGD + ATP12A-/-
+ NaHCO3
+ vATPase_si
+ Mll4_si
D
Relative mitochondrial membrane potential
E
MSCs PGC1A-ESRRA-MLL4
log p-val
FC gene expression
Day 20
F
Phenotype Recall B3-agonist
OCR fold-change over control
F-test p-val <0.0001
days
G
Differentiated Adipocytes
H3K4me3 levels in mito-HAR hub promoters mean fold-change variation
3TR expression
3TR in ATP12A-/-
+ IGD
+ IGD in ATP12A-/-
H
Differentiated Adipocytes
Expression of mito-HAR hub genes (mean fold-change variation)
I
Differentiated Adipocytes
OCR fold-change (beta-3 agonist treated)
J
nutrient deficit
catabolic reactions
H+
vATPase
Lysosome
Cell membrane
Tmem175
Atp12a
K+
pH fluctuations
Nuclear droplet dynamic behavior
Epigenetic activity
Phenotype adaptations

**Fig. S10. Modulating drive fluctuations enhances specialization and homeostasis.** *(related to Fig. 6)***.**

**(A)** Schematic illustration of perturbation experiments during cell culture. MSCs cells transfected with PGC1A, ERRA and MLL4 in combination (here representing control; 3TR, three transcriptional regulators), with normal (5mM glucose) and low glucose media (2mM) during overexpression (12h), and during the days before subculturing (every third day). This group was called intermittent glucose deficit (IGD). ATP12A-/- MSCs were also used with or without IGD. During the days before subculturing, we corrected pH by adding $NaHCO_3$ (1mM) and used Mll4-siRNA and vATPase-siRNA perturbation.
On day 20, MSCs as described in **A**, were collected and assayed for **(B)** basal oxygen consumption rate (OCR), **(C)** Relative mitochondrial/nuclear DNA ratio, and **(D)** relative mitochondrial membrane potential.
**(E)** Transcription factor expression profile in SCAT-MSCs transfected with transcriptional regulators as in **A,** in combination with IGD protocol (red color) or IGD in ATP12A-/- MSCs (green color).
**(F)** Fold-change variation of oxygen consumption rate (OCR) in the same conditions as in **A** and after treatment with beta-3 agonist on day 30 of cell culture.
Cells, treated as in **A**, were induced to adipocyte differentiation on day 9. On day 10 post-induction, adipocytes from engineered SCAT-MSCs were collected and assayed for (**G**) H3K4me3 (fold change variation in promoters of at least 40 genes per group), (**H**) gene expression (fold change variation of at least 40 genes per group), and (**I**) oxygen consumption rate (OCR) after beta-3 adrenergic receptor agonists CL-316,243 (10nM, Tocris) treatment (12h).
**(J)** Summary illustration of Atp12a perturbation effect on lysosome function, pH fluctuations and nuclear plasticity in MSCs during nutrient deficiency. Downstream role of fluctuating pH drives improving the dynamic behavior of droplets, epigenetic activity and phenotypic adaptations.
For all panels, cell experiments were done with 3 independent replicates. Data show mean values and SEM. Unpaired, two-tailed student's t-test was used when two groups were compared, and ANOVA followed by Fisher's least significant difference (LSD) test for *post hoc* comparisons for multiple groups. Nonlinear regression (Lorentzian-Cauchy model) and extra sum-of-squares F test to compare models between conditions for time-frame experiments. * indicates p-value <0.05.

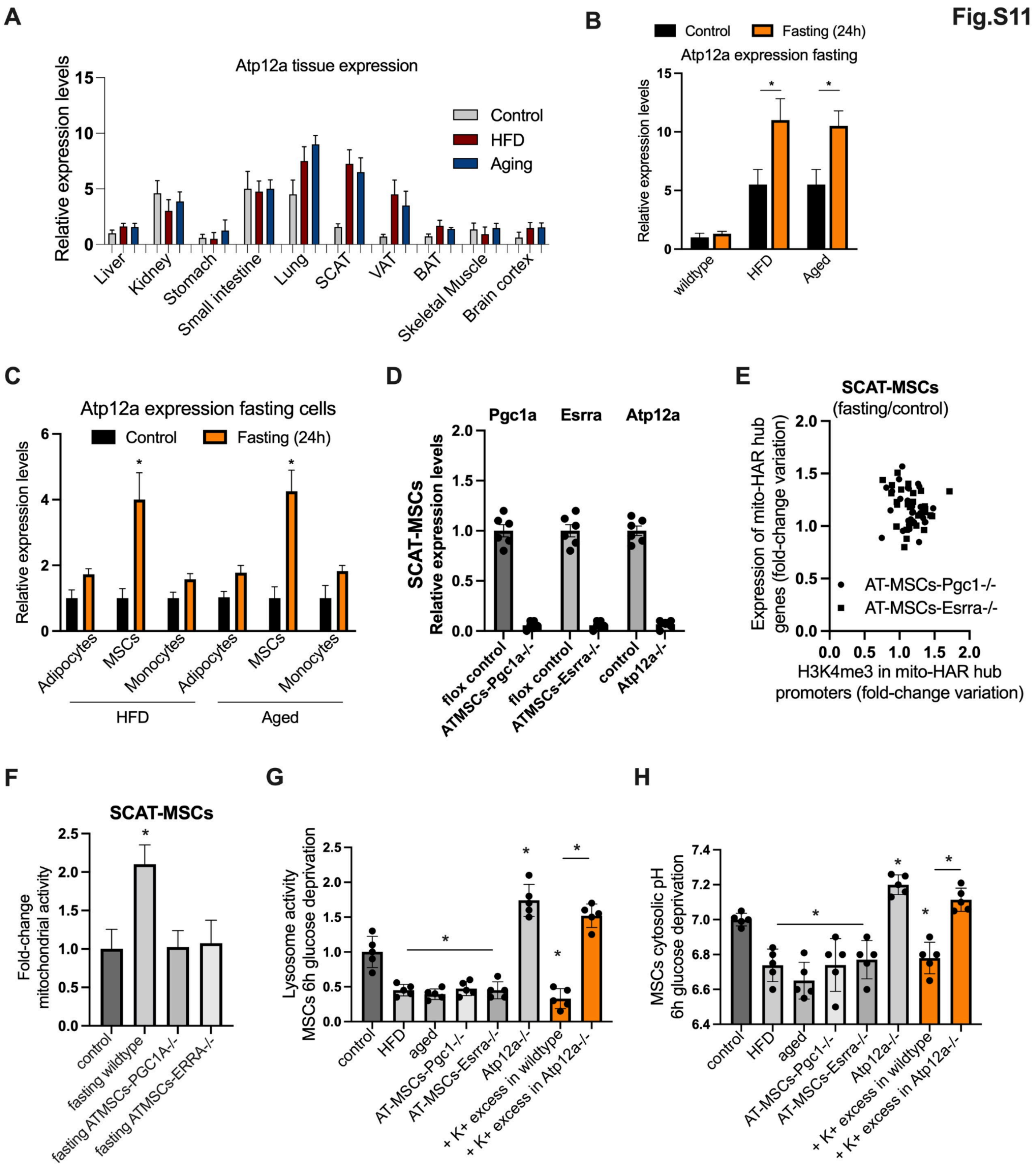
Fig.S11
A
Atp12a tissue expression
Relative expression levels
Control
HFD
Aging
Liver
Kidney
Stomach
Small intestine
Lung
SCAT
VAT
BAT
Skeletal Muscle
Brain cortex
B
Control
Fasting (24h)
Atp12a expression fasting
Relative expression levels
wildtype
HFD
Aged
C
Atp12a expression fasting cells
Control
Fasting (24h)
Relative expression levels
Adipocytes
MSCs
Monocytes
HFD
Aged
D
Pgc1a
Esrra
Atp12a
SCAT-MSCs
Relative expression levels
flox control
ATMSCs-Pgc1a-/-
ATMSCs-Esrra-/-
control
Atp12a-/-
E
SCAT-MSCs
(fasting/control)
Expression of mito-HAR hub genes (fold-change variation)
AT-MSCs-Pgc1-/-
AT-MSCs-Esrra-/-
H3K4me3 in mito-HAR hub promoters (fold-change variation)
F
SCAT-MSCs
Fold-change mitochondrial activity
control
fasting wildtype
fasting ATMSCs-PGC1A-/-
fasting ATMSCs-ERRA-/-
G
Lysosome activity MSCs 6h glucose deprivation
control
HFD
aged
AT-MSCs-Pgc1-/-
AT-MSCs-Esrra-/-
Atp12a-/-
+ K+ excess in wildtype
+ K+ excess in Atp12a-/-
H
MSCs cytosolic pH 6h glucose deprivation
control
HFD
aged
AT-MSCs-Pgc1-/-
AT-MSCs-Esrra-/-
Atp12a-/-
+ K+ excess in wildtype
+ K+ excess in Atp12a-/-

**Fig. S11. Evaluation of transcriptional compartment components *in vivo*** *(related to Fig. 7)***.**

**(A)** Relative Atp12a gene expression in different tissues from wildtype, HFD (8 weeks), and aged (18 months) mice models.
**(B)** Relative Atp12a gene expression in subcutaneous adipose tissue (SCAT) from mice as in A control and fasted for 24 hours.
**(C)** Relative Atp12a gene expression in sorted cells from SCAT from mice as in B.
**(D)** Relative gene expression in SCAT-MSCs from mice models including ATMSCs specific Pgc1a-/- and Esrra (driven by the Prx-CRE) and Atp12a-/- mice models.
**(E)** Fold-to-fold change plot of mitochondrial-HAR hub gene expression and H3K4me3 levels in regulated genes after fasting in sorted cells from subcutaneous adipose tissue from AT-MSCs-Pgc1a-/- and AT-MSCs-Esrra-/- mutant mice (fold-change variations of at least 20 genes per group).
**(F)** Mitochondrial activity (OCR) in sorted cells from mice described in **E**.
**(G)** Lysosome activity in extracted MSCs cells from subcutaneous adipose tissue from lean, HFD (8 weeks) and aged (18 months) wildtype mice, along with AT-MSCs-Pgc1a-/-, AT-MSCs-Esrra-/-, and Atp12a-/- mutant mice. Lysosome activity in MSCs was tested *ex vivo* after 6 hours of glucose deprivation. Atp12a-/- MSCs were additionally treated with potassium excess (25mM) during the challenge. Extracted cells from at least 4 mice per group were plated and assayed.
**(H)** Cytosolic pH after glucose deprivation challenge in MSCs from mice models as in **E**.
For all panels, animal experiments were done with n=5-8 mice per group, both female and male mice were included. Data show mean values and SEM. Unpaired, two-tailed student's t-test was used when two groups were compared, and ANOVA followed by Fisher's least significant difference (LSD) test for *post hoc* comparisons for multiple groups. * indicates p-value <0.05.

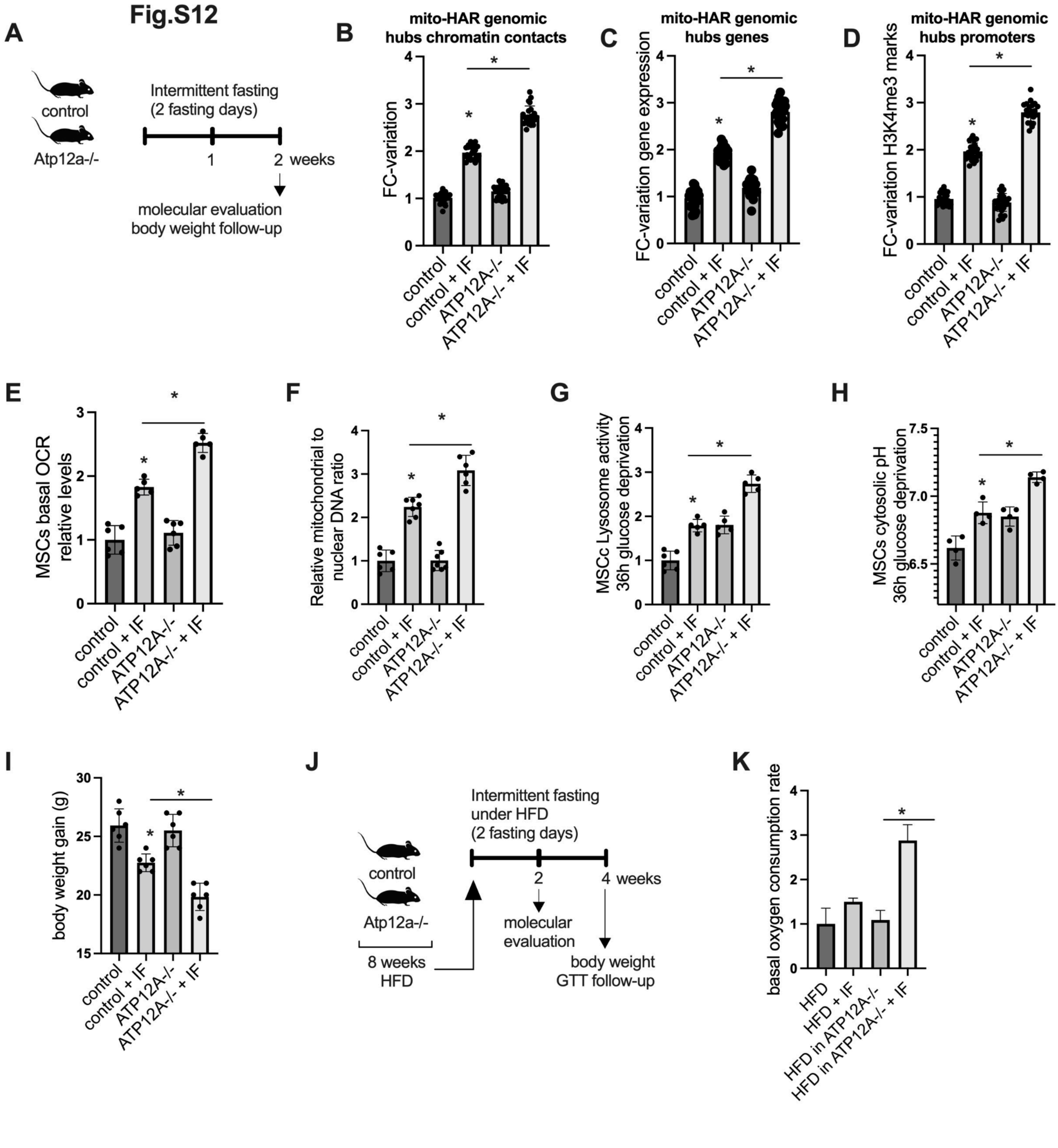

**Fig. S12. Nonequilibrium tuning enhances metabolic specialization.** *(related to Fig. 7)*.

**(A)** Schematic illustration of intermittent fasting (IF) intervention in lean and Atp12-/- knockout mice. Mice were fasted for 24 h on day 2 and day 5 of each of the two weeks. Molecular assays were performed after 2 weeks of IF intervention. Body weight was measured after the IF.

Sorted MSCs were evaluated for **(B)** 3C chromosome conformation assays of mito-HAR genomic hubs chromatin contacts (mean fold change variation of at least 15 contacts per group), **(C)** mito-HAR hub gene expression (mean fold-change variation of at least 20 genes per group), **(D)** H3K4me3 levels in mito-HAR genomic hub regulated genes (mean fold-change variation of H3K4me3 levels in promoters of at least 20 genes per group), and **(E)** basal oxygen

consumption rate (OCR, extracted cells from each mouse were counted, plated and assayed). Data shown as mean fold-change variation over control cells. (**F**) Relative mitochondrial/nuclear DNA ratio in sorted SCAT-MSCs. As in **E**, plated cells were assayed for lysosome activity (**G**) and cytosolic pH (**H**) after 36 h of in *ex vivo* glucose deficiency challenge (low glucose media).
(**I**) Body weight gain after IF intervention.
(**J**) Schematic illustration of intermittent fasting (IF) intervention in lean and Atp12-/- knockout mice receiving a high fat diet (8 weeks). Mice were fasted for 24 h on day 2 and day 5 of weeks 9 and 10. Mice models were still receiving HFD during this intervention period. Molecular assays were performed after 2 weeks of IF intervention (10 weeks total study time). Body weight was measured during 4 weeks after IF (12 weeks total study time).
**(K)** Stromal vascular fraction from each mouse as in **E** assayed for basal OCR.
For all panels, animal experiments were done with n=5-8 mice per group, both female and male mice were included. Cell experiments were done with 3 independent replicates. Data show mean values and SEM. Unpaired, two-tailed student's t-test was used when two groups were compared, and ANOVA followed by Fisher's least significant difference (LSD) test for *post hoc* comparisons for multiple groups. * indicates p-value <0.05.

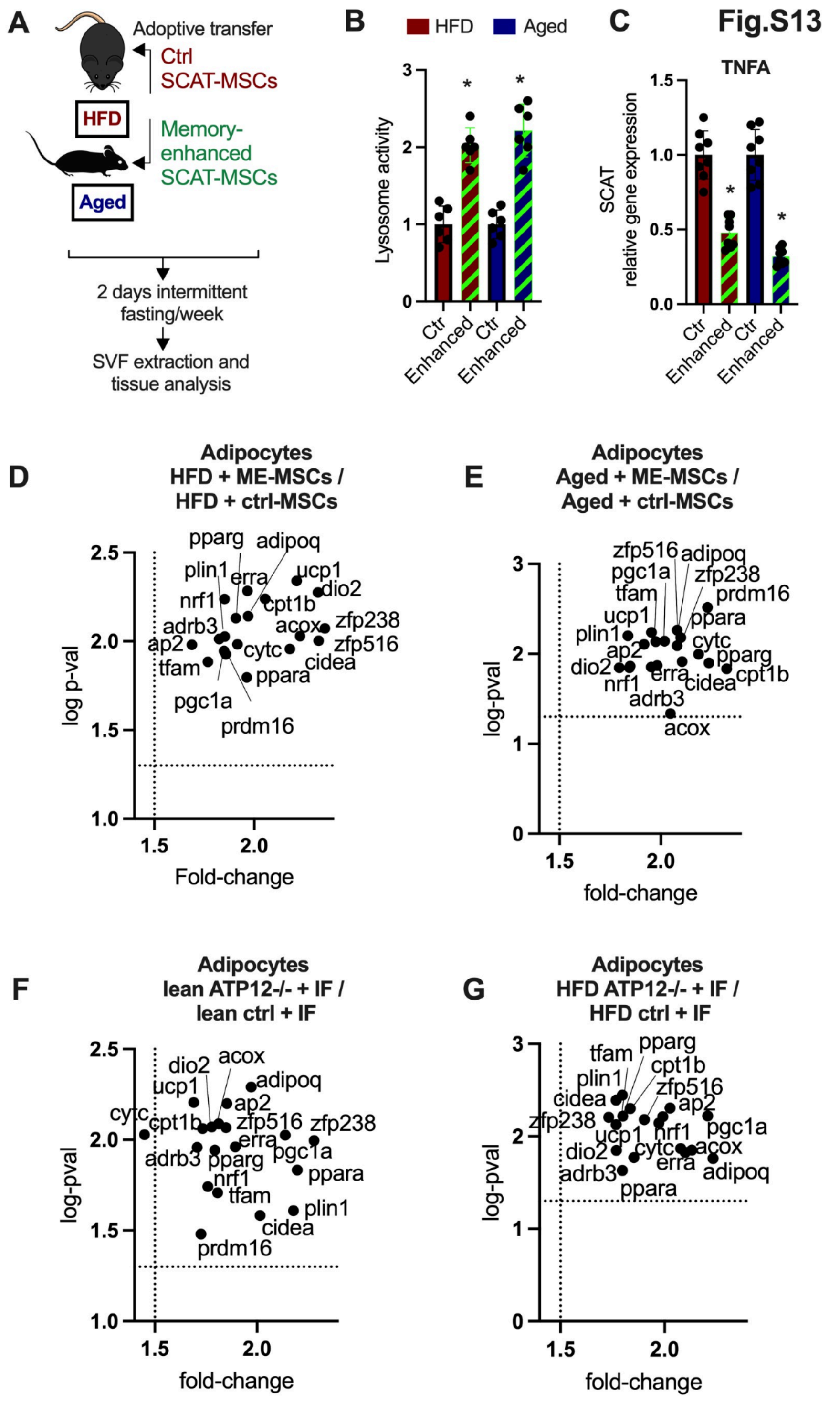

**Fig. S13. Nonequilibrium tuning, lysosome function, adipocyte thermogenic programs** *(related to Fig. 7)*.

**(A)** Nonequilibrium tuning strategy for evaluation of adoptive cell transfer to SCAT of HFD and aged wildtype mice. Memory-enhanced (ME) SCAT-MSCs are Atp12a-/-, overexpressed Pgc1a, Erra and Mll4, and underwent intermittent glucose deprivation (12h) previous each cell culture passage. Once transferred to SCAT, host mice received 2 days intermittent fasting per week.

**(B)** Stromal vascular fraction from each mouse assayed for lysosome activity. Data shown as mean fold change variation over control group.
**(C)** As in **A**, gene expression of tumor necrosis factor alpha (TNFA).
**(D, E)** Gene expression analysis in extracted adipocytes from mice models receiving treatments as in **A**.
**(F)** Gene expression analysis in extracted adipocytes from lean mice with cell-specific deletion of Atp12a over floxed (control) animals and after 2 weeks of intermittent fasting.
**(G)** Gene expression analysis in extracted adipocytes from HFD mice (8 weeks) with cell-specific deletion of Atp12a over floxed (control) animals and 2 weeks of intermittent fasting.
For all panels, animal experiments were done with n=5-8 mice per group, both female and male mice were included. Cell experiments were done with 3 independent replicates. Data show mean values and SEM. Unpaired, two-tailed student's t-test was used when two groups were compared, and ANOVA followed by fisher's least significant difference (LSD) test for post hoc comparisons for multiple groups. * indicate p-value <0.05.

#### Supplementary tables

##### Table S1. Metabolic phenotype comparison and human specific molecular signatures

1.1.A. HAR-associated genes clustered in genomic hubs and linked to metabolic function and nuclear-encoded mitochondrial genes from mitocarta (last version)
1.1.B. Mitochondrial and metabolic HAR genomic hubs
1.1.C. link

##### Table S2. Intcyt evaluation.

| Method (raw pixels) | MNIST test recall | Fashion-MNIST test recall |
|---|---|---|
| K-NN | 96.9% | 85.4% |
| Linear SVM | 91.7% | 83.6% |
| Random forest | 97% | 87.3% |
| *2-conv CNN (<100k params; DL)* | *99%* | *93%* |
| **Intcyt (ours)** | **98%** | **92%** |
| ***Intcyt structural ablation*** | **~65-90%** | **~45-70%** |
| ***Intcyt residual ablation*** | **~85-94%** | **~75-82%** |

**Table S2. Test recall evaluation**: Baseline results (K‑NN, linear SVM, random forest, 2‑conv CNN (<100k params)) are approximate (≈) representative values from published benchmarks on MNIST and Fashion‑MNIST using the standard train/test split and raw‑pixel inputs (see references; protocols vary by source). Intcyt was evaluated on the held‑out test set and results are shown as mean over independent runs. "Test recall" denotes single‑label multi‑class recall (macro-averaged recall or mean per class recall; equivalent to overall accuracy in this setting). Intcyt values at baseline (no ablation) are ± SD 0.6. Ablation experiments were conducted with K=10 and estimate variation is represented in the table. Benchmark references: K-NN and Linear SVM on MNIST and Fashion-MNIST; (Xiao, H., Rasul, K., & Vollgraf, R., Arxiv, 2017), (LeCun, Y., Bottou, L., Bengio, Y., & Haffner, P., Proceedings of the IEEE, 1998); Random forest on MNIST and fashion-MNIST (Wang, Y., et al., Scientific Data, 2022), (Xiao et al, Arxiv, 2017); 2-conv CNN (LeCun Y. et al., 1995), (Benchmarking computer vision algorithm for Fashion MNIST and MNIST dataset at https://github.com/timothylimyl/FASHION-MNIST).

#### Biological and Statistical Materials and methods

##### Animal studies

All animal experiments were performed in accordance with internationally accepted guidelines and principles for the use of laboratory animals, and were approved by the Novo Nordisk Research Center Seattle Institutional Animal Care and Use Committee, Novo Nordisk research center Lexington Massachusetts, the Novo Nordisk Ethical Review Committee, Novo Nordisk Research Center in China, MIT committee of animal care, and regional ethics committee of Alicante, and Universidad Miguel Hernández-CSIC, Spain. Adult wild-type male C57BL/6J mice aged 6-12 weeks or 18 months were obtained from Jackson Laboratories (Stock # 000664). Perturbation in adipose precursor cells was generated as previously described (*1*). Prrx1-CRE mouse line (jackson laboratories, strain # 005584) was bred with Pgc1a loxp mice (jackson laboratories, strain #:009666), and Esrra loxp mice (jackson laboratories, strain #034713). Atp12a knockout mice were generated as previously described (*2*) and are part of the knockout mouse project (MMRRC_046720-UCD, C57BL/6N-Atp12atm1(KOMP)Vlcg/MbpMmucd).

Mice were housed 4-8 per cage with ad libitum access to water and food (Type IV Scanbur Cages; 1800 $cm^2$ floor area). Animal holding rooms temperature ranged from 20-26 °C and humidity ranged from 30-70%. Lighting was on an automated light/dark cycle with lights on between 6:00 AM and 6:00 PM. Mice were provided with Alpha Dri bedding material to just cover the bottoms of the cage, and one Nestlet square. Environmental enrichment included a plastic hut, a paper hut was provided. Mice received a hydrogel cup upon arrival until familiarized with the automatic water system. Standard rodent chow diet (PicoLab Rodent Diet 20 Lab Diet #5R53) was provided to the mice upon arrival until the diet was switched from chow to high fat (described below). All food and water were available ad libitum unless indicated. Mice were acclimated to housing for one week.

High fat diet: At 6 weeks of age wild-type mice were placed on a 60% high fat diet (High-Fat Diet #D12492 (20.0% Protein, 60% Fat, 20% Carbohydrate) during 8 weeks until they reached dietary induced obese status (DIO). Similarly, wild-type, and Atp12-/- mice received a high-fat diet for 8 weeks. Mice protocols and treatments: wild-type, mutant mice, aged mice (18 months) and mice undergoing HFD were used for a fasting challenge of 24h, after which tissue extraction was performed. Control and Atp12-/- mice undergoing HFD of 8 weeks were subjected to an intermittent fasting protocol (2-4 weeks), consisting of 2 intercalated days per week (Tuesdays and Fridays) without food access (24 hours each day).

##### Cell isolation and sorting

Adipocyte progenitors and mesenchymal stem cells from adipose tissue: To isolate adipose progenitors from subcutaneous adipose tissues, adipose tissue dissociation kit (Miltenyi Biotec) and adipose tissue progenitor isolation kit (Magnetic-activated cell sorting, Miltenyi Biotec) were used following manufacturer's instructions. The CD271 microbead kit (Miltenyi biotec) was used for isolation of adipose tissue MSCs (AT-MSCs) following manufacturer's instructions.. Adipocytes: Isolation of adipocytes was performed during the SVF extraction with some modifications during the pelleting steps. In brief, after incubating in a vigorous shaking water bath for 20-30 minutes at 37 C, cells disintegrate into individual cells. Then, centrifugation at 500g (2200 RPM) for 10 minutes was used to separate floating adipocytes from pelleted SVF. Aspiration of the floating fat cell layer using 1000 uL pipette tip (cut), very carefully remove the floating adipocyte layer to a fresh tube with a warm cell buffer (if performing further experiments) or SDS-lysis buffer. Cd11b+ monocyte/macrophage cells from adipose: After adipose tissue dissociation, samples are processed as described below following magnetic-activated cell sorting strategies. Monocytes/macrophages are isolated by the digestion with pronase and collagenase digestion followed by discontinuous gradient ultracentrifugation as described above. After initial centrifugation at 600 rpm, the supernatant cell fraction (containing non-parenchymal cells including immune fraction) is further centrifuged at 150rpm for 5 minutes, and washed with DMEM medium. After washing and centrifugation, the pellet cell fraction is suspended in 90 µl MACS buffer (1xPBS, 0.5% BSA) containing 0.6% citrate-dextrose solution (Sigma). Then, 10 µl of CD11b MicroBeads (Miltenyi Biotec, #130-049-601) are added to the cell suspension and incubated for 20 minutes at 4 degree celsius. During incubation, place a MACS LS column (Miltenyi Biotec, #130-042-401) on MACS MultiStand and wash the column with 3 ml PBS twice and then with 3 ml MACS buffer. The cell suspension is passed to the column to remove CD11b-negative cells. 5 ml of MACS buffer is applied onto the column to collect CD11b+ cell fraction.

**Mammalian cell culture and treatments**

Murine subcutaneous adipocytes: SVF from inguinal fat depots of C57BL/6J wild-type and mutant mice (when indicated) were isolated and prepared as previously described (*4*). In brief, at 90% confluence, cell were induced with adipogenic induction media containing 10% FBS, 1% pen-strep, 100 nM dexamethasone (Sigma), 0.4 isobutyl methyl xanthine (IBMX, Sigma), 3 ug/ml insulin (Sigma) and 1uM rosiglitazone (Tocris). The differentiation cocktail was used during the first 2 days of culture, followed by rosiglitazone and insulin for 6 to 10 days.

Murine subcutaneous adipose tissue mesenchymal stem cell cultures: Once AT-MSCs were extracted, cells were plated in collagen coated 12 well plates (2x10e5 cell/well), they were maintained in StemPRO medium (Thermofisher) supplemented with glucose (5.5mM). Cells were maintained for 4 to 12 passages (when indicated). Passaging was performed every 72h during which cells were counted and re-plated to the same density. Cells were maintained as previously described (*5*), and AT-MSCs were tested for their cell differentiation ability into adipocyte lineages.

Cell treatments: Indicated cell cultures were kept in different condition media: Dulbecco's with low glucose (2mM). For RNAi screens cells were transfected using lipofectamine 3000 (Invitrogen, Thermo fisher technologies) with siRNA for histone methyltransferases (Set1a, setdb1, set7, setd4, mll1, mll2, mll3, mll4, nsd1, ezh1), histone demethylases (kdm2b, kdm6a, kdm6b, kdm4b), proton pumps and channels (vATPase, tmem175, hvcn1, atp12a, slc9a1, slc9a2, slc9b1, slc9b2), and scramble control (Applied biosystems).
When indicated, cells were treated with oligomycin (1uM; complex V inhibitor, Sigma Aldrich), 1µM FCCP (maximal respiration inducer, Sigma Aldrich), or 1µM antimycin (respiration inhibitor, Sigma Aldrich), beta-3 adrenergic receptor agonists CL-316,243 (10nM, Tocris), potassium chloride solution (25mM potassium, Tocris), and sodium bicarbonate (1mM, Sigma aldrich).

Constructs and plasmids for transfection: wild-type expressing plasmids for murine GFP-PGC1A (GFP-PGC1 was a gift from Bruce Spiegelman, Addgene plasmid # 4, RRID:Addgene_4). Wild-type expressing plasmids for murine ERRA (pcDNA3.1(+)-C-6His-mEsrra was a gift from Brijesh Kumar Singh (Addgene plasmid # 173152; RRID:Addgene_173152). For MLL4 (kmtd2) constructs, cDNAs encoding kmt2d were amplified from murine cells, and cloned into pcDNA3.1 (+)-C-6His backbone plasmid (addgene). For mutant PGC1A constructs, cDNAs for encoding regions of interest were amplified from murine cells and cloned into pEGFP-C1 backbone plasmid (addgene). These mutant constructs exclude IDR2 regions identified in silico. For PGC1A constructs without GFP-tag, cDNAs encoding full PGC1A sequence was amplified and cloned into pcDNA3.1 (+)-C-6His backbone plasmid (addgene; for molecular experiments testing the effect of GFP on protein interactions). For ATP12A and TMEM175 expression constructs, cDNAs were amplified from murine cells, and cloned into pcDNA3.1 (+)-C-6His backbone plasmid (addgene). The sequences of all plasmids were confirmed by Sanger sequencing.
Plasmid transfection: plasmids were transfected to adipose MSCs cells using Amaxa-nucleofector and nucleofector kits for transfection of primary cell lines following manufacturer's protocols (Lonza). Cells were transfected with different plasmids when they reached 70-80% confluence after 1-3 passaging. For co-transfection of multiple plasmids, premixing of plasmid DNAs (same concentration) was performed prior to transfection. siRNA molecules were transfected using Lipofectamine 3000 (Invitrogen, Thermo Fisher Scientific) and removed 24 h post-transfection, according to standard manufacturer protocols. Transfected cells were assayed post-transfection with protein identification and other functional assays when indicated.

SCAT-MSCs transcriptional engineering: AT-MSCs from mice models were isolated and maintained in vitro for 3 days in 12 well plates to cell density of 2x10e5 cell/well. On day 3 cells were dissociated with Trypsin (Sigma aldrich) and resuspended in solution. AT-MSCs cells were transfected in suspension with the Amaxa-nucleofector system (Lonza) and recommended primary cell nucleofector kits (Lonza). Cells were transfected with expressing plasmid vectors for PGC1A, ERRA and MLL4 individually or in combination. When indicated, 12 hours after transfection the media was changed to low glucose media (2mM) for 12h. Protein variation and transfection efficiency were tested using proximity ligation assays as indicated (see PLA assays). When indicated, cells were maintained 4-12 passages, with passaging (sub-culturing) days every 3rd day, where the cells were resuspended, counted and plated. When indicated, 12h prior sub-culturing, the media of the cells was changed to low glucose media. When indicated, cells were transfected with si_RNA during subculturing. Control cells were maintained as indicated and were transfected with empty expression vectors. Molecular and functional assays were performed in modified and control cells.

**Analysis of Gene Expression**

Total RNA was isolated from cells or tissues using Isol-RNA Lysis Reagent (5 PRIME) according to manufacturer's instructions. Amplification Grade DNase I (Life Technologies) was used to treat 1 μg of RNA, from which, 500 ng were used for cDNA preparation following Applied Biosystem Reverse Transcription Kit (Life Technologies). Quantitative Real-Time PCR was performed in a ViiA 7 and QuantStudio Real-Time PCR system thermal cycler with SYBR Green PCR Master Mix (both Applied Biosystems). Analysis of gene expression was performed using the ΔΔCt method and relative gene expression was normalized to hypoxanthine phosphoribosyltransferase (HPRT) mRNA levels. When indicated gene expression analyses were shown as mRNA levels relative to controls. To evaluate transcriptional compartment activity of genes within genomic hubs, mean fold-change variation over control for each gene was used to compare overall effects (displayed as a module) for different biological conditions. Primer sequences available upon reasonable request.

**Chromatin immunoprecipitation**

ChIP experiments were performed as previously described (*6*) with the following modifications: Cells were plated and (2-10 million) pooled before homogenization. Cells were homogenized in PBS and crosslinked with 1% formaldehyde during 10min. For tissue, 50mg were homogenized in PBS and crosslinked with 1% formaldehyde for 10 min. Glycine (125 mM) was added, followed by centrifugation at 4000 rpm for 5 min. After aspirating the supernatant, the pellet was washed with 1 ml cold PBS followed by centrifugation at 4000 rpm for 5 min.

This step was repeated with buffer 1 (0.25% Triton X-100, 10mM EDTA, 0.5mM EGTA, 10mM HEPES, pH 6.5) and buffer 2 (200mM NaCl, 1mM EDTA, 0.5mM EGTA, 10mM HEPES, pH 6.5). The pellet was then resuspended in 300 μl lysis buffer (1% SDS, 10mM EDTA, 50mM Tris-HCl, 1X protease cocktail inhibitor, pH 8). Samples were sonicated on ice using a Bioruptor (Diagenode) 30 times for 30 seconds ON and OFF intervals. After the lysates were sonicated to shear the DNA to fragment lengths of 300bp, the complexes were immunoprecipitated with specific antibodies. 10-20 ul of antibody was used for each immunoprecipitation. No antibody controls were also included for each ChIP assay and no precipitation was observed by quantitative Real-Time PCR (qPCR) analysis.

Input samples were processed in parallel. The antibody/protein complexes were collected by either salmon sperm DNA/protein A agarose slurry or Protein A/G PLUS agarose beads (Santa Cruz sc-2003) and washed several times. The immuno-complexes were eluted with 1% SDS and 0.1 M NaHCO3 and samples were treated with proteinase K for 1 hour and DNA was purified by phenol/chloroform extraction, ethanol precipitation and resuspended in 20 ul H2O. Input DNA and immunoprecipitated DNA were analyzed by quantitative PCR. Primer sequences used to detect each DNA region are available upon reasonable request. To evaluate transcriptional compartment activity in the form of promoter binding by transcriptional regulators within genomic hubs, mean fold-change variation over control for each promoter binding was used to compare overall effects (displayed as a group) for different biological conditions. Primer sequences available upon reasonable request.

**Chromosome conformation assays**

<u>In situ ChIP-loop immunoprecipitation of chromatin interactions:</u> Targeted ChIP-loop-qPCR protocol is based on ChIP immunoprecipitation assay previously described (*7*) with the following modifications: In brief, 1 to 15 million crosslinked cells were resuspended in lysis buffer ( 60mM Tris pH 7.5 0.5% Igepal, 0.25% Sodium-deoxycholate 0.1% SDS, 150mM NaCl, and protease inhibitors) and rotated at 4° C for 30 minutes. Nuclei were pelleted at 4° C for 5 minutes at 2500 rcf. Pelleted nuclei were washed once with 500 μL of ice-cold lysis buffer. Supernatant was removed again and the pellet was resuspended in 100 μL of 0.4% SDS and incubated at 62° C for 30 minutes. To quench the SDS, 10% Triton X-100 were added and samples were rotated at 37° C for 15 minutes. NEB Buffer 2 and 15 μL of 25 U/μL HindIII (NEB; following manufacturer's recommendation were then added and samples were rotated at 37° C for 2 hours (15 μL was used for 10–15 million cells, 8 μL for 5 million cells, and 4 μL for 1 million cells).

The enzyme was then heat-inactivated at 62° C for 20 minutes. 52ul incorporation master mix was then added: 37.5 μL of 0.4 mM biotin-dATP (Thermo Fisher 19524016), 4.5 μL of a dCTP, dGTP, and dTTP mix at 10 mM each, and 10 μL of 5 U/μL DNA Polymerase I, Large (Klenow) Fragment (NEB, M0210). The reactions were rotated at 37° C for 1 hour. 948 μL of ligation master mix was then added: 150 μL of 10X NEB T4 DNA ligase buffer with 10 mM ATP (NEB, B0202), 125 μL of 10% Triton X-100, 3 μL of 50 mg/mL BSA (Thermo Fisher AM2616), 10 μL of 400 U/μL T4 DNA Ligase (NEB, M0202), and 660 μL of water. The reactions were then rotated at room temperature for 4 hours.

After proximity ligation, the nuclei with in-situ generated contacts were pelleted at 2500 rcf for 5 minutes at room temperature. As described above, the pellet was then resuspended in 300 μl lysis buffer (1% SDS, 10mM EDTA, 50mM Tris-HCl, 1X protease cocktail inhibitor, pH 8). Samples were sonicated on ice using a Bioruptor (Diagenode) 20 times for 15 seconds ON and OFF intervals. Clarify samples for 15 minutes at 16100 rcf at 4 degrees and w2X volume of ChIP Dilution Buffer (0.01% SDS, 1.1% Triton X-100, 1.2 mM EDTA, 16.7 mM Tris-HCl pH 7.5, 167 mM NaCl) was added.

The complexes were immunoprecipitated with 10-20 ul of specific antibodies (for each immunoprecipitation). Complexes were immunoprecipitated with PGC1A or ERRA specific antibodies when individual transfections were performed. For combined transfections, digested complexes were immunoprecipitated with PGC1A (see chromatin immunoprecipitation and antibodies sections). 60 μL of Protein A beads (Thermo Fisher) were precleared. Beads were washed and resuspended in ChIP Dilution Buffer to a volume of 50 μL per tube (100 μL per sample), then added to samples and rotated at 4° C for 1 hour. Samples were placed on a magnet and supernatants were transferred into fresh tubes. Beads were washed three times each with Low Salt Wash Buffer (0.1% SDS, 1% Triton X-100, 2 mM EDTA, 20 mM Tris-HCl pH 7.5, 150 mM NaCl), High Salt Wash Buffer (0.1% SDS, 1% Triton X-100, 2 mM EDTA, 20 mM Tris-HCl pH 7.5, 500 mM NaCl), and fresh LiCl Wash Buffer (10 mM Tris-HCl pH 7.5, 250 mM LiCl, 1% NP-40, 1% sodium deoxycholate, 1 mM EDTA).

The antibody/protein complexes were collected by Protein A/G PLUS agarose beads (Santa Cruz) and washed several times. The immuno complexes were eluted with 1% SDS and 0.1 M $NaHCO_3$ and samples were treated with proteinase K for 1 hour and DNA was purified by phenol/chloroform extraction, ethanol precipitation and resuspended in purified H2O. After reverse cross-linked, ligated DNA was purified and sheared to a length of roughly 250-500 base pairs, at which point ligation junctions were pulled down with streptavidin beads and prepped for targeted quantitative measurement of interactions using qPCR assays comparing different samples. Ligation efficiencies were determined using qPCR assays comparing control templates. Specific primers for computationally selected genomic regions were designed to validate chromatin-chromatin contacts. This was performed using top active genes in each hub, followed by targeted 3C evaluation of promoter-promoter interactions, guided by GeneHancer or NCBI promoter coordinates. To evaluate transcriptional compartment activity in the form of chromatin contacts between genomic hubs, mean fold-change variation over control for each chromatin contact was used to compare overall effects (displayed as a group) for different biological conditions. Primer sequences available upon reasonable request.

<u>In situ 3C chromosome conformation assays</u>: Between 15 thousand and 2 million cells were used for in situ chromosome conformation assays as previously described (*8*) with some modifications. Briefly, cells were crosslinked with 1% formaldehyde for 10-15 minutes at room temperature. Cells were lysed with 60mM Tris pH 7.5 0.5% Igepal, 0.25% Sodium-deoxycholate 0.1% SDS, 150mM NaCl, and protease inhibitors. Pelleted nuclei were resuspended in 0.4% SDS and incubated 30 min 60°C for permeabilization. After quenching with 1% Triton-X for 12 min at 37°C, nuclei were digested with 5U/μl DpnII in 1x DpnII buffer overnight at 37°C. Cells were then pelleted (2500g 10min), and the restriction enzyme was inactivated at 65°C for 20 min. For the 1.5hr fill-in at 37°C, biotinylated dATP was used for ligation at 25°C for 4 hours with rotation.

Nuclei were pelleted and sonicated in 10mM Tris pH 7.5, 1mM EDTA, 0.25% SDS for 16 min, 100 intensity, 200 cycles per burst, and max temperature of 5°C. DNA was reverse cross-linked overnight at 65°C with proteinase K and RNAse A. Each experiment was performed in biological replicates. After reverse cross-linked, ligated DNA was purified and sheared to a length of roughly 250-500 base pairs, at which point ligation junctions were pulled down with streptavidin beads and prepped for targeted quantitative measurement of interactions using qPCR assays comparing different samples. Primer sequences available upon reasonable request.

Data was represented as the fold-change of interaction over control (e.g. fasting/control), where statistically significance and fold-variations were shown as indicated in figure legends. To evaluate transcriptional compartment activity in the form of chromatin contacts between genomic hubs, mean fold-change variation over control for each chromatin contact was used to compare overall effects (displayed as a module) for different biological conditions. This was performed using top active genes in each hub, followed by targeted 3C evaluation of promoter-promoter interactions, guided by GeneHancer or NCBI promoter coordinates. Primer sequences available upon reasonable request.

**Antibodies**

Anti-PGC1A (sigma-aldrich calbiochem, catalog No. ST1204), Anti-PGC1 alpha antibody N-terminal (abcam, ab191838), anti-ERRA antibody N-terminal (Abcam, anti-ERRA EPR46Y, catalog No. ab76228), anti-KMT2D/MLL4 (antibodies online, catalog No. ABIN6944095), anti-H3K4me3 (Diagenode, catalog No. C15410030), anti-H3K27me3 (recombinant anti-Histone H3 (tri methyl K27) antibody EPR18607, catalog No. ab192985), Anti-GFP antibody (ab290), Anti-6X His tag antibody [HIS.H8] (ab18184).

**Commercial assays**

Triglyceride assay kit (Abcam ab65336), intracellular pH indicator (pHrodo Red AM, fluorescent, Thermo fisher), hydroxyproline assays (ab222941), mitochondrial isolation kit (ab110168), lysosome activity assay (ab234622), Mitochondrial membrane potential (JC-1, ab113850), Lipid oil red staining kit (plate reader cat#MAK194), Senescence assay kit (beta galactosidase, fluorescence, abcam catalog No, ab228562), and potassium fluorometric assay kit (Abcam, ab252904). Measuring glucose levels in media from cell cultures was performed using the GlucCell monitoring system (Gluccell, CLS-1322-02).

**Proximity ligation assays**

Cell or tissues lysates were obtained using RIPA buffer (50 mM tris pH 7.5, 150 mM NaCl, 1 mM EDTA, 1% (w/v) Triton-X-100, 0.5% (w/v) Na-deoxycholate, 0.1% (w/v) sodium dodecyl sulfate, 20 mM glycerol-2-phosphate, 5 mM sodium pyrophosphate) freshly supplemented with 1 mM DTT and 0.5 mM PMSF. Antibodies were conjugated with oligonucleotides using the oligonucleotide conjugation kit for proximity ligation assays (PLAs, ab218260). Protein concentration was determined by Bradford Protein Assay following manufacturer's instructions (BioRad). Protein extracts and conjugated antibodies were used for PLA in which qPCR on conjugated products was performed following manufacturer's instructions. For homogeneous PLA assay (individual proteins) antibodies against tag proteins (GFP or His tags) and proteins of interest (PGC1A, ERRA, and MLL4) were conjugated with 3' or 5' oligonucleotide probes each, followed by immuno-qPCR according to manufacturer's instructions.

**Live cell imaging and Fluorescence Recovery After Photobleaching (FRAP)**

FRAP was performed using an inverted confocal super-resolution microscope Zeiss LSM 880-Airyscan Elyra PS.1 with a 63x oil immersion objective. Live-cells were maintained in the microscope chamber at 37ºC and 5% $CO_2$. A circular region of interest (ROI) 23 mm was selected on GFP-PGC1 nuclear condensates and bleached with Argon laser at 488 nm wavelength by 10 consecutive bleaching iterations (1ms). Fluorescence intensity was recorded at baseline, during and after the bleaching and corrected by ROI corresponding to the background. After subtraction of background intensity, intensities were normalized in each experiment for global fluorescence (from unbleached droplets). In selected droplets, fluorescence intensity before bleaching was used for normalization (as 1) to compare variation post-bleaching and in different experiments.

Mean bleached fluorescence intensity was obtained with (1) mean fluorescence intensity during bleaching divided by (2) the mean pre-bleaching intensity. Mean recovery of fluorescence intensity was obtained with mean fluorescence intensity post-bleaching (10 seconds post-bleaching) recorded every 2 seconds until 30 seconds post-bleaching. These values were divided by the mean pre-bleaching intensity and mean recovery was calculated by the mean. This process was performed in replicates and compared with 3 different experiments, which were used to calculate the mean and the standard deviations. When indicated, bar plots were created to compare the degree of droplet recovery between conditions.

Classification of material properties in droplets was estimated as previously described (*9*). This follows the degree of fluorescence recovery to infer droplet viscosity. Condensates of similar size were selected for each experiment. Image processing and analysis was performed with ImageJ. Pictures of PGC1A droplets were taken when comparing different cellular interventions.Different media conditions were used as indicated; Dulbecco's with low glucose or Dulbecco´s high glucose medium. When indicated, cells were transfected with siRNA for knockdown experiments, expression plasmid vectors, and chemical treatments. Treatment 1,6 Hexanediol at 2.5mM was used as negative control (240117, sigma aldrich). Low glucose (2 mM), normal glucose (5 m), NaHCO3 (1mM) were used to modify cellular conditions.

##### Extracellular Flux Analysis (Seahorse) Assays

Cells were maintained and treated as described above. When indicated, mitochondrial oxidative phosphorylation from cells was analyzed using extracellular flux analysis (XF24; Seahorse Biosciences) in DMEM buffer (pH 7.4; Sigma Aldrich). Baseline oxygen consumption rates (OCR) were measured every 7 min for control and cells in different biological conditions as previously described.

##### Oil Red O staining

Cells were grown, maintained, differentiated and treated as described above. Cell Quantification of Oil Red O staining in SCAT-MSCs and cells treated with adipocyte differentiation cocktail was performed using the Lipid oil red staining kit (plate reader cat#MAK194) and following manufacturer's instructions. In brief, cells grown in a 24 well plate were fixed with formalin (10%) for 1h, washed twice with water. They were incubated with 60% isopropanol for 5min and incubated with oil red o working solution for 15 minutes. After washing with water 3 times, hematoxylin was added for 1min and washed 3 times with water. Stain was extracted in 250ul isopropanol, which was used to measure oil red O stain in a 96 well plate reader at 492 nm. Absorbance was normalized relative to control cells and fold-change variation between groups was compared.

##### Mitochondria/genomic DNA ratio

DNA was extracted using Isol-RNA Lysis Reagent (5 PRIME) according to the manufacturer's instructions. After DNA quantification, 3 nmol of DNA were used to quantitative real-time PCR using SYBR-green. We used primers for mitochondrial DNA and primers for hypoxanthine-guanine phosphoribosyltransferase (HPRT) as a gene specifically transcribed in the nucleus (available upon request). The ratio between the mitochondrial DNA and genomic DNA gives an indication of the number of mitochondria.

##### Colony-forming fibroblast assays

Adipose tissue MSCs were expanded in culture to 70%–80% confluence, harvested with trypsin-EDTA (GIBCO, Invitrogen, Germany) and counted. Cells were diluted in culture media and plated in 96 wells plate at a cell seeding density of 500 cells. Each biological condition was plated in independent 96 well plates. After incubation for either 3, 7, and 14 days at 37°C in 5% humidified $CO_2$, cells were washed with PBS and stained with 0.5% Crystal Violet (Sigma-Aldrich) for 15 min at room temperature. Cells were washed with PBS and the number of wells with visible colonies were counted from 3 plates (triplicate) per condition. Data was represented for each time point and biological condition in triplicates as the average percent of wells (from 96 wells) with visible colonies.

##### Extracellular potassium measurements

SCAT-MSCs were maintained and treated as indicated before (see mammalian cell cultures). Cellular media was used to determine relative extracellular potassium levels using the potassium fluorometric assay kit (Abcam, ab252904) following manufacturer's instructions.

##### Adoptive cell transfer

Adipose tissue mesenchymal stem cells from 6-12 weeks wild type mice were extracted and maintained as indicated (see mammalian cell culture). These cells are used for ex vivo engineering and subsequently transfer to host mice. AT-MSCs from wild type control and ATP12A-/- mice were isolated and maintained in vitro for 3 days in 12 well plates to cell density of 2x10e5 cell/well. ATP12A-/- cells were transfected with the nucleofector (Lonza) with expressing plasmid vectors for PGC1A, ERRA and MLL4 in combination. 12 hours after transfection media was changed to low glucose media (2mM) for 12h. Protein variation and transfection efficiency were tested using proximity ligation assays as indicated (see PLA assays). These cells were maintained during 6 days where, in the passaging day (every 3rd day), the media was changed to low glucose media (12h). These cells were called memory enhanced (ME) SCAT-MSCs.

Control cells were maintained as indicated (see mammalian cell culture) and were transfected with empty expression vectors. Transcriptional compartment activity was checked in similar experiments in both cell groups. In the last passaging day, cells were dissociated (trypsin, sigma aldrich) from cell plates, counted and used for transfer. A total of 2x10e6 cells/mice were resuspended in 200 ul of PBS and were injected by subcutaneous injections to the inguinal area of host mice. Recipient mice were 2 groups: wild type HFD (8 weeks prior cell transfer) and aged mice (18 months). One week before cell transfer, host mice received one dose of 100 mg/kg of cyclophosphamide to facilitate engrafting as previously described (*10*). Each group of host mice received either control or NE SCAT-MSCs. 4 days after injections, host mice underwent 2 days of intermittent fasting protocol per week (intercalated days) during 2 weeks. After this, animals were sacrificed, and SCAT, SCAT-SVF were collected for downstream molecular analysis.

**Intraperitoneal glucose tolerance test (IP-GTT)**

Intraperitoneal GTT was performed on overnight fasted mice. Blood glucose levels were measured at basal state (0 min) and then at 30, 60, and 120 min after i.p. injection of glucose (1.5 mg/kg body weight). Blood glucose concentrations were measured using the Accu-Chek Aviva monitoring system (Roche, Basel, Switzerland).

**Analysis of protein disordered regions**

Disorder scores from amino acid sequences were obtained in PONDR (stored at http://www.pondr.com/). IDR regions were defined using VSL2 scores.

**Identification of genomic hubs by positional gene enrichments (PGE)**

To identify genomic hubs or regions with positional gene enrichments (PGEs) from the location of genes of interest, we used previously established PGEst methods (*11*, *12*). Briefly, PGEs exploit topological gene locations to determine genomic regions enriched in gene sets. This approach is based on calculating the hypergeometric distribution along genomic distances for gene sets. For a given region, it determines the probability of having observed genes in that region. To test statistical significance, cumulative p-value distributions are used on random simulations or false discovery rate on large gene-sets as previously described (*11*, *12*). The probability of achieving p-value enrichments that are better than chance estimates are used to define genomic hub enrichments.

The additional constraints to categorize a genomic region displaying positional enrichments for a given set of genes are as follows: having >4 genes; that no smaller region was found by random permutations and cumulative p value distributions; that no larger regions with more target genes were found. This algorithm defines genomic regions by the distance between queried genes, followed by estimation of p-value distribution to compare chance expectation. Genomic ranges for positional enrichments were set to 10 million base pairs where PGEs with larger domains were filtered out. To obtain qualitative PGEs we used regions that were enriched within cytogenetic bands (*13*) (curated list archived in http://www.gsea-msigdb.org/gsea/msigdb/index.jsp). We additionally performed permutations with random gene sets (same number of inputs), followed by statistical comparisons using Chi-square test and p-value.

**Intracellular pH measurement**

Cellular models were maintained and treated as indicated (see mammalian cell culture). Cytosolic pH in different conditions were obtained with the intracellular pH indicator kit (pHrodo Red AM, fluorescent, Thermo fisher) following manufacturer’s instructions and with a multimode plate reader (Agilent).

**Intracellular pH fluctuations and Monte Carlo models**

We used dynamic time-dependent pH variations between challenged (i.e. low glucose) and control states to evaluate and model context-dependent fluctuations of cytosolic pH. These dynamic variations in pH were used to build nonlinear statistical models followed by Monte Carlo modeling. In brief, cell culture for dynamic measurements of pH were as follows: cells were plated in 24 well plates in equal cell numbers (1x10e5 cells/well). Cell maintenance and treatments as previously described (see mammalian cell culture). Cells were treated as indicated (e.g., control cells and cells in low glucose during 12h or 36h) after which, we started to estimate pH every 10-15 min, during 60-90 minutes (evaluation window), and where T = time interval so that T0 refers to the beginning of the evaluation period in control or treated cells. Thus, T1, …, Tn corresponds to interval time assessment every 15 mins for at least 4 to 6 intervals.

To avoid variations given by the light indicator during pH measurements, cells were plated and used for evaluation of only one time interval so that every plate tracked 1 different time point. Experimental conditions were performed in replicates and were repeated at least 3 times. Fluorescent readouts from each condition were transformed to relative cytosolic pH levels following manufacturer's instructions. These values were then used to build nonlinear models for comparison across conditions.

Nonlinear sine-wave models: Experimental observations showed an asymmetric oscillatory pattern of pH variation. Their behavior is similar to sinusoidal spectral patterns where fluctuations capture discrepancies at different frequencies. This oscillatory behavior has been observed in complex biological systems where this pattern has been used to evaluate principles for non-stationary variability (*14*). Moreover, asymmetric fluctuations have been observed in different biological systems (*15–17*). To compare this behavior across biological conditions, we used nonlinear sine-wave regression (sine-wave with non zero baseline model in graph pad prism) to model our experimental observations.

We generated models for each condition (across interval measurements) to assess their best-fit parameter. Nonlinear least squares model was used to fit the regression without weighting the data points as previously described (*18*). To discriminate the best-fit parameters between models we used the Akaike's information criteria (AICc), which assumes non-nested inferences. Goodness-of-fit was evaluated by $R^2$ and standard error of the estimate. Parameters obtained from the sine-wave models were as follows: amplitude defines the height of the waves from the baseline; frequency is the number of cycles per time unit; baseline is the Y value where the curve oscillates. We compared the best-fit parameters from biological experiments with randomly generated values that preserved amplitude distributions.

We generated random values per time point followed by nonlinear sine-wave regression, and used AICc to evaluate goodness-of-fit between them. This included basal pH detection in unstimulated conditions. This showed discrepancies between the models, with amplitude of fluctuation displaying the largest variation. We then used these quantified parameters to simulate sinewave behavior constrained by these model variations. We used an in-built simulation option in graph pad where XY table values were selected. Time-series variations were on the X values with similar intervals as our experimental data set (>1000 interval simulations). Y values were generated using sinewave non zero baseline model equations with amplitude, frequency, phaseshift and baseline constants from experimentally derived models. Gaussian random error was used to generate random scatter values. Once the simulations were obtained we performed nonlinear sinewave regression on simulated values to get a better estimation of confidence intervals and estimate best-of-fit and oscillatory parameters. We used these statistical approximations for Monte-Carlo simulation followed by outcome model discrimination.

Monte Carlo modeling of pH fluctuations**:** To estimate the distribution and range of variation among oscillatory parameters in models from different experimental biological observations, we used a Monte-Carlo approach on large-scale simulated data from our previously described nonlinear sine-wave models. From the simulated sine-wave models, we obtained best-fit estimates of models constrained by experimental observations. Using these best-fit estimates, we modeled 500 paralleled models using Monte-Carlo (graph pad prism in-built monte carlo implementation).

We defined sinewave equation parameters for each group followed by nonlinear sinewave regression. Outcome discrimination on each model was carried out with AICc and standard error of the estimate from experimental groups and random generated models. We next tested distribution discrepancies for oscillatory parameters such as amplitude from the Monte-Carlo models. Mean distributions between biological groups from which the models were obtained, were used to make comparisons between model fluctuations and estimate p-value for model assumptions.

**Nonlinear sine-wave and Monte Carlo models from** Intcyt **learning parameters**

We used time-dependent data variations from parameters generated by Intcyt throughout the learning process (see computational framework). These dynamic variations were used to build nonlinear statistical models followed by Monte Carlo modeling. In brief, compartments and stress activity parameters through the learning cycles of Intcyt were used and categorized in 2 groups: pre-clustering (noise and information [1]) and self-organized learning of information (self-organized learning [2]). These values were then used to build nonlinear models for comparison across categories.

Nonlinear sine-wave models: Learning parameters in Intcyt showed an asymmetric oscillatory pattern, with increased amplitude of variation, and similar to asymmetric fluctuations observed in biological systems (*15–17*). To compare this behavior across the two groups, we used nonlinear sine-wave regression (sine-wave with non zero baseline model in graph pad prism) to model these observations. We generated models for each category (across interval measurements) to assess their best-fit parameter.

The non-linear least squares model was then used to fit the regression without weighting the data points as previously described (*18*). To discriminate the best-fit parameters between models we used the Akaike's information criteria (AICc). Goodness-of-fit was evaluated by $R^2$ and standard error of the estimate. Parameters obtained from the sine-wave models were as follows: amplitude defines the height of the waves from the baseline; frequency is the number of cycles per time unit; baseline is the Y value where the curve oscillates. We compared the best-fit parameters from learning parameters representing both categories with randomly generated values that preserved amplitude distributions.

We generated random values per time point followed by nonlinear sine-wave regression, and used AICc to evaluate goodness-of-fit between them. This showed discrepancies between the models, with amplitude of fluctuation displaying the largest variation. We then used these quantified parameters to simulate sinewave behavior constrained by these models. We used an in-built simulation option in graph pad where XY table values were selected. Time-series variations were on the X values with similar intervals as our experimental data set (>1000 interval simulations). Y values were generated using sinewave non zero baseline model equations with amplitude, frequency, phaseshift and baseline constants from experimentally derived models. Gaussian random error was used to generate random scatter values. Once the simulations were obtained we performed nonlinear sinewave regression on simulated values to get a better estimation of confidence intervals and estimate best-of-fit and oscillatory parameters. We used these statistical approximations for Monte-Carlo simulation followed by outcome model discrimination.

Monte Carlo modeling**:** To estimate the distribution and range of variation among oscillatory parameters in models from different experimental observations, we used a Monte-Carlo approach on large-scale simulated data from the nonlinear sine-wave models. From the simulated sine-wave models, we obtained best-fit estimates of models constrained by experimental observations. Using these best-fit estimates, we modeled 500 paralleled models using Monte-Carlo (graph pad prism in-built monte carlo implementation). We defined sinewave equation parameters for each group followed by nonlinear sinewave regression. Outcome discrimination on each model was carried out with AICc and standard error of the estimate from experimental groups and random generated models. We next tested distribution discrepancies for oscillatory parameters such as amplitude from the Monte-Carlo models. Mean distributions between groups from which the models were obtained, were used to make comparisons between models and to estimate p-value for model assumptions.

**Statistical Analysis**

Statistical analyses related to experimental procedures were performed using GraphPad Prism Software (San Diego, CA) and all parameters are indicated in the corresponding figure legend. Quantitative data are presented as the mean ± SEM and n is indicated for each experiment. All experiments were carried out with 3 biological replicates and 3 independent times. Animal experiments were performed in more than 3 animals per group, both female and males were used when indicated. Unpaired Student's t test was used to determine statistical significance when two groups were compared. One-way ANOVA followed by Fisher's least significant difference (LSD) test for post hoc comparisons was used to determine statistical significance when multiple groups were compared. Two-way ANOVA was used to estimate significance between groups constrained by time-based or concentration-based measurements, followed by Tukey test for post-hoc multiple comparisons. Hypergeometric test enrichments and Chi-square tests were used to determine the statistical significance between expected and observed frequencies. Correlations were calculated by Spearman rank and Pearson correlation with a two-tailed test. Statistical significance was defined as p value < 0.05 by either test and is denoted with an asterisk as follows: * p-value 0.05-0.01, ** p-value 0.01-0.001 and *** < 0.001 when indicated.

**Disclosure: use of generative AI**

During the preparation of this work, the authors used the assistance of LLMs such as GPT5, Claude 4.5, Gemini 2.5 pro to edit and improve readability and grammar. All suggestions of text edits were edited, reviewed, validated, and manually corrected by the authors prior to submission of this work. The authors take full responsibility for the originality and content of this work. In addition, during preparation of specific parts, the authors used the same LLM models for code assistance. The authors reviewed and edited the content as needed and take responsibility for the content of the code.

### Supplementary text (S1): mathematical and computational framework

#### S1. Conceptual Foundations Why this Approach?

##### S1.0. The CellDa framework

Cell Dynamic Specialization (CellDa, **Fig. 1B**), is an iterative cross-domain framework that integrates experimental biology, mathematical formalization, computational and machine learning implementation to identify general principles of specialization (**Fig. 1B**). CellDa operates through four integrated cycles or phases:

[1] Biological discovery (Steps 1-5; **Fig. 1B**): This phase characterizes specialization of cell function in cells responsive to environmental changes, mapping transcriptional and phenotypic adaptations to find operational features (nested architecture, dynamic restructuring, input-driven responses, and optimization through environmental oscillation **Fig. 1C**).

[2] Mathematical formalization (Steps 6-8; **Fig. 1B**): This phase extracts and formalizes these operational rules of cellular specialization using a principled approach using operadic algebra and nonequilibrium optimization theory, creating a thermodynamically consistent framework that captures nested composition, structural dynamics and entropy accounting (**Fig. 1D**).

[3] Computational implementation (Steps 9-10; **Fig. 1B**): We implement and evaluate the behavior of these principles in machine learning paradigms, revealing emergent behaviors and modulators of specialization that would be hard to predict from biological observation alone (**Fig. 1E, F**).

[4] Biological validation (Steps 11; **Fig. 1B**): Computational modeling and prediction guide the identification and perturbation of biological analog, in particular molecular modulators that can be targeted for enhanced cell programming and disease therapeutics (**Fig. 1F**).

##### S1.1. Oscillatory Optimisation and the Maxwell's Demon Analogy

Current theories on nonequilibrium within dissipative systems have not been able to account for the heterogeneity observed in biological systems (1-6). Current work advocates the emergence of specialized functions as the unifying ground for understanding out-of-equilibrium systems, including biological ones (4, 6). Specialization of function is associated with compartmentalization and its plasticity (7-11). In living systems, both too little and too much specialization can compromise survival (12, 13). This lets us conceptualize specialization as a temporal state in which living systems are compelled to converge given a particular environment.

On the other hand, absence of specialization could be compared, for example, to a diffuse mixture of particles, all available within the same space. This means that non-specialization can also be seen as a temporal state (14), and that the survival of living systems is influenced by the ability to stay out of equilibrium (15). Interestingly, the oscillation resulting from this nonequilibrium tuning reminds us about metabolic states sustaining cellular processes. To better appreciate this, we can model living systems as compartments containing specific chemical attributes, in which case the game of interactions happening in their dynamic system (16) is similar to Maxwell's demon thought experiment (17) (**Fig. 5A**).

We are therefore inspired by Maxwell's demon thought experiment, which serves as an intuitive paradigm for what it means to be a compartmentalized system that restores as much specialization as possible without the apparent violation of the laws of thermodynamics. In the original thought experiment of Maxwell a demon is used to control a gate between two compartments such that it allows molecules to pass through it according to their velocities. This sorting process goes from high-entropy (disorganized) chambers to the generation of low-entropy (organized) compartments, with an apparent breakdown of the Second Law of Thermodynamics. The resolution of the paradox by Landauer and Bennett showed that information processing itself has thermodynamic cost: the demon has to compute information recalling past measurements, thus, information is erased while creating entropy, returning the Second Law balance.

We used this framework to simulate cellular specialization. Living systems can be modeled as collective assemblies or compartments, containing specific molecular features, where specific dynamic rules create functional relationships that perform Maxwell-demon-like optimization: particular compartments selectively concentrate certain molecules (transcription factors, metabolites, epigenetic modifiers) while rejecting others. However, and contrary to Maxwell's passive demon, cellular compartments are actively ensuring their own specialization by means of internal gradients, chemical affinities, chemical repellent, oscillations, molecular interactions, energy-consuming enzymatic activities, internal passive flows, and even metabolic state fluctuations. One thing is certain, however, there is an inherent ability to oscillate between functional compartmentalized and mixed states, which is a feature of living nonequilibrium systems.

The key insight is that the ability to oscillate between specialized and non-specialized states determines survival in living systems instead of remaining in either extreme. Too much specialization compromise adaptation while too little specialization compromise function. This dynamic equilibrium in which systems are forced to converge towards intermediate specialized states, given certain environmental conditions, gave the conceptual basis for our optimization principle.

**S1.2. Why Transcriptional Adaptations and Metabolic Stress as a Model System**

We decided on transcriptional adaptations linked to metabolic function to define operational rules of cell specialization for five reasons:

(i) A direct readout of specialization: Transcriptional programs predict subsets of gene networks that define phenotypic cell function; they also offer a direct bond between molecular causes and phenotypic outcomes.

(ii) Multi-scale nested architecture: Transcriptional regulation is hierarchical and nested. There are genomic hubs containing functional units of regulation such as enhancers and genes, which are used to recruit transcription regulatory complexes, which form phase-separated condensates, and modulate expression of gene programs linked to cell fate or function transitions. The observation that nested products of compartmentalization in the nucleus occur naturally reproduces operadic formalisms (see mathematical section).

(iii) Integration between static and dynamic aspects of genome organization: Genome organization offers stable/consistent spatial templates (chromatic loops, TADs, functionally enriched genomic hubs) which are highly stable, whereas the transcriptional condensates present an aspect of active self-organization (formation, fusion, disintegration). This combination of structural constraint and dynamic plasticity is ideal for testing and exploring optimization principles.

(iv) Thermodynamic sensitivity: Transcriptional condensates are sensitive to changes in environmental variations including metabolic states (nutrient availability, pH, ionic strength, crowding etc) and are directly associated with functional outputs (gene programs, epigenetic modifications, cell function). This allows us to directly study how external drivers modulate internal structural organization and specialization outcomes.

(v) Experimental tractability: Unlike many nonequilibrium cellular processes, transcriptional compartments can be visualized (live imaging), perturbed (genetic, pharmacological, environmental manipulations), and quantified (ChIP, Hi-C, sequencing, FRAP) with existing technologies, which can allow contextual tests of theoretical predictions.

These features made transcriptional compartments and metabolic specialization a paradigmatic system for generalization of principles to broader contexts, which could be used for a wide range of phenomena, both biological (neural networks (32), immune systems, developmental programmes (36)) and computational (adaptive and continuous learning systems; **Fig. 2-6**).

**S1.3. Nonequilibrium States and Survival**

The lack of specialization can be represented by a diffuse mixture of molecular components in a uniform spatial distribution (high entropic state of thermodynamic equilibrium). On the other hand, specialization is a compartmentalised organization with low local entropy (**Fig. 5A**). Both are temporal states, not stable configurations in active collectives, such as components in a cell.

The first concept to our framework is the recognition that biological systems need to be at far-from-equilibrium to survive. A system at true thermodynamic equilibrium has no concentration gradients, no chemical potential differences and no capacity for work. In short, the system is dead. Living systems are sustained by the continual dissipation of free energy within them in order to retain organizational gradients against the decay over time by entropy.

A self-sustaining process of metabolic states cycling constantly creates temporal windows and different thermodynamic conditions. These conditions can be also associated with specific switches between more-specialized and less-specialized states. Cells are exposed to rhythmic nutrient availability (feeding/fasting) and energetic states (ATP/AMP) and redox environment, which leads to the generation of time-dependent gradients and periods where different organizational states are thermodynamically favored.

Our framework formalizes the way that systems harness these types of oscillations: cell compartments are formed during high energy states, when costs are incurred by investing in organizational work. During low energy states, partial relaxation in organization is allowed while retaining key modifications (**Fig. 3A-B, 6K**). As that happens over and over again, they achieve seeds of durable specialization (**Fig. 6K, fig. S9F-J**).

This oscillating logic resolves two apparently conflicting needs; the need for stability of specialization (in order to maintain function) yet the need for adaptive flexibility (in order to respond to environmental change). Very rigid specialization would be fragile; constant exploration would be inefficient. Oscillation between these extremes, with asymmetric rectification favoring productive organizational states, is an energy efficient solution (**Table 2, Fig. 6E, L**).

#### S2. Mathematical concepts: Why Operadic Algebra?

##### S2.1. The Challenge with Multi-Scale Biological Modelling

It is well-known that biological systems organize on a number of different spatial and temporal scales (**Fig. 1A, 4A-B**). That is why many modeling approaches for biological principles encounter problems when trying to integrate quantitative differences between scales. For example, reductionist approaches are able to represent fine-scales (molecular dynamics, biochemical networks), but are computationally intractable when they scale up. Other methods, including differential equations and agent-based models, describe macroscopic behaviour but lose ground in mechanistic insights. Multi-scale techniques attempt to merge scales through boundary conditions (20-23), but they only perform well in isolated scales, lacking natural composability among them and sometimes incorporating specific models that do not easily compose.

We therefore need a mathematical language that would enable us to (**Fig. 4A-C**) (i) express hierarchical nesting (not added on), (ii) dynamically restructure across scales, (iii) consistently facilitate the exchange of information across scales with structured and well-defined operations (21), and (iv) help us in analysis (decomposing systems) and synthesis (build systems from constituent parts).

##### S2.2. Operadic Algebra: natural language for hierarchical composition.

These abilities are provided by operadic algebra, which comes from algebraic topology and category theory (18, 19) (Fig. 4A–B). An operad is defined as a mathematical object in which the composition of operations is formalized in a way that allows to compose complex objects from simpler ones, while taking into account hierarchical relationships. The benefits of operadic algebra for biological modelling are:

Compositional closure: composition operations in operadic systems produce new and valid operations (**Fig. 4D**). Biologically, this means that combining sub-cellular processes (transcription, translation, metabolic reactions) generates cellular-level behaviors (growth, division, differentiation) in a formal consistent way. This is different to ad-hoc multi-scale models where different mathematical tasks are called at each level, while operadic frameworks have the same logic across scales.

Explicit hierarchy: they offer a natural way to encode parent-child nested relationships by tree structures (**Fig. 4B-C, 5C-D**). As parent operations "contains" child operations as inputs, such that combining operations resembles biological nested organization (nucleus contains chromatin, chromatin contains genes, genes contain regulatory elements). The topology of the tree enforces consistency across levels as changes in one level automatically pass through its hierarchy based on compositional rules.

Factorization as decomposition: operads allow us to build up (composition **Fig. 4D**), and to break apart components (factorisation, **Fig. 4D**). The ability to perform operation bidirectionally enables both forward and inverse modelling of complex relationships (29). In our model then, factorisation processes represent compartment dissolution while composition is compartment formation.

Relational ontology: operadic formalism specifies accurate relationships between scales via maps of composition (Fig. 4D-G). This is a problem that Longo and Montnevril call the "relational ontology" problem (24). It states that it is difficult to formalize the meaning of one organizational level giving rise to another. In our framework, we answer this issue explicitly as high-level behaviors are defined as compositions of low-level operations and, vice versa. While residual functions measure the entropic cost of these compositions (25) (**Fig. 5B, eq. 3.4.5-A**).

**S2.3. Why the use of operads has not permeated biological modeling**

Despite these benefits, operadic approaches are still rare in biological modelling. There are several reasons for this:

Mathematical abstraction: Operads emerged from pure mathematics (homotopy theory, algebraic topology) using notations and concepts completely unfamiliar to computational engineers and biologists. Indeed, our work required translating biological principles and experimental observations to operadic machinery, and the to biologically motivate the interpretation (cells, organelles, compartments instead of morphisms, objects, categories; **Fig. 4A-B, Fig. S5A-E**).

Lack of biological motivation: Previous implementations of operads dealt with physics (quantum field theory, string theory) or computer science (computer semantics, database theory). The peculiarities of biological systems, specifically (nested organisation, dynamic restructuring and thermodynamic coupling) do not suggest by themselves that they are well-suited to operadic formalism.

Computational implementation issues: Abstract mathematical representations do not readily become executable algorithms. Developing new kinds of computational representations (tree data structures and operadic operations; **Fig. 5C-D**) and optimisation methods (minimisation and search residual accounting; Fig. 5B) that obey axioms of operadics formalism is challenging. All these issues while simultaneously remaining computationally tractable, was necessary for our work.

Incomplete theory: Hiroaki Kitano wrote in 2002 that nonequilibrium dissipative systems theory (30) "have not quite taken into consideration the heterogeneity and structured nature of biological systems." This critique has generally been a valid one. An example is the dissipative structure theory of Prigogine (28, 29), that accounts for pattern formation but not hierarchical compartmentalisation. Our work fills, to some extent, this gap by embedding nonequilibrium optimisation in operadic structure (**Fig. 5A-B**).

**S2.4. What Our Framework Recovers**

By integrating operadic algebra with the concepts of nonequilibrium thermodynamics (as explained in the previous section), our formalism automatically recovers features that are characteristic of living systems (**Fig. 4A-C, 8A-B**):

(i) Functional specialisation: Compartments specialise thanks to $U^2$ minimisation (**Fig. 5B-D**), as they discover structures that optimize input-output relationships. (ii) Local-to-global organisation: Actions diffuse from the leaf cells to the root by compositional hierarchy (**Fig. 4G, Fig. S5E**), thus allowing for emergent, system level behaviours from local interactions. (iii) Robustness: Incorporation of tree substructures isolates failure in specific scales, such that compartment failures do not necessarily damage the whole system (**Fig. 5D**). (iv) Plasticity: Adaptation of the structure after changing inputs are achieved by factorisation and fusion processes (**Fig. 4E-F, 5D**). (v) Autonomy: Decentralized control of the entire process is allowed as it is locally optimised with each cell without global coordination. (vi) Emergence: Complex behaviours (specialisation, memory, adaptation) do not have to be explicitly programmed (**Fig. 5E-F, Fig. S6-S7**) but emerge from simple rules (composition, factorisation, $U^2$ minimisation; **Fig. 5E-F**).

These emergent properties do not result from design but appear directly from operational and structural properties coupled with thermodynamic constraints. This suggests that the framework represents fundamental organisational principles, rather than surface level patterns.

**S2.5. Multi-Scale Compartmentalisation vs. Previous Approaches**

Previous mathematical work on biological modelling has concentrated mostly on compartmentalisation at single scales:

Reaction-diffusion pathways (28) describe the formation of spatial patterns; however, the nature of the compartments is modeled as an emergent steady state rather than a functional hierarchy. Compartmental models; Ordinary differential equations (ODE) models of cellular pathways contain compartments as parameters (26) (nuclear vs. cytoplasmic concentrations), but do not model compartment formation/dissolution. Sections of agent-based models can depict compartments at a single scale (single cells in the tissue) but fail with nested hierarchies.

To our knowledge, our framework is the first to (**Fig. 4, Table 3**): (i) model multiple nested scales of compartmentalisation within a single formalism; (ii) consider compartment structure as dynamic and optimizable rather than fixed; (ii) account for thermodynamic costs of compartment formation and information processing; (iv) enable bidirectional analysis to answer questions why compartments acquire a specific distribution and architecture. This positions our work as a foundational contribution toward a rigorous theory of nonequilibrium multi-scale biological organization: a gap identified twenty years ago that is still unfulfilled.

#### S3. Developmental and Evolutionary Dynamics in Intcyt

##### S3.1. Cell-like processes in adaptive machine learning.

Intcyt forms organizational dynamics remarkably similar to biological development (**Fig 5D, fig. S6A-B**). Starting with a single undifferentiated cell, the system evolves complex hierarchical architectures endogenously with the iterative implementation of its operadic operations. This process displays certain properties of cells:

- Compartmentalization: Upon incorporating inputs with different statistical distributions, Intcyt spontaneously compartmentalizes into specialized compartments (organelles) that select progressively input subclasses (**Fig. 5D, fig. S6C**). In MNIST, for example, early learning makes separate compartments that discriminate between curved digits (e.g., 0, 3, 8, 9) and angular digits (e.g., 1, 4, 7); later refinement separates these into digit-specific sub compartments (**fig. S6C**).
- Fission: When heterogeneous information is in an organelle (high cytosolic content), then the system factorises the information into multiple daughter organelles with more homogeneous components (**Fig 4E**). This process is similar to the organelle fission event in cells (mitochondrial fission, endosome budding) or when compartments are divided by fission to maintain optimal size and composition.
- Fusion: Organelles with redundant or very similar information fuse by composition operations (Fig. 4F), decreasing the complexity of the tree without decreasing the function. In many ways, this is similar to organelle fusion processes (mitochondria fusion for metabolic efficiency; endosome-lysosome fusion for degradation) in which coordination of function is achieved by fusing compartments.
- Division and proliferation: Further proliferation and division of the root cell give rise to new tree branches (**Fig. 5D**, learning cycle 5000). This results in different branches of independent lineages that specialize for specific input categories, and is similar to evolutionary or developmental dynamics of distinct lineages or daughter cells.

Content exchange: exchange of information between parents and children involves a form of intercellular signalling or compartment communication. Child cells report their content to parents cells, while parent cells distribute their cytosolic content to child cells. This two-way flow keeps homeostasis across the hierarchy.

##### S3.2. Plasticity as a learning principle in Neural Networks

##### S3.2. Plasticity as a learning principle in Neural Networks

Unlike artificial neural networks (ANNs) that maintain constant architectures throughout training (number of layers, neurons per layer, patterns of connection), Intcyt constantly adjusts its architecture to the data (**Fig. 5D**), showing that this architectural plasticity takes place in three different aspects:

(i) Depth adaptation: tree depth increases when existing compartments are not able to effectively partition input space. While the exact number of layers in the system is not a parameter to be explicitly defined as they are given by the factorization, creating hierarchical layers automatically (**Fig. 5D**). (ii) Width adaptation: The number of sibling organelles in each level increases or decreases by fission/fusion operations according to the diversity of inputs (Fig. 5D, fig. S6A-B). Heterogenous data initiate more compartments and broader architectures while low variety data consolidate. (iii) Topology adaptation: The tree structure itself changes, it branches and merges depending on the optimisation (**Fig. 5D, fig. S6D**). This enables continuous learning in which new information creates new branches without the need to replace old knowledge.

Performance implications: Structural plasticity offers multiple benefits compared to static architectures (**fig. S6D**): (i) Efficiency: Only storage for functionally relevant compartments are retained and updated. Redundant structure is pruned so there is fewer computational cost as compared to an over-parameterised static network. (ii) Continual learning: New categories of input cause new branches to form without catastrophic forgetting (**Fig. 5D**). Old knowledge is retained in established compartments, new knowledge creates new compartments. This architectural isolation reduces the interference of tasks with each other. (iii) Interpretability: The evolved tree structure offers a natural taxonomy of concepts learned (**fig. S6C**). Related inputs are clustered in branches near each other; however, hierarchical relations (subordinate/superordinate concepts) translate into parent-child relations in the tree. (iv) Robustness: noise in one part of the tree will not affect the entire system. Tree topology isolates failed states, which promote graceful degradation instead of direct pruning.

**S3.3. Comparison to Static Architectures**

Current deep learning architectures (ResNets, Transformers, CNNs) have great performance, but struggle with adaptation as they rely on rigid architectures. Some of the limitations they exhibit are:

Depth parameters need to be specified before training. The number of layers for machine learning algorithms need to be pre-determined by engineers (33). This leads to some caveats such as (i) using few layers simply does not have enough representational power while (ii) too many layers leads to problems in optimization (vanishing gradients (35)) and overfitting. The optimal depth needs to be determined through trial-and-error or hyper-parameter search.

Fixed connectivity: Connection patterns (fully connectivity, convolutional, attention) are manually designed and stay fixed during training. The weights are adaptive, but the internal blueprint is not, which limits performance when distribution shifts.

Catastrophic forgetting: To train on new tasks requires re-training from scratch or the use of advanced methods like elastic weight consolidation or progressive neural networks to avoid overwriting old knowledge.

Hyperparameter sensitivity: Performance depends on the architectural decisions beforehand (number of layers, number of neurons, size of kernels), which require extensive tuning. Optimal configurations are data set specific, and are determined through costly hyperparameter search

The solution to these limitations has been proposed to be dynamic structural plasticity (**Fig. 5D and Table 3**), which is what Intcyt does, by determining its own width and connectivity structure based on operadic optimisation. No need for hyperparameter search as structure is data-driven. New learning generates new structure for endless adaptation without forgetting. This results in an algorithm that discovers architecture instead of being a fixed learning algorithm.

**S4. Integration with Existing Machine Learning Paradigms**

**S4.1. Compatibility with Convolution Neural Networks (CNNs)**

Although there are fundamental differences between the learning and interpretation of "filters" in CNNs and Intcyt, their memory structures are compatible with each other. For example, standard CNN filters in conventional layers is a small weight matrix for each filter (e.g. 3x3), which is processed over input images by performing component-wise multiplication and summation to detect local features (edges, textures, patterns). While these filters are optimized through back-propagation to reduce classification errors, the learned filters are often not interpretable. Visualization requires special operations such as gradient-ascent or activation maximization, such that the interpretations are essentially heuristic.

Intcyt operadic filters: Each organelle has a content vector that can interact with other inputs by component-wise multiplication (**Fig. 4G Eq. 3.3.2-A**), similar to CNN filters. However, Intcyt filters are different in several ways:

Direct interpretability: The organelle content vector is explicitly encoding input distribution (cluster centroids, category prototypes; **Fig. S6C**). Displaying the content vector of an organelle is sufficient to visualize what that organelle has learned.

Hierarchical organization: Filters for Intcyt are organized in a tree-structured database (**Fig. 5C-D**), in which compartmentalization can be viewed as reflecting functional relationships. Related features are located at nearby organelles, whereas abstract features are located at higher parent cells. This structure results in a built-in hierarchy without having to design layers.

Dynamic filter creation: New filters are created from fission operations as illustrated in **Fig. 4E** when existing filters are not adequate to partition the input space. This is different from CNNs, where the number of filters is set and is not adaptable during the training process.

Local optimization: Either, instead of performing global back-propagation, local U2 minimization is used to optimize each filter (organelle) individually (Fig. 5B). Consequently, the contribution of each filter can be estimated separately, eliminating the credit assignment problem.

**Potential hybrid architectures:** In future work, CNN feature extraction can be combined with the structural organization of Intcyt. CNNs might give low-level feature maps from raw images while Intcyt clusters and organizes them into interpretable hierarchies adaptively. This combination would offer complementary strengths, for instance, CNNs are good at recognizing patterns in pixels, while Intcyt specializes in abstracting structures and continuous learning.

##### S4.2. Tree-Based Database for Learned Features

The hierarchical memory structure of Intcyt is a kind of self-organizing database where learned concepts change according to statistical similarity and functional relationships (see Fig. 5C-D, see Fig. S6C).

Database properties:

Hierarchical indexing: Parent cells are categories of abstractions while child cells contain specific instances related to the abstractions (Fig. 5C - D). For example, the root node of the MNIST dataset has the abstract "curved digits" and its subclasses are "8 vs.0" and so on. This natural taxonomy is generated automatically without the need for labeling data.

Content-addressable memory: The incoming queries are forward propagated to the appropriate and representative compartments (Fig. 4G, Fig. S5D-E). The tree allows efficient search, you don't need to compare with all stored examples, but only relevant branches.

Dynamic schema: In contrast to traditional databases with fixed schemas, Intcyt has a flexible structure that adapts to data complexity (see Fig. 5D). Simple datasets result in shallow narrow trees, while complex ones result in deep trees. The schema is automatically updated as new data is coming in.

Lossless representation: The tree retains fine prolific detail while supporting coarse-grained abstractions (Fig. 5C). High resolution information is encoded in the leaf cells, while low resolution summaries are encoded in ancestral nodes. Then, this shows that multiple abstraction levels are present at the same time.

Tracking provenance: Each cell has a record of residual value (Fig. 5D, red shading; Fig. S6D), which represents the relative amount of organizational effort for that compartment. High-residual cells represent well-established knowledge, while low-residual cells represent recent learning. This metadata supports the quantification of uncertainty and to guide active learning strategies.

##### S4.3. Intcyt machine interaction and interpretability

The transparent memory organization of Intcyt offers new forms of interaction as follows:

Exploring learned concepts: The users are able to browse the tree to understand the internal representations of the system (**Fig. S6C**). Analyses of the content of the organelles provide acquired prototypes and analyses of the tree topology yields conceptual relationships. The result of this direct interpretability is that there is no need for post hoc explanation methods.

Correction of errors: If the system makes an error in classification of an input, users can define which organelle was responsible, manually modify its content or require the re-distribution to another branch. This human in the loop refinement is not possible with opaque neural networks.

Querying: Rather than performing exhaustive input-output tests, users can query knowledge by asking, for example, whether the system has learned about a specific category, which would help finding the relevant compartments and examining their contents.

Guiding learning: Users can guide the structural adaptation process by creating compartments themselves, initialising organelles with prior knowledge, or constraining fusion/fission operations, making it possible to inject knowledge beyond standard supervised learning. This also offers the possibility of incorporating additional learning policies for intcyt itself or for its usage in hybrid ai architectures.

Dynamic visualization: Real-time visualizations of trees (**Fig. 5D**), residual accumulation (**Fig. S6D)**, and the evolution of adaptive optimization (Fig. S7D) can be used to observe the learning progress and where any problems may be arising (e.g., over-fitting, under-fitting, inefficient exploration).

These capabilities make the computational model  a platform for collaborative and adaptive intelligence– systems that integrate machine learning performance with human semantic knowledge and control.

#### S5. Solving Challenges in Deep Learning

##### S5.1. The Lottery Ticket Hypothesis

Problem: The lottery ticket hypothesis (Frankle & Carbin, 2019 (34)) states that densely initialized neural networks contain sparse sub-networks which, when trained separately from their initialization, can achieve similar performance as the whole network. This indicates that the challenge of identifying "winning tickets", requires exhaustive testing of exponential sub-network configurations or the use of sophisticated iterative pruning strategies.

Why this is important: Finding optimal sub-networks before costly training would significantly save on computational costs. However, predicting which initial configurations will work involves predicting the path of optimization that the network will take when starting from a given initial configuration. In other words, solving the optimization path before running the network.

The solution of our approach relies on emergent sub-network discovery. Intcyt finds efficient sub-networks as emergent phenomena as a result of its structural plasticity (**Fig. 5D, fig. S6A-B**). Instead of searching over possible prunings of a predetermined architecture, Intcyt only grows the structure that is necessary to capture input statistics and input distribution. It shows the following features:

Incremental growth: In Intcyt, compartments (organelles, sub-trees) are only added when they are really needed **(Fig. 5D**) such that they are added only when $U^2$ minimisation is not satisfied by  the current structure. This ensures every part has a functional use so that vestigial connections or relations are eliminated.

Functional pruning: Fusion operations (**Fig. 4F**) fuse together organelles that are redundant (i.e. which contain similar information or produce similar outputs). This pruning is done automatically and avoids magnitude-based heuristics as well as sensitivity analysis.

Dynamic reallocation: In case new inputs show that the current structure is sub-optimal, fission/fusion can reallocate and redistribute capacity (**Fig. 5D**). The system is not fixed to early architectural choices but can be restructured as the learning takes place progressively.

Optimal by construction: Because the structure is obtained by optimisation ($U^2$ minimisation with residual constraints; **Fig. 5B**), the final architecture is inherently efficient. Every compartment exists because it cuts $U^2$ more than its residual cost. Thus, the architecture discovery is carried out by principled optimisation as opposed to random search.

Quantitative comparison: On MNIST, Intcyt produces classification accuracy with emergent architectures having 15-20 adaptive compartments (final tree depth 3-4 levels; **Fig. 5D, fig. S7A-B**). A similar feed forward neural network needs ~100 hidden units in one hidden layer or ~50 units per layer in two hidden layers networks in order to achieve accuracy. Intcyt compartment counts are not based on hyper-parameter tuning, but instead on adaptive data structure.

Generalisation to ANNs: This leads to the idea of developing hybrid approaches, where one can use Intcyt structural learning to determine the best architecture topology, and then train conventional ANN weights in that topology. Operadic structure identifies where the neurons should be while gradient descent focuses on what the weights should have.

##### S5.2. Vanishing and Exploding Gradients

Problem: In deep neural networks that use backpropagation, gradient signals have to flow from output layers to input layers. Because gradients are computed by chain multiplication and local derivatives (gradient at layer n = gradient at layer n+1 local derivative), they can decay or grow exponentially with network depth. Vanishing gradient causes problems with learning in initial layers, whereas exploding gradients destabilise the process in the training by causing the weights to update in divergent ways.

This issue persists even if mitigations techniques are performed. For example, even if initialization improvements and architectural modifications (Xavier, He init, ResNets with skip connections, LSTM gated cells, gradient clipping) make better the learning issues they fail to address the root cause. The global nature of backpropagation ties all layers together, which makes early layers too sensitive to the magnitude of gradients in downstream layers.

The solution offered by Intcyt relies on local to global optimisation. Our computational model works around gradient problems by using fundamental local learning (**Fig. 5B, fig. S6D**) with key features:

Local loss functions: Each cell optimises its own $U^2$ objective, such that each cell measures only the discrepancy between its compartmentalised and non compartmentalised outputs. They do not need a global loss function that must be propagated through the entire tree.

Limited gradient paths: This is due to the fact that $U^2$ is calculated locally (comparing the value of a parent cell vs. its direct children; **Fig. 5B**), which means that gradient paths are short which prevents scaling exponentially with overall tree depth.

Independent learning rates: Each organelle can have its own learning rate which is dependent on local properties (residual levels, recent $U^2$ level changes etc). This fine grained control alleviates the homogeneous learning-rate issue that is known to cause gradient issues in regular networks.

Structural adaptation: This can be considered as an alternative to gradient-based exploration. When there is no possibility to minimize $U^2$ in the current structure (e.g. too small gradient magnitude), the system uses fission/fusion operations (Fig. 4E-F) to explore other structures, which represents a non-gradient-based escape mechanism from local minima.

Comparison to biological learning: Biological neural networks do not use global error backpropagation (39). Instead, learning seems to require local rules of plasticity (Hebbian learning, spike-timing-dependent plasticity) coupled with neuromodulatory commands. Local optimization by Intcyt is very similar to this biological paradigm. Quantitative stability analysis: in MNIST, training gradients do not decay or explode systematically, gradient magnitudes throughout training are fairly stable within depths (distance from root). In contrast, there is 10-100 times as much variation in the magnitude of gradients between early and late layers in the typical feed forward neural network (before applying mitigation techniques).

##### S5.3. The Credit Assignment Problem (CAP)

Problem: The credit assignment problem asks how to assign responsibility/accountability, either positive or negative, to each of the parts of a complex system. In end-to-end trained neural networks with a global loss function, the contribution of individual neurons or layers is not well-defined, and performance is a complicated nonlinear function of the sum of all the weights.

This has several consequences, such as inability to interpret what the individual units have learned, lack of selective capacity to enhance poor components without disturbing the strong ones, challenges in debugging including finding out which parts fail in a case of poor performance, and limited ability to transfer learning between components among models.

Intcyt approaches this issue using local interpretability, as it provides credit assignment in a transparent manner through three mechanisms (**fig. S6C-D**) as shown below:

Local loss functions: $U^2$ minimization-values within each cell (**Fig. 5B**) defines the extent to which the specialization of the cell improves the system performance. High $U^2$ is indicative of essential compartmentalisation; low $U^2$ is indicative of redundancy, which offers the possibility of assessing the contribution of each component for the global structural and learning representation in the tree.

Organelle content vectors are directly interpretable: Vectors in organelles represent cluster centroids, category prototypes, or feature detectors in input space, which is referred to as content interpretability (**fig. S6C**). Examining content allows one to know what each organelle has learnt, allowing semantic credit assignment or the identification of more complex relationships (this organelle recognizes curved shapes).

Residual accounting: Residual value for each cell (**Fig. 5D; fig. S6D**) is a cumulative record of organisational work done to bring about that structure. High residual cells represent stable and well established knowledge for auditable trusting; low residual cells represent recent tentative learning and signal a need for further scrutiny.

Selective improvement: Organelles that have high $U^2$ but suboptimal clustering (i.e. inadequate capacity) are able to undergo fission (**Fig. 4E)** with no effect on other compartments. This localized modulation is not possible in monolithic ANNs where all parameters are dependent on each other (40). In addition, this enables local-to-global adaptation as local tuning will constitutively improve the structural representation in the whole tree as well as abstract feature extraction in higher levels of the tree.

Transfer learning: Well-known compartments (high residual, low $U^2$, stable content) may be frozen and re-used in related projects. For example, the recognition of digits may have low-level feature organelles that are common to the recognition of handwritten letters, but the task-specific organelles are kept separate.

Debugging: In the case of misclassified inputs by Intcyt, the organelle to which the input was routed can be traced. The routing decision (action propagation; **Fig. 4G**) can be then further examined and whether the error is due to mis-routing or due to local deficits in discrimination can be determined.

Comparison with attention mechanisms: Attention mechanisms in Transformer architecture provide some credit assignment to input tokens with high attention weight. However, attention does not occur at the internal representation level, but at the input level. The assignment of credit in Intcyt goes down to the level of internal structure, making every compartment role transparent (**fig. S6C-D**).

**S5.4. Comparison with LSTM and other architectures**

Long Short-term Memory (LSTM) networks address the vanishing gradient problem of sequential data using a gated mechanism for controlling the information flow (37). Although this approach is effective, LSTMs demonstrate that deep-learning problems cannot be solved by only tweaking the architecture. LSTMs reduce but do not completely remove the gradient problems. Long-range dependencies are bypassed on the vanishing gradients through the use of gradient highways, which are provided by memory mechanisms (called forget, input, and output gates). Still, in extremely deep networks, gradients can still vanish over multiple LSTM cells, such that exploding gradients still occur, and have to be explicitly clipped. The global nature of backpropagation still continues and the errors keep propagating through the entire sequence.

Complementary LSTM strengths of Intcyt reside in for instance, LSTMs are very good at picking up temporal dependencies of the sequential data, while Intcyt is very good at structural organisation and hierarchical abstraction. Future hybrid architectures (42) can use LSTMs for sequential feature extraction, outputs for Intcyt for structural organisation and also long term memory. Intcyt's tree structure (**Fig. 5C-D**) could be used to implement episodic memory systems in which sequences are organised according to context (38) such that different branches are allocated to different scenarios.

LSTM, ResNet, Transformer, among others, all consist of advanced engineering solutions to specific deep learning problems (vanishing gradients, long range dependencies, parallel processing). Each solves a specific problem using intelligent architecture but still suffers from fundamental limitations of static structure and global optimisation.Intcyt uses a different learning approach in which instead of engineering ad hoc-solutions to pre-defined problems, it derives structure from first principles (operadic composition with thermodynamic constraints; **Fig. 4,5A-B**) and allows solutions to emerge. This generative approach can be better prepared to encounter unforeseen problems, adaptively adjusting the structure of the system to the problems rather than depending on system design parametrization.

#### S6. Future Directions for Operadic Learning Systems.

##### S6.1. Expansions to Other Machine Learning Paradigms

The operadic framework introduced here has been implemented as a self-organized clustering and generative reconstruction algorithm (**Fig. 5E-F, fig. S7**), which can be generalized to other learning paradigms:

Supervised learning: External labels could exert a direct effect on the decision of promoting fission/fusion or other structural configurations. Rather than only using data-based compartment formation (**Fig. 5D**), label information can help the tuning structure towards discriminative partitions more rapidly. For example, one can enforce different branches for different classes when working on a classification problem or use regression targets to constrain the partition of the continuous feature space.

Reinforcement learning: Actions, states and rewards can be grouped into tree hierarchies. High-level policies (parent cells) break down into low level sequences of action (child cells). Hierarchical reinforcement learning has a natural operadic structure mapping, where temporal abstraction can be represented as compositional hierarchy. This could also be implemented with LLM architecture to improve reasoning traces.

Semi-supervised learning: Initial compartments may be seeded by a few labelled examples, which leads to the subsequent growth via unsupervised fission as more and more unlabelled data come in. This hybrid approach combines the oversight of supervision and the scalability of unsupervised approaches.

Meta-learning: Each task is modelled as an organelle in a meta-learning tree. Similarity and hierarchical relations between tasks organise naturally among the tree structure. Transfer learning arises organically: parent-cell representations are task-shared between related problems, whereas the child-cell representations are task-specific. This could be implemented in multi-agent systems requiring coordination and shared hierarchies (current issues with agentic systems).

Continual learning: Intcyt has inherent support for online learning through structural adaptation (**Fig. 5D**). New data streams are able to cause fission/fusion without re-training from scratch. The main issue is scaling the growing size of trees, which, with a periodic check of the problem and aggressive fusion, may keep the size of the problem tractable.

##### S6.2. Operads as a fundamental Units for Learning Networks

The introduction of the concept of operadic algebra as a building block for machine learning opens both theoretical and practical routes (**Table 3**).

**Theoretical implications**

A (unifying) theory of learning can be tested empirically: operadic structure provides a genuine common language for various paradigms (supervised, unsupervised, reinforcement, meta-learning). Instead of having a theory for each paradigm, the operadic framework is expressed in general principles such as composition (**Fig. 4D**), factorisation (**Fig. 4D**) and optimization under thermodynamic constraints (**Fig. 5A-B**), which specialise to specific paradigms depending on the input structure and optimisation goals.

Sample-complexity bounds are expected: operadic hierarchy is expected to induce inductive biases (compositionality, hierarchized abstraction) that should decrease sample complexity compared to flat models. This is also more efficient in terms of computational needs. Additional work developing new learning bounds or Rademacher complexity for the operadic learners might help formalise this intuition.

Thermodynamic constraints: because of explicit tracking of the information processing cost with residuals (Fig. 5D, fig. S6D; eq. 3.4.5-A), operadic representations allow empirical research of fundamental thermodynamic limitations on learning. We might be able to query the energy needs it takes to learn or represent a concept or whether learning efficiency scales with the temperature (**Table 2**).

**Practical implications**

Modularity and composability: Operadic structure in our framework makes it possible to have a modular design, where independently trained components compose to form larger systems. This allows distributed learning (different teams learn different sub-trees), reuse of well-proven sub-trees from different applications, as well as fast prototyping through component library-based assembly.

Hardware mapping: The natural map of operadic operations in our framework (composition, factorisation, $U^2$ evaluation, etc) enables the co-design and test of hierarchical hardware structures. Neuromorphic circuits, for example, with tree-structured interconnections, distributed memory hierarchies or reconfigurable circuits have the potential to realize operadic learning more efficiently than the standard matrix-based GPU systems.

Formal verification: The mathematical formalization of our operadic framework provides the foundation for future rigor evaluation and verification of learning systems. Other properties that determine whether parts of the tree or subtree always display consistent outputs [0, 1]" or whether some organelles never interact or other always associate can be empirically tested, which is a prerequisite for safety-critical and auditable applications.

**S6.3. Open Issues and Future Research Agenda**
In spite of the progress, however, there are still a number of significant open questions:

Scalability: How easy is it for operadic frameworks/learners to scale to modern architectures/benchmarks (e.g. ImageNet, GPT scale language models). Current demonstrations are using small datasets to evaluate key emergent learning principles. Scaling may require algorithmic novelties (more efficient computation, parallel tree operations, approximate homeostasis) or hybrid paradigms combining operadic structure with neural-network efficiency (as previously discussed).

Theoretical guarantees: can convergence of operadic optimisation be proved? What are tight sample complexity bounds? How do thermodynamic constraints affect learnability in an information-theoretic sense? Rigorous theory would be used to design additional algorithms and establish realistic performance expectations.

Best scheduling of operations: When should fission vs. fusion vs. composition vs. numerical optimisation be performed by the system? The current framework uses heuristic scheduling rules. Principled scheduling based on the optimisation theory or learned meta-policies could help improve efficiency.

Cross-domain transfer: How well does knowledge about operadic structures acquired in one domain generalize to other related domains? If our framework learns visual concepts from images, can the structure learned help to speed up the learning of visual-reasoning tasks? Systematic transfer learning studies will learn some about generality.

Biological correspondence: Is operadic operation implemented generally in biological systems? A study on whether genome organisation, neural circuits, or immune systems have compositional structure analogous to our framework could support or refute the biological relevance of the framework.

Thermodynamic validation: The residual accounting gives lower bounds for entropy production (**eq. 3.4.5- A, Table 1**). The thermodynamic consistency of physical hardware implementing operadic learning would be experimentally verified by direct calorimetric measurements which would also serve to calibrate the LND units to absolute energy costs.

**S6.4. Concluding Perspective**
By introducing operads as a basic building block for machine learning (**Fig. 4, 8B**), this work provides a new dimension for innovation in machine intelligence. The essence of neural networks (perceptrons, layers, backpropagation) have all remained relatively unchanged since the 1980s despite much progress in architectures, optimisation methods, and scale. Operadic frameworks represent a new computational unit: not only as a new way to connect neurons, but as a new ontology for learning systems. For example, compositional hierarchies that represent the efficient optimization, not just of parameters, but of structure, under thermodynamic constraints (**Fig. 5, Table 3**).

This first paper just scratches the surface. This showed that it is feasible (Intcyt is functional; **Fig. 5D-F**), interpretable (the tree structures encode concepts learned; **fig. S6C**) and thermodynamically consistent (residuals are dissipation; **Table 1, Fig. 6, eq. 3.4.5-A**). Future work should address scalability, theoretical foundations, and integration with current techniques in hybrid architectures. Nevertheless, the core insight, that learning can be formalised as operadic composition under nonequilibrium optimisation offers a principled basis for thermodynamically informed artificial intelligence.

The convergence between operadic learning and biological specialization is suggestive that these principles are independent of implementation (**Table 1, 2; Fig. 7M, 8**). Whether systems are made of proteins or transistors, if they are under thermodynamic constraints in their processing of information, they seem to move to an identical organisational logic: hierarchical structure, dynamic adaptation, oscillatory optimisation and explicit accounting for dissipation. This unity of principles on different substrates is suggestive of basic laws of adaptive intelligence, laws we are only just beginning to articulate.

**S6.5. Alternative interpretations and discriminating tests**
We argued in our work that asymmetric oscillations are controllable drivers that can be rectified into lasting organization, having a sign level thermodynamic closure and a greater drive-to-organization efficiency (DOE) than continuous forcing (Tables 1-2). In the next sub-sections, we proceed to analyze alternative explanations and the empirical data against them.

1. Do the asymmetric oscillations described in our work cause or just correlate with specialization outcomes? Alternative: The fluctuations may be a byproduct of cellular stress or cell-cycle state, and organization may occur independently.

Evidence against: (i) Bidirectional control: Oscillation amplification through ATP12A perturbation increases specialization, and attenuation of amplification through vATPase/TMEM175 RNAi or pH buffering, in all molecular readouts, reduces it (Fig. 6 and related supplementary figures). (ii) Temporal specificity: The time interval peak of H3K4me3 deposition and droplet remodelling are similar and overlap maximal pH fluctuation changes; not at six or thirty-six hours despite similar cumulative exposure (Fig. 6J). (iii) Dose–response: Organizational gains commensurately increase under graded variations/perturbations of the amplitude of oscillation (Fig.6). (iv) Computational parity: Periods of enhanced compartment oscillations co-vary with residual increments and consolidation in Intcyt (Fig.6A -D); no residual accumulation or compartment oscillation abolish completely the optimization mechanism. This creates a mechanistic causal association in the algorithm. Combined, timing precision and bidirectional control are cases in favor of causality.

2. Is the effect of ATP12A pH-specific or due to off-target alternatives? Alternative: ATP12A can have an effect on other ionic or transport processes, or promote additional byproducts that impact key cell functions.

Evidence against: (i) pH rescue: The benefits of ATP12A knockout are collapsed by buffering cytosolic pH. (Fig. 6F–I). Extracellular potassium excess (channel via ATP12A) leads to pH imbalances that are rescued by perturbing ATP12A. (ii) Lysosomal axis coherence: Interacts with a canonical proton-pump pathway (vATPase/ TMEM175) in line with a pH-dependent lysosome pathway (Fig. 6H -I). (iii) Biophysical mechanism: modulating pH amplitude changes droplet viscosity in coactivator (FRAP), which affects MLL4 interaction and enzymatic activity dynamics (Fig. 6J). These observations all lead to pH-dependent control and not some other signal off target.

3. Are computational parallels designed or discovered? Alternative: There is a chance that our computational

adaptive ML models mimic biology, simply because it was designed to.

Evidence against: (i) Emergence, not scripting: The amplification of the fluctuation amplitude in the course of SOL and the hierarchical abstractions/reconstruction are all the results of the nonequilibrium $U^2$ minimization/optimization, just as it is for the residual accounting; the oscillation timing and a tree shape are not hard-coded (Fig. 4-6 and computational supplementary figures). (ii) Task generality.: Reconstruction and imputation problems (MNIST, DREAM3) do not contain biological details but exhibit the same oscillation-organization coupling (Fig. 5E-F). (iii) Prospective prediction: It was predicted by the model that specialization would increase with an increase in amplitude amplification. This led us to explore analog drives and modulators, which led to ATP12A. These arguments are against post hoc mimicry.

4. Is the thermodynamic consistency only directional and coincidental (sign-level)? Alternative: Directional concordance can be too weak to claim thermodynamic consistency and relevance.

Evidence against: (i) TUR-style scaling: Scales of specialization and dissipation occur in a way that is consistent with TUR-like bounds, which we show as heuristic and not absolute values (§ Eq. 3.4.5-D and tables 1 and 2). (ii) Quantified efficiency: In both domains, systematic and reproducible gains in oscillatory conditions are demonstrated by DOE (Table 2 and supplementary methods) and not just matched signs. (iii) Common units: Landauer-normalized dissipation (LND) only provides a conservative lower-bound, and substrate-neutral ledger that links organization to information-processing cost proxies (§ 3.4.5,4.5). Although an absolute calorimetry would optimize absolute values, the cumulative constraints exceed sign-only evidence.

5. Are the principles system-specific instead of general? Alternative: The results might be limited to MSC programs and the implementation of Intcyt.

Evidence against: (i) Cross-domain convergence: The overlap across substrates, scales, and tasks (cells ↔ computation) is indicative of common constraints and not idiosyncrasies (Tables 1-2 ). (ii) Ubiquity of oscillators: The ubiquity of oscillators, circadian, cell-cycle, neural, indicate a general rectification plan. We also clearly state this as a testable generalization not as a universal proof; its extrapolation to other specializations is one of the fundamental priorities going forward (Discussion 8.5).

### Theoretical, Mathematical, and Computational Framework

**Box 1: Summary of mathematical framework (for in depth, see Appendix A)**
The goal is to formalize an adaptive specialization framework using a mathematical language, which could represent hierarchical nested structures (compartments within compartments), allow reversible reorganization (structures can form and dissolve), and take into account the costs of information processing (dissipation cost from irreversible changes).

Systems are represented as hierarchical trees of "operadic cells," a mathematical algebraic structure that readily supports nested compartments, and their interactions with each other. Each cell includes a cytosol with a common pool of undifferentiated resources, organelles as specialised compartments performing specific functions, and a residual budget, which keeps track of the rising cost of information processing that is produced by logically irreversible operations. This tree structure allows multiscale nesting and recursive application of the operations at any hierarchical level

**Core operations.** There are two complementary processes of structural evolution and adaptation. The first one is composition (assembly) which combines components into compartments, while the second one is factorisation (disassembly) which breaks down compartments to simpler constituents. Additional processes include fission/fusion operations (multi-level tuning of compartments) and content exchange between compartments and cells to regulate granularity and maintain homeostatic balance.

External drives and system-level action are defined by organelle operations when they are affected by environmental inputs. The output of the system is an aggregate weighted sum by the total compartment tree of the system with signals propagating upward from child to parent nodes to deliver the system level responses.

**Optimization objective ($U^2$).** The objective function $U^2$ is defined as the square difference between the response of the system under its current compartmental configuration and the response when all compartments can become homogenized (i.e. no specialization or mixed). Structural $U^2$ is a measure of its objective function, where a large $U^2$ implies that compartments have some functional benefit, and are therefore important enough to be preserved and refined into the improved structural function. A small $U^2$ implies that compartments are redundant and can be simplified or lost (dissolve). Repeated execution of this policy will move the system towards the configuration that matches the inputs to the system.

**Information processing accounting.** The total cost of information processing splits into two components ($S_{\text{total}} = \Delta S_{\text{comp}} + \Delta S_{\text{res}}$), a delta Scomp which decreases with increasing compartmental organisation (corresponding to local increase in order), and a delta Sres which increases with each logically irreversible operation (i.e. information erasure, discarded content), where every update made irreversibly advances the residual ledger. These costs are stated in Landauer-normalised units (LND; bits-equivalent dissipation), providing a conservative, physically-based, lower bound proxy on energy requirements.

**Learning dynamics.** Each input cycle is a cycle through several steps: (1) evaluation of $U^2$, (2) readjustment of the compartment contents based on changes in $U^2$ using a gradient-like optimization, (3) reorganization through composition/factorisation if changes in $U^2$ are required, (4) restoration of homeostasis through content and cytosolic exchanges, and (5) clear out cytosolic byproduct while augmenting the residual ledger. Over repeated cycles, results show that the fluctuations in the structure increase during the productive learning period, which is similar to the oscillatory dynamics reported in biological cells.

**Key insight.** This operadic nonequilibrium optimisation paradigm brings together structure discovery and rigorous information processing accounting. Specialisation emerges by oscillatory drives temporarily reducing barriers to irreversible structural changes while retaining functional benefits between oscillatory cycles.

**Computational: MNIST and Fashion-MNIST recall benchmark**
MNIST and Fashion-MNIST were tested using the standard train/test splits (10 classes). The inputs were grayscale 28×28 images that were not processed (as raw pixels; no handcrafted feature extraction), as described in the table description.

Intcyt evaluation: Intcyt was trained on the training set using its self-organization procedure to form an adaptive hierarchy of compartments. For all of the ablation experiments (and their matched controls), Intcyt was initialized with K = 10 terminal compartments (leaf compartments). After the training, each terminal compartment (leaf) was assigned a class label using a training set using majority vote between the labels of samples routed to that compartment. At test time, each image was passed through the hierarchy that was learned and was predicted as the label that was given to its terminal compartment. Test recall was calculated on the held out test set as the macro-averaged recall over the 10 classes (expressed as a percentage). Reported Intcyt values are mean ± standard deviation over independent runs (different random seeds and/or presentation orders), with all other hyperparameters fixed.

Residual ablation (residuals removed during learning): To test the role of residual accumulation in intcyt, we performed an ablation by which residual ledger accumulation was disabled throughout the training. Concretely, in the cytosolic cleaning/spontaneous reaction step where residual would normally have been accumulating, the residual variable was forced to stay at zero (avoiding residual bookkeeping/accumulation) without changing anything else (numerical updates and, when enabled, structuring operations). The evaluation protocol (majority voting readout and test set recall computation) was the same as the baseline Intcyt evaluation.

Structural ablation (no structural operations; fixed hierarchy): To test the contribution of structural reorganization to specialization, we performed a structural ablation, in which structural operations were disabled at the time of training, while keeping numerical updates active. Specifically, with K = 10 initial terminal compartments, we disabled fission, fusion and operadic composition events throughout training, causing the compartment structure to be frozen, and only learn about the numerical compartment dynamics. The evaluation protocol (majority voting readout and test set recall computation) was the same as the baseline Intcyt evaluation.

Baselines. Baseline results from k-NN, linear SVM, random forest, and a compact 2-convolution CNN (<100k parameters) were taken as representative from published benchmark values under comparable dataset conditions (standard splits; raw-pixel inputs). The specific sources for each baseline are well established in the field and are provided in the table legend. Yet, because reported baseline results can vary with preprocessing, augmentation, and hyperparameter tuning, these values are intended as contextual benchmarks rather than a strictly controlled head-to-head re-implementation. The code and the documentation for Intcyt is hosted at the following address**: https://github.com/daicelabs/intcyt-v2**

**Supplementary Methods (M): Thermodynamic foundations for the operadic framework**

**M0. Notation bridge (from math to methods)**
Purpose. One-to-one traceability is provided by this bridge between symbols defined in the Mathematical framework (see supplementary appendix A) and quantities computed and reported in this Method section. This ensures accordance between mathematical constructs and the theoretical implementation.

Objects: The notation for cells and super-cells are adopted from the mathematical framework. Individual cells and super-cells are denoted as shown below and according to Definitions 1.2 and 2.1. In addition, as defined in Definitions 2.3 - 2.7, the hierarchical structure is broken down into the categories "leaves" and "junctions."

Individual cells $c \in \mathcal{C}N(n)$, super-cells $\hat{c} \in \mathcal{C}N, q(n)$ (M0.1, cell membership)

The content of each cell is measured by the content vector K(c) (Definition 1.21), which is the allocation of the resources over different dimensions. The fitness-gap $\delta_k(c, d)$ (Definition 1.22) measures how closely a cell d is satisfying the requirements of an organelle k within a cell c. When the pair (c,d) satisfies the condition d(k,c) = 0 then we say that the organelle k "fits" the cell c (see supplementary appendix A; Convention 1.23), which indicates perfect compatibility.

$K(c) \in \mathbb{R}^N$, $\delta_k(c,d) = 0$ (M0.2, d fits organelle k of c)

In order to guarantee mathematical agreement we apply the cleaning operation clean(c) (see supplementary appendix A; Definitions 2.13, 2.16) which imposes non-negative compartment contents and well-defined sum properties (see supplementary appendix A; Definitions 2.14-2.15). This cleaning step is important as it allows physically meaningful states to be maintained. The existence and uniqueness of the solution to the homeostasis problem is established in Theorem 1.42 and the problem is formally defined in Definition 1.40.

The behavior of the system in response to inputs from the environment is expressed by the cell action c · a and the algebra operator U(c,d)(a) (see supplementary appendix A; Definition 1.30 and 1.34), which has closed forms that were given in Proposition 1.36. The algebra operator is a measure of how far the functional difference is between a compartmentalised structure, arranged hierarchically, and a flat, unspecialised one.

$U(c,d)(a);=;(c \circ d)\cdot a;-;c\cdot\big(d_1\cdot a_1,\ldots,d_n\cdot a_n\big)$ (M0.3, algebra operator)

This operator is at the heart of our proposed framework due to the Specialisation Theorem (see supplementary appendix A; Theorem 1.38) which links the minimisation of U to the refinement of the partition of resources within compartments in the form of an entropy ledger decrease, a direct measure of the level of refinement within the system organisation.

For adaptive tuning we use the allostatic differential (see supplementary appendix A; Definition 1.48; formula in Proposition 1.49) that is applied with the help of the γ-weight (see supplementary appendix A; Definitions 2.18-2.19). This mechanism exhibits a gradient-like nature, and leads the system to progress towards configurations that better fit the demands from the environmental inputs.

Evaluation rule. All the thermodynamic proxies such as dissipative features and residuals are calculated after cleaning is done in order to operate on non-negative content vectors that meet our algebraic constraints. This is cited in the main text sections 2.4 (operator U) 2.4.1, 2.4.2 (operator U), and 4 (implementation).

**M1. Compartment and residual (erasure) entropy-like proxies**

Goal. We provide operational measures of two complementary aspects of system organisation, namely that compartment entropy measures the degree of specialisation inside compartments while residual entropy measures the cumulative dissipation cost of irreversible computations. Throughout the process, we keep units explicit in order to allow reporting as proxies in either bits (information-theoretic) or J K-1 (thermodynamic).

**M1.1 Compartment entropy proxies $S_{comp}$**

After the action of the cleaning operation (Definitions 2.13/2.16) the content vector K(c) has non-negative entries with the total content S K(c) >0. We define normalised probabilities *pi(c)=Ki(c)/SK(c)*, which represent the fraction of total resources in dimension i. The compartment entropy is therefore the Shannon entropy of this probability distribution:

$S_{\text{comp}}(c) = -\sum_{i=1}^{N} p_i(c), \log_2 p_i(c)$ (M1.1., Compartment entropy)

This quantity is between zero (perfect specialization, all resources collected in one dimension) and $\log_2 N$ (maximum disorder, equal distribution across all dimensions). Theorem 1.38 ensures that the specialisation is sharper (i.e. lower Scomp) under the minimisation of U (see supplementary appendix A).

Worked mini-example: Consider for example a system with 3 dimensions and content K(c) = (0.6,0.3,0.1) giving directly probabilities p = (0.6,0.3,0.1) because the sum of the probabilities is unity. Then, the compartment entropy is: Scomp = −(0.6$\log_2$0.6 + 0.3$\log_2$0.3 + 0.1$\log_2$0.1) ≈1.295 bits

**M1.2 Residual (erasure) information processing cost $S_{res}$**

Let $\widetilde{R}(c)$ be a cumulative number of bits erased by logically irreversible operations from cell c (detailed accounting in M3-M4); Following Landauer's principle, each bit of information erased must be used up by dissipating a minimum of $k_{\mathrm{B}}T \ln 2 \cdot \Delta H_{\text{erase}}$ of entropy to the environment (The minimum entropy cost for erasing $\Delta H_{\text{erase}}$ bits). This leaves us with the residual entropy (in J K -1) and the minimal dissipated heat (in J):

$$S_{\text{res}}(c) = k_B \ln 2\widetilde{R}(c) \text{ , } Q_{\text{diss}} = T, S_{\text{res}}(c) \text{(M1.2)}$$

In practice, $\widetilde{R}$ is reported in bits, and joule calculations are used when reporting explicitly lower-bound energy statements when being associated with additional physical measurements. Time-series decomposition and phase segmentation are shown in **Supplementary Methods (M) Figure. 1 (A–B)**.

##### M1.3 Total entropy of a tree $S_{total}$

For a hierarchical super-cell tree with leaves nodes Leaf(T), the total entropy is the sum of compartment organisation and irreversible dissipation at all the leaves:

$$S_{\text{total}}(T) = \sum_{c \in \text{Leaf}(T)} \Big( S_{\text{comp}}(c) + S_{\text{res}}(c) \Big) \text{(M1.3)}$$

However, this decomposition keeps separated the entropy linked to internal organization (which may decrease locally) from the entropy generated by irreversible processes (which must increase), which in overall gives a thermodynamic lower-bound accounting of proxy parameters. Where cited in main text section 2.4.5 in Fig. 5D & Fig. 6A-D (residual ledger), and section 5 (closure). Time-series decomposition and phase segmentation are shown in **Supplementary Methods (M) Figure. 1 (A–B)**.

##### M2. Diagnostic sign-level closure

Update cycle. Each computational cycle is carried out in a series of stages: Allostasis -> (Fission/Fusion) -> Composition -> Cleaning. Non-negative states and well-defined summations are enforced by the cleaning function (supplementary appendix A; Definitions 2.14-2.16), while preserving sign constraints and making sure content matches organelle requirements is enforced by the homeostatic solutions (supplementary appendix A; Theorem 1.42; Proposition 1.43). The system tends to be biased towards lower U owing to the allostatic updates and differentials (supplementary appendix A; Proposition 1.49) which, by Theorem 1.38, implies that Scomp tends to decrease (supplementary appendix A)

Using the entropy decomposition of M1.3 we can check at every stage thermodynamic consistency is in place:

$$\Delta S_{\text{total}} = \Delta S_{\text{comp}} + \Delta S_{\text{res}} \geq 0 \text{ (M2.1)}$$

But this inequality must hold for the total change in entropy. Specifically, when the system becomes more organised (compartments specialize) such that the compartment entropy decreases:

$$\Delta S_{\text{comp}} < 0 \Longrightarrow \Delta S_{\text{res}} \geq \left| \Delta S_{\text{comp}} \right| \text{ (M2.2)}$$

That is, for every unit of entropy reduction, by means of structural organisation, there must be at least an equal amount of entropy increase, by means of irreversible operations. Time-series decomposition and phase segmentation are shown in **Supplementary Methods (M) Figure. 1 (A–B)**.

Operational test. In any window, where U decreases, or partitions sharpens (measured in M5), we check for sign-level consistency: $S_{\text{comp}}$ should decrease when the residual/LND (in computational systems) or pH-amplitude (in biological cells) increases. These directional correspondences are summarised on Table 1 (see M8). Where given in the main text as section 2.4.5; Section 5.2; Table 1.

##### M3. How we log logically irreversible steps

Not all computational operations (information processing) have a thermodynamic cost. Specifically, we consider the following to be Landauer-eligible erasure events that enter into the residual counter $\widetilde{R}$.

First, cleaning takes place via cytosolic zeroing then transferring to residual (see supplementary appendix A; Definition 2.13). When the shared cytosolic pool is reset, the information content of the pool is effectively erased, needing then dissipation. Second, the merge-and-discard operations during the fusion of organelles or during factorisation eliminate computational registers from the working system (see supplementary appendix A; Section 1.7). Theorem 1.58 describes the circumstances under which such fusion/fission events can change the measure of U. Third, auxiliary buffer resets (see supplementary appendix A; Definitions 2.18 - 2.19) after allostatic updates which empty transient computational storage.

Each of these events produces an erasure counter. When there is dimensional metadata, we determine the number of erased coordinates as an estimate of bit cost per event. As a conservative default, we allocate one bit for each event when there is no available information on the dimension of the events. Where quoted in main text: Section 2.4.5 (residuals); Section 3.5 (ledger); Methods (LND).

**M4. Lower-bound Landauer-Normalized Dissipation (LND) calibration**
Goal: We are looking to map internal signals of the system to bits per irreversible operation in a way that is in line with our algebraic framework and thermodynamic features, without resorting to absolute Joules claims requiring specific hardware for calibration.

Inputs per cycle *t*: At each time step, we keep track of four key quantities. The stress $S_t$ is a signed adjustment or work signal derived from the allostatic step (see supplementary appendix A; Proposition 1.49; Definitions 2.18-2.19), that is the magnitude of adaptive pressure. The total recorded residual after cleaning and erasure operations is accumulated in a residual store $r_t \geq 0$. The number of compartments $C_t$ represents refers to the number of active compartments in the hierarchy. All of this data is reported in the event log if available. Specifically, the event log gives the following counts: $NZ_t$ , $NM_t$ , $NR_t$ for zeroing, merge, and reset operations respectively.

Example of phase segmentation: We divide simulation runs into two phases, an early exploratory phase, and self-organised learning (SOL) phase. The transition point is detected by a strong threshold on the compartment count $C_t$, where the median absolute deviation (MAD) rule is applied in order not to be sensitive to outliers. This phase difference is very important since the dynamics and dissipation patterns are very different between exploration and refinement regimes.

Operation count refers to the total count of irreversible operations per cycle:

$$E_t = N_Z(t) + N_M(t) + N_R(t) \quad \text{(M4.1)}$$

When detailed event logs are not available, we use a conservative estimate:

$\widehat{E}_t = |c_t - c_{t-1}| + \mathbf{1} r_t - r_{t-1} > 0$, which includes structural changes as well as any remaining increment.

Structural bits lower bound: For each cycle we calculate the minimum number of bits erased by the structural operations:

$$B_{\text{struct},t} = \sum_{e \in Z,M,R} w_e, N_e(t) \quad \text{(M4.2; bits erased per cycle)}$$

where $W_e$ are weights defaulting to unity. When we have dimensional information (i.e. we know that a register contains d bits), we can let $W_e$ equal the actual amount of information that is being erased.

Stress-to-bits slope: In order to obtain a bit equivalent to dissipation measure from the abstract stress signal, we fit a single calibration slope (stress units per bit) over a predefined calibration window:

(M4.3; Slope and signal bits)

$$\alpha = \frac{\sum\limits_{t \in \mathcal{W}} |s_t|}{\sum\limits_{t \in \mathcal{W}} B_{\text{struct},t}},$$

$$B_{\text{signal},\mathcal{A}} = \frac{1}{\alpha}, \sum_{t \in \mathcal{A}} |s_t|$$

This slope can be used to convert accumulated stress during any analysis window A to an equivalent lower-bound bit count *B signal ,A*. Calibration linearity, slope estimation, and phase-resolved LND distributions are shown in **Supplementary Methods (M) Figure. 1 (C-D)**.

LND metrics per phase (P) where P $\in$ {early, SOL}. We now define the Landauer-normalized Dissipation as bits per irreversible operation:

$$\text{LNDstruct}, \mathcal{P} = \frac{\sum t \in \mathcal{P} B_{\text{struct},t}}{\sum\limits_{t \in \mathcal{P}} E_t},$$

$$\text{LNDsignal}, \mathcal{P} = \frac{1}{\alpha}, \frac{\sum t \in \mathcal{P} |s_t|}{\sum\limits_{t \in \mathcal{P}} E_t}$$ , (M4.4; Bit per erasure of operation)

The first measure (LNDstruct) is directly based on counted erasure events, the second measure (LNDsignal) is based on calibrated stress signals. Both give conservative lower bounds on the information processing cost. When appropriate these can be normalised by an effective dimensionality $N_{eff}$ to account for the size of the system.

Sensitivity and uncertainty: We use bootstrap resampling to estimate uncertainty and provide confidence intervals of parametrization. The paired bootstrap tests are used for phase or window comparisons. Sensitivity analysis consists of changing the weights of measured events, $W_e$, detection thresholds, procedures for handling outliers and refits within the SOL phase. Free estimates of event-logs are by necessity more conservative, and tend to overestimate LND (**Supplementary Methods (M) Figure. 1 C-D**). Where mentioned in the main text: Section 2.4.5 (ledger+Landauer-style diagnostics), Section 5.3 (DOE uses LND).

**M5. Diagnostic Drive-to-Organization Efficiency (DOE)**
Concept: The DOE parameter is the slope of organization gain divided by the dissipation cost in matched time periods (windows). This metric exposes a proxy for how much the system transforms irreversible operations into functional structure.

Organization *O:* We first choose a measure of scalar organization and there are three natural choices are taken by the framework: (i) compartmental sharpening by $S_{\text{comp}} \downarrow$, where a decrease compartment entropy indicates strong specialization; (ii) decrease of U-loss, defined as $(-\Delta \|U(c,d)(a)\|_2^2)$, corresponding to decreased structural differences in compartment comparison via optimization; or (iii) organelle clusters, measuring the match between compartment contents and functional requirements (directly related to Theorem 1.38).

Dissipation *D*: For the dissipation axis we use either the actual structural bit count *Bstruct* or the signal based bit count *Bsignal* from M4 which is consistent with the calibration of the LND.

DOE *w* = robust slope of *O* on *D*; (M5.1; DOE slope in window $(w)$), such that we obtained a window-level matched slope as $\text{DOE}_w = \frac{\Delta O}{\Delta D}$

Window-level DOE slopes were calculated using robust regression methods (Theil-Sen estimator; Theil, 1950; Sen, 1968) in order to reduce sensitivity to outliers. For windows with low density of data it is sufficient to use finite-difference approximations. For cross domain visualisation comparing computational and biological systems, we use the z-scored axes to give a dimensionless, standardised DOE (sDOE) for easy comparison. Representative scatter and Theil–Sen fit used to estimate DOE across windows are shown in **Supplementary Methods (M) Figure. 1(E)**. The methodology is mentioned in the main text (see below, 5.3, oscillatory efficiency; Table 2).

**(M5.1) Relative DOE variants**

$$\text{DOE}^{\text{LND}}\text{rel} = \frac{\left(\frac{\Delta\text{Organization}}{\text{LND}}\right)}{\left(\frac{\Delta\text{Organization}}{\text{LND}}\right)\text{baseline}}$$ , (INTCYT; DOE-LND)

$$\mathrm{DOE}^{\mathrm{proxy}}\mathrm{rel} = \frac{\left(\frac{\Delta\mathrm{Organization}}{A\mathrm{pH,rel}}\right)}{\left(\frac{\Delta\mathrm{Organization}}{A_{\mathrm{pH,rel}}}\right)_{\mathrm{baseline}}}$$ , (Cells; DOE-proxy)

**M5.2. Discussion on DOE: organization proxy is discrete and step-like**

Motivation. In this work, Drive-to-Organization Efficiency (DOE) is intended to quantify how effectively irreversible operations (dissipation/drive) are converted into persistent structure (organization/specialization). We define DOE as a marginal efficiency,

$$\mathrm{DOE} \equiv \frac{dO}{dL},$$

where O(t) is a scalar organization proxy and $L(t) = \sum_{\tau \le t} \ell(\tau)$ is a cumulative dissipation proxy (or its calibrated lower-bound proxy such as $B_{\mathrm{struct}}$ or $B_{\mathrm{signal}}$; see M4).

Why a slope estimator can look "modest" for discrete organization proxies. In the computational system, one natural and easily interpretable organization proxy is a discrete structural observable, such as compartment count (or, more generally, a count of discrete reorganizations). In this case, O(t) is piecewise constant and changes only at discrete reorganization events (e.g., fission/fusion/composition). So, even when dissipation L(t) accumulates continuously, the organization data series may exhibit long plateaus with $\Delta O = 0$. Over these intervals, any local derivative $dO/dL$ is naturally near zero, and a windowed slope estimator can appear sparse even when the system is doing work. This regime should be interpreted as maintenance dissipation (cost to preserve and stabilize structure) rather than net organization growth.

This effect is amplified when DOE is estimated via robust typical-slope methods (e.g., Theil–Sen). In a sliding window, the Theil–Sen estimator is the median of pairwise slopes,

$$\widehat{\mathrm{DOE}}_w = \mathrm{median}_{i<j} \left(\frac{O_j - O_i}{L_j - L_i}\right).$$

When O(t) is stepwise, a large fraction of pairs satisfy $O_j = O_i$, producing slopes of 0. If these pairs dominate, the median slope is conservative and can under-represent episodic bursts of organization associated with discrete reorganizations. In other words: when O(t) is stepwise, the instantaneous derivative dO/dL is concentrated at reorganization events; thus a typical-slope statistic (median Theil–Sen) can be small even when gains occur.

Implication for interpretation. A modest or intermittently zero window-slope DOE does not imply the absence of rectification; it can instead reflect that (i) organization gains arrive in bursts, and (ii) the system expends dissipation between these events to maintain and stabilize the already-selected structure. This is consistent with the general view that nonequilibrium systems may dissipate continuously while only occasionally crossing barriers into new structural states.

**M6. IB-style diagnostic proxy for section 2.4.4**

Goal: We confirm that specialization retains predictive information about future inputs while rejecting non-predictive memory–in line with features from the Information Bottleneck framework. The latter pays an unavoidable cost of dissipation, in the form of erasing extraneous state.

Representation *Z*: The representation is built from the specialized action of the current super-cell, c, applied on the input a, which is composed of expected or specialized signals (Definition 2.12) and organelle outputs after allostatic adaptations:

$Z = \mathrm{act}(\hat{c} \mid a)_I$, (M6.1; Representation) where depending on the research granularity, we choose either a single index I, or aggregate group of indices.

Target *Y*: The target constitutes features from subsequent inputs, classification labels or the following specialised signal, whichever the system is trying to predict and/or respond to in order to adapt.

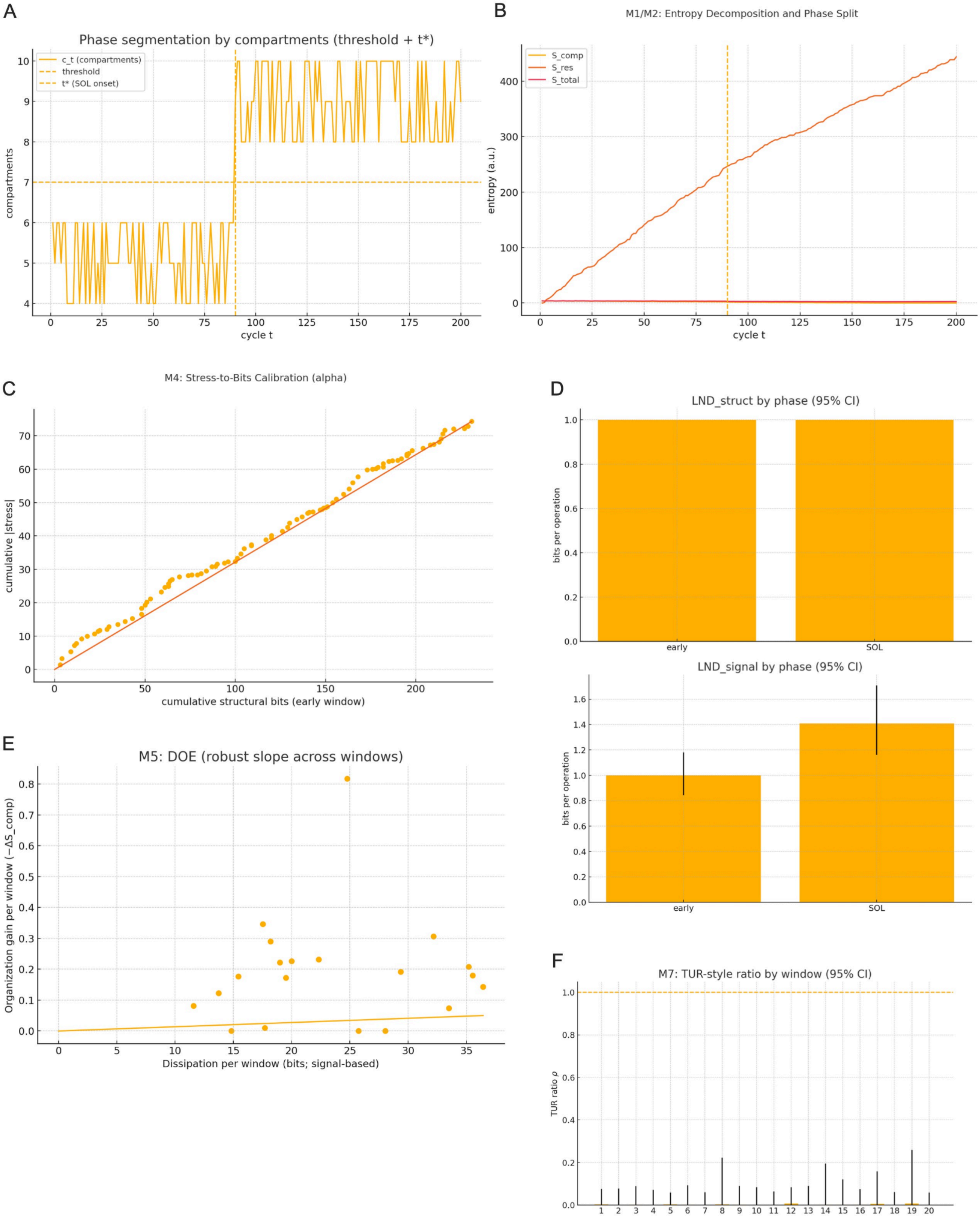
A
Phase segmentation by compartments (threshold + t*)
c_t (compartments)
threshold
t* (SOL onset)
compartments
cycle t
B
M1/M2: Entropy Decomposition and Phase Split
S_comp
S_res
S_total
entropy (a.u.)
cycle t
C
M4: Stress-to-Bits Calibration (alpha)
cumulative |stress|
cumulative structural bits (early window)
D
LND_struct by phase (95% CI)
bits per operation
early
SOL
LND_signal by phase (95% CI)
bits per operation
early
SOL
E
M5: DOE (robust slope across windows)
Organization gain per window (−ΔS_comp)
Dissipation per window (bits; signal-based)
F
M7: TUR-style ratio by window (95% CI)
TUR ratio ρ

**Supplementary Methods Fig. 1. Diagnostics for the operadic framework**
**(A)** Phase segmentation by compartmentalization. The compartment number ct is plotted for 200 learning cycles. A robust threshold (median + 2 x the median absolute deviation (MAD) of early cycles) and its crossing point t* represent the transition between the early and the self-organized learning (SOL) phases. This segmentation marks the beginning of the point at which stable compartmental consolidation takes place, which corresponds to the emergence of specialization dynamics used in downstream analyses (Methods M2).

**(B)** Entropy decomposition and phase split. Cycle-resolved decomposition of total entropy $S_{\text{total}} = S_{\text{comp}} + S_{\text{res}}$ shows that as the structural entropy Scomp decreases residual entropy Sres increases and the total entropy remains not decreasing. The vertical line represents t* from panel A. This confirms the consistency with the generalized second law: organizational gains are compensated by dissipation through irreversible operations (Methods M1-M2).

**(C)** Stress-to-bits calibration ($\alpha$). Within the initial window, the total magnitude of stress $\sum|s_t|$ is linearly related to the cumulative structural erasures $\sum B_{\text{struct},t}$. A through origin linear fit gives the slope $\alpha$, hence the conversion of stress to bits-equivalent dissipation. Near zero intercept and linearity behavior assert that residual increments represent actual erasure events and not random arbitrary variations (Methods M4)
**(D)** Landauer-normalized dissipation (LND) by phase. Top: structural LND (bits per operation) calculated from erasure events that were logged. Bottom: LND obtained from the signal-derived stress signal and calibrated using the $\alpha$-calibrated signal. The early and SOL phases have overlapping 95% bootstrap confidence intervals, which are consistent in the magnitude of dissipation between the different estimation methods. This establishes a reliable dimensionless measure of energy cost in line with Landauer's bound (Methods M4).

**(E)** Drive-to-organization efficiency (DOE). A scatter plot of organization gain per window (-$\Delta S_{\text{comp}}$) vs. bits dissipated per window Dw. This shows a positive Theil-Sen slope that proves that enhanced nonequilibrium drive leads to proportionally greater structural specialization, confirming that oscillatory fluctuations are rectified into net organizational work (Methods M5). Note: In Table 2 we report a phase-level DOE computed from net changes ($\Delta O/\Delta L$), whereas slope-based DOE in sliding windows provides a local diagnostic that can appear conservative when the organization proxy is stepwise (see M5.2).

**(F)** Thermodynamic-uncertainty-relation (TUR-style heuristic) diagnostic. For each window, the ratio

$$\rho = \frac{2\mu_J^2}{\sigma_J^2 \Sigma_w}$$

relates learning precision ($\mu_J\ \sigma_J$) to dissipation $\Sigma_w$. Most $\rho_w$ values are less than or close to 1 as would be expected in the limit of stationary driven processes. Occasional excursions are an indication of windows with transient non-stationarity or sparse erasure counts. Together with DOE and LND analyses, this provides support to a physically consistent learning-dissipation trade-off (Methods M7).
Comment & overall interpretation: Together, panels A-F support the assumption that asymmetric fluctuations work in an oscillatory fashion as a driving resource for efficient specialisation. Consistent with these results, structural order (Scomp decreasing) always correlates with quantifiable dissipation (Sres increasing), and efficiency metrics (DOE, LND, TUR) show that organization arises under conserved thermodynamic constraints instead of arbitrary optimization. These findings are cross-domain evidence that the principles of self-organized computation embrace energy-consistent principles of nonequilibrium adaptation.

---

IB-style quantities (estimators): We estimate two important mutual information quantities. The predicted information $(I(Z;Y))$ of the latent representations Z about the target $Y$ (characterizes to what extent the representation Z can predict *Y*), and estimated by variational lower bounds (Barber-Agakov) or InfoNCE applied on pairs of trajectories:

$I(Z;Y) = H(Y) - H(Y \mid Z)$, (M6.2; Predictive MI (conceptual))

The higher predictive information suggests that more representation is preserving task-relevant structure. The compression I(Z;X) is the measure of the amount of raw input information X retained in Z, which is estimated as Gaussian MI Proxy or decoder log-likelihoods:

$I(Z;X) = H(Z) - H(Z \mid X)$, (M6.3; Compression MI (conceptual))

The non-predictive memory is approximated as the difference between the information gain of Z given X and the information gain of Z given Y (see equation M6.4). During efficient learning, this quantity should follow dissipation measures via diagnostic metrics such as the DOE and thermodynamic uncertainty relations (TUR), as the system loses memory that is not useful or contributes to prediction.

$\Delta\Big(I(Z;X) - I(Z;Y)\Big)$, (M6.4; non-predictive memory (proxy))

Naturally, at a fixed representational complexity, the lower U and the sharpening guaranteed by Theorem 1.38 align with increased I(Z;Y). The cleaning and homeostasis operations limit unnecessary state variables, which allows compression operations to eliminate degrees of freedom which do not contribute to the performance of the function. Where cited in the main text section 2.4.4.

**M7. Learning-dissipation trade-off: TUR-style heuristic (subsection 3.4.5a)**

Purpose: This is meant as a diagnostic tool instead of a statement that the system reaches or saturates thermodynamic bounds. We relate fluctuations of a learning current to the accumulated bits of dissipation and provide insight to the thermodynamic efficiency of the specific learning process.

Observable $(J_t)$: Given a suitable organisation measure, a learning current is defined as $J_t=\Delta O_t$ where O may be an organisation gain such as -$\Delta S_{\mathrm{comp}}$capturing higher specialization or$(-\Delta\|U(c,d)(a)\|_2^2)$ (decreased functional divergence), or may be a matched performance gain such as Δlog-likelihood on a held-out set.

Window $(w)$: Inside each window of analysis, we calculate three statistical parameters:

$\mu_J = \langle J_t \rangle_w, \qquad \sigma_J^2 = \mathrm{Var}w(Jt), \qquad \Sigma_w = B_{\mathrm{signal},w}$, (M7.1; window statistics), where $\mu_J$ is the average learning progress, $\sigma_J^2$ is the variance quantifying fluctuations, and $\Sigma_w$ calculates the dissipation in bits from M4.

TUR ratio (heuristic): The thermodynamic uncertainty relation for driven systems proposes:

$\rho_w = \dfrac{2, \mu_J^2}{\sigma_J^2}, \Sigma_w \lesssim 1$, (M7.2; TUR-style diagnostics)

For near-stationary driven stochastic processes, $\rho_w$ approximation metric should be bounded by unity, or $\rho_w< 1$. Values of $\rho_w>1$ indicate either an under-parameterized dissipation in our accounting or model mismatches (non-stationary dynamics or missing dissipation pathways). We present the value of $\rho_w$ together with confidence intervals and discuss deviations not as hard constraints (or strict violations) but with diagnostic purposes. Per-window values of the TUR-style ratio with 95% bootstrap CIs are presented in **Supplementary Methods (M) Figure. 1(F)**. Where cited in the main text as section 3.4.5a. Concept.

**M8. Panel of cross-domain closure (Table 1)**

The panel shows consistency across substrates, which offers a unified verification for readouts from computational and biological systems. The table shows (1) dissipation proxy, (2) specialization readout, (3) directional relationship; columns represent experimental conditions: control, low-glucose, low-glucose + $NaHCO_3$, vATPase - RNAi, TMEM175 - RNAi, ATP12A - RNAi, IGD (iron - glucose deprivation), ATP12A-/- + IGD. Our computational system (Intcyt) exhibits two example phases: early phase vs. SOL phase. Each cell is annotated with directional arrows (up/down arrow) according to: dissipation proxies measured as pH- amplitude oscillations in biological cells (M9) and LND /residual accumulation in INTCYT simulations (M4); specialization readouts: compartment entropy Scomp, organelle consolidation and cluster purity (via U minimization) and biological markers including OCR, FRAP and H3K4me3 at mitochondrial associated loci.

Specialization is higher when dissipation proxies are running high and lower when proxies are being averaged out, as is typical of generalized second-law accounting (Maxwell closure). This panel shows that the directional predictions in the mathematical framework hold across the computations as well as the biological instantiations.

**M9. Biological proxy methods**

pH-amplitude estimation: Ratiometric cytosolic pH imaging is used in order to capture oscillations of the protons. For each cell trace a quasi-periodic template p(t) is fitted and the peak-to-trough amplitude extracted for every cycle. The amplitude estimates are accompanied by confidence intervals when Monte-Carlo sampling of the fit parameters is performed.

$A_{\mathrm{pH}} = \frac{\mathrm{peak}-\mathrm{trough}}{\mathrm{baseline}}$, (M9.1; amplitude (one cycle) Normalized)

The normalized amplitude is used as a biological proxy for the rate of dissipation following the hypothesis that the larger the oscillation in pH the greater proton-pumping work and, therefore, the greater the energy dissipation.

Organization readouts: Biological specialization is evaluated using a variety of complementary measurements: oxygen consumption rate which is a measure of mitochondrial activity, fluorescence recovery after photobleaching studies (FRAP), which provide recovery time constants that are indicative of organelle mobility and functional compartmentalization, and epigenetic marks (in this case H3K4me3 enrichment at mitochondrial associated promoters in hypoxia associated regulatory regions (mito-HARs)), which report on transcriptional states associated with metabolic reorganization. For DOE and closure plots, all the measurements are standardized to z-scores or are percent change from control conditions.

Perturbations: Experimental manipulations involve the RNA interference or the knockout of specific genes using molecular techniques (e.g. vATPase, TMEM175, ATP12A), pharmacological manipulations, temperature variation, and variable exposure times. Experimental groups, each has n biological replicates providing appropriate statistical power.

Quality control: Strict quality control is enforced: traces showing photobleaching artifacts or baseline drift are rejected; image analysis pipelines are pre registered; and extraction parameters are held constant across conditions so as to avoid batch effects. For purely computational studies, the analogous experimental structure is reflected in systematic ablations: stress signals are eliminated, event logging is turned off, noise schedules are changed or the operation rules of the structures are altered. The impact on measures of LND, DOE and TUR are determined using the same statistical framework. In the main text in Fig. 6 F-K as cited in Section 5-6 (biology), Section 7 (cross-domain).

**M10. Statistics, uncertainty, robustness**

Bootstrap resampling: bootstrap resamples are created for each dataset and bias-corrected and accelerated (BCa) 95% confidence intervals are computed for all derived quantities such as LND, DOE, and mutual-information bounds.

Randomization tests: Statistical significance is measured using randomization procedures: labels of phase (early vs. SOL) are randomized to detect the authenticity of the differences of DOE; indices of time within windows are randomised to obtain sanity tests for TUR calculations by destroying the temporal correlations that are expected to occur within real learning dynamics.

Sensitivity analyses: Robustness is checked by systematically varying methodological choices: window width is varied by +-50%: event weights are tested with bracket uncertainty in dimensional content; outlier trimming is used to eliminate outliers; alternative organization measures (entropy vs. U-loss vs. cluster purity) are compared for consistency. Additionally, the stress -to- bits slope a is recalibrated only inside the SOL phase as a sensitivity check on the calibration procedure.

Multiple comparisons: When comparing multiple hypothesis tests (across DOE windows, MI estimators, TUR checks), the false discovery rate (FDR) is regulated from within each test family in order to ensure that the Type I error rates are appropriate.

Limitations: Several caveats need to be noted. LND gives a lower bound on the bit per operation that is associated with our specific counting rules: for other definitions of irreversibility, different numbers may result. The TUR diagnostic is not an absolute, but rather a heuristic, and is provided mainly as a test of consistency. Mutual-information-bounds are estimator quality-dependent and may be biased in the case of high dimensional representations. Finally, the conversion of bits to absolute Joules has to be done by direct calorimetry or hardware specific calibration, and our Landauer bound calculations are conservative theoretical minima.

#### Supplementary Appendix A: Mathematical and computational framework

The computational framework and documentation for Intcyt is hosted at the following github address:
https://github.com/daicelabs/intcyt-v2

### Part 1. Mathematical Model

1.1. **Compartmentalization.** In this section, we define the concept of a cell as a finite collection of real numbers representing various amounts of chemicals or energetic components. Note that the notations $\mathsf{res}(c)$, $\mathsf{org}(c)$ and $\mathsf{cyt}(c)$ used in the main text will only be introduced in this text from section 1.4. For the sections preceding section 1.4 – including the present one – we will use more explicit notations in order to facilitate the definition of certain operations such as compositions (Definition 1.6) and the statements of Theorem 1.8 and Theorem 1.17. These notations will also be relevant for the discussion of section 3.

**Convention 1.1** (Notation)**.** For every non-negative integer $n$, we will denote by $[n]$ the set of integers ranging from 1 to $n$. If $n$ is zero, then $[n]$ is empty. We will also denote by $\mathbb{R}$ the set of real numbers and by $\mathbb{R}_+$ the set of non-negative real numbers.

**Definition 1.2** (Cells)**.** Let $N$ be a positive integer. A *cell* of *dimension* $N$ consists of

- a non-negative integer $n \geq 0$, called the *number of organelles*;
- a non-negative real number $C$, called the *residual*;
- an element $x$ of $\mathbb{R}^N$, called the *cytosolic content*;
- a $n$-tuple $(x_i)_{i\in[n]}$ of elements in $\mathbb{R}_+^N$, where each $x_i$ is called the *i-th organelles*;

*Remark* 1.3 (Empty cell). By definition, any cell $c = (n, C, x, (x_i)_i)$ for which the natural number $n$ is equal to 0 must be equipped with an empty tuple $(x_i)_{i\in[n]}$ – meaning that the tuple does not contain any vector $x_i$.

**Example 1.4** (A model for cellular structures)**.** The following pictures represent two cells of dimension 3. The leftmost cell has 2 organelles encoded by the vectors $x_1 = (9, 7, 0)$ and $x_2 = (8, 0, 1.2)$ and its cytosolic content is encoded by the vector $x = (.2, 0, 1.4)$. The residual, indicated in brackets, is equal to 0.

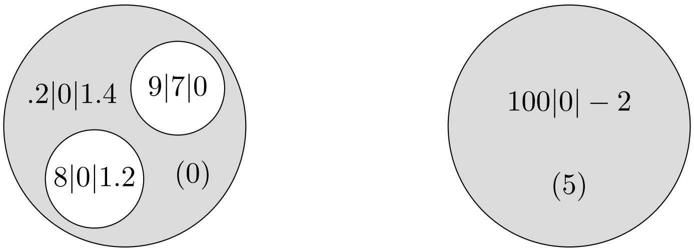

The rightmost cell does not possess any organelle (*i.e.* $n = 0$ and $(x_i)_i$ is empty). However, the cell has a non-trivial cytosolic content encoded by the vector $x = (100, 0, -2)$. The residual is equal to 5.

**Definition 1.5** (The set of cells as a topological space)**.** For convenience, we will denote by $\mathcal{C}^N(n)$ the set of cells of dimension $N$ with $n$ organelles. We deduce from Definition 1.2 that this set is in bijection with the topological space $\mathbb{R}_+ \times \mathbb{R}^N \times \mathbb{R}_+^{N\times n}$.

We now want to be able to nest cells within the organelles of other cells. To this end, we introduce a composition operation modeled on the type of composition used for operads, as shown in the picture below. Our interest in defining such a nesting operation comes from our desire to better conceptualize the sequence of mechanisms that allows cells to find optimal compartmentalization configurations.

As will be seen, we will model compartment formation in terms of a decomposition problem (*i.e.* a factorization problem) whose solutions are characterized in Theorem 1.8.

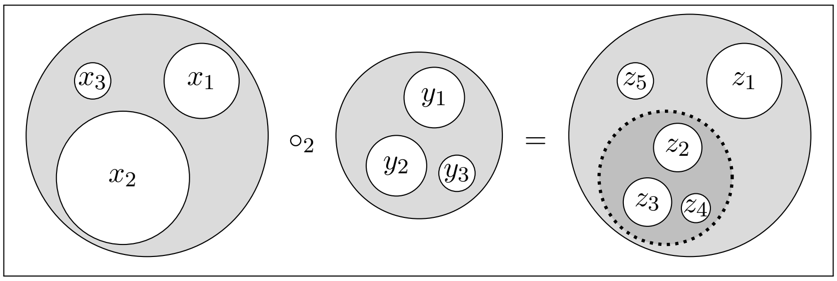

The previous picture, which is often the type of pictures used in operad theory, can also be given a more biological representation. Specifically, the dotted line shown in the previous picture can be seen as a disassembling membrane allowing its content to be mixed with the outer environment.

**Definition 1.6** (Composition)**.** Let $c = (n, C, x, (x_i)_i)$ and $d = (m, D, y, (y_j)_j)$ be two cells of dimension $N$ and let $k \in [n]$. The composition of $c$ with $d$ at organelle $k \in [n]$ is defined by the following cell:

$$c \circ_k d = (n + m - 1, C + D, x + y, x \circ_k y)$$

where $x \circ_k y = (x_1, \ldots, x_{k-1}, y_1, y_2, \ldots, y_m, x_{k+1}, \ldots, x_n)$.

**Example 1.7** (Composition)**.** Equation (1.1) shows an example of a composition of two cells at the second organelle $x_2 = (11, 13)$ of the leftmost cell. The resulting cell, shown on the right-hand side, possesses the two organelles of the middle cell, which replace the second organelle of the leftmost cell, as well as the first organelle $x_1 = (0, 1)$ of the leftmost cell.

(1.1)

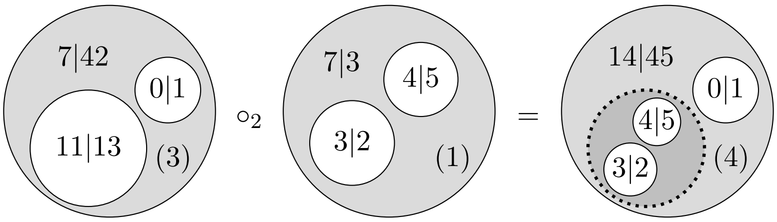

Note that composing cells do not require the cells to satisfy any condition. However, we will later see that the work produced by the cells can be affected by the way the cells are composed between each other. In section 1.4, we will introduce the concept of fitness to distinguish certain types of optimal compositions from others.

Definition 1.6 is simple enough to enable us to solve factorization problems (see Theorem 1.8). Factorizations will be used in section 1.6 to define homeostasis.

**Theorem 1.8** (Factorization problem)**.** *Let $c = (n, C, x, (x_i)_i)$ be a cell of dimension $N$ and let $k \in [n]$. The equation $X \circ_k Y = c$ holds, if and only if, there exist a non-negative integer $\mu$, a non-negative real number $\alpha$ and two vectors $\kappa$ and $\lambda$ in $\mathbb{R}_+^N$ such that the cells $Y$ and $X$ are of the following form:*

$$\begin{aligned} X &= \big(n + 1 - \mu, C - \alpha, x - \kappa, (x_1, \ldots, x_{k-1}, \lambda, x_{k+\mu}, \ldots, x_n)\big) \\ Y &= (\mu, \alpha, \kappa, (x_k, \ldots, x_{k+\mu-1})) \end{aligned}$$

*Proof.* Directly follows from Definition 1.6. □

Below, we introduce the concept of identity cells, which will play an important role in section 2.1 (see Convention 2.10). The importance of these cells lies in their property with respect to compositions.

**Definition 1.9** (Identities)**.** For every positive integer $N$ and every vector $a$ in $\mathbb{R}^N$, we will denote the cell $(1, 0, 0, a)$ of $\mathcal{C}^N(n)$ as $\mathsf{id}_a$.

**Proposition 1.10** (Identities)**.** *For every cell $c = (n, C, x, (x_i)_i)$ be a cell of dimension $N$, the following equations hold for every vector $a$ in $\mathbb{R}^N$:*

$$c \circ_k \mathsf{id}_{x_k} = c \qquad\qquad \mathsf{id}_a \circ_1 c = c$$

*Proof.* By taking the parameters $\mu = 1$, $\alpha = 0$, $\kappa = 0$ and $\lambda = x_k$ in the factorization $X \circ_k Y = c$ of Theorem 1.8, we obtain an equation $c \circ_k \mathsf{id}_{x_k} = c$. Similarly, by taking $\mu = n$, $\alpha = C$, $\kappa = x$ and $\lambda = a$ in the factorization $X \circ_k Y = c$ of Theorem 1.8, we obtain the equation $\mathsf{id}_a \circ_1 c = c$ for any vector $a$ in $\mathbb{R}^N$. □

1.2. **Shuffling organelles.** In the sequel, we will prove results that are invariant up to shuffling the organelles of the cells. As a result, we will only prove the result for one configuration and all the other configuration will follow from the invariance by symmetries. To be able to say when certain results are invariant by symmetry, we define, below, the symmetric action of a bijection on a cell.

**Definition 1.11** (Symmetric action)**.** Let $c = (n, C, x, (x_1, x_2, \ldots, x_n))$ be a cell of dimension $N$. For every bijection $\sigma : [n] \to [n]$, we will denote by $\sigma \odot c$ the cell $(n, C, x, (x_{\sigma(1)}, x_{\sigma(2)}, \ldots, x_{\sigma(n)}))$.

**Proposition 1.12** (Symmetric action)**.** *Let $c \in \mathcal{C}^N(n)$ and $d \in \mathcal{C}^N(m)$. For every index $k \in [n]$ and every bijection $\sigma : [n] \to [n]$, the following identity holds: $(\sigma \odot c) \circ_k d = \sigma \odot (c \circ_{\sigma^{-1}(k)} d)$.*

*Proof.* This is a direct application of Definition 1.11 and Definition 1.6. □

**Convention 1.13** (Symmetric action on vectors)**.** In addition to acting on cells, we shall also need to make bijection act on vectors, as is already the case in Definition 1.11. In this respect, for every bijection $\sigma : [n] \to [n]$ and every vector $v = (v_1, v_2, \ldots, v_n)$, we will denote by $\sigma \odot v$ the vector $(v_{\sigma(1)}, v_{\sigma(2)}, \ldots, v_{\sigma(n)})$.

1.3. **Simultaneous compositions.** So far, our cell compositions have only dealt with one organelle at a time. However, it is possible to define a composition operation that simultaneously composes cells at multiple organelles. A very intuitive way to do so is shown in Definition 1.14.

**Definition 1.14** (Simultaneous composition)**.** Let $c$ be a cell in $\mathcal{C}^N(n)$ and let $d$ denote a $n$-tuple $(d_1, \ldots, d_n)$ of cells $d_k$ in $\mathcal{C}^N(m_k)$. We define the simultaneous composition of $c$ with $d$ as the following cell in $\mathcal{C}^N(m_1 + m_2 + \cdots + m_n)$:

$$c \circ d = ((\ldots(c \circ_n d_n) \circ_{n-1} d_{n-1} \circ_{n-2} \ldots) \circ_1 d_1)$$

**Example 1.15** (Composition)**.** Below, equation (1.2) shows an example of a simultaneous composition of cells. The cell given in the first component of the tuple is composed at the first organelle $x_1 = (0, 1)$ of the lefmost cell and the cell given in the second component of the tuple is composed at the second organelle $x_2 = (11, 13)$ of the leftmost cell. The resulting cell, shown on the right-hand side, possesses all the organelles shown in the tuple.

(1.2) 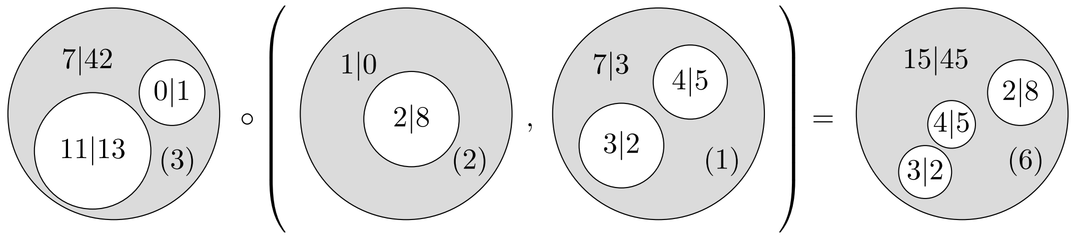

The reader may have noticed the composition of the second cell of the tuple does not affect the composition of the first cell of the tuple, which suggests that that cells could be composed in any order provided that we figure out the index at which the composition occur (see Theorem 1.17)

The following proposition uses the symmetric action on vectors defined in Convention 1.13.

**Proposition 1.16.** *Let $c$ be a cell in $\mathcal{C}^N(n)$ and let $d$ denote a $n$-tuple $(d_1, \dots, d_n)$ of cells $d_k$ in $\mathcal{C}^N(m_k)$. For every bijection $\sigma : [n] \to [n]$, the following identity holds: $(\sigma \odot c) \circ d = \sigma \odot (c \circ (\sigma^{-1} \odot d))$.*

*Proof.* This is a direct application of Definition 1.14 and Proposition 1.16. □

The order of the composition chosen in Definition 1.14 let us wonder whether a different order would provide a different composition. The answer is negative and is shown by Theorem 1.17, which states that the composed cells can be permuted, provided that the index of the composition is shifted accordingly.

**Theorem 1.17** (Compositional properties)**.** *Let $c \in \mathcal{C}^N(n)$, $d \in \mathcal{C}^N(m)$ and $e \in \mathcal{C}^N(p)$ be three cells of dimension $N$ and let $k \in [n]$ and $r \in [m+n-1]$. The following equations hold.*

$$(c \circ_k d) \circ_r e = \begin{cases} (c \circ_r e) \circ_{k+p-1} d & \text{if } r < k \\ c \circ_k (d \circ_{r-k+1} e) & \text{if } k \leq r \leq k+m-1 \\ (c \circ_{r-m+1} e) \circ_k d & \text{if } k+m \leq r \end{cases}$$

*Proof.* Directly follows from Definition 1.6 since the addition of vectors in $\mathbb{R}^N$ is associative and commutative and so does the nesting of tuples. □

In section 1.6, simultaneous compositions will play a important role in defining homeostasis.

**Theorem 1.18** (Factorization problem)**.** *Let $c = (\ell, C, x, (x_i)_i)$ be a cell of dimension $N$. The equation $X \circ (Y_1, \dots, Y_n) = c$ holds, if and only if, for every $i \in [n]$, there exist a non-negative integer $\mu_i$, a non-negative real number $\alpha_i$ and two vectors $\kappa_i$ and $\lambda_i$ in $\mathbb{R}^N_+$ such that the equation $\ell = \sum_{i=1}^n \mu_i$ holds and the cells $Y_1, \dots, Y_n$ and $X$ are as follows:*

$$\begin{aligned} X &= \big(n, C - \textstyle\sum_{i=1}^n \alpha_i, x - \sum_{i=1}^n \kappa_i, (\lambda_i)_i\big) \\ Y_i &= (\mu_i, \alpha_i, \kappa_i, (x_{L(i)}, \dots, x_{L(i)+\mu_i-1})) \end{aligned}$$

*where, for every $i \in [n]$, we let $L(i) := \sum_{k=1}^{i-1} \mu_k$.*

*Proof.* Follows from Definition 1.14 and $n$ applications of Theorem 1.8. □

**Convention 1.19** (Notations)**.** For the same notations as those used in the statement of Proposition 1.18, let us denote the vectors $(\mu_i)_i$, $(\alpha_i)_i$, $(\kappa_i)_i$ and $(\lambda_i)_i$ as $\mu$, $\alpha$, $\kappa$ and $\lambda$, respectively. From now on, we will denote the solutions $Y_1, \dots, Y_n$ and $X$ of the equation $X \circ (Y_1, \dots, Y_n) = c$ as follows:

$$\begin{aligned} \mathsf{left}(c)(\alpha, \kappa, \lambda) &:= X = \big(n, C - \textstyle\sum_{i=1}^n \alpha_i, x - \sum_{i=1}^n \kappa_i, (\lambda_i)_i\big) \\ \mathsf{right}(c)(\mu, \alpha, \kappa)_i &:= Y_i = (\mu_i, \alpha_i, \kappa_i, (x_{L(i)}, \dots, x_{L(i)+\mu_i-1})) \end{aligned}$$

The vector $(Y_1, \dots, Y_n)$ will therefore be denoted as $\mathsf{right}(c)(\mu, \alpha, \kappa)$.

1.4. **Compositional fitness.** We introduce the notion of fitness to distinguish those cells that perfectly compose for the purpose of homeostasis (see section 1.6). The idea is that the organelle at which the composition is done should be equal to the content of the cell by which it is replaced.

**Convention 1.20** (Notation)**.** From now on, for more clarity, we will tend to refer to the structure of a cell $c = (n, C, x, (x_i)_i)$ through standard notations. Specifically, for such a cell $c$, we will denote the quantity $C$ as $\mathsf{res}(c)$, the vector $x$ as $\mathsf{cyt}(c)$ and the tuple $(x_i)_i$ as $\mathsf{org}(c)$ such that the vector $x_i$ will be referred to as $\mathsf{org}(c)_i$ for every $i \in [n]$.

**Definition 1.21** (Content)**.** For every cell $c \in \mathcal{C}^N(n)$, we define the *content of* $c$ as the quantity:

$$K(c) := \begin{cases} \mathsf{cyt}(c) + \sum_{i=1}^n \mathsf{org}(c)_i & \text{if } n > 0 \\ \mathsf{cyt}(c) & \text{if } n = 0 \end{cases}$$

**Definition 1.22** (Fitness)**.** Let $c \in \mathcal{C}^N(n)$ and $d \in \mathcal{C}^N(m)$ be two cells and let $k \in [n]$. We define the *fitness* $\delta_k(c, d)$ of the cell $d$ for the $k$-th organelle of $c$ as the vector of $\mathbb{R}^N$ defined by the following formula.

$$\delta_k(c, d) := \mathsf{org}(c)_k - K(d)$$

**Convention 1.23** (Fitting)**.** A cell $d$ will be said to *fit* the $k$-th organelle of a cell $c$ if the fitness vector $\delta_k(c,d)$ is equal to the vector 0 in $\mathbb{R}^N$.

**Proposition 1.24** (Contents and fitness)**.** *Let $c \in \mathcal{C}^N(n)$ and $d \in \mathcal{C}^N(m)$ be two cells and let $k \in [n]$. The identity $\delta_k(c,d) = K(c) - K(c \circ_k d)$ holds.*

*Proof.* It suffices to verify that the equation of the statement holds. We will do so by simplifying the terms in the difference $K(c \circ_k d) - K(c)$. By definition, we have the following equation.

$$K(c \circ_k d) - K(c) = \mathsf{cyt}(c) + \mathsf{cyt}(d) + \sum_{i \in [n]-\{k\}} \mathsf{org}(c)_i + \sum_{j=1}^{m} \mathsf{org}(d)_j - (\mathsf{cyt}(c) + \sum_{i=1}^{n} \mathsf{org}(c)_i)$$

Simplifying the terms of the right-hand side gives the following series of equations

$$K(c \circ_k d) - K(c) = \mathsf{cyt}(d) + \sum_{j=1}^{m} \mathsf{org}(d)_j - \mathsf{org}(c)_k = K(d) - \mathsf{org}(c)_k = -\delta_k(c,d)$$

The last equality shows that the identity of the statement holds. □

*Remark* 1.25 (Content of simultaneous compositions)*.* In terms of a simultaneous composition, Proposition 1.24 implies that for every cell $c \in \mathcal{C}^N(n)$ and $n$-tuples $d = (d_1, \ldots, d_n)$ of cells of dimension $N$, the following equation holds.

$$K(c \circ d) = K(c) - \sum_{k=1}^{n} \delta_k(c, d_k)$$

1.5. **Specialization theorem.** In this section, we prove our most important theorem (Theorem 1.38), which is explains the learning capacities of the algorithm presented in the main text (see section 2).

**Definition 1.26** (Sum)**.** For any vector $v = (v_1, \ldots, v_N)$ in $\mathbb{R}^N$, we will denote the sum of all the components of $v$ as $\mathcal{S}(v)$ (as shown below).

$$\mathcal{S}(v) := \sum_{u=1}^{N} v_u$$

For the sake of conciseness, for any function $f : U \to \mathbb{R}^N$ sending an element $x \in U$ to a vector $(f_1(x), \ldots, f_N(x))$, we will shorten the composition $\mathcal{S}(f(x)) = \sum_{u=1}^{N} f_u(x)$ as $\mathcal{S}f(x)$.

**Definition 1.27** (Inputs)**.** For every positive natural number $N$ and real number $A$, we will denote by $\Delta_N(A)$ the space defined by the following specification.

$$\{p \in \mathbb{R}_+^N \mid \mathcal{S}(p) \leq A\}$$

This space describes a higher dimensional pyramid in $\mathbb{R}_+^N$.

**Definition 1.28** (Empty and non-empty cells)**.** A cell $c \in \mathcal{C}^N(n)$ will be said to be *empty* if the sum $\mathcal{S}K(c)$ is equal to 0. Conversely, a cell $c \in \mathcal{C}^N(n)$ will be said to be *non-empty* if $\mathcal{S}K(c) \neq 0$.

*Remark* 1.29 (Entropy)*.* At the end of this section, we will interpret the concept of entropy in every non-empty cell $c \in \mathcal{C}^N(n)$ as a level of homogenization of its associated numerical values. To give an example, a cell in which the proportions $\mathsf{cyt}(c)_u/\mathcal{S}K(c)$ and $\mathsf{org}(c)_{i,u}/\mathcal{S}K(c)$ are relatively close to each other for every $u \in [N]$, as illustrated below, on the right, will be associated with a high entropy.

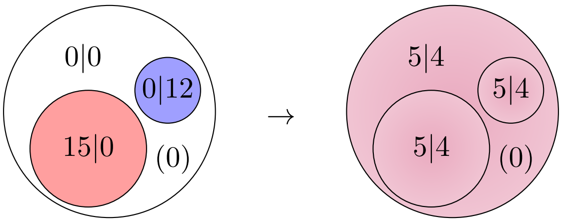

In what follows, we will use the organelles of a cell to reduce its associated entropy. Specifically, we will use the organelles of the cell to either keep specific quantities together or to isolate them from others (with which they should not interact). In Definition 1.30, we associate cells with an action operation that promotes this type of behavior through optimization mechanisms such as gradient descent, cell fusion and cell fission.

**Definition 1.30** (Action of a cell)**.** The *action of a non-empty cell* $c \in \mathcal{C}^N(n)$ on a vector $a = (a_i)_{i\in[n]}$ in $\Delta_N(A)^n$ is defined by the following vector:

$$c \cdot a = \Big( \sum_{i=1}^{n} a_{i,u} \frac{\mathsf{org}(c)_{i,u}}{\mathcal{S}K(c)} \Big)_{u\in[N]}$$

Note that, if the sum $\mathcal{S}\mathsf{cyt}(c)$ is positive, then the quantity $\mathsf{org}(c)_{i,u}$ is always less than or equal to $\mathcal{S}K(c)$(see the formula given in Definition 1.21). As a result, the ratio of $\mathsf{org}(c)_{i,u}$ over $\mathcal{S}K(c)$ can be interpreted as a conditional probability, allowing us to interpret the action of a cell as a vector of conditional expected values. The following proposition formalizes this idea in terms of a stability property.

**Proposition 1.31** (Stability)**.** *For every non-empty cell* $c \in \mathcal{C}^N(n)$ *and vector* $a \in \Delta_N(A)^n$*, the action* $c \cdot a$ *belongs to* $\Delta_N(A - \lambda A)$ *where* $\lambda = \mathcal{S}\mathsf{cyt}(c)/\mathcal{S}K(c)$*.*

*Proof.* The proof follows from the series of relations displayed below.

$$\mathcal{S}(c \cdot a) = \sum_{u=1}^{N}\sum_{i=1}^{n} a_{i,u} \frac{\mathsf{org}(c)_{i,u}}{\mathcal{S}K(c)} \leq A \sum_{u=1}^{N}\sum_{i=1}^{n} \frac{\mathsf{org}(c)_{i,u}}{\mathcal{S}K(c)} = A\frac{\mathcal{S}K(c) - \mathcal{S}\mathsf{cyt}(c)}{\mathcal{S}K(c)}$$

□

*Remark* 1.32 (Spontaneous reactions). For every non-empty cell $c$ of dimension $N$, the quantity $\mathcal{S}\mathsf{cyt}(c)$ may not always be positive. Indeed, the expression $\mathcal{S}\mathsf{cyt}(c)$ can be the sum of both positive and negative values, as illustrated below.

$$\mathcal{S}\mathsf{cyt}(c) = a + b - c + d - e - f + g$$

Our interest in modeling biology motivates the following interpretation: if the quantity $\mathcal{S}\mathsf{cyt}(c)$ is negative, then the cytosolic content of $c$ is missing resources. With such an interpretation, we could ask whether one can compensate this lack by looking for resources elsewhere in the cell. A solution could be the residual $\mathsf{res}(c)$, which we would like to see as an energetic resource.

More specifically, the idea is that, if the inequality $\mathsf{res}(c) + \mathcal{S}\mathsf{cyt}(c) \geq 0$ holds, there are enough positive values both in the residual and the cytosol to compensate the negative values in the cytosol. We could see this compensation as an overall reaction happening in the cytosol, taking elements with positive values to create elements with negative values. This would go as follows:

$$\mathsf{res}(c)\mathsf{Energy} + a\mathsf{A} + b\mathsf{B} + g\mathsf{G} \to c\mathsf{C} + e\mathsf{E} + f\mathsf{F} + \mathcal{S}\mathsf{cyt}(c)\mathsf{Energy}$$

As shown above, the quantities $\mathsf{res}(c)$ and $\mathcal{S}\mathsf{cyt}(c)$ play the role of the energy being used and released by the reaction. In section 2.2, we will use this kind of l ogic to define the 'cleaning' of a cell (Definition 2.13), in which we will i ncrement the variable $\mathsf{res}(c)$ by the value $\mathcal{S}\mathsf{cyt}(c)$ and set the variable $\mathsf{cyt}(c)$ to zero.

In the rest of this section, our goal i s to define an operator that will allow us to control what we could see as the level of entropy of a cell (see Remark 1.29).

**Convention 1.33** (Notations)**.** For every collection $\{m_k\}_{k\in[n]}$ of non-negative integers and every collection $(a_k)_{k\in[n]}$ of vectors $a_k = (a_{k,i})_{i\in[m_k]} \in \Delta_N(A)^{m_k}$, we will denote by $a_1 \quad a_2 \cdots \odot a_n$ the vector concatenation

$$(a_{1,1}, \dots, a_{1,m_1}, a_{2,1}, \dots, a_{2,m_2}, \dots, a_{n,m_n}, \dots, a_{n,m_n})$$

which lives in the space $\Delta_N(A)^L$ where $L = \sum_{k=1}^{n} m_k$

**Definition 1.34** (Algebra operator)**.** Let $c$ and $d_1, \dots, d_n$ be non-empty cells of dimension $N$. For every collection $a = (a_k)_{k\in[n]}$ of vectors $a_k \in \Delta_N(A)^{m_k}$, we will denote by $U(c,d)(a)$ the following difference:

$$(c \circ d) \cdot (a_1 \odot a_2 \odot \cdots \odot a_n) - c \cdot (d_1 \cdot a_1 \odot d_2 \cdot a_2 \odot \cdots \odot d_n \cdot a_n)$$

The quantity $U(c,d)(a)$ given in Definition 1.34 assesses the effect of creating compartments $d_1, \dots, d_n$ within a cell $c \circ d$ on the action operation. In the remainder of the article, we will refer to the function $a \mapsto U(c,d)(a)$ as the *algebra operator* of $c$ and $d$.

*Remark* 1.35 (Invariance by symmetry)*.* Let $c \in \mathcal{C}^N(n)$, $a$ be a vector in $\Delta_N(A)^n$ and $\sigma : [n] \to [n]$ be a bijection. We can easily deduce from Definition 1.30 that the equation $(\sigma \odot c) \cdot a = c \cdot (\sigma^{-1} \odot a)$ holds. It then follows from Proposition 1.16 that the following identity holds.

$$\Big((\sigma \odot c) \circ (\sigma \odot d)\Big) \cdot (\sigma \odot a) = \Big(\sigma \odot (c \circ d)\Big) \cdot (\sigma \odot a) = c \cdot a$$

In the same fashion, we can deduce the following equation, which says that the algebra operator is invariant by symmetry.

$$U(\sigma \odot c, \sigma \odot d)(\sigma \odot a) = U(c,a),$$

This fact will be used in Theorem 1.58, in which we will show that the statement holds for one index configuration and deduce that the other index configurations hold after permutation along bijections $\sigma : [n] \to [n]$.

**Proposition 1.36** (Formula for the algebra operator)**.** *Let $c$ be a non-empty cell in $\mathcal{C}^N(n)$ and $d = (d_1, \dots, d_n)$ be a $n$-tuple of non-empty cells $d_k$ in $\mathcal{C}^N(m_k)$. For every collection $a = (a_k)_{k\in[n]}$ of vectors $a_k \in \Delta_N(A)^{m_k}$, the following identity holds:*

$$U(c,d)(a) = \left(\sum_{k=1}^{n}\Big(\frac{1}{\mathcal{S}K(c \circ d)} - \frac{\mathsf{org}(c)_{k,u}}{\mathcal{S}K(c)\mathcal{S}K(d_k)}\Big)\Big(\sum_{i=1}^{m_k} a_{k,i,u}\mathsf{org}(d_k)_{i,u}\Big)\right)_{u\in[N]} . \tag{1.3}$$

*If, for every $k \in [n]$, the identity $\mathcal{S}\delta_k(c, d_k) = 0$ holds, then we obtain the following expression:*

$$U(c,d)(a) = \left(\sum_{k=1}^{n}\frac{(d_k \cdot a_k)_u}{\mathcal{S}K(c)}\Big(\mathcal{S}\mathsf{org}(c)_k - \mathsf{org}(c)_{k,u}\Big)\right)_{u\in[N]} . \tag{1.4}$$

*Proof.* By Definition 1.14 and Definition 1.30, we have the following formula.

$$(c \circ d) \cdot (a_1 \odot a_2 \odot \cdots \odot a_n) = \left(\sum_{k=1}^{n}\sum_{i=1}^{m_k}\frac{a_{k,i,u}\mathsf{org}(d_k)_{i,u}}{\mathcal{S}K(c \circ d)}\right)_{u\in[N]}$$

Similarly, by applying Definition 1.30 twice, we obtain the following formula.

$$c \cdot (d_1 \cdot a_1 \odot d_2 \cdot a_2 \odot \cdots \odot d_n \cdot a_n) = \left(\sum_{k=1}^{n}\frac{\mathsf{org}(c)_{k,u}}{\mathcal{S}K(c)}\sum_{i=1}^{m_k}\frac{a_{k,i,u}\mathsf{org}(d_k)_{i,u}}{\mathcal{S}K(d_k)}\right)_{u\in[N]}$$

The difference of the two vectors gives us expression (1.3). Factorizing by the inverse of $\mathcal{S}K(d_k)$ in each summand, we obtain the expression

$$U(c,d)(a) = \left(\sum_{k=1}^{n}\Big(\frac{\mathcal{S}K(d_k)}{\mathcal{S}K(c \circ d)} - \frac{\mathsf{org}(c)_{k,u}}{\mathcal{S}K(c)}\Big)(d_k \cdot a_k)_u\right)_{u\in[N]} \tag{1.5}$$

By Remark 1.25 and Definition 1.22, the identity $\mathcal{S}\delta_k(c, d_k) = 0$ holding for every $k \in [n]$ gives us the identities $\mathcal{S}K(c \circ d) = \mathcal{S}K(c)$ and $\mathcal{S}\mathsf{org}(c)_k = \mathcal{S}K(d_k)$. Using these identities in expression (1.5) gives us expression (1.4). ☐

*Remark* 1.37 (Lowerbound for the algebra operator). While the formula given in (1.3) allows the quantity $U(c,d)(a)$ to be either negative, positive or zero, the formula given in (1.4) makes the quantity $U(c,d)(a)$ non-negative. The reason for this lowerbound comes from the fact that the quantity $\mathsf{org}(c)_{k,u}$ is always smaller than the sum $\mathcal{S}\mathsf{org}(c)_k$ of positive terms $\mathsf{org}(c)_{k,1}, \dots, \mathsf{org}(c)_{k,N}$.

Hence, when each cell $d_i$ of the tuple $d$ fits the organelle of $c$ with which it is associated, the algebra operator satisfies the inequality $U(c,d)(a) \geq 0$ for every vector $a$. Theorem 1.38, given below, studies the upperbounds of $U(c,d)(a)$.

**Theorem 1.38** (Specialization theorem). *Let $c$ be a non-empty cell in $\mathcal{C}^N(n)$ and $d = (d_1, \dots, d_n)$ be a $n$-tuple of non-empty cells $d_k$ in $\mathcal{C}^N(m_k)$ for which the identities $\mathcal{S}\delta_k(c, d_k) = 0$ hold. For every collection $a = (a_k)_{k\in[n]}$ of vectors $a_k \in \Delta_N(A)^{m_k}$, the implication*

$$U(c,d)(a)_u \leq \eta \qquad \Rightarrow \qquad \mathcal{S}\mathsf{org}(c)_k - \eta\frac{\mathcal{S}K(c)}{(d_k \cdot a_k)_u} \leq \mathsf{org}(c)_{k,u} \leq \mathcal{S}\mathsf{org}(c)_k \tag{1.6}$$

*holds for every $u \in [N]$ and $k \in [n]$ if $(d_k \cdot a_k)_u \neq 0$.*

*Proof.* The upper bounds comes from the expression $\mathcal{S}\mathsf{org}(c)_k = \mathsf{org}(c)_{k,1} + \dots + \mathsf{org}(c)_{k,N}$, which implies that $\mathcal{S}\mathsf{org}(c)_k$ must be greater than $\mathsf{org}(c)_{k,u}$ for every $u \in [N]$ since the organelles of $c$ are all vectors of non-negative values. The lower bound can be deduced from expression (1.4), given in Proposition 1.36, as follows. First, since the inequality $\mathsf{org}(c)_{k,u} \leq \mathcal{S}\mathsf{org}(c)_k$ holds, we can use the non-negativeness of the summands to deduce the following inequalities.

$$\frac{(d_k \cdot a_k)_u}{\mathcal{S}K(c)}\Big(\mathcal{S}\mathsf{org}(c)_k - \mathsf{org}(c)_{k,u}\Big) \leq \sum_{k=1}^{n} \frac{(d_k \cdot a_k)_u}{\mathcal{S}K(c)}\Big(\mathcal{S}\mathsf{org}(c)_k - \mathsf{org}(c)_{k,u}\Big) = U(c,d)(a)_u \leq \eta$$

Since $(d_k \cdot a_k)_u$ is non-negative and, in fact, by assumption, non-zero, a simple algebra argument between the extreme ends of the previous relation finally provides the lower bound. □

*Remark* 1.39 (Specialization against entropy). Theorem 1.38 says that if one tries to minimize the values of the algebra operator of a non-empty cell $c$ as well as optimize its fitness with respect to a collection of non-empty cells $d = (d_1, \dots, d_n)$ at a dimension $u$ in which the input $a$ sends a strong signal $(d_k \cdot a_k)_u$, then the component of the organelle $\mathsf{org}(c)_k$ in that dimension $u$ tries to converge to the value $\mathcal{S}\mathsf{org}(c)_k$. Interestingly, this can create a 'competition' between the variables $\mathsf{org}(c)_{k,u}$, because the quantity $\mathcal{S}\mathsf{org}(c)_k$ is the sum of all variables $\mathsf{org}(c)_{k,u}$. More specifically, if there existed two variables $\mathsf{org}(c)_{k,u_0}$ and $\mathsf{org}(c)_{k,u_1}$ that were equal to $\mathcal{S}\mathsf{org}(c)_k$, then the following inequality would hold, which is impossible when $\mathcal{S}\mathsf{org}(c)_k \neq 0$.

$$\mathcal{S}\mathsf{org}(c)_k = \mathcal{S}\mathsf{org}(c)_{k,1} + \dots + \mathcal{S}\mathsf{org}(c)_{k,N} \geq 2\mathcal{S}\mathsf{org}(c)_k$$

Hence, the only possible scenario is when there is only one index $u_0$ for which the identity $\mathsf{org}(c)_{k,u_0} = \mathcal{S}\mathsf{org}(c)_k$ holds. In addition, all the other variables $\mathsf{org}(c)_{i,u}$, for which $u \neq u_0$ and $i \in [n]$, must then be equal to zero. In other words, the variable $\mathsf{org}(c)_{k,u_0}$ takes over all the other variables $\mathsf{org}(c)_{i,u}$.

To conclude, reducing the values of the algebra operator promotes specialization in the components $u$ whose associated signals $(d_k \cdot a_k)_u$ are the highest. Interestingly, if the values of the algebra operator can be reduced in more than one dimension, then this specialization process can occur at various components simultaneously.

1.6. **Homeostasis as a back propagation mechanism.** In the preent section, the concept of homeostasis mainly refers to the ability of a system to maintain fitness between its components.

**Definition 1.40** (Homeostasis problem). Let $c \in \mathcal{C}^N(n)$ and let $d = (d_1, \dots, d_n)$ be a tuple of $n$ cells $d_k$ in $\mathcal{C}^N(m_k)$. Below, we denote $m = (m_i)_{i\in[n]}$ and $\mathsf{res}(d) = (\mathsf{res}(d_i))_{i\in[n]}$.

▷ We define a *homeostasis problem* for $(c, d)$ as a collection $\lambda = (\lambda_i)_{i\in[n]}$ of vectors in $\mathbb{R}_+^N$.

▷ We define a *homeostasis solution* for the previous homeostasis problem as a collection $\kappa = (\kappa_i)_{i \in [n]}$ of vectors in $\mathbb{R}^N$ such that, for every $i \in [n]$, the cells

$$c' = \mathsf{left}(c \circ d)(\mathsf{res}(d), \kappa, \lambda)$$
$$d'_i = \mathsf{right}(c \circ d)(m, \mathsf{res}(d), \kappa)_i$$

exist and the equation $\delta_i(c', d'_i) = 0$ holds. Such a solution will be denoted as a triple $(\kappa, c', d')$ where $d'$ denotes the tuple $\mathsf{right}(c \circ d)(m, \mathsf{res}(d), \kappa) = (d'_1, \ldots, d'_n)$.

**Example 1.41** (Homeostasis)**.** At a conceptual level, homeostasis ensures that the fitness requirement imposed by the outer cell to its inner cells holds. If there is underfitting of an inner cell, then the outer cell makes sure to complete what is missing with new resources. If there is overfitting of an inner cell, then the outer cell makes sure to take the generated surplus. Let us consider the triple of cells $(c, d_1, d_2)$ shown below, where $c$ plays the role of the outer cell and $d_1$ and $d_2$ play the role of inner cells nested in the first organelle $x_1 = (0, 1)$ and the second organelle $x_2 = (11, 13)$ of $c$, respectively.

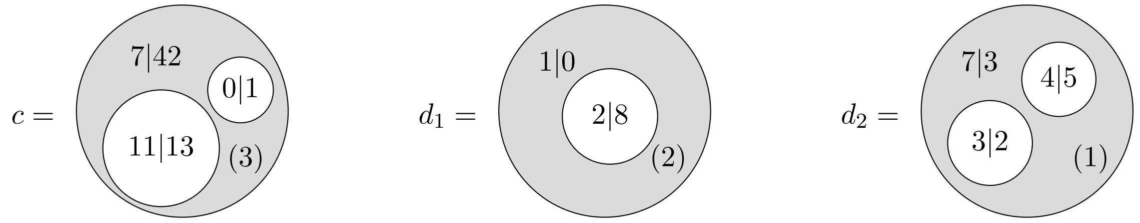

The first organelle of $c$ sets a state requirement of $(0, 1)$ to the inner cell $d_1$. Similarly, the second organelle of $c$ sets a state requirement of $(11, 13)$ to the inner cell $d_2$. The states of the inner cells $d_1$ and $d_2$ are given by their contents $K(d_1) = (3, 8)$ and $K(d_2) = (14, 10)$. Here we see that $d_1$ works too much in both components while $d_2$ works too much in the first component, but not enough in the second component.

Let us now consider the homeostasis problem that consists of the two 2-tuples $\lambda_1 = (0, 1)$ and $\lambda_2 = (11, 13)$. Solving the homeostasis problem for this transformation would amount to determining the state taken by the cell $c$ in order for the inner cell to fit the underlying composition. Intuitively, this means that information is exchanged by the two cells through the inner cell membrane. Letting the information go through the membrane is translated into the composition $c \circ d$ and closing the flux of information is translated into a factorization problem of the form $c' \circ (d'_1, d'_2) = c \circ (d_1, d_2)$. A solution of this problem is given below.

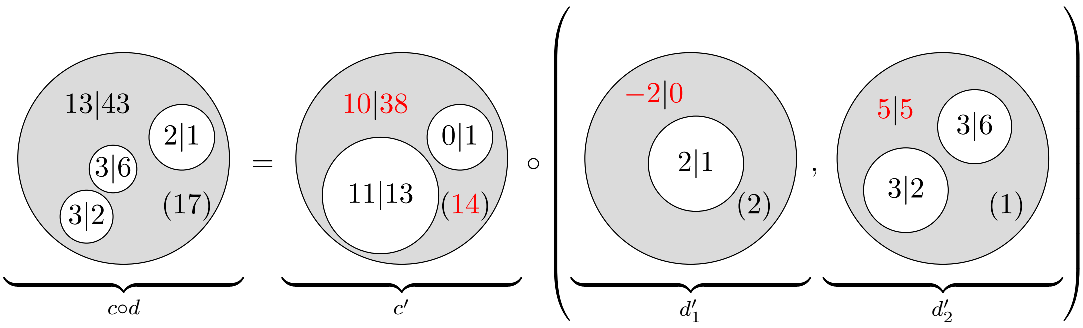

Compared to its previous state $c$, the outer cell $c'$ has taken 1 unit from its second organelle, in its first component, and has given 4 units to its second organelle, in its second component.

**Theorem 1.42** (A unique solution)**.** *The solution $(\kappa, c', d')$ of any homeostasis problem $(c, d, \lambda)$, as described in Definition 1.40, exists, is unique and is determined by the identity*

$$\kappa_i = \lambda_i - K(d_i) + \mathsf{cyt}(d_i)$$

*for every $i \in [n]$. In addition, the equations $\mathsf{res}(c') = \mathsf{res}(c)$ and $c' \circ d' = c \circ d$ hold.*

*Proof.* Let $L(i)$ denote the sum $\sum_{k=1}^{i-1} m_k$. By Definition 1.22, the condition $\delta_i(c', d'_i) = 0$ can be reformulated as follows:

$$\lambda_i = \mathsf{org}(c')_i = K(d'_i) = \kappa_i + \sum_{i=1}^{m_i} \mathsf{org}(c \circ d)_{L(i),i}$$

Since $K(d_i)$ is equal to $\mathsf{cyt}(d_i) + \sum_{i=1}^{m_i} \mathsf{org}(c \circ d)_{L(i),i}$, the previous relation can be derived into the formula given in the statement. In particular, this shows that uniqueness of the solution. To show that the solution exists, we need to check that the parameters defining $c'$ and $d'_i$ that are all well-defined. It turns out that we only need to check that the residual of $c'$ is non-negative. By Theorem 1.18, we have the following equation:

$$\mathsf{res}(c') = \mathsf{res}(c \circ d) - \sum_{i=1}^{n} \mathsf{res}(d_i) = \mathsf{res}(c) + \sum_{i=1}^{n} \mathsf{res}(d_i) - \sum_{i=1}^{n} \mathsf{res}(d_i) = \mathsf{res}(c)$$

Since the residual of $c$ is non-negative, the existence of the solution is proven. Finally, to show that the equation $c' \circ d' = c \circ d$ holds, we can use Theorem 1.18. For this, we need to make sure that the number of organelles of $c \circ d$ is $\sum_{i=1}^{n} m_i$, which directly follows from Definition 1.14. □

In the rest of this section, we study the variations of the algebra operator in terms of a differential over infinitesimal homeostatic exchanges (Definition 1.44). To do so, we will need Proposition 1.43 and Proposition 1.45, which will help us compute useful formulas for these variations.

**Proposition 1.43** (Properties of homeostasis solutions)**.** *Let $(\kappa, c', d')$ be the solution of a homeostasis problem $(c, d, \lambda)$ as described in Definition 1.40. The following equations hold:*

1) $K(c' \circ d') = K(c \circ d)$;
2) $\mathsf{org}(d'_k) = \mathsf{org}(d_k)$, *for every* $k \in [n]$.
3) $K(c') = K(c' \circ d')$;
4) $K(c) = K(c \circ d) + \sum_k \delta_k(c, d_k)$;
5) $K(d'_k) = \mathsf{org}(c')_k$, *for every* $k \in [n]$;
6) $K(d_k) = \mathsf{org}(c)_k - \delta_k(c, d_k)$;

*Proof.* Item 1 comes from the second statement of Theorem 1.42. Item 2 is a direct consequence of Definition 1.19. Item 3 directly follows from the equation $\delta_k(c', d'_k) = 0$, which, by Definition 1.40, holds for every $k \in [n]$. Item 4 follows from the equation given in Remark 1.25 and the fact that $\delta_k(c', d'_k) = 0$ for every $k \in [n]$. Finally, Item 5 is shown in Remark 1.25 and item 6 directly follows from Definition 1.22. □

**Definition 1.44** (Infinitesimal homeostatic solutions)**.** Let $(c, d, \lambda)$ be a homeostasis problem as described in Definition 1.40. For every real number $\varepsilon$, integer $v \in [N]$ and integer $j \in [n]$, a homeostasis solution $(\kappa, c', d')$ will be said to be $(\varepsilon, j, v)$-*infinitesimal* if

- ▷ $\lambda_{j,v} = \mathsf{org}(c)_{j,v} + \varepsilon$;
- ▷ $\lambda_{k,u} = \mathsf{org}(c)_{k,u}$ for every pair $(k, u) \neq (j, v)$ such that $k \in [n]$ and $u \in [N]$;
- ▷ the identity $\mathcal{S}\delta_k(c, d_k) = 0$ holds for every $k \in [n]$.

**Proposition 1.45.** *Let $(\kappa, c', d')$ be an $(\varepsilon, j, v)$-infinitesimal solution for a homeostasis problem $(c, d, \lambda)$ as described in Definition 1.40. The following equations hold:*

1) $\mathcal{S}K(c' \circ d') - \mathcal{S}K(c \circ d) = 0$;
2) $\mathcal{S}K(c') - \mathcal{S}K(c) = 0$;
3) $\mathcal{S}K(d'_j) - \mathcal{S}K(d_j) = \varepsilon$;
4) $\mathcal{S}K(d'_k) - \mathcal{S}K(d_k) = 0$ *for every* $k \in [n]$ *that is not* $j$.

*Proof.* Item 1 follows from the first item of Proposition 1.43. To show item 2, use the third and fourth items of Proposition 1.43 to deduce the equation $\mathcal{S}K(c') - \mathcal{S}K(c) = \sum_k \mathcal{S}\delta_k(c, d_k)$, and use the last item of Definition 1.44 to conclude. To show item 3 and item 4, use the fifth and sixth items of Proposition 1.43 to deduce the equation $\mathcal{S}K(d'_j) - \mathcal{S}K(d_j) = \mathcal{S}\mathsf{org}(c')_j - \mathcal{S}\mathsf{org}(c)_j + \mathcal{S}\delta_k(c, d_k)$ and use the items of Definition 1.44 with the identity $\lambda_j = \mathsf{org}(c')_j$ (Definition 1.40) to conclude. □

**Proposition 1.46** (Effects)**.** *Let $(\kappa, c', d')$ be an $(\varepsilon, j, v)$-infinitesimal solution for a homeostasis problem $(c, d, \lambda)$ where $c$ and $c'$ are non-empty cells in $\mathcal{C}^N(n)$ and, for every $k \in [n]$, the $k$-th components $d_k$ and $d'_k$ of $d$ and $d'$ are non-empty cells in $\mathcal{C}^N(m_k)$. For every collection $a = (a_k)_{k\in[n]}$ of vectors $a_k \in \Delta_N(A)^{m_k}$, the following equations hold.*

$$U(c', d')(a)_u - U(c, d)(a)_u = \frac{\varepsilon}{\mathcal{S}K(c)} \times \begin{cases} (d_j \cdot a_j)_u \dfrac{\mathsf{org}(c)_j}{\mathcal{S}\mathsf{org}(c)_j + \varepsilon} & \text{if } u \neq v \\ -(d_j \cdot a_j)_v \displaystyle\sum_{u=1,\neq v}^{N} \frac{\mathsf{org}(c)_{j,u}}{\mathcal{S}\mathsf{org}(c)_j + \varepsilon} & \text{if } u = v \end{cases}$$

*Proof.* By Definition 1.40 and Definition 1.44, the identities $\mathcal{S}\delta_k(c', d'_k) = 0$ and $\mathcal{S}\delta_k(c, d_k) = 0$ hold for every $k \in [n]$. By Proposition 1.36, this means that we have the following formulas for every $u \in [N]$:

$$U(c', d')(a)_u = \sum_{k=1}^{n} \frac{(d'_k \cdot a_k)_u}{\mathcal{S}K(c')} \Big(\mathcal{S}\mathsf{org}(c')_k - \mathsf{org}(c')_{k,u}\Big) \qquad U(c, d)(a)_u = \sum_{k=1}^{n} \frac{(d_k \cdot a_k)_u}{\mathcal{S}K(c)} \Big(\mathcal{S}\mathsf{org}(c)_k - \mathsf{org}(c)_{k,u}\Big)$$

According to the third and fourth items of Proposition 1.45 and the second item of Proposition 1.43, the following equations hold:

$$(d'_k \cdot a_k)_u = (d_k \cdot a_k)_u \times \begin{cases} 1 & \text{if } k \neq j \\ \dfrac{\mathcal{S}K(d_j)}{\mathcal{S}K(d_j) + \varepsilon} & \text{if } k = j \end{cases}$$

According to Definition 1.44 and Definition 1.40, the following identities hold:

$$\mathsf{org}(c')_{k,u} = \mathsf{org}(c)_{k,u} + \begin{cases} 0 & \text{if } (k,u) \neq (j,v) \\ \varepsilon & \text{if } (k,u) = (j,v) \end{cases} \qquad \mathcal{S}\mathsf{org}(c')_k = \mathcal{S}\mathsf{org}(c)_k + \begin{cases} 0 & \text{if } k \neq j \\ \varepsilon & \text{if } k = j \end{cases} \tag{1.7}$$

We can use the equations and the second item of Proposition 1.45 to show that the difference $U(c', d')(a)_u - U(c, d)(a)_u$ is equal to the following expression for every $u \in [N]$:

$$\frac{(d_j \cdot a_j)_u}{\mathcal{S}K(c)} \frac{\mathcal{S}K(d_j)}{\mathcal{S}K(d_j) + \varepsilon} \Big(\mathcal{S}\mathsf{org}(c)_j + \varepsilon - \mathsf{org}(c')_{j,u}\Big) - \frac{(d_j \cdot a_j)_u}{\mathcal{S}K(c)} \Big(\mathcal{S}\mathsf{org}(c)_j - \mathsf{org}(c)_{j,u}\Big)$$

Let us factorize the previous expression by the inverse of $\mathcal{S}K(d_j) + \varepsilon$ and let us replace the quantity $\mathcal{S}K(d_j)$ with $\mathcal{S}\mathsf{org}(c)_j$ since $\mathcal{S}\delta_j(c, d_j) = \mathcal{S}K(d_j) - \mathcal{S}\mathsf{org}(c)_j = 0$ to obtain the following expression:

$$\frac{1}{\mathcal{S}K(c)} \frac{(d_j \cdot a_j)_u}{\mathcal{S}\mathsf{org}(c)_j + \varepsilon} \Big(\mathcal{S}\mathsf{org}(c)_j(\mathcal{S}\mathsf{org}(c)_j + \varepsilon - \mathsf{org}(c')_{j,u}) - (\mathcal{S}\mathsf{org}(c)_j + \varepsilon)(\mathcal{S}\mathsf{org}(c)_j - \mathsf{org}(c)_{j,u})\Big)$$

We can simplify the right-hand side bracket of the previous expression as follows:

$$\frac{1}{\mathcal{S}K(c)} \frac{(d_j \cdot a_j)_u}{\mathcal{S}\mathsf{org}(c)_j + \varepsilon} \Big(\mathcal{S}\mathsf{org}(c)_j(\mathsf{org}(c)_{j,u} - \mathsf{org}(c')_{j,u}) + \varepsilon \mathsf{org}(c)_{j,u}\Big)$$

Using the left-hand side equations of (1.7) and the fact that $\mathcal{S}\mathsf{org}(c)_j$ is the sum $\sum_{u=1}^{N} \mathsf{org}(c)_{j,u}$, we can show that the previous expression is equal to the one given in the statement. This finishes the proof. □

**Definition 1.47** (Squared algebra operator)**.** Let $c$ and $d_1, \dots, d_n$ be non-empty cells of dimension $N$. For every collection $a = (a_k)_{k\in[n]}$ of vectors $a_k \in \Delta_N(A)^{m_k}$, we will denote by $U^2(c, d)(a)$ the scalar product of the vector $U(c, d)(a)$ with itself.

**Definition 1.48** (Allostasis)**.** Let $(\kappa, c', d')$ be an $(\varepsilon, j, v)$-infinitesimal solution for a homeostasis problem $(c, d, \lambda)$ where $c$ and $c'$ are non-empty cells in $\mathcal{C}^N(n)$ and, for every $k \in [n]$, the $k$-th components $d_k$ and $d'_k$ of $d$ and $d'$ are non-empty cells in $\mathcal{C}^N(m_k)$. For every collection $a = (a_k)_{k\in[n]}$ of vectors $a_k \in \Delta_N(A)^{m_k}$, we will denote:

$$\frac{\partial_{j,v} U^2(c, d)}{\partial(c, d)}(a) := \lim_{\varepsilon \to 0} \frac{U^2(c', d')(a) - U^2(c, d)(a)}{2\varepsilon}.$$

We will call the previous quantity the *allostatic differential* at $(j, v)$.

**Proposition 1.49** (Formula for allostatic differentials)**.** *Let $(\kappa, c', d')$ be an $(\varepsilon, j, v)$-infinitesimal solution for a homeostasis problem $(c, d, \lambda)$ where $c$ and $c'$ are non-empty cells in $\mathcal{C}^N(n)$ and, for every $k \in [n]$, the $k$-th components $d_k$ and $d'_k$ of $d$ and $d'$ are non-empty cells in $\mathcal{C}^N(m_k)$. For every collection $a = (a_k)_{k\in[n]}$ of vectors $a_k \in \Delta_N(A)^{m_k}$, the following equation holds.*

$$\frac{\partial_{j,v}U^2(c,d)}{\partial(c,d)}(a) = \frac{-1}{\mathcal{S}K(c)} \sum_{u=1,\neq v}^{N} \frac{\mathsf{org}(c)_{j,u}}{\mathcal{S}\mathsf{org}(c)_j}\Big((d_j \cdot a_j)_v \times U(c,d)(a)_v - (d_j \cdot a_j)_u \times U(c,d)(a)_u\Big)$$

*Proof.* By Definition 1.48 and Definition 1.47, the allostatic differential of the statement is equal to the limit

$$\frac{\partial_{j,v}U^2(c,d)}{\partial(c,d)}(a) = \lim_{\varepsilon\to 0} \sum_{u=1}^{N} \frac{U(c',d')(a)_u^2 - U(c,d)(a)_u^2}{2\varepsilon}. \tag{1.8}$$

It is easy to check that the following ientity holds:

$$\frac{U(c',d')(a)_u^2 - U(c,d)(a)_u^2}{2\varepsilon} = U(c,d)(a)_u \frac{U(c',d')(a)_u - U(c,d)(a)_u}{\varepsilon} + \frac{(U(c',d')(a)_u - U(c,d)(a)_u)^2}{2\varepsilon}$$

By Proposition 1.36, we can see that the difference $U(c',d')(a)_u - U(c,d)(a)_u$ takes the form of a multiplication $\varepsilon f(\varepsilon)$ where the limit of $f(\varepsilon)$ exists when $\varepsilon \to 0$. In particular, this gives us the following equation:

$$\lim_{\varepsilon\to 0} \frac{U(c',d')(a)_u^2 - U(c,d)(a)_u^2}{2\varepsilon} = U(c,d)(a)_u \lim_{\varepsilon\to 0} f(\varepsilon)$$

It follows from the expression of $f(\varepsilon)$ given in Proposition 1.36 that the following identities hold:

$$\lim_{\varepsilon\to 0} \frac{U(c',d')(a)_u^2 - U(c,d)(a)_u^2}{2\varepsilon} = U(c,d)(a)_u \frac{1}{\mathcal{S}K(c)} \times \begin{cases} (d_j \cdot a_j)_u \dfrac{\mathsf{org}(c)_j}{\mathcal{S}\mathsf{org}(c)_j} & \text{if } u \neq v \\ -(d_j \cdot a_j)_v \displaystyle\sum_{u=1,\neq v}^{N} \frac{\mathsf{org}(c)_{j,u}}{\mathcal{S}\mathsf{org}(c)_j} & \text{if } u = v \end{cases} \tag{1.9}$$

Since equation (1.8) is a limit of a finite sum whose terms admit limits, the allostatic differential of (1.8) is equal to the sum of these limits, all described in (1.9). Computing this sum gives us the equation of the statement. □

1.7. **Cell fission and cell fusion.** The goal of this section is to show that we can regulate the variations of the algebra operator though composition and factorization of cells (see Theorem 1.58).

**Definition 1.50** (Tensor operator)**.** For every positive integer $n$ greater than 2 and collection $\lambda = (\lambda_i)_{i\in[n]}$ of vectors in $\mathbb{R}_+^N$, we will denote the cell $(n, 0, 0, \lambda)$ as $\xi_n^N(\lambda)$. The map $\lambda \mapsto \xi_n^N(\lambda)$ will be called the *$n$-tensor operator.*

**Convention 1.51** (Tensors)**.** Let $n$ be a positive integer and $d = (d_1, \dots, d_n)$ be a $n$-tuple of cells $d_k$ in $\mathcal{C}^N(m_k)$. By Theorem 1.18, for every collection $\lambda = (\lambda_i)_{i\in[n]}$ of vectors in $\mathbb{R}_+^N$, the composition $\xi_n(\lambda) \circ d$, living in $\mathcal{C}^N(m_1 + \cdots + m_n)$, does not depend on $\lambda$ and yields a unique cell whose collection of organelles is the concatenation of the organelles of the cells $d_1, \dots, d_n$, as shown below.

$$\xi_n(\lambda) \circ d = \left(\sum_{k=1}^{n} m_k, \sum_{k=1}^{n} \mathsf{res}(d_k), \sum_{k=1}^{n} \mathsf{cyt}(d_k), \Big(\mathsf{org}(d_1), \mathsf{org}(d_2), \dots, \mathsf{org}(d_n)\Big)\right) \tag{1.10}$$

We will denote this cell as $d_1 \otimes d_2 \otimes \cdots \otimes d_n$ and call it the *$n$-tensor of $d$.*

**Example 1.52** (Using tensors to model fission and fusion of cells)**.** A tensor of cells can be seen as a fusion of the two cells, were the cytosol are added together but the organelles are put next to each other. Alternatively, expressing a given cell as a tensor is a way to model cell division, were the residual and the organelles of the cell are split between the two child cells.

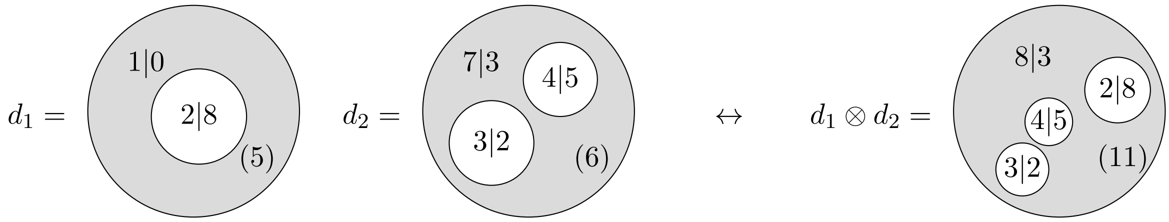

In the remainder of this section, we use tensors to study how cell division and fusion can decrease the values of the algebra operator.

**Example 1.53** (Fission and fusion via compartment formation)**.** The point of defining cell fission and cell fusion by using a tensor operator (Definition 1.50) is that it emphasizes compartment formation, where the created compartment is the tensor operator cell $\xi_n^N(\lambda)$. Of course, as seen in Example 1.52, we cannot explicitly see the formation of this compartment, because this cells is directly composed with other cells. Nevertheless, we can visualize the compartment formation if we make the factorization and composition operations defining (1.10) more explicit (through a two-step operation). More specifically, a fusion can be represented by

- an outer factorization (in the environment) to create the future merged cell;
- an inner composition (in the appeared compartment) to pop-up the inner membranes;

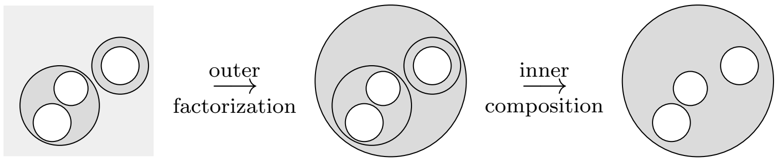

Similarly, a fission can be represented by

- an inner factorization (in the cell) to create the future divided cells;
- an outer composition (in the environment) to pop-up the old membrane;

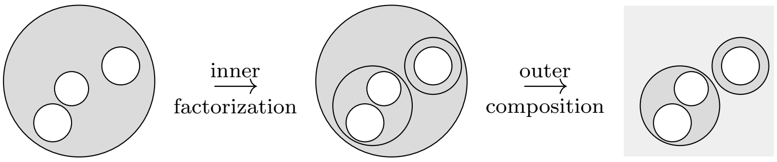

These types of decomposition will be used in section 2.1 to trigger compartment formation while minimizing the values of algebra operator via fission and fusion of cells.

**Proposition 1.54** (Content and tensors)**.** *Let $d_1 \in \mathcal{C}^N(m_1)$ and $d_2 \in \mathcal{C}^N(m_2)$ be two cells. The identity $K(d_1 \otimes d_2) = K(d_1) + K(d_2)$ holds.*

*Proof.* We can deduce the following identities from formula (1.10) given in Convention 1.51.

$$K(d_1 \otimes d_2) = \mathsf{cyt}(d_1 \otimes d_2) + \sum_{i=1}^{m_1+m_2} \mathsf{org}(d_1 \otimes d_2)_i = \mathsf{cy}(d_1) + \mathsf{cyt}(d_2) + \sum_{i=1}^{m_1} \mathsf{org}(d_1)_i + \sum_{i=1}^{m_2} \mathsf{org}(d_2)_i$$

We can then rearrange the terms of the right-hand side to express the sum $K(d_1) + K(d_2)$. $\square$

**Proposition 1.55** (Action and tensors)**.** *Let $d_1 \in \mathcal{C}^N(m_1)$ and $d_2 \in \mathcal{C}^N(m_2)$ be two cells such that their 2-tensor $d_1 \otimes d_2$ is a non-empty cell. For every pair of vectors $a_1 \in \Delta_N(A)^{m_1}$ and $a_2 \in \Delta_N(A)^{m_2}$, the following identity holds for every $u \in [N]$.*

$$((d_1 \otimes d_2) \cdot (a_1 \odot a_2))_u = (d_1 \cdot a_1)_u \frac{\mathcal{S}K(d_1)}{\mathcal{S}K(d_1 \otimes d_2)} + (d_2 \cdot a_2)_u \frac{\mathcal{S}K(d_2)}{\mathcal{S}K(d_1 \otimes d_2)}$$

*Proof.* By Definition 1.30, we have the equation

$$((d_1 \otimes d_2) \cdot (a_1 \odot a_2))_u = \sum_{i=1}^{m_1+m_2} (a_1 \odot a_2)_{i,u} \frac{\mathsf{org}(d_1 \otimes d_2)_{i,u}}{\mathcal{S}K(d_1 \otimes d_2)}$$

Because the vector $a_1 \odot a_2$ is the concatenation of $a_1$ and $a_2$ and the vector $\mathsf{org}(d_1 \otimes d_2)$ is the concatenation of $\mathsf{org}(d_1)$ and $\mathsf{org}(d_2)$, we can rewrite the previous expression as follows.

$$((d_1 \otimes d_2) \cdot (a_1 \odot a_2))_u = \sum_{i=1}^{m_1} (a_1)_{i,u} \frac{\mathsf{org}(d_1)_{i,u}}{\mathcal{S}K(d_1 \otimes d_2)} + \sum_{i=1}^{m_2} (a_2)_{i,u} \frac{\mathsf{org}(d_2)_{i,u}}{\mathcal{S}K(d_1 \otimes d_2)}$$

Factorizing the two sums, on the right-hand side, by $\mathcal{S}K(d_1)/\mathcal{S}K(d_1 \otimes d_2)$ and $\mathcal{S}K(d_2)/\mathcal{S}K(d_1 \otimes d_2)$, we obtain the following expression.

$$((d_1 \otimes d_2) \cdot (a_1 \odot a_2))_u = \frac{\mathcal{S}K(d_1)}{\mathcal{S}K(d_1 \otimes d_2)} \left( \sum_{i=1}^{m_1} (a_1)_{i,u} \frac{\mathsf{org}(d_1)_{i,u}}{\mathcal{S}K(d_1)} \right) + \frac{\mathcal{S}K(d_2)}{\mathcal{S}K(d_1 \otimes d_2)} \left( \sum_{i=1}^{m_2} (a_2)_{i,u} \frac{\mathsf{org}(d_2)_{i,u}}{\mathcal{S}K(d_2)} \right)$$

Finally, by Definition 1.30, this last expression is equivalent to the one given in the statement. □

*Remark* 1.56 (Barycenter). Proposition 1.55 and Proposition 1.54 say that the action of a tensor $(d_1 \otimes d_2)$ on a concatenation $(a_1 \odot a_2)$ is the barycenter of the actions $d_1 \cdot a_1$ and $d_1 \cdot a_2$ where the weights are given by the contents of $d_1$ and $d_2$. More generally, when fitness is assumed and the values of the cytosol are ignored, the action of a cell can be seen as a barycenter of the actions of its organelles. Interestingly, the barycenter of a set of real values can be used as a 'pivot' to divide the space of these real values into two parts: the sets of real values that are below it and those that are above it. In section 2.5, we will use this criterion to model mergings and divisions of cells: specifically, we will look at the barycenter of the organelles of a cell and gather together those organelles that tend to be grouped in the same parts (see Definition 2.30).

**Proposition 1.57** (Fitness and tensor cells)**.** *Let $c \in \mathcal{C}^N(n+1)$ and $c' \in \mathcal{C}^N(n)$ be two cells such that there exits two cells $d_1$ and $d_2$ of dimension $N$ for which the equation $c = c' \circ_n \xi_2(K(d_1), K(d_2))$ holds. In this case, the following identity holds:*

$$K(c) - K(c') = \delta_n(c', d_1 \otimes d_2)$$

*Proof.* First, we apply Proposition 1.24 to the composition $c = c' \circ_n \xi_2(K(d_1), K(d_2))$ to deduce the expression of $K(c')$ in terms of $K(c)$: this gives us the equation $K(c) - K(c') = \mathsf{org}(c')_n - K(d_1) - K(d_2)$. Then, Proposition 1.54 allows us to turn this equation into the equation $K(c) - K(c') = \mathsf{org}(c')_n - K(d_1 \otimes d_2)$. Now, since Definition 1.22 gives us the identity $\delta_n(c', d_2 \otimes d_2) = \mathsf{org}(c')_n - K(d_1 \otimes d_2)$, the statement follows. □

The following theorem assesses by how much the values of the algebra operator are reduced and enhanced when compartment fission or fusion happens in a cell. Since the algebra operator is invariant by symmetry (see Remark 1.35), we will consider only one index configuration for the fusion (or fission) of the organelles, namely only the last two organelles $d_n$ and $d_{n+1}$ will be tensored. See Remark 1.59 for further discussion.

**Theorem 1.58** (Compartment theorem)**.** *Let $c \in \mathcal{C}^N(n+1)$ and $c' \in \mathcal{C}^N(n)$ be two non-empty cells, let $d = (d_1, \ldots, d_{n+1})$ be a $(n+1)$-tuples of non-empty cells where, for every $k \in [n+1]$, the cells $d_k$ is in $\mathcal{C}^N(m_k)$ and let $a = (a_k)_{k \in [n+1]}$ be a collection of vectors $a_k \in \Delta_N(A)^{m_k}$. Let us denote*

$$d' = (d_1, \ldots, d_{n-1}, d_n \otimes d_{n+1}) \qquad \text{and} \qquad a' = (a_1, \ldots, a_{n-1}, a_n \odot a_{n+1}).$$

*If the equation $c = c' \circ_n \xi_2(K(d_n), K(d_{n+1}))$ holds and if we assume that $\mathcal{S}\delta_k(c, d_k) = 0$ for every $k \in [n+1]$ and $\mathcal{S}\delta_k(c', d'_k) = 0$ for every $k \in [n]$, then the following identity holds for every $u \in [N]$.*

$$U(c', d')(a')_u - U(c, d)(a)_u = \Big((d_n \cdot a_n)_u - (d_{n+1} \cdot a_{n+1})_u\Big) \left(\frac{\mathsf{org}(c)_{n,u}}{\mathcal{S}\mathsf{org}(c)_n} - \frac{\mathsf{org}(c)_{n+1,u}}{\mathcal{S}\mathsf{org}(c)_{n+1}}\right) \frac{\mathcal{S}\mathsf{org}(c)_n \mathcal{S}\mathsf{org}(c)_{n+1}}{\mathcal{S}K(c)\mathcal{S}\mathsf{org}(c')_n}$$

*Proof.* According to Proposition 1.36, since $\mathcal{S}\delta_k(c, d_k) = 0$ for every $k \in [n+1]$ and $\mathcal{S}\delta_k(c', d'_k) = 0$ for every $k \in [n]$, we have the following expressions:

$$U(c, d)(a)_u = \sum_{k=1}^{n} \frac{(d_k \cdot a_k)_u}{\mathcal{S}K(c)} \Big(\mathcal{S}\mathsf{org}(c)_k - \mathsf{org}(c)_{k,u}\Big) \quad U(c', d')(a')_u = \sum_{k=1}^{n} \frac{(d'_k \cdot a'_k)_u}{\mathcal{S}K(c')} \Big(\mathcal{S}\mathsf{org}(c')_k - \mathsf{org}(c')_{k,u}\Big)$$

Before evaluating the difference between the previous terms, let us make some observations. First, Proposition 1.57 gives us the equation $K(c) - K(c') = \delta_n(c', d_n \otimes d_{n+1})$. We can couple this equation with the fact that the summed fitness $\mathcal{S}\delta_n(c', d_n \otimes d_{n+1})$ is zero to deduce that the equation $\mathcal{S}K(c) = \mathcal{S}K(c')$ holds.

Second, it follows from Definition 1.19 and the equation $c = c' \circ_n \xi_2(K(d_n), K(d_{n+1}))$ that the identity $\mathsf{org}(c')_k = \mathsf{org}(c)_k$ holds for every $k \in [n-1]$. As a result, the difference $U(c', d')(a')_u - U(c, d)(a)_u$ is equal to the following expression:

$$\begin{aligned} U(c', d')(a')_u - U(c, d)(a)_u &= \frac{(d'_n \cdot a'_n)_u}{\mathcal{S}K(c)} \Big(\mathcal{S}\mathsf{org}(c')_n - \mathsf{org}(c')_{n,u}\Big) \\ -\frac{(d_n \cdot a_n)_u}{\mathcal{S}K(c)} \Big(\mathcal{S}\mathsf{org}(c)_n - \mathsf{org}(c)_{n,u}\Big) &- \frac{(d_{n+1} \cdot a_{n+1})_u}{\mathcal{S}K(c)} \Big(\mathcal{S}\mathsf{org}(c)_{n+1} - \mathsf{org}(c)_{n+1,u}\Big) \end{aligned} \tag{1.11}$$

We now want to factorize expression (1.11). First, according to Proposition 1.54, the identity $\mathcal{S}K(d'_n) = \mathcal{S}K(d_n) + \mathcal{S}K(d_{n+1})$ holds. Since the quantities $\mathcal{S}\delta_n(c, d_n)$, $\mathcal{S}\delta_{n+1}(c, d_{n+1})$ and $\mathcal{S}\delta_n(c', d'_n)$ are all zeros, Definition 1.22 implies that the previous identity is equivalent to the identity

$$\mathcal{S}\mathsf{org}(c')_n = \mathcal{S}\mathsf{org}(c)_n + \mathcal{S}\mathsf{org}(c)_{n+1}. \tag{1.12}$$

We can use equation (1.12) in each of the brackets of (1.11) as well as the identity of Proposition 1.55 in the topmost summand of (1.11), where $d'_n = d_n \otimes d_{n+1}$ and $a'_n = a_n \odot a_{n+1}$, to obtain the following new expression:

$$\begin{aligned} U(c', d')(a')_u - U(c, d)(a)_u &= \frac{(d_n \cdot a_n)_u}{\mathcal{S}K(c)} \left(\frac{\mathcal{S}K(d_n) - \mathcal{S}K(d'_n)}{\mathcal{S}K(d'_n)}\right) \Big(\mathcal{S}\mathsf{org}(c)_n - \mathsf{org}(c)_{n,u}\Big) \\ &+ \frac{(d_{n+1} \cdot a_{n+1})_u}{\mathcal{S}K(c)} \left(\frac{\mathcal{S}K(d_{n+1}) - \mathcal{S}K(d'_n)}{\mathcal{S}K(d'_n)}\right) \Big(\mathcal{S}\mathsf{org}(c)_{n+1} - \mathsf{org}(c)_{n+1,u}\Big) \\ + \frac{(d_n \cdot a_n)_u}{\mathcal{S}K(c)} \frac{\mathcal{S}K(d_n)}{\mathcal{S}K(d'_n)} \Big(\mathcal{S}\mathsf{org}(c)_{n+1} - \mathsf{org}(c)_{n+1,u}\Big) &+ \frac{(d_{n+1} \cdot a_{n+1})_u}{\mathcal{S}K(c)} \frac{\mathcal{S}K(d_{n+1})}{\mathcal{S}K(d'_n)} \Big(\mathcal{S}\mathsf{org}(c)_n - \mathsf{org}(c)_{n,u}\Big) \end{aligned} \tag{1.13}$$

Again, we can use the fact that $\mathcal{S}\delta_{n+1}(c, d_{n+1})$, $\mathcal{S}\delta_n(c, d_n)$ and $\mathcal{S}\delta_n(c', d'_n)$ are zeros to further simplify the dividends of expression (1.13). Specifically, we use the identities $\mathcal{S}\mathsf{org}(c)_{n+1} = \mathcal{S}K(d_{n+1})$, $\mathcal{S}\mathsf{org}(c)_n = \mathcal{S}K(d_n)$ and $\mathcal{S}\mathsf{org}(c')_n = \mathcal{S}K(d'_n)$ as well as equation (1.12) to turn each of the dividend of (1.13) whose expressions use terms of the form $\mathcal{S}K(\_)$ into expressions using terms of the form $\mathcal{S}\mathsf{org}(\_)_\_$, as shown below.

$$\begin{aligned} -\frac{(d_n \cdot a_n)_u}{\mathcal{S}K(c)} \left(\frac{\mathcal{S}\mathsf{org}(c)_{n+1}}{\mathcal{S}\mathsf{org}(c')_n}\right) \Big(\mathcal{S}\mathsf{org}(c)_n - \mathsf{org}(c)_{n,u}\Big) &- \frac{(d_{n+1} \cdot a_{n+1})_u}{\mathcal{S}K(c)} \left(\frac{\mathcal{S}\mathsf{org}(c)_n}{\mathcal{S}\mathsf{org}(c')_n}\right) \Big(\mathcal{S}\mathsf{org}(c)_{n+1} - \mathsf{org}(c)_{n+1,u}\Big) \\ + \frac{(d_n \cdot a_n)_u}{\mathcal{S}K(c)} \frac{\mathcal{S}\mathsf{org}(c)_n}{\mathcal{S}\mathsf{org}(c')_n} \Big(\mathcal{S}\mathsf{org}(c)_{n+1} - \mathsf{org}(c)_{n+1,u}\Big) &+ \frac{(d_{n+1} \cdot a_{n+1})_u}{\mathcal{S}K(c)} \frac{\mathcal{S}\mathsf{org}(c)_{n+1}}{\mathcal{S}\mathsf{org}(c')_n} \Big(\mathcal{S}\mathsf{org}(c)_n - \mathsf{org}(c)_{n,u}\Big) \end{aligned}$$

Observe that the first and third summands as well as the second and fourth summands of the previous expression possess common factors, namely $(d_n \cdot a_n)_u/\mathcal{S}K(c)$ or $(d_{n+1} \cdot a_{n+1})_u/\mathcal{S}K(c)$. We can factorize these summands with respect to their common factors and obtain the following expression after minor simplifications.

$$\frac{(d_n \cdot a_n)_u}{\mathcal{S}K(c)} \frac{\mathcal{S}\mathsf{org}(c)_n \mathcal{S}\mathsf{org}(c)_{n+1}}{\mathcal{S}\mathsf{org}(c')_n} \left( \frac{\mathsf{org}(c)_{n,u}}{\mathcal{S}\mathsf{org}(c)_n} - \frac{\mathsf{org}(c)_{n+1,u}}{\mathcal{S}\mathsf{org}(c)_{n+1}} \right)$$
$$+ \frac{(d_{n+1} \cdot a_{n+1})_u}{\mathcal{S}K(c)} \frac{\mathcal{S}\mathsf{org}(c)_n \mathcal{S}\mathsf{org}(c)_{n+1}}{\mathcal{S}\mathsf{org}(c')_n} \left( \frac{\mathsf{org}(c)_{n+1,u}}{\mathcal{S}\mathsf{org}(c)_{n+1}} - \frac{\mathsf{org}(c)_{n,u}}{\mathcal{S}\mathsf{org}(c)_n} \right)$$

Finally, factorizing the previous expression by the obvious common factor gives the following expression:

$$U(c', d')(a')_u - U(c, d)(a)_u = \Big( (d_n \cdot a_n)_u - (d_{n+1} \cdot a_{n+1})_u \Big) \left( \frac{\mathsf{org}(c)_{n,u}}{\mathcal{S}\mathsf{org}(c)_n} - \frac{\mathsf{org}(c)_{n+1,u}}{\mathcal{S}\mathsf{org}(c)_{n+1}} \right) \frac{\mathcal{S}\mathsf{org}(c)_n \mathcal{S}\mathsf{org}(c)_{n+1}}{\mathcal{S}K(c)\mathcal{S}\mathsf{org}(c')_n}$$

This finishes the proof. □

*Remark* 1.59 (Increasing specialization through compartment flexibility). Theorem 1.58 assesses the variations of the algebra operator when an organelle of a cell is divided or, conversely, when two organelles are merged. More specifically, the formula of Theorem 1.58 implies that the values of the algebra operator are reduced at a given dimension $u$ through compartment division, meaning that the inequality

$$U(c', d')(a')_u - U(c, d)(a)_u > 0$$

holds, if the division of the cell $d'_n = d_n \otimes d_{n+1}$ into the two cells $d_n$ and $d_{n+1}$ is such that either condition (1.14) or condition (1.15) is satisfied:

$$(d_n \cdot a_n)_u > (d_{n+1} \cdot a_{n+1})_u \qquad \frac{\mathsf{org}(c)_{n,u}}{\mathcal{S}\mathsf{org}(c)_n} > \frac{\mathsf{org}(c)_{n+1,u}}{\mathcal{S}\mathsf{org}(c)_{n+1}} \tag{1.14}$$

$$(d_n \cdot a_n)_u < (d_{n+1} \cdot a_{n+1})_u \qquad \frac{\mathsf{org}(c)_{n,u}}{\mathcal{S}\mathsf{org}(c)_n} < \frac{\mathsf{org}(c)_{n+1,u}}{\mathcal{S}\mathsf{org}(c)_{n+1}} \tag{1.15}$$

In other words, the variation between the signals sent by $d_n$ and $d_{n+1}$ to $c$ at dimension $u$ agrees with the variation between the proportions of the $u$-th component of the content between $d_{n+1}$ and $d_n$. Similarly, the values of the algebra operator is reduced at a given dimension $u$ through compartment merging, meaning that the inequality

$$U(c', d')(a')_u - U(c, d)(a)_u < 0$$

holds, if the merging of the two cells $d_n$ and $d_{n+1}$ into the cell $d'_n = d_n \otimes d_{n+1}$ is such that either condition (1.16) or condition (1.17) is satisfied:

$$(d_n \cdot a_n)_u > (d_{n+1} \cdot a_{n+1})_u \qquad \frac{\mathsf{org}(c)_{n,u}}{\mathcal{S}\mathsf{org}(c)_n} < \frac{\mathsf{org}(c)_{n+1,u}}{\mathcal{S}\mathsf{org}(c)_{n+1}} \tag{1.16}$$

$$(d_n \cdot a_n)_u < (d_{n+1} \cdot a_{n+1})_u \qquad \frac{\mathsf{org}(c)_{n,u}}{\mathcal{S}\mathsf{org}(c)_n} > \frac{\mathsf{org}(c)_{n+1,u}}{\mathcal{S}\mathsf{org}(c)_{n+1}} \tag{1.17}$$

In other words, the variation between the signals sent by $d_n$ and $d_{n+1}$ to $c$ at dimension $u$ is opposed to the variation beween the proportions of the $u$-th component of the content between $d_{n+1}$ and $d_n$.

### Part 2. Implementation: the algorithm and why it works

The goal of the present section is to describe, from a mathematical standpoint, the learning algorithm described in the result section of the main text. To facilitate the reader's understanding, we will also use pseudo code to describe the main steps of the learning algorithm. We will refer to the documentation of the code, available at `https://github.com/remytuyeras/intcyt-library`, for further details. Note that this documentation thoroughly explains the content of the present section, from a programming standpoint, through a tutorial. Therefore, we strongly encourage the reader who desires to fully understand the results shown in the main text of this article to have the documentation at hand.

The learning algorithm that we are about to describe is a combination of four main steps, shown in red, in the following diagram.

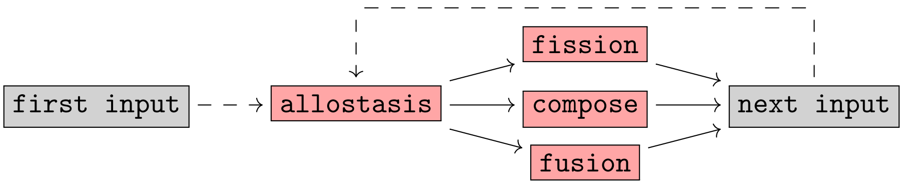

Every input given to the algorithm goes through the step `allostasis` (see diagram above), but depending on the iteration at which the algorithm is, the algorithm can go through either the step `fusion`, the step `fission` or the step `compose`. After any of these steps, the algorithm switches to the next input and restarts the same process from the step `allostasis`, as shown above.

The idea behind using the steps `fusion`, `fission` and `compose` at different iteration lies in Remark 1.53, in which we notice that fusion and fission of cells can be seen as two-step processes. Specifically, for a given *super cell* $\hat{c}$ (see Definition 2.1), the steps `fusion` and `fission` initiates fusion and fission events in $\hat{c}$, while the role of `compose` is to complete these steps. Even though there are actual formal reasons for using such a two-step process (see section 2.4), delaying the completion of fusions and fissions in $\hat{c}$ can also let the algorithm decides whether the chosen fusion or fission events are actually advantageous for $\hat{c}$. When fission and fusion events are not completed through `compose`, new compartments are created in $\hat{c}$. These new compartments can represent a new advantage to organize and classify the information learned. On the other hand, if these compartments turn out to be too disadvantageous for $\hat{c}$, the algorithm will eventually make them disappear.

2.1. **Super cells.** The following definition formalizes the concept of a tree of cells, which was used in the main text of this article, in terms of an object called a 'super cell'. In a few words, the concept of a super cell is an extension of the concept of a cell through a tree-like structure whose leaves and junctions are each associated with a cell.

**Definition 2.1** (Super cells)**.** For every every positive integer $N$ and every non-negative integers $q$ and $n$, we define the set $\mathcal{C}_q^N(n)$ by the recursive formulas given below, on the left (in which $\coprod$ and $\prod$ denote a coproduct and a product of sets, respectively):

$$\begin{cases} \mathcal{C}_0^N(n) = \mathcal{C}^N(n) \\ \mathcal{C}_{q+1}^N(n) = \coprod_{p\geq 1} \coprod_{\substack{(q_1,\dots,q_p):\\ \max(q_i)_{i=1}^p = q}} \coprod_{\substack{(m_1,\dots,m_p):\\ \sum_{i=1}^p m_i = n}} \left( \mathcal{C}_0^N(p) \times \prod_{i=1}^p \mathcal{C}_{q_i}^N(m_i) \right) \end{cases}$$

tree representation of an element in $\mathcal{C}_{q+1}^N(n)$

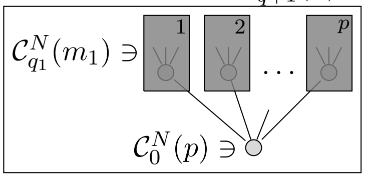

For every pair of non-negative integers $(q, n)$, an element $\hat{c} \in \mathcal{C}_q^N(n)$ is referred to as a *super cell* and said to be *of dimension* $N$, *of height* $q$ and to *possess* $n$ *organelles*. As shown above, on the right, a super cell in $\mathcal{C}_q^N(n)$ can be viewed as a tree-like structure whose leaves and junctions are labeled by cells of dimension $N$. This is further discussed in Convention 2.2.

**Convention 2.2** (Tree structure)**.** Let $\hat{c}$ be a super cell in $\mathcal{C}_q^N(n)$ for which $q \geq 1$. If we unpack the recursive structure of $\hat{c}$ specified in Definition 2.1, then we can express $\hat{c}$ in terms of a collection of tuples of cells and super cells (of lower height) as follows:

$$(2.1) \quad \begin{cases} \hat{c} = (c, (\hat{c}_1, \hat{c}_2, \dots, \hat{c}_p)) & \text{where } c \in \mathcal{C}^N(p) \text{ and } \hat{c}_i \in \mathcal{C}_{q_i}^N(m_i) \\ \hat{c}_{i_1\dots i_s} = (c_{i_1\dots i_s}, (\hat{c}_{i_1\dots i_s 1}, \dots, \hat{c}_{i_1\dots i_s p_{i_1\dots i_s}})) & \text{where } c_{i_1\dots i_s} \in \mathcal{C}^N(p_{i_1\dots i_s}) \text{ and } \hat{c}_{i_1\dots i_s i} \in \mathcal{C}_{q_i}^N(m_{i_1\dots i_s i}) \end{cases}$$

To simplify the previous relations, we can replace any long sequence of indices $i_1 \dots i_s$ with the corresponding collection $I = \{i_1, \dots, i_s\}$. Note that the collection $I$ can be empty, as in the case of the super cell $\hat{c}$ and the cell $c$ given above. Using the notation $I$, we can rewrite the relations of (2.1) more concisely, as shown below, on the left.

$$\text{(2.2)} \qquad \begin{cases} \hat{c} = (c, (\hat{c}_1, \hat{c}_2, \dots, \hat{c}_p)) \\ \hat{c}_I = (c_I, (\hat{c}_{I1}, \hat{c}_{I2}, \dots, \hat{c}_{Ip_I})) \end{cases}$$

$\hat{c}_I =$ 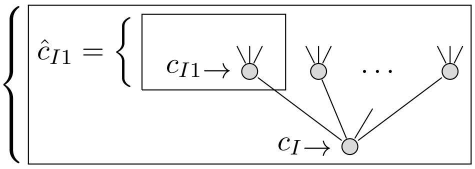

Each of the equations given in (2.2) can be visualized as a branching from a cell $c_I$ to a collection of super cell $\hat{c}_{I1}, \dots, \hat{c}_{Ip_I}$, as shown above, on the right. As a result, the total structure encoded by $\hat{c}$ can be represented as a tree of $q$ levels, where each of the equations of the form $\hat{c}_I = (c_I, (\hat{c}_{I1}, \dots, \hat{c}_{Ip_I}))$ represents a junction in the tree. On the other hand, the super cells $\hat{c}_I$ that cannot be expressed as a tuple of super cells must be of height 0 and correspond to the leaves of the tree structure associated with $\hat{c}$.

Later, given any super cell $\hat{c}$, we will refer to any associated super cell $\hat{c}_I$ of height greater than 1 as a *junction of* $\hat{c}$ and any associated super cell $\hat{c}_I$ of height 0 as a *leaf of* $\hat{c}$.

**Definition 2.3** (Indexing collections for junctions and leaves)**.** For every $\hat{c} \in \mathcal{C}_q^N(n)$, we denote by $\mathsf{Junc}(\hat{c})$ the set of collections indexing the junctions of $\hat{c}$ and by $\mathsf{Leaf}(\hat{c})$ the set of collections indexing the leaves of $\hat{c}$. More formally, these sets are defined by the following specifications:

$$\begin{cases} \mathsf{Junc}(\hat{c}) = \{I \mid \hat{c}_I \text{ is a super cell of } \hat{c} \text{ whose height is not } 0\} \\ \mathsf{Leaf}(\hat{c}) = \{I \mid \hat{c}_I \text{ is a super cell of } \hat{c} \text{ whose height is } 0\}. \end{cases}$$

We will denote by $\mathsf{Index}(\hat{c})$ the union $\mathsf{Junc}(\hat{c}) \cup \mathsf{Leaf}(\hat{c})$.

**Proposition 2.4** (Number of leaves)**.** *For every $\hat{c} \in \mathcal{C}_q^N(n)$, the set $\mathsf{Leaf}(\hat{c})$ possesses $n$ elements.*

*Proof.* Follows from an induction using the recursive structure given in Definition 2.1. □

**Definition 2.5** (Linear order)**.** For every super cell $\hat{c}$ in $\mathcal{C}_q^N(n)$, we define a linear order on $\mathsf{Index}(\hat{c})$ by taking the lexicographical order on integers, namely:

$$I = \{i_1, \dots, i_s\} \prec J = \{j_1, \dots, j_t\} \Leftrightarrow \exists u < \mathsf{min}(s,t) \text{ such that } i_k = j_k \text{ for every } k \in [u] \text{ and } i_{u+1} < j_{u+1}$$

**Convention 2.6** (Indexing shifts)**.** For every $\hat{c} \in \mathcal{C}_q^N(n)$ and every indexing collection $I \in \mathsf{Leaf}(\hat{c})$, we denote as $m(I)$ the number of organelles of $\hat{c}_I$. If $I \in \mathsf{Junc}(\hat{c})$, then we denote the number of organelles of $\hat{c}_I$ as $p(I)$. Finally, if we denote by $I_1 \prec \cdots \prec I_i$ the ordered sequence of elements of $\mathsf{Leaf}(\hat{c})$ that are less than $I$, then we denote by $L(I)$ the sum $m(I_1) + \cdots + m(I_i)$.

**Definition 2.7** (Organelles for a super cell)**.** For every $\hat{c} \in \mathcal{C}_q^N(n)$, any vector of the form $\mathsf{org}(\hat{c}_I)_i$, where $I \in \mathsf{Leaf}(\hat{c})$ and $i \in [m(I)]$, will be called an *organelle* of $\hat{c}$. If we denote by $I_1 \prec \cdots \prec I_n$ the ordered sequences of the elements of $\mathsf{Leaf}(\hat{c})$, then we denote by $\mathsf{org}(\hat{c})$ the concatenation $\mathsf{org}(\hat{c}_{I_1}) \odot \cdots \odot \mathsf{org}(\hat{c}_{I_n})$, which comprises all the organelles of $\hat{c}$.

**Proposition 2.8** (Retrieval)**.** *For every $\hat{c} \in \mathcal{C}_q^N(n)$, let $(v_1, v_2, \dots, v_n)$ denote the tuple $\mathsf{org}(\hat{c})$ of the organelles of $\hat{c}$. For every $I \in \mathsf{Leaf}(\hat{c})$, the vector $(v_{L(I)+1}, v_{L(I)+1}, \dots, v_{L(I)+m(I)})$ corresponds to $\mathsf{org}(\hat{c}_I)$.*

*Proof.* Follows from the construction of $\mathsf{org}(\hat{c})$ given in Definition 2.7 and the definition of $L(I)$ given in Convention 2.6. □

**Definition 2.9** (Pre-actions and actions)**.** Let $\hat{c}$ be a super cell in $\mathcal{C}_q^N(n)$ and let $a$ be a vector in $\Delta_N(A)^n$. For every $I \in \mathsf{Index}(\hat{c})$, we define the *pre-action* $\mathsf{preact}(\hat{c}|a)_I$ and the *action* $\mathsf{act}(\hat{c}|a)_I$ of the super cell $\hat{c}$ on the vector $a$ at the indexing collection $I$ by the following recursive formulas:

- ▷ if $I \in \mathsf{Index}(\hat{c})$, then $\mathsf{act}(\hat{c}|a)_I := c_I \cdot \mathsf{preact}(\hat{c}|a)_I$;
- ▷ if $I \in \mathsf{Leaf}(\hat{c})$, then $\mathsf{preact}(\hat{c}|a)_I = (a_{L(I)+1}, \dots, a_{L(I)+m(I)})$;

▷ if $I \in \mathsf{Junc}(\hat{c})$, then $\mathsf{preact}(\hat{c}|a)_I = \mathsf{act}(\hat{c}|a)_{I1} \odot \cdots \odot \mathsf{act}(\hat{c}|a)_{Ip(I)}$;

The leaves of any super cell are special super cells because their height is 0, meaning that they are proper cells. As a result, these super cells do not have enough structure to allow us to interpret fission and fusion events, because these events require two layers of cells, hence a super cell of height 1 (see section 1.6 and section 1.7). A solution to this problem will be to interpret each leaf of $\hat{c}$ as a super cell of height 1 by replacing their organelles with identity cells (see Definition 1.9).

**Definition 2.10** (Direct children and basal pruning)**.** For every $\hat{c} \in \mathcal{C}_q^N(n)$, we define the *direct children* of $\hat{c}$ as the following tuple of cells:

$$d(\hat{c}) = \begin{cases} (\mathsf{id}_{\mathsf{org}(c)_1}, \dots, \mathsf{id}_{\mathsf{org}(c)_n}) & \text{If } \hat{c} \text{ is a leaf, meaning that } \hat{c} = c \\ (c_1, \dots, c_p) & \text{If } \hat{c} \text{ is a junction, meaning that } \hat{c} = (c, (\hat{c}_1, \dots, \hat{c}_p)) \end{cases}$$

We define the *basal pruning* of $\hat{c}$ as the super cell $\mathsf{base}(\hat{c}) := (c, d(\hat{c}))$ where $c$ is the obvious root cell associated with $\hat{c}$.

**Definition 2.11** (Homeostatic state)**.** A super cell $\hat{c}$ in $\mathcal{C}_q^N(n)$ will be said to be *in a homeostatic state* (or to have *reached homeostasis*) if for every $I \in \mathsf{Junc}(\hat{c})$ such that $\hat{c}_I = (c_I, (\hat{c}_{I1}, \dots, \hat{c}_{Ip(I)}))$, each cell of the form $c_{Ii}$ (associated with the super cell $\hat{c}_{Ii}$) fits the $i$-th organelle of the cell $c_I$ (see Convention 1.23). In other words, the equation $\delta_i(c_I, c_{Ii}) = 0$ holds for every $I \in \mathsf{Junc}(\hat{c})$ and every index $i \in [p(I)]$.

Motivated by Convention 2.10, we diversify the pre-action of a super cell (Definition 2.9) in two ways, namely as an 'expected signal' and a 'specialized signal' (see Definition 2.12). In the spirit of the discussion following Definition 1.30, we think of the former as an expected value of the input, while we think of the latter as a measurement of how well the input matches each of the organelles of the super cell.

**Definition 2.12** (Expected and specialized signal)**.** Let $\hat{c}$ be a super cell in $\mathcal{C}_q^N(n)$ and let $a$ be a vector in $\Delta_N(A)$. For every $I \in \mathsf{Index}(\hat{c})$, we define

▷ the *expected signal* $\mathsf{esgn}(a|\hat{c})_I$ of the vector $a$ through $\hat{c}$ at the indexing collection $I$ as the pre-action $\mathsf{preact}(a'|\hat{c})_I$ on the concatenation $a' = a \odot a \odot \cdots \odot a$ of $n$ copies of $a$.

▷ the *specialized signal* $\mathsf{ssgn}(a|\hat{c})_I$ of the vector $a$ through $\hat{c}$ at the indexing collection $I$ as the pre-action $\mathsf{preact}(a'|\hat{c})_I$ on the concatenation $a' = (\mathsf{id}_{\mathsf{org}(\hat{c})_1} \cdot a) \odot (\mathsf{id}_{\mathsf{org}(\hat{c})_2} \cdot a) \odot \cdots \odot (\mathsf{id}_{\mathsf{org}(\hat{c})_n} \cdot a)$.

2.2. **Spontaneous reactions and homeostatic state.** Before describing the three steps of our algorithm, we must describe an intermediate step that is essential to the three steps. This intermediate step consists in making sure that a given super cell has reached homeostasis (Definition 2.11). This is done by going through successive steps of normalization called 'cleanings'. These cleaning procedures essentially implements the 'overall reaction' described in Remark 1.32.

**Definition 2.13** (Cleaning)**.** For every cell $c \in \mathcal{C}^N(n)$, we call the *cleaning of* $c$ the cell $\mathsf{clean}(c)$ of $\mathcal{C}^N(n)$ determined by the following equations:

1) *Energy storage:* $\mathsf{res}(\mathsf{clean}(c)) = \mathsf{res}(c) + \mathcal{S}\mathsf{cyt}(c)$
2) *Disorder cleaning:* $\mathsf{cyt}(\mathsf{clean}(c)) = (0, 0, \dots, 0)$
3) *Compartmentalization:* $\mathsf{org}(\mathsf{clean}(c)) = \mathsf{org}(c)$

If we interpret the cleaning of a cell as the result of an 'overall chemical reaction', then we can interpret the three items of Definition 2.13 as follows: item 1 states that the cytosolic content is converted into energetic content, which is to be used in later similar chemical reactions; item 2 states that the cytosolic content is cleaned from its components, which have been turned into energetic content; and item 3 states that the organelles of the cells are untouched, mainly because they are separated from the cytosol by membranes.

By Definition 1.2, the cleaning of a cell is defined if, and only if its residual is non-negative. This means that a cell admits a cleaning if, and only if, it is well-defined according to the following definition.

**Definition 2.14** (Well-definedness)**.** A cell $c \in \mathcal{C}^N(n)$ will be said to be *well-defined* if the inequality $\mathsf{res}(c) + \mathcal{S}\mathsf{cyt}(c) \geq 0$ is satisfied.

The previous notion of well-definedness can be extended to super cells as follows.

**Definition 2.15** (Well-definedness)**.** A super cell $c \in \mathcal{C}_q^N(n)$ will be said to be *well-defined* if it satisfies the following inductive conditions:

1) If $\hat{c}$ is of height 0, then $\hat{c}$ is well-defined as a cell (Definition 2.14);
2) If $\hat{c} = (c, (\hat{c}_1, \dots, \hat{c}_p))$ is of height $q > 0$, then $c$ is well-defined as a cell and each super cell $\hat{c}_i$ is a well-defined super cell of height $q - 1$ by induction.

We also define the cleaning of a super cell by induction. In our implementation, the cleaning of a super cell is implemented through the method `spontaneous_reaction` (see the documentation).

**Definition 2.16** (Cleaning)**.** For every well-defined super cell $\hat{c} \in \mathcal{C}_q^N(n)$, we define the *cleaning of* $\hat{c}$ as the super cell $\hat{e}$ of $\mathcal{C}_q^N(n)$ that possesses the same indexing structure as $\hat{c}$ and is determined by the following equations:

1) If $I \in \mathsf{Leaf}(\hat{c})$, then $\hat{e}_I = e_I$ is equal to the cell $\mathsf{clean}(c_I)$ (in the sense of Definition 2.13);
2) If $I \in \mathsf{Junc}(\hat{c})$, then $\hat{e}_I$ is equal to the super cell $(e_I, (\hat{e}_{I1}, \dots, \hat{e}_{Ip(I)}))$ where each super cell $\hat{e}_{Ii}$ is defined by induction and where the cell $e_I$ is determined by the following equations:
   2.1) *Energy storage:* $\mathsf{res}(e_I) = \mathsf{res}(c_I) + \mathcal{S}\mathsf{cyt}(c_I)$
   2.2) *Disorder cleaning:* $\mathsf{cyt}(e_I) = (0, 0, \dots, 0)$
   2.3) *Homeostatic state:* $\mathsf{org}(e_I) = (K(e_{I1}), \dots, K(e_{Ip(I)}))$

From now on, the cleaning $\hat{e}$ of $\hat{c}$ will be denoted as $\mathsf{clean}(\hat{c})$.

*Remark* 2.17 (Hierarchical learning and data integration). In our implementation, the ability to clean a super cell is essential to making the algorithm to learn and classify data. More specifically, cleaning allows a cell to communicate their learning progress to their parent cell in the tree structure. To understand why this is the case, we will use the following schematic, which represents various update states through which a cell and its parent go during learning.

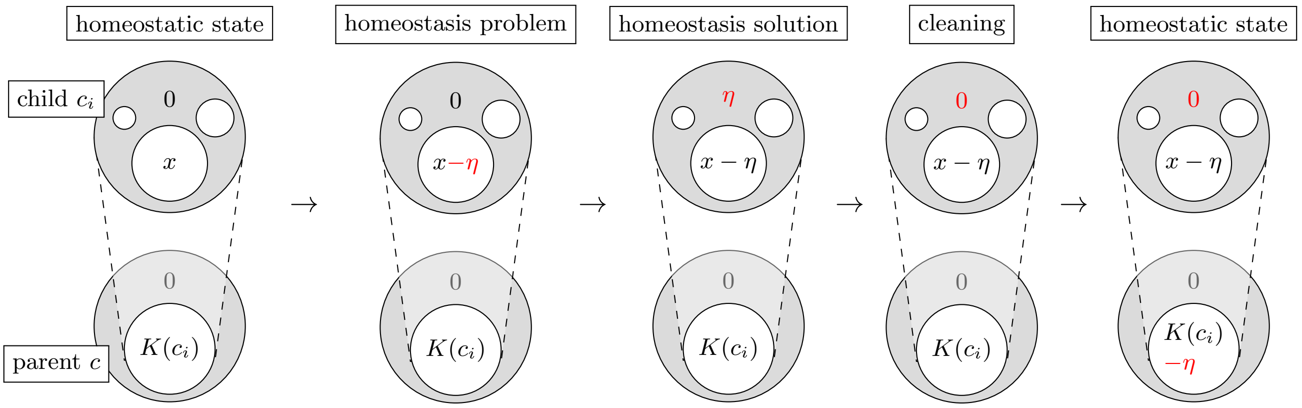

The first stage, from the left, represents a cell and its parent whose associated ambient super cell (not represented) is in a homeostatic state (Definition 2.11). Because the cytosolic content of the child cell $c_i$ is 0, the corresponding organelle $\mathsf{org}(c)_i$ of the parent cell $c$ is equal to the content $K(c_i)$ of the child cell (Definition 1.22). The second stage, from the left, shows a situation in which the organelle $x$ of the child cell $c_i$ undergoes a small change $-\eta$ (this kind of modification will be used in section 2.3 to make a super cell learn). After such a modification, the super cell might no longer be in a homeostatic state, so we re-establish fitness between the cells by solving the homeostasis problems induced by these modifications (see Definition 1.40). In the third stage, we can see that solving the homeostasis problems changes the values of the cytosolic contents. While these added values ensure fitness, they prevent the organelle $\mathsf{org}(c)_i$ of the parent cell $c$ to report what has been learned in its child cell $c_i$. We therefore proceed to a cleaning of the super cell, which we illustrate in the last two stages: the second last stage illustrates the cleaning of the child cell $c_i$ (Definition 2.13) and the last stage ensures that the child cell $c_i$ fits the organelle $\mathsf{org}(c)_i$. As a result, the organelle $\mathsf{org}(c)_i$ accounts for the information learned by its associated child cell $c_i$.

2.3. **Allostasis.** The first step of our algorithm, called `Allostasis`, is strongly inspired from gradient descent techniques. The goal of this step is to *minimize* the values of the algebra operators (Definition 1.34) associated with each of the junctions of a super cell. To do so, we use homeostasis problems as back propagation mechanisms (section 1.6). The main difference with the usual gradient descent is that we use a different concept of differentials, namely the concept of allostatic differential given in Definition 1.48. As with usual gradient descent, we will use a "gamma parameter". In our case, the gamma parameter is a function of the following form:

$$\gamma : \mathcal{C}^N(m) \times \Delta_N(A)^p \times [p] \times [N] \to \mathbb{R}$$

In Definition 2.18, given below, we use a parameter gamma whose third argument is the pre-action associated with an expected signal (Definition 2.12). In the spirit of gradient descent, the output of the gamma parameter should be viewed as an intensity, while the allostatic differential associated with it (see Definition 2.12) should be viewed as a directionality (through its positive and negative values).

**Definition 2.18** (Allostatic matrix)**.** Let $\hat{c}$ be a super cell in $\mathcal{C}_q^N(n)$ and let $a$ be a vector in $\Delta_N(A)^n$. For every $I \in \mathsf{Index}(\hat{c})$ such that $\mathsf{base}(\hat{c}_I) = (c_I, (d_1, d_2, \dots, d_p))$, we denote by $\mathsf{All}(\hat{c}|a)_I$ the $p \times N$-matrix whose $(j, v)$-coefficient is equal to the following multiplication:

$$\gamma(d_j, \mathsf{esgn}(\hat{c}|a')_I, j, v) \times \left(\frac{\partial_{j,v} U^2(\mathsf{base}(\hat{c}_I))}{\partial(\mathsf{base}(\hat{c}_I))}\right). \tag{2.3}$$

The following definition formalizes the tuning procedure illustrated in Remark 2.17.

**Definition 2.19** (Allostatic tuning)**.** Let $\hat{c}$ be a super cell in $\mathcal{C}_q^N(n)$, let $a$ be a vector in $\Delta_N(A)^n$ and let $I$ be in $\mathsf{Index}(\hat{c})$. Suppose that we are given a homeostasis problem $\lambda$ for $\mathsf{base}(\hat{c}_I) = (c_I, d(\hat{c}_I))$ such that:

$$\lambda_{i,u} \text{ is equal to either } \mathsf{org}(c_I)_{i,u} \text{ or } \mathsf{org}(c_I)_{i,u} - \mathsf{All}(\hat{c}|a)_{I,i,u}. \tag{2.4}$$

If we let $(\kappa, e_I, d_I)$ denote the homeostasis solution of $\lambda$ (see the equations below), then we define the *allostatic tuning* $\mathsf{tune}(\hat{c}|\lambda)_I$ of $\hat{c}$ *relative to* $(\lambda, I)$ as the super cell $(e_I, d_I)$ if $I \in \mathsf{Index}(\hat{c})$ and as the cell $e_I$ if $I \in \mathsf{Leaf}(\hat{c})$.

$$\begin{cases} e_I &= \mathsf{left}\big(c_I \circ d(\hat{c}_I)\big)\big(\mathsf{res}(d(\hat{c}_I)), \kappa, \lambda\big) \\ d_I &= \mathsf{right}\big(c_I \circ d(\hat{c}_I)\big)\big(m(\hat{c}_I), \mathsf{res}(d(\hat{c}_I)), \kappa\big) \end{cases}$$

*Remark* 2.20 (Homeostatic state). In our implementation, we compute formula (2.3) by using the formula of Proposition 1.49. In order to use Proposition 1.49, we need to make sure that the super cell $\hat{c}$ is in a homeostatic state (in our implementation, this is done through the methods `spontaneous_reaction` associated with the classes `Cell` and `SuperCell`). Such a condition is necessary because Proposition 1.49 relies on formula (1.4), which requires the super cell $\hat{c}$ to be in a homeostatic state. Note that this homeostatic state is always maintained through allostatic tuning due to the fitness properties satisfied by homeostasis solutions (see Definition 1.40).

*Remark* 2.21 (Existence of a tuning). While Theorem 1.42 states that homeostasis solutions always exist and are unique, it is important to note that a homeostasis problem $\lambda$ may be ill-defined when some of the coefficients $\lambda_{i,u}$ are negative (which is forbidden by Definition 2.19). As a result, the allostatic tuning of a super cell (Definition 2.19) may not always exist. This conflict can be solved algorithmically by setting the corresponding coefficients $\mathsf{All}(\hat{c}|a)_{I,i,u}$ to 0 so that $\lambda_{i,u}$ stays non-negative (see formula (2.4)). In our implementation, we use this kind of practice in order to find a well-defined allostatic tuning (in the sense of Definition 2.15); for more detail, see the methods `compute_variable`, `homeostasis` and `allostasis` associated with the class `SuperCell`.

In our implementation, we use gamma parameters of the form shown below, where $c \in \mathcal{C}^N(n)$ and $a \in \Delta_N(A)^n$, where the function $\mathsf{agr}$ is defined in Definition 2.22 (in our implementation, the function $\mathsf{agr}$ is implemented through the method `agreement`) and where the function $\mathsf{magr}$ is defined in Definition 2.23.

$$\gamma(c,a,j,u) = \begin{cases} 0 & \text{if learning should not occur according to specific criteria;} \\ 10^E \times \left(\dfrac{\mathsf{agr}(c,j,a_j)}{\mathsf{magr}(c,a)}\right)^F & \text{otherwise.} \end{cases}$$

The idea behind the previous formula is that the quantity $\mathsf{agr}(c,j,a_j)$ measures a correlation between $c$ and $a_j$ (a value between 0 and 1), the quantity $\mathsf{magr}(c,a)$ makes sure that the correlation measured by $\mathsf{agr}(c,j,a_j)$ is not biased towards a certain type of input $a$, the exponent $F$ attenuates the influence of the normalized correlation $\mathsf{agr}(c,j,a_j)/\mathsf{magr}(c,a)$ when its values are low and the power $10^E$ makes sure that the values $\gamma(c,j,a)$ land within a specific numerical interval. Note that the scenarios in which $\gamma(c,a,j,u)$ is set to 0 are relatively easy to formalize in our case due to the interpretability of the information learned by the cells (see the tutorial of the documentation).

**Definition 2.22** (Agreements)**.** Below, for every vector $x = (x_1,\dots,x_N)$, we let $\|x\|$ denote the norm $\sqrt{\sum x_u^2}$ of $x$. Let $c$ be a cell in $\mathcal{C}^N(n)$ and $a$ be a vector in $\Delta_N(A)$. For every integer $j \in [N]$, we define *the $j$-th agreement* $\mathsf{agr}(c,j,a)$ *of $c$ with $a$* as the following normalized scalar product

$$\mathsf{agr}(c,j,a) := \sum_{u=1}^{N} \left( \frac{\mathsf{org}(c)_{j,u}}{\|\mathsf{org}(c)_j\|} \times \frac{a_u}{\|a\|} \right)$$

The agreement of $c$ with $a$ can be interpreted as the cosine of the angular distance between $\mathsf{org}(c)_j$ and $a$

**Definition 2.23** (Max-agreements)**.** Let $c$ be a cell in $\mathcal{C}^N(n)$ and $a$ be a vector in $\Delta_N(A)^n$. We define *the max-agreement* $\mathsf{magr}(c,a)$ *of $c$ with $a$* as the maximum of the values $\mathsf{agr}(c,j,a_j)$ for every $j \in [N]$.

$$\mathsf{magr}(c,a) := \max\{\mathsf{agr}(c,j,a_j) \mid j \in [N]\}$$

We finally give the pseudo code for the step `Allostasis`. In our implementation, this code is implemented through the methods `compute_variable`, `homeostasis`, and `allostasis` (see the documentation).

| `Allostasis` |
|---|
| **Input:** $(\hat{c}, a, \gamma)$ where $\hat{c}$ is in a homeostatic state |
| **For** every indexing collection $I \in \mathsf{Index}(\hat{c})$ taken in the decreasing order **do:** |
| **Compute** $\mathsf{All}(\hat{c}\|a)_I$ |
| **Set** $\lambda_{I,i,u} := \mathsf{org}(c_I)_{i,u} - \mathsf{All}(\hat{c}\|a)_{I,i,u}$ |
| **For** every indexing collection $I \in \mathsf{Index}(\hat{c})$ taken in the decreasing order **do:** |
| **If** $I \in \mathsf{Leaf}(\hat{c})$ **do:** |
| **Update** $(c_I, (c_{I1}, \dots, c_{Ip(I)})) \leftarrow \mathsf{tune}(\hat{c}\|\lambda_I)_I$ |
| **If** $I \in \mathsf{Leaf}(\hat{c})$ **do:** |
| **Update** $c_I \leftarrow \mathsf{tune}(\hat{c}\|\lambda_I)_I$ |
| **While** the cells of $\hat{c}_I$ are not well-defined (Definition 2.15) **do:** |
| **Update** the conflicting $\lambda_{I,i,u}$ to $\mathsf{org}(c_I)_{i,u}$ (see Remark 2.21) |
| **If** $I \in \mathsf{Leaf}(\hat{c})$ **do:** |
| **Update** $(c_I, (c_{I1}, \dots, c_{Ip(I)})) \leftarrow \mathsf{tune}(\hat{c}\|\lambda_I)_I$ |
| **If** $I \in \mathsf{Leaf}(\hat{c})$ **do:** |
| **Update** $c_I \leftarrow \mathsf{tune}(\hat{c}\|\lambda_I)_I$ |
| **Return** $\hat{c}$ |

2.4. **Composition of cells.** The present section describes the step `Compose` of our algorithm. This step is used to complete the other two steps `Fusion` and `Fission` described in section 2.5. The main reason for this two-step completion is to avoid composing both leaf and non-leaf super cells within the same cell, because this operation would not return a super cell (see Definition 2.1).

**Definition 2.24** (Binary configurations)**.** A *binary configuration* for a super cell $\hat{c}$ in $\mathcal{C}_q^N(n)$ is a function $\beta : \mathsf{Index}(\hat{c}) \to \{0,1\}$. We organize the binary values $\beta_I$ of $\beta$ as a tree structure of the same form as the underlying tree structure encoding $\hat{c}$, namely we define by induction:

1) $\hat{\beta}_I := \beta_I$ for every $I \in \mathsf{Leaf}(\hat{c})$.
2) $\hat{\beta}_I := (\beta_I, (\hat{\beta}_{I1}, \dots, \hat{\beta}_{Ip(I)}))$ for every $I \in \mathsf{Junc}(\hat{c})$;

Each number 1 and 0 of the binary configuration specifies whether the outer cell of the super cell is to be composed within its outer environment or not. For instance, if a cell $c_I$ is associated with a number $\beta_I = 1$, then this means that we intend to compose the cell $c_I$ within its parent cell (see Definition 2.25).

As mentioned at the beginning of this section, the main reason for completing fusion and fission events in a separate composition step is that fusion and fission events may require to handle simultaneous compositions of leaf and non-leaf cells within the same cells. These mixed compositions can conflict with the definition of a super cell, in which the children of a parent cell are either all proper super cells (all of height greater than 0) or all leaf super cells (all of height equal to 0). We handle these potential conflicts through Definition 2.25, in which all-leaf-compositions are treated in item 2.1 and mixed-compositions are treated in item 2.2.

**Definition 2.25** (configured composition)**.** Let $\beta$ be a binary configuration for a super cell $\hat{c}$ in $\mathcal{C}_q^N(n)$. We define the *configured composition of $\hat{c}$ relative to $\beta$* as the super cell $\mathsf{comp}(\hat{c}|\beta)$ defined recursively as follows:

1) If $\hat{c}$ is a leaf (*i.e.* a cell), then $\mathsf{comp}(\hat{c}|\hat{\beta}) = \hat{c}$;
2) If $\hat{c} = (c, (\hat{c}_1, \dots, \hat{c}_p))$ is a junction and:
   2.1) all super cells $\hat{c}_1, \dots, \hat{c}_p$ are leaves for which $\beta_1 = \cdots = \beta_p = 1$, then we define:
   $$\mathsf{comp}(\hat{c}|\hat{\beta}) = c \circ (\hat{c}_1, \dots, \hat{c}_p)$$
   2.2) otherwise, we proceed as follows: for every $i \in [p]$, we define the numerical variable
   $$\nu_i = \begin{cases} 0 & \text{if } \mathsf{comp}(\hat{c}_i|\hat{\beta}_i) \text{ is a cell } c_i' \\ 1 & \text{if } \mathsf{comp}(\hat{c}_i|\hat{\beta}_i) \text{ is a super cell } (c_i', (\hat{c}_{i1}', \dots, \hat{c}_{ip(i)}')) \end{cases}$$
   and we define
   $$\begin{cases} \mathsf{comp}(\hat{c}|\hat{\beta}) = (c \circ \mathsf{val}(\hat{c}|\hat{\beta}), (\mathsf{coval}(c|\hat{\beta})_1, \dots, \mathsf{coval}(c|\hat{\beta})_p)) \\ \mathsf{val}(\hat{c}|\hat{\beta}) = (\mathsf{val}(\hat{c}|\hat{\beta})_1, \dots, \mathsf{val}(\hat{c}|\hat{\beta})_p) \end{cases}$$
   where we denote:
   $$\mathsf{val}(\hat{c}|\hat{\beta})_i = \begin{cases} c_i' & \text{if } \beta_i = 1 \\ \mathsf{id}_{K(c_i')} & \text{if } \beta_i = 0 \end{cases} \qquad \mathsf{coval}(c|\hat{\beta})_i = \begin{cases} \hat{c}_{i1}', \dots, \hat{c}_{ip(i)}' & \text{if } \beta_i = 1 \text{ and } \nu_i = 1 \\ (\mathrm{id}_{\mathsf{org}(c_i')_1}, \dots, \mathrm{id}_{\mathsf{org}(c_i')_p}) & \text{if } \beta_i = 1 \text{ and } \nu_i = 0 \\ \mathsf{comp}(\hat{c}_i|\hat{\beta}_i) & \text{if } \beta_i = 0 \end{cases}$$

The pseudo-code of the step `compose` is straightforward and mainly relies on Definition 2.25. In our implementation, this code is implemented through the method `compose` of the class `SuperCell` (see the documentation).

| Compose |
| --- |
| **Input:** $(\hat{c}, \hat{\beta})$ |
| **Return** $\mathsf{comp}(\hat{c}|\hat{\beta})$ |

2.5. **Merging and dividing cells.** The present section describes the steps `fusion` and `fission`. Our implementation of these steps mainly relies on the statement of Theorem 1.58 and the interpretation given in Remark 1.59. To give an intuition, we illustrate an example of the step `fission` below, on the left, and an example of the step `fusion` on the right. As can be seen, both steps consists in adding new compartments (in darker gray) to the original cell. The integers 0 and 1 next to each compartment represent the values of an associated binary configuration (Definition 2.24), which we construct according to the intuition described in Example 1.53.

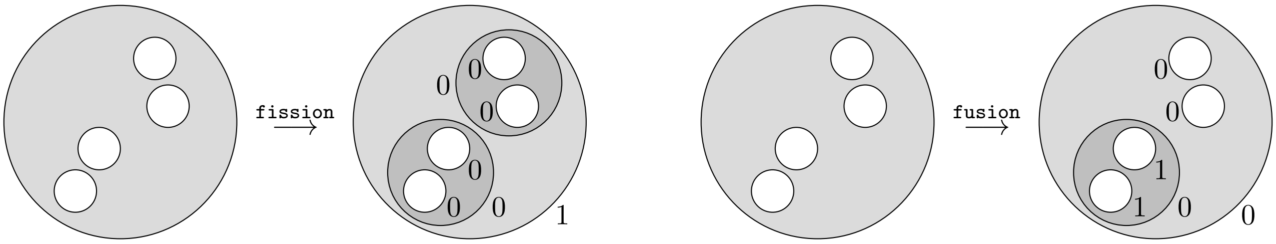

For `fusion`, we separate the organelles of the cell in two groups, distributed within two new compartments, and we compose the outer membrane of the super cell within the environment – when is possible. For `fission`, we also separate the organelles of the cell in two groups, but we only create a compartment for one of the groups, and we compose the super cells associated with the organelles of that group within the new compartment.

The following definition simulates a cell division at the root of a super cell.

**Definition 2.26** (Division)**.** Let $\hat{c}$ be a super cell in $\mathcal{C}_q^N(n)$ in a homeostatic state such that $q > 0$ and $\beta$ be a binary configuration for $\hat{c}$. Supposing that $\hat{c} = (c, (\hat{c}_1, \dots, \hat{c}_p))$, we let $S = \{i_1 < i_2 < \dots < i_k\}$ be a subset of $[p]$. We define the *basal division* of $(\hat{c}, \hat{\beta})$ *relative to* $S$ as a pair $(\hat{e}, \hat{\tau})$ where $\hat{\tau} = (\tau, (\hat{\tau}_1, \hat{\tau}_2))$ is a binary configuration for the super cell $\hat{e} = (e, (\hat{e}_1, \hat{e}_2))$ defined as follows:

1) the cell $e$ is in $\mathcal{C}^N(p-k+1)$ and is defined as the tensor cell

$$\xi_2(\sum_{j \in S} \mathsf{org}(c)_j, \sum_{j \notin S} \mathsf{org}(c)_j)$$

while we have the identity $\tau = 1$;

2) the super cells $\hat{e}_1$ and $\hat{e}_2$ are determined by the following relations

$$\hat{e}_i = \begin{cases} (e_1, (\hat{c}_j)_{j \in S}) & \text{if } i = 1 \\ (e_2, (\hat{c}_j)_{j \notin S}) & \text{if } i = 2 \end{cases}$$

where the cells $e_1$ and $e_2$ are determined by the equations

$$\begin{array}{lll} \mathsf{res}(e_1) = \mathsf{res}(c)/2, & \mathsf{cyt}(e_1) = 0, & \mathsf{org}(e_1) = (\mathsf{org}(c)_j)_{j \in S}, \\ \mathsf{res}(e_2) = \mathsf{res}(c)/2, & \mathsf{cyt}(e_2) = 0, & \mathsf{org}(e_2) = (\mathsf{org}(c)_j)_{j \notin S}, \end{array}$$

while we have the identity $\hat{\tau}_i = \hat{\beta}_i$ for every $i \in [p-k+1]$.

The following definition simulates a cell merging at the root of a super cell.

**Definition 2.27** (Merging)**.** Let $\hat{c}$ be a super cell in $\mathcal{C}_q^N(n)$ in a homeostatic state such that $q > 0$ and $\beta$ be a binary configuration for $\hat{c}$. Supposing that $\hat{c} = (c, (\hat{c}_1, \dots, \hat{c}_p))$, we let $S = \{i_1 < i_2 < \dots < i_k\}$ be a subset of $[p]$. We define the *basal merging* of $(\hat{c}, \hat{\beta})$ *relative to* $S$ as a pair $(\hat{e}, \hat{\tau})$ where $\hat{\tau} = (\tau, (\hat{\tau}_1, \dots, \hat{\tau}_{p-k+1}))$ is a binary configuration for the super cell $\hat{e} = (e, (\hat{e}_1, \dots, \hat{e}_{p-k+1}))$ defined as follows:

1) the cell $e$ is in $\mathcal{C}^N(p-k+1)$ and is determined by the relations $\mathsf{res}(e) = \mathsf{res}(c)$, $\mathsf{cyt}(e) = 0$ and

$$\mathsf{org}(e)_i = \begin{cases} \mathsf{org}(c)_i & \text{if } i < i_1 \\ \sum_{j \in S} \mathsf{org}(c)_j & \text{if } i = i_1 \\ \mathsf{org}(e)_{i-q} & \text{if } i_q < i < i_{q+1} \end{cases}$$

while we have the identity $\tau = \beta$;

2) the super cells $\hat{e}_1, \dots, \hat{e}_p$ are determined by the following relations

$$\hat{e}_i = \begin{cases} \hat{c}_i & \text{if } i < i_1 \\ (\xi_k(K(c_{i_1}), K(c_{i_2}), \dots, K(c_{i_k})), \hat{c}_{i_1}, \dots, \hat{c}_{i_k}) & \text{if } i = i_1 \\ \hat{c}_{i-q} & \text{if } i_q < i < i_{q+1} \end{cases}$$

while we have the following identities:

$$\hat{\tau}_i = \begin{cases} \hat{\beta}_i & \text{if } i < i_1 \\ (0, (\hat{\tau}_{i_1}, \dots, \hat{\tau}_{i_k})) & \text{if } i = i_1 \\ \hat{\beta}_{i-q} & \text{if } i_q < i < i_{q+1} \end{cases} \qquad \text{where} \qquad \hat{\tau}_{i_j} = (1, (\hat{\beta}_{i_j 1}, \dots, \hat{\beta}_{i_j p(i_j)})).$$

Thus, for both steps `fusion` and `fission`, we divide the organelles in two groups with respect to a subset $S$ of the set of the organelles indices. In the rest of this section, our goal is to explain how we compute the set $S$ for our implementation. We proceed in two steps:

1) we first compute a weighted adjacency matrix on the set of organelle indices (Definition 2.30);
2) and we take $S$ to be one of the maximal connected component of the graph associated with the matrix (see Definition 2.33).

To do so, we use the idea of barycenters discussed in Remark 1.56, which we formalize through Definition 2.29. We start with a notation for tensors of child cells in a super cells.

**Convention 2.28** (Tensor of child cells)**.** Let $\hat{c}$ be a super cell in $\mathcal{C}_q^N(n)$. For every $I \in \mathsf{Junc}(\hat{c})$ such that $\hat{c}_I = (c_I, (\hat{c}_{I1}, \dots, \hat{c}_{Ip}))$ and every subset $S = \{i_1, i_2, \dots, i_k\}$ of $[p]$, we will denote by $c_{I,S}$ the tensor $c_{Ii_1} \otimes c_{Ii_2} \cdots \otimes c_{Ii_k}$.

Definition 2.29 introduces weighted sums of pre-actions, which we use to retrieve the type of actions computed in Proposition 1.55.

**Definition 2.29** (Tensored pre-action)**.** Let $\hat{c}$ be a super cell in $\mathcal{C}_q^N(n)$ and $a$ be a vector in $\Delta_N(A)$. For every $I \in \mathsf{Junc}(\hat{c})$ such that $\hat{c}_I = (c_I, (\hat{c}_{I1}, \dots, \hat{c}_{Ip}))$, let us denote by $(b_1, b_2, \dots, b_p)$ the specialized signal $\mathsf{ssgn}(\hat{c}|a)_I$. We define the *tensor of* $\mathsf{ssgn}(\hat{c}|a)_I$ as the following weighted sum (or barycenter).

$$\mathsf{center}(\hat{c}|a)_I := \sum_{i \in S} b_i \times \frac{\mathcal{S}K(c_{Ii})}{\mathcal{S}K(c_{I,[p]})}$$

The following definition implements the criterion discussed in Remark 1.56, namely the construction of an adjacency matrix that indicates which of the organelles are on the same side of the barycenter $\mathsf{center}(\hat{c}|a)_I$.

**Definition 2.30** (Adjacency matrices)**.** Let $\hat{c}$ be a super cell in $\mathcal{C}_q^N(n)$ and $a$ be a vector in $\Delta_N(A)$. For every $I \in \mathsf{Junc}(\hat{c})$ such that $\hat{c}_I = (c_I, (\hat{c}_{I1}, \dots, \hat{c}_{Ip}))$, let us denote by $(b_1, b_2, \dots, b_p)$ the specialized signal $\mathsf{ssgn}(\hat{c}|a)_I$. For every postive real number $\nu$, we define two $(p \times p)$-matrices $\pi_\nu^-(I)$ and $\pi_\nu^+(I)$ as follows (where $\#$ denotes the cardinality operation):

$$\pi_\nu^-(I,a)_{i,j} := \#\{u \in [N] \mid b_{i,u} \text{ and } b_{j,u} \text{ are less than } \nu \times \mathsf{center}(\hat{c}|a)_{I,u}\}$$
$$\pi_\nu^+(I,a)_{i,j} := \#\{u \in [N] \mid b_{i,u} \text{ and } b_{j,u} \text{ are greater than } \nu \times \mathsf{center}(\hat{c}|a)_{I,u}\}$$

From now on, the two weighted adjacency matrices $\pi_\nu^-(I,a)$ and $\pi_\nu^+(I,a)$ will be viewed as their associated weighted graphs. Below, we use the matrices $\pi_\nu^-(I,a)$ and $\pi_\nu^+(I,a)$ for cell fusion and cell fission, respectively.

**Definition 2.31** (Connected components)**.** Let $\hat{c}$ be a super cell in $\mathcal{C}_q^N(n)$ and $a$ be a vector in $\Delta_N(A)$. For every $I \in \mathsf{Junc}(\hat{c})$, every $\varepsilon \in \{+,-\}$ and non-negative integer $\omega$, we denote by $\mathsf{clust}_\nu^\varepsilon(I, a|\omega)$ the set of subsets $S \subseteq [p]$ such that

1) the maximum weight $m = \mathsf{max}\{\pi_\nu^\varepsilon(I,a)_{i,j} \mid i,j\}$ in the graph $\pi_\nu^\varepsilon(I,a)$ is greater than or equal to $\omega$;
2) there is a path linking the edges in $S$ through vertices of maximum weight $m$ in the graph $\pi_\nu^\varepsilon(I,a)$;
3) $S$ has a maximal cardinal among all subsets satisfying item 1.

While the elements of $\mathsf{clust}_\nu^\varepsilon(I, a|\omega)$ allow us to select groups of organelles with respect to one of the inequalities discussed in Remark 1.59, namely the 'signal comparison', it does not tell us anything about the second type of comparison, which compares the proportions in the organelles of the cell. We take care of this second step through the scoring system defined in Definition 2.32. In this definition, we replace the organelle vectors with content vectors, assuming that the underlying super cell in a homeostatic state (see

Definition 2.11). Also, for every subset $S \subseteq [p]$, we will write $[p]\backslash S$ to denote the complement of $S$ within $[p]$.

**Definition 2.32** (Quality)**.** Let $\hat{c}$ be a super cell in $\mathcal{C}_q^N(n)$ and $a$ be a vector in $\Delta_N(A)$. For every $I \in \mathsf{Junc}(\hat{c})$, every $\varepsilon \in \{+,-\}$ and non-negative integer $\omega$, we define the *quality* of a set $S \in \mathsf{clust}_\nu^\varepsilon(I, a|\omega)$ as the sum given below, on the left.

$$\mathsf{qual}(S) := \sum_{u \in \mathbf{B}(I,S)} \frac{K(c_{I,S})_u}{\mathcal{S}K(c_{I,S})} \times \frac{\mathcal{S}K(c_{I,[p]\backslash S})}{K(c_{I,[p]\backslash S})_u} \quad \text{where} \quad \mathbf{B}(I,S) = \left\{ u \in [N] \;\middle|\; \frac{K(c_{I,S})_u}{\mathcal{S}K(c_{I,S})} > \frac{K(c_{I,[p]\backslash S})_u}{\mathcal{S}K(c_{I,[p]\backslash S})} \right\}.$$

**Definition 2.33** (Best-compartments)**.** Let $\hat{c}$ be a super cell in $\mathcal{C}_q^N(n)$ and $a$ be a vector in $\Delta_N(A)$. For every $I \in \mathsf{Junc}(\hat{c})$, every $\varepsilon \in \{+,-\}$ and non-negative integer $\omega$, we say that a set $S \in \mathsf{clust}_\nu^\varepsilon(I, a|\omega)$ is a *best choice* if the quality $\mathsf{qual}(S)$ is maximal among all the qualities of the elements of $\mathsf{clust}_\nu^\varepsilon(I, a|\omega)$.

We now give the pseudo-code of the steps `fusion` and `fission`. In our implementation, this code is implemented through the methods `best_compartment` and `proposed_clustering` associated with the class `Cell` and through the methods `merge_base`, `fusion`, `divide_base`, and `fission` associated with the class `SuperCell`.

`Fusion` (if $\varepsilon = -$) / `Fission` (if $\varepsilon = +$)

**Input:** $(\hat{c}, \beta, a, \varepsilon, \nu, \omega)$ where $\beta$ is a binary configuration for $\hat{c}$
**Update** $\hat{c} \leftarrow \mathsf{clean}(\hat{c})$
**For** every indexing collection $I \in \mathsf{Index}(\hat{c})$ taken in the decreasing order **do:**
  **Compute** $\mathsf{clust}_\nu^\varepsilon(I, a|\omega)$
  **Select** a best compartment $S$ in $\mathsf{clust}_\nu^\varepsilon(I, a|\omega)$
  **If** $\varepsilon = +$ **do:**
    **Set** $(\hat{e}, \hat{\tau})$ as the basal division of $(\hat{c}_I, \hat{\beta}_I)$ relative to $S$ (Definition 2.26)
  **If** $\varepsilon = -$ **do:**
    **Set** $(\hat{e}, \hat{\tau})$ as the basal merging of $(\hat{c}_I, \hat{\beta}_I)$ relative to $S$ (Definition 2.27)
  **Update** $\hat{c}_I \leftarrow \hat{e}$
  **Update** $\hat{\beta}_I \leftarrow \hat{\tau}$
**Return** $(\hat{e}, \hat{\beta})$

#### 3. Link with operad theory

While we never really tried to formalize our model in categorical terms, it may have occurred to the learned reader that our model takes place within the world of 'colored operad' [28]. Specifically, for a given positive integer $N$, the collection of sets $\mathcal{C}^N(1)$, $\mathcal{C}^N(2)$, ... can be organized into a colored operad $\mathcal{D}^N$ on $\mathbb{R}_+^N$. This colored operad is given by a collection of sets $\mathcal{D}^N(x_1, \dots, x_n; y)$ containing all those cells $c = (n, C, x, (x_i)_i)$ for which $y = K(c)$ is in $\mathbb{R}_+^N$. These sets are equipped with identities $\mathsf{id}_a \in \mathcal{D}^N(a; a)$ (see Proposition 1.10) and satisfy the usual composition axioms for operads (see Theorem 1.17). From this point of view, we have used the operad $\mathcal{D}^N$ every time we composed a cell that fitted the organelle of another cell.

Ideally, we could use the operadic point of view to further simplify our model. For instance, the operad $\mathcal{D}^N$ can better be described by the colored operad $\mathcal{D}_n^N : (\mathbb{R}_+^N)^n \times \mathbb{R}_+^N \to \mathbf{Set}$ mapping every tuple $(x_1, \dots, x_n, y)$ in $(\mathbb{R}_+^N)^n \times \mathbb{R}_+^N$ to the set $\mathbb{R}_+$ such that its identities are the 0 elements of the sets $\mathcal{D}_1^N(a, a) = \mathbb{R}_+$ and its compositions are given by the sums of real numbers of the following type:

$$\mathcal{D}_n^N(x_1, \dots, x_n, y) \times \prod_{i=1}^{n} \mathcal{D}_{m_i}^N(z_{i,1}, \dots, z_{i,m_i}, x_i) \to \mathcal{D}_{m_1+\dots+m_n}^N(z_{1,1}, \dots, z_{n,m_n}, y).$$

Interestingly, the tensor operations defined in Convention 1.51 can also be defined as sum operations of the following form:

$$\otimes : \mathcal{D}_n^N(x_1, \dots, x_n, y) \times \mathcal{D}_m^N(x'_1, \dots, x'_m, y') \to \mathcal{D}_{n+m}^N(x_1, \dots, x_n, x'_1, \dots, x'_m, y + y').$$

Such a reformulation could be useful to better understand the mechanisms pertaining to the evolution of super cells during a learning phase. Specifically, with this formalism, super cells can be defined in terms of a *free* colored operad on $\mathcal{D}^N$. This kind of formalization could be useful to clarify the relationships that exist between cell fission, cell fusion, and homeostasis. In particular, this clarification could be used to investigate potential improvements of our current implementation.
